# Unscored variants leave many comparisons of variant effect predictors undecided: evidence from ClinVar, published benchmarks and genomic language models

**DOI:** 10.64898/2026.11.23.768553

**Authors:** Varunkumar Asediya, Udayaraja GK, Haja N. Kadarmideen, Chandra S. Pareek

## Abstract

**Background:** Variant effect predictors leave different variants unscored, so reported accuracies often rest on different variants, and which predictor is better can depend on variants never scored.

**Results:** Manski’s sharp bounds contain each predictor’s accuracy and area under the receiver operating characteristic curve (AUROC) whatever its unscored variants would score. A comparison is decided when the two bounds are disjoint, which cannot happen when the two shares scored (of variants or, for AUROC, pathogenic–benign pairs) sum to one or less. For 49 dbNSFP predictors on 328,328 ClinVar substitutions, 745 of 1,176 comparisons are undecided inside declared scopes and 1,093 on the whole panel. In ten benchmarks from one PubMed query, 1,045 of 2,842 stated orderings (36.8%) are undecided. Yet most orderings between well-known predictors are decided (177 of 287), among them AlphaMissense above REVEL in the sixth Critical Assessment of Genome Interpretation. AlphaMissense’s canonical-transcript release scores no missense substitution in secondary-findings genes CALM1, CALM2 or WT1, and missense variants that REVEL’s and AlphaMissense’s releases leave unscored are pathogenic less often than the panel’s (likelihood ratios 0.18 and 0.25, describing each release). Across nine species, the genomic language model Evo 2 does not outperform gene annotation (species-mean AUROC 0.943 against 0.958); its 8,192-bp margin is resolved over Ensembl’s conservation track (+0.108) but not over the 241-mammal phyloP (+0.055).

**Conclusions:** Benchmarks should report each predictor’s reach (share of variants scored) per label class and which orderings the bounds decide; the deposited reach matrices and glmtrust package allow these checks without rescoring.

## Background

Published accuracy comparisons guide which variant effect predictor a diagnostic laboratory runs and which is trusted in species without clinical genetics. Yet predictors leave different variants unscored. A conservation score is undefined where its alignment fails, a missense predictor such as Sorting Intolerant From Tolerant (SIFT) [1] or Polymorphism Phenotyping v2 (PolyPhen-2) [2] scores only missense substitutions, and a splicing model’s precomputed files skip genes outside its transcript set [3]. Two predictors are therefore compared either on the variants both score or each on its own variants, and a printed accuracy need not say which.

Benchmarks settle this by convention: they compare each pair on shared variants [4,5], fill gaps with each predictor’s least deleterious score to check a ranking [5], report only the models able to score a mutation type [6], or score each predictor on the variants it covers [7]; dbNSFP’s maintainers benchmark 49 of the scores they distribute on nine annual ClinVar releases and print each score’s coverage beside its accuracy, with a minimum-coverage filter [8]. Earlier predictors’ incomplete coverage motivated a genome-wide protein language model, Evolutionary Scale Modeling 1b (ESM1b) [9]. The Critical Assessment of Functional Annotation (CAFA) charges a function predictor, in its full mode, for every target it leaves unanswered [10,11], and diagnostic-accuracy standards ask how non-evaluable results were handled [12,13]. We found no benchmarking guideline for variant effect predictors that does either. Benchmarks guard against circularity [14] by excluding training variants or scoring against deep mutational scans [15], yet variant type [16] and gene identity [17] alone account for much of the accuracy they report.

In the clinic, the Clinical Genome Resource (ClinGen) has calibrated score intervals of several predictors as computational evidence of stated strength [18,19]. One calibration could not recommend thresholds for the evolutionary model of variant effect (EVE) [20]: its predictions left about half of the benign calibration variants unscored, in genes with a skewed ratio of pathogenic to benign variants [19].

Outside genomics, the accuracy an abstaining classifier would have had on every item is identified when its abstentions are stochastic and the evaluation data are independent of its training, but not when it abstains deterministically [21], as a predictor with a declared scope does. Grading by accuracy alone rewards a language model for guessing over abstaining [22], and when only the items on which two models disagree are labelled, partial identification can certify, or fail to certify, their F1 ranking [23]. The tools are established. With nothing assumed about missing outcomes, a mean lies in Manski’s sharp bound [24], whose form for an average treatment effect (each unit seen under one treatment only) always contains zero. A contaminated share, under Horowitz and Manski’s contaminated-sampling model [25], gives an interval, and for untied scores the arithmetic is that of the interval-based area under the curve [26].

Alignment-free genomic language models [27,28] score any variant they can centre in a sequence window, so against a conventional predictor nearly every missing score belongs to the other side. They learn constraint, as models trained on explicit alignments do [29,30], so conservation is the baseline that matters [30,31]. The authors of the Genomic Pretrained Network with Species Tree and Alignment Representations (GPN-Star) note that standard genomic language models still fall short of classical evolutionary ones [30]. Jiang and colleagues report in a preprint that Evo 2 [31] separates the catalogued alleles of Online Mendelian Inheritance in Animals (OMIA) [32,33] from population variants in eight non-human species [34] (Supplementary Note S1).

We show when a comparison is decided. For any metric averaged over items, and for the area under the receiver operating characteristic curve (AUROC), two predictors are ordered, whatever their unscored variants would score, exactly when their sharp bounds are disjoint, which cannot happen when the two shares scored (of variants or, for AUROC, of pathogenic–benign pairs) sum to one or less. We add a proof that both ends of the AUROC bound are attained, the breakdown point in closed form and the rule’s application to the orderings that benchmarks print; the bounds themselves and their comparison are established [24]. For 49 predictors on ClinVar [35], as the database for nonsynonymous single-nucleotide polymorphisms’ functional predictions (dbNSFP) distributes them, the bounds leave 745 of 1,176 comparisons undecided inside declared scopes and 1,093 on the whole panel. In ten published benchmarks they leave 36.8% of stated orderings undecided, most of them in two benchmarks’ sorted figure legends, while most orderings between well-known predictors are decided. Among ClinVar variants of uncertain significance, we count those a calibrated tool leaves unscored separately from those it scores as indeterminate; the first calibration study tabulates missing predictions per tool [18]. Across nine species, Evo 2 does not outperform annotation, and its lead over conservation holds only against Ensembl’s track and only at the window mean its developers released. The same rule applies to genomic language model benchmarks and a language-model factuality benchmark. We release the reach matrices (reach being the share of variants scored, which other literatures call coverage), a nine-species atlas and the glmtrust package, which computes the bounds [36].

## Data Description

We assembled the panels below and deposited them as per-variant tables that record which variants each predictor scored and which it left unscored, so a comparison can be recomputed with unscored variants charged rather than dropped. Which variants a predictor leaves unscored is itself the measurement; unscored variants are given scores only in the conventions set against the bound (Supplementary Note S37). Additional file 1 maps each variant panel and published benchmark from source to figure (Supplementary Figure S10). Additional file 2 holds the code and the glmtrust package; Additional file 3 holds per-variant tables and the recompute scripts.

Additional file 4 holds 328,328 ClinVar substitutions, in 15,606 genes, that dbNSFP annotates as nonsynonymous or splice-site, with 49 predictors’ reach indicators but not their scores. Additional file 5 holds the 1,484,836 ClinVar single-nucleotide variants classified pathogenic or benign, before the review-status filter. It carries review status, consequence class, phyloP100way [37] and phastCons100way [38] values, and the reach, but not the scores, of Combined Annotation Dependent Depletion (CADD) [39], the Rare Exome Variant Ensemble Learner (REVEL) [40] and AlphaMissense [41]. The ClinVar release behind it also holds 2,232,064 single-nucleotide variants of uncertain significance at one review star or better (Supplementary Note S38).

The cross-species atlas (Additional file 3) holds 11,130 single-nucleotide variants in nine species, eight of them domestic animals. Its 2,240 positives are catalogued disease- or trait-associated alleles, from OMIA in eight species and ClinVar in human. Its 8,890 negatives are population variants matched on trinucleotide context at 10:1 (9.92:1 in cat) and, in human, ClinVar benign or likely benign records at 1:1. Each row carries Evo 2’s scores at two readouts and, where defined, the Genomic Evolutionary Rate Profiling (GERP) [42,43] value and the Zoonomia 241-mammal phyloP [44,45]. A consequence-matched arm adds 9,119 negatives for 2,237 positives, matched on consequence class as well as context: 5,002 share their positive’s class and 4,117 are synonymous. It carries the 241-mammal phyloP, GERP and, in chicken and human, GPN-Star scores, and no Evo 2 score (Supplementary Note S44).

The splicing panel holds 3,912 variants with splice-disruption labels from six datasets (Supplementary Note S2). Additional file 6 holds the 500,000-variant simulated background for the splice predictors. Three further panels are a BRCA1 saturation-editing panel (3,893 substitutions), 5,000 fine-mapped porcine cis-expression quantitative trait loci (cis-eQTLs) with 15,000 controls, and 7,479 single-nucleotide ClinGen Evidence Repository classifications [46]. The last Analyses subsection reads published values only: ten benchmarks returned by one PubMed query (Supplementary Note S24), the score files released with GPN-Star [30] and TraitGym [47], and the results table of SimpleQA, a language-model factuality benchmark [48].

## Analyses

### The accounting, and when an ordering is decided

A predictor leaves unscored, or declines, the variants it returns no score for, whether outside its declared scope, unaligned or missing from its files, and a printed accuracy is silent about them (Fig. 1a). The covered AUROC, A_cov_, is its AUROC on the pathogenic–benign pairs in which it scores both variants; its pair coverage, ρ, is their share of all pairs. With nothing assumed about declined variants, the AUROC it would attain had it scored every variant lies in the sharp bound [ρA_cov_, ρA_cov_ + 1 − ρ], whose ends count every unranked pair wrong or right (Fig. 1b; Methods). The must-answer AUROC, 0.5 + (A_cov_ − 0.5)ρ, counts each as a tie; it is the bound’s midpoint, and A_cov_ minus it the must-answer penalty.

**Figure 1.**
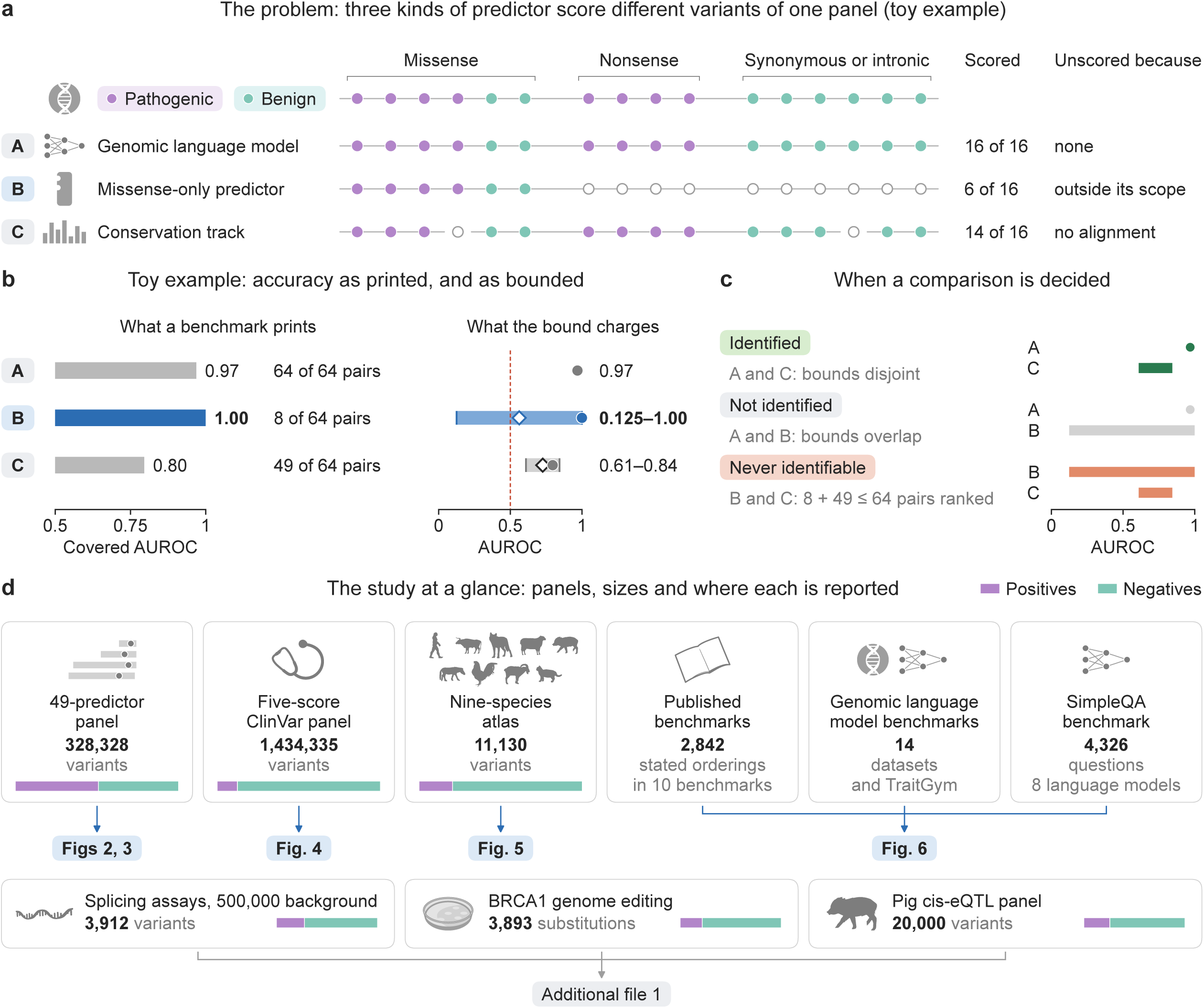
The problem, the accounting and the study at a glance. (a) A toy panel of eight pathogenic and eight benign variants, grouped by consequence, and three kinds of predictor: A scores every variant, B only the four pathogenic and two benign variants inside its declared scope, C all but two that lack an alignment (open, unscored). (b) Left, what a benchmark prints: each predictor’s area under the receiver operating characteristic curve (AUROC) on the pairs it ranks, its covered AUROC (bars start at chance, 0.5); blue, predictor B. Right, the sharp bound on the AUROC it would attain had it scored every variant (band), with the covered AUROC (circle) and the must-answer AUROC (diamond), which counts each unranked pair as a tie; dashed, chance. (c) The three verdicts on the toy’s three pairs: identified, the two bounds are disjoint; not identified, they overlap; never identifiable, the two pair coverages sum to one or less. (d) The study’s panels: size, share of positives (bar) and the figure that reports each. The toy’s values illustrate the accounting and are not results about any kind of predictor.

Alt text: A toy panel of sixteen variants and three predictors: A scores every variant, B only six inside its declared scope, C all but two. B’s covered AUROC of 1.00 on 8 of 64 pairs fits any whole-panel AUROC from 0.125 to 1.00. Comparisons are identified, not identified or never identifiable; cards map the study’s panels to the figures. A comparison is identified, or decided, when the two predictors’ bounds do not overlap (Fig. 1c). The bound and the comparison of two bounds are Manski’s [24]. The rule is exact for metrics that average a bounded contribution over items, such as accuracy, and for AUROC, but not for metrics that are not averages, such as the F-score (Supplementary Note S36). Where the rule is exact, it cannot be sharpened: one predictor’s missing scores do not restrict another’s, so without a known relation between the two, no analysis of the variants both score decides a comparison whose bounds overlap. Coverage alone can preclude ordering: two predictors whose coverages ρ sum to one or less cannot be ordered, whatever their covered accuracies (never identifiable; Fig. 1c). Fig. 1d maps the panels analysed.

### 49 predictors as dbNSFP distributes them: reach and bounds

As dbNSFP 5.3.1a distributes them, most of 49 published predictors rank under half of a ClinVar panel’s pathogenic–benign pairs, with bounds that contain chance. The panel holds that release’s rank scores for 328,328 ClinVar substitutions (166,149 pathogenic, 162,179 benign; Fig. 2; key, Fig. 2a). A decline in this panel pools three causes: the variant lies outside the predictor’s scope, is absent from the tool’s own release, or was scored by the tool only on a transcript that dbNSFP’s gene model has since retired [49]. Reach and verdicts therefore belong to this source and move with it (Supplementary Note S5).

**Figure 2.**
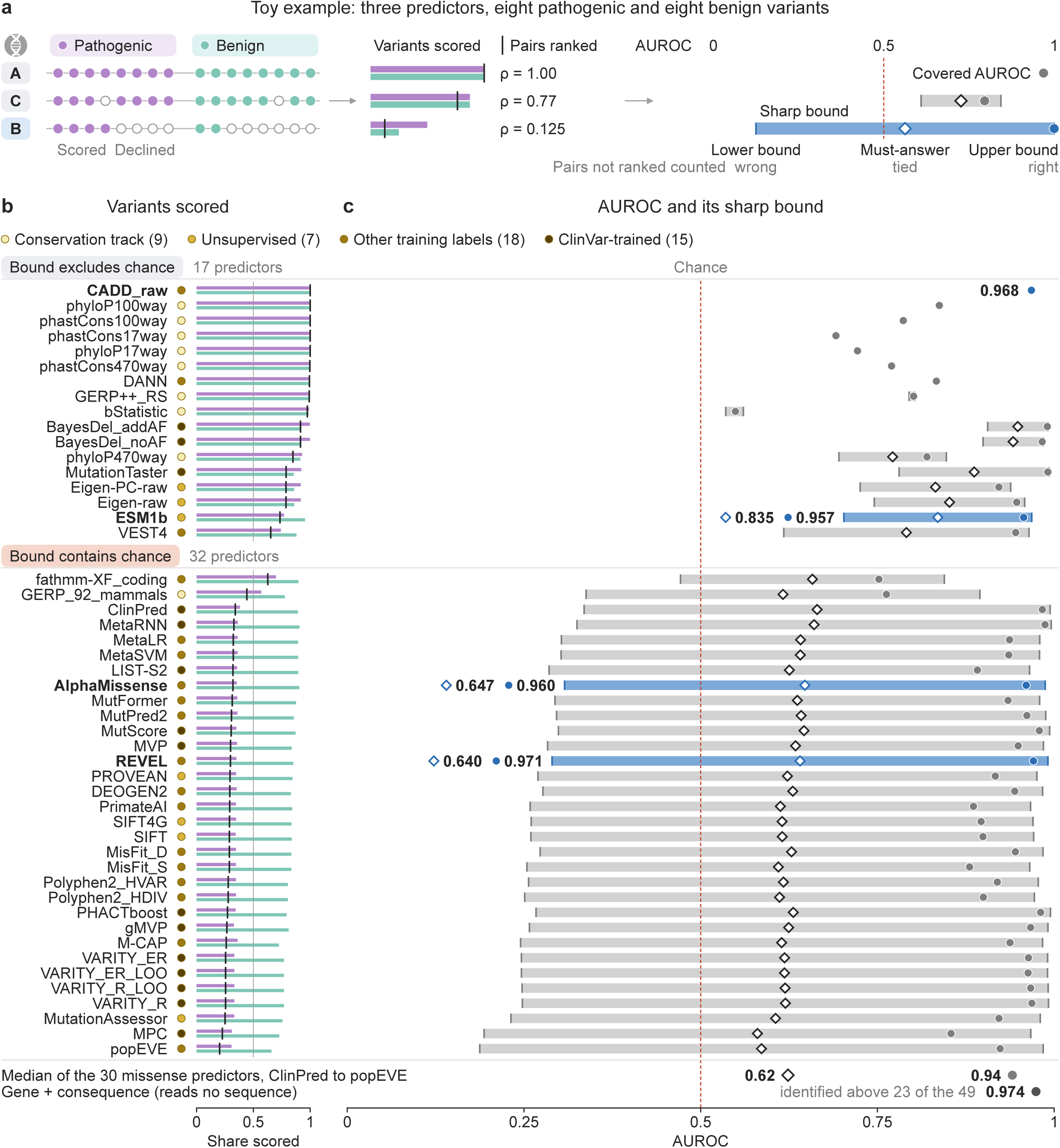
Reach and sharp bounds of 49 published predictors on 328,328 ClinVar substitutions in dbNSFP 5.3.1a. (a) Key, on the toy of Figure 1a: its three predictors score or decline eight pathogenic and eight benign variants (open, unscored). The shares scored (bars) multiply to the share of pairs ranked, ρ (tick); counting the unranked pairs as wrong, tied or right gives the lower bound, the must-answer area under the receiver operating characteristic curve (AUROC; diamond) and the upper bound; circle, covered AUROC; blue, predictor B as in Figure 1. (b) Each predictor’s class by training labels (dot), the shares of pathogenic and benign variants it scores, and ρ, in order of ρ; hairline, half of the pairs. (c) Sharp bound (band, not drawn where the circle covers it), covered AUROC (circle) and must-answer AUROC (diamond, omitted under the circle); dashed, chance; blue, four widely used predictors (CADD raw, ESM1b, AlphaMissense and REVEL); bottom rows, medians of the 30 missense predictors that rank under half their pairs, and gene + consequence, which reads no sequence, with the number of the 49 it is identified above.

Alt text: Reach and bounds of 49 predictors on 328,328 ClinVar substitutions. Five answer over 99.9% of variants; 31 rank under half of the pathogenic–benign pairs, and 32 bounds contain chance. The 30 missense predictors among the 31 have median covered and must-answer AUROCs of 0.94 and 0.62. Gene and consequence, which read no sequence, have an AUROC of 0.974, identified above 23 of the 49.

Scope shows as a wall: 36.7% of the variants (1.0% of them missense) are answered by at most 20 of the 49 predictors and hold 63.6% of the panel’s pathogenic variants (Supplementary Figure S15). Inside missense scope, declines largely span whole genes, and most genes that dbNSFP’s REVEL column leaves empty are almost wholly scored by REVEL’s own release (Supplementary Notes S5 and S39).

Reach falls into three bands: five predictors (CADD and four conservation tracks) answer over 99.9% of the variants, 13 answer 70% to 99.9% and 31 fewer than 70% (Fig. 2b; Supplementary Table S9). Those 31 score both variants of under half of the pathogenic–benign pairs (Supplementary Figure S9), and at such coverage a bound contains chance whatever the covered AUROC (Supplementary Note S36). In all, 32 bounds contain chance and 17 exclude it (Fig. 2c). For the 30 missense predictors among the 31, the median covered AUROC is 0.94 and the median must-answer AUROC 0.62.

### Which comparisons are decided, what conventions assume, and how far an ordering survives

On the whole panel, where each predictor is charged for variants outside its declared scope, the bounds identify 83 of 1,176 pairwise comparisons; 557 are not identified and 536 never can be (Fig. 3b, c). Inside declared scopes most comparisons stay undecided: 745 of 1,176. On the narrower of each pair’s two scopes, 431 of the 1,176 comparisons are identified, and 377 also pass a gene-clustered test of overlap at the 5% level [50,51] (Supplementary Notes S3 and S4). On the panel’s 202,643 missense substitutions, 429 are identified, 376 pass that test (confidence-set coverage 93.3%) and 316 Holm’s correction (Fig. 3b, c; Supplementary Note S3). Of the 83 identified on the whole panel, the test keeps 74 and Holm’s correction for the 1,176 comparisons 67. The lower ends of the bounds rest on a median of 14.5 effective genes on the missense subset (AlphaMissense’s on 1.5, 80% in TTN), yet removing the five genes that carry most endpoint variance drops only 2 of the 74 and 7 of the 377. Twenty-three of the 83 compare two conservation tracks, 47 one such track with another predictor, and only 13 involve none (Fig. 3d). Sixteen involve bStatistic, a regional background-selection statistic that reads near chance.

**Figure 3.**
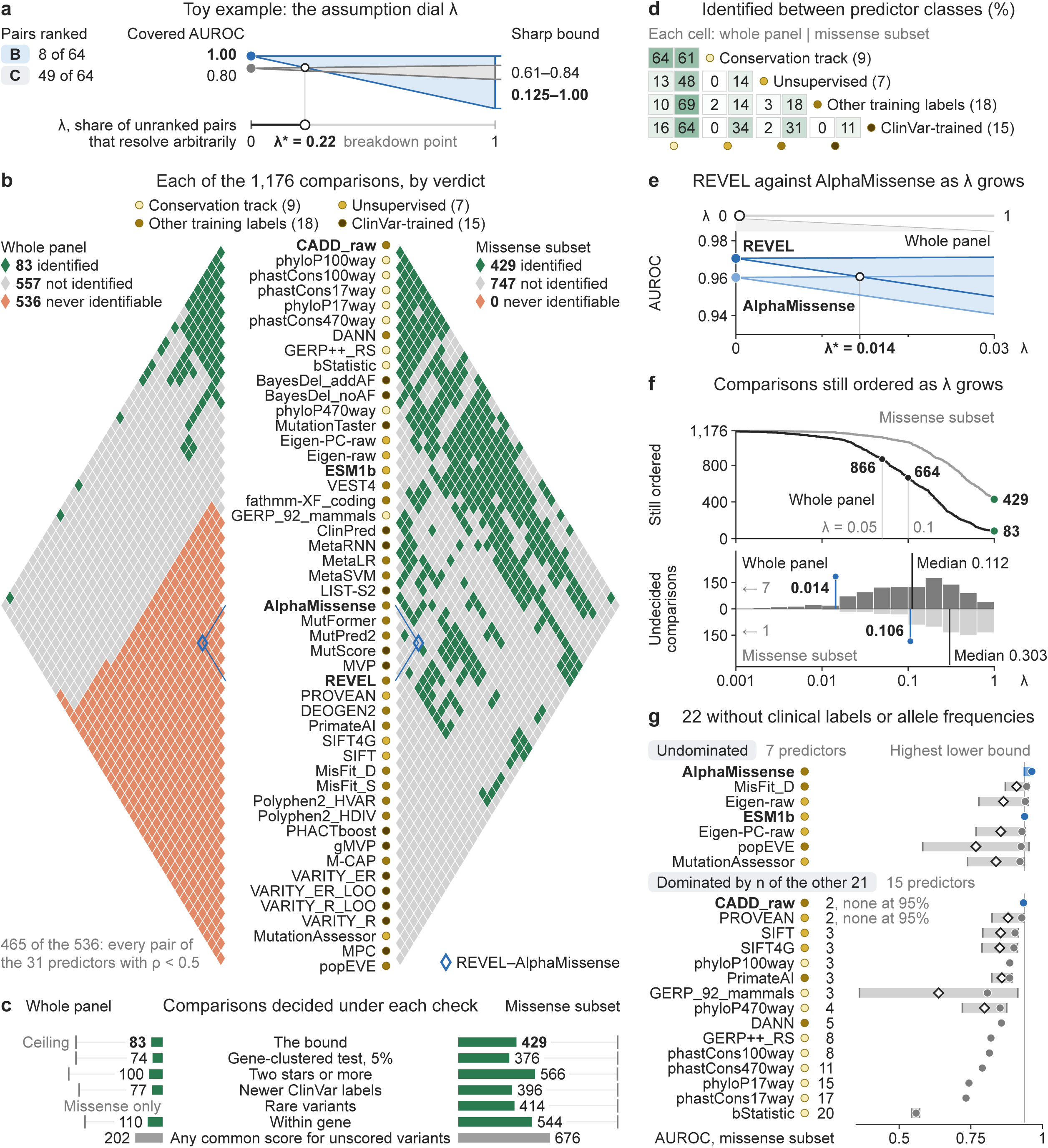
Which comparisons are decided, and how far they survive. (a) Figure 1a’s toy predictors B and C: as a share λ of unranked pairs resolves arbitrarily, each one’s interval grows from the covered area under the receiver operating characteristic curve (AUROC) at λ = 0 to the sharp bound at λ = 1; the two meet at the breakdown point λ*. (b) One cell per pair of predictors, in Figure 2’s order: whole panel left, missense subset right; verdicts in Figure 1’s colours; dots, predictor class; blue, the Rare Exome Variant Ensemble Learner (REVEL) against AlphaMissense. (c) Comparisons decided by the bound and under each check: gene-clustered 95% confidence sets (Supplementary Note S3), two or more review stars, newer ClinVar labels, rare variants (missense only, so no whole-panel bar; Supplementary Note S5) and within gene; end ticks, the pair-coverage ceiling; grey, orders holding whatever common score unscored variants get. (d) Percentage of pairs identified between predictor classes. (e) REVEL against AlphaMissense, whole panel, from Figure 2c’s covered AUROCs. (f) Comparisons still ordered against λ; below, undecided comparisons’ λ* (rules, medians; pins, REVEL–AlphaMissense; arrows, λ* below the axis). (g) Missense subset: bound, covered and must-answer AUROC of the 22 predictors trained without clinical labels or allele-frequency inputs, and how many of the other 21 lie wholly above (“none at 95%” marks a predictor none of whose dominations survives c’s gene-clustered confidence sets); line, the highest lower bound.

Alt text: The 1,176 comparisons among 49 predictors, one cell per pair. On the whole panel the bound identifies 83 and 536 never can be; on the missense subset, 429. At λ = 0.05, 866 stay ordered; undecided comparisons break at a median λ of 0.112, REVEL’s lead over AlphaMissense at 0.014. Seven of 22 predictors trained without clinical labels are undominated.

At two review stars or more, 100 are identified rather than 83, and 566 rather than 429 (Fig. 3c). As prospective assessments do [7,52], we also evaluate on the 232,563 labels newer than the predictors’ dated ClinVar training snapshots (four of the 15 ClinVar-trained predictors name none). On them, 77 whole-panel comparisons are identified, all among the 83 and none reversed, and 396 on the missense subset (Supplementary Note S5; Supplementary Table S26). Within genes rather than pooled [17,53], one of the 400 comparisons identified both ways reverses (Supplementary Notes S5 and S6). The predictors are correlated (211 of the 1,176 pairs rank-correlate above 0.80), so the counts describe this set.

Conventions decide what the bound leaves open, each by assumption (Supplementary Note S37; Supplementary Table S29). Covered AUROCs printed side by side rest on different variants: head to head on the variants both score, 58 of the 1,093 undecided comparisons reverse, 35 of them resolved by gene-clustered 95% intervals and 22 under Holm’s correction with the 1,093 as the family and only the reversals tested (Supplementary Note S4; Supplementary Table S29). Whatever common score declined variants are given, 202 of the 1,176 comparisons keep one order, 676 against 429 identified on the missense subset (the 202 on the whole panel, where 83 are identified; Fig. 3c). Filling each predictor’s unscored variants with its least deleterious score, as a 97-predictor benchmark does to check its ranking [5], orders all 1,176 comparisons on the missense subset. The ranking largely survives the opposite fill (Spearman 0.809), yet 205 of those orders reverse. Comparison on shared variants resolves 1,113 at 95% confidence, and its order is the opposite of the fill’s for 188 pairs, 162 of them resolved on shared variants: REVEL leads AlphaMissense by 0.0100 there and trails by 0.0178 under the fill. Whether declined variants are harder is mostly unresolved (median interval width 0.048). Most predictors read higher on the variants all score, as the sixth Critical Assessment of Genome Interpretation (CAGI6) reported [7] and as 41 of the 49 do. Yet declined variants are more often benign, and eleven near-complete predictors separate a predictor’s declined in-scope variants worse than its scored ones for 3 of 42, better for 7 and indistinguishably for 32 (Supplementary Note S37).

Let a share λ of each predictor’s unranked pathogenic–benign pairs resolve arbitrarily, the rest like its ranked pairs [25]: the λ at which the two predictors’ intervals meet is the comparison’s breakdown point λ* [54]; no smaller share can reverse it, and λ* exceeds one exactly when it is identified (Fig. 3a). On the whole panel 866 comparisons stay ordered at λ = 0.05, 664 at 0.1 (of 1,176), though there the unranked pairs include variants outside each predictor’s scope. Undecided comparisons break at a median λ of 0.112 (0.303 on the missense subset; Fig. 3f). At the reach dbNSFP distributes, REVEL’s lead over AlphaMissense breaks at λ = 0.014 on the whole panel and 0.106 on missense variants (Fig. 3e, f), and at 0.397 with the tools’ own releases (Supplementary Note S5). Of the 22 predictors trained without clinical labels or allele-frequency inputs, 15 are worse on the missense subset than another of the 22, 13 of them passing the gene-clustered test (Fig. 3g; Supplementary Note S7).

### Five widely used scores, a score that reads no sequence, and what a missing score means in the clinic

On the 1,434,335 ClinVar single-nucleotide variants (SNVs) at one review star or better, CADD [39] scores 99.88% and phyloP [37] and phastCons [38] 99.997%, but the missense-only REVEL [40] and AlphaMissense [41] 17.64% and 13.49% (Fig. 4a–c). Their covered AUROCs, 0.9575 and 0.9631, fall to must-answer AUROCs of 0.5256 and 0.5168, within bounds of [0.054, 0.998] and [0.035, 0.999], against CADD’s [0.989, 0.991] (Supplementary Table S8). On the variants both score, with genes resampled for each 95% confidence interval (CI), REVEL leads CADD by 0.0091 (95% CI 0.0028 to 0.0146) on 253,041 variants, and AlphaMissense leads CADD by 0.0316 (0.0279 to 0.0353) on 193,533 (Fig. 4e).

**Figure 4.**
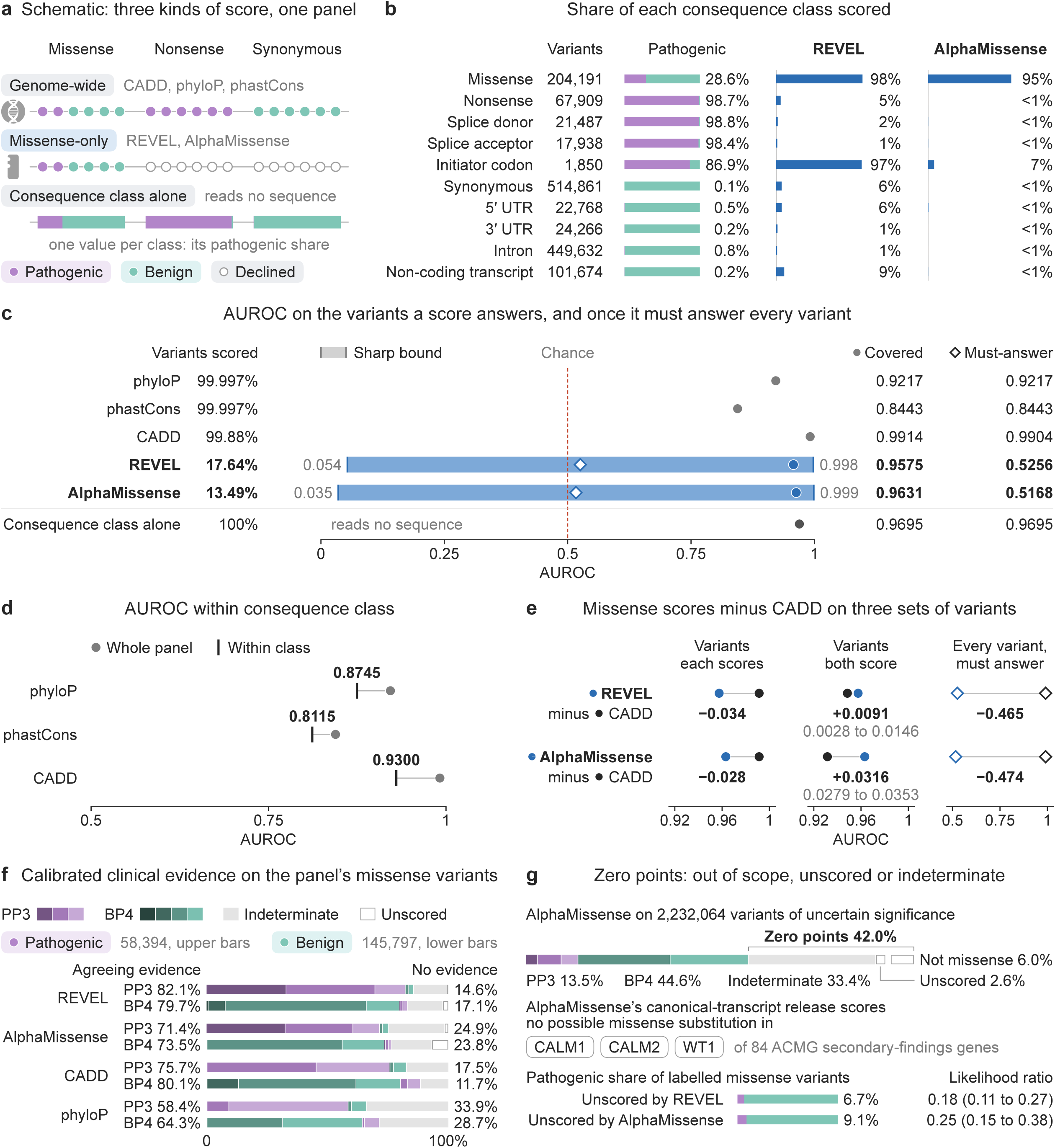
Five widely used scores on 1.43 million ClinVar variants, consequence class alone and calibrated evidence. Single-nucleotide variants at one review star or better (170,680 pathogenic, 1,263,655 benign); own-release scores. (a) Genome-wide and missense-only scores (open, unscored), and consequence class alone. (b) Ten classes (UTR, untranslated region): size, pathogenic share and share scored by the Rare Exome Variant Ensemble Learner (REVEL) and AlphaMissense. (c) Variants scored, sharp bound (band), covered area under the receiver operating characteristic curve (AUROC; circle) and must-answer AUROC (diamond); below, consequence class alone, out of fold. (d) c’s covered AUROC against AUROC within class (tick; pair-weighted mean of class AUROCs); REVEL and AlphaMissense omitted (too few classes); classes in Supplementary Figure S16a. (e) Each missense score minus Combined Annotation Dependent Depletion (CADD): c’s covered (left) and must-answer (diamonds) AUROCs, and on the variants both score (middle; 95% interval, genes resampled). (f) Calibrated pathogenic (PP3) and benign (BP4) evidence on missense variants, strongest first (PP3 strong to supporting; BP4 very strong to supporting), then indeterminate, unscored; upper bars, pathogenic, lower, benign; columns, shares agreeing with the label and with no evidence. (g) AlphaMissense on 2,232,064 ClinVar variants of uncertain significance (PP3, BP4, indeterminate, unscored, outside missense scope); secondary-findings genes its canonical-transcript release leaves unscored; pathogenic share of labelled missense variants a score leaves unscored, with likelihood ratios (95% intervals, 2,000 gene resamples) describing each release, not a variant.

Alt text: Five scores on 1.43 million ClinVar variants. REVEL and AlphaMissense score 17.64% and 13.49%, so covered AUROCs of 0.9575 and 0.9631 fall to must-answer AUROCs of 0.5256 and 0.5168, while consequence class alone reads 0.9695. AlphaMissense leaves 42.0% of variants of uncertain significance at zero points: 6.0% outside scope, 2.6% unscored, 33.4% indeterminate.

A score that reads no sequence sets the scale of these accuracies, as it does in published benchmarks [14,16,17]. Consequence class alone, estimated out of fold, reads 0.970 on the ClinVar panel, within 0.022 of CADD’s 0.991 (0.9695 against 0.9914 before rounding; Fig. 4c). Splice-donor and nonsense variants are 98.8% and 98.7% pathogenic, and synonymous and intronic variants 0.1% and 0.8% (Fig. 4b), shares partly reflecting how such labels are assigned [55]. Within consequence class, CADD reads 0.9300, phyloP 0.8745 and phastCons 0.8115 (Fig. 4d). On the dbNSFP panel, gene and consequence together read 0.974 but are identified above only 13 to 23 of the 49 predictors across rate estimators (Fig. 2c; Supplementary Note S9).

Under ClinGen’s calibrations [18,19], a missense score inside a calibrated interval earns pathogenic (PP3) or benign (BP4) computational evidence (Fig. 4f; Supplementary Figure S13; Supplementary Table S22). Removing the expert-panel missense labels that rest on such evidence [46,56] lowers each calibrated tool’s AUROC by 0.004 to 0.008 (Supplementary Note S8). A score between the intervals is indeterminate and earns no PP3 or BP4 points; nor does a missing score, which the calibration studies count within that interval [18,19]. Among the 84 secondary-findings genes [57], AlphaMissense’s canonical-transcript release, read by genomic position, scores no possible missense substitution in CALM1, CALM2 or WT1, unlike its isoform release, and leaves 10,572 of 235,209 unscored in TTN; CADD leaves none (Supplementary Table S31). Of the 2,232,064 ClinVar SNVs of uncertain significance at one review star or better, 42.0% receive no PP3 or BP4 points from AlphaMissense: 6.0% are not missense by ClinVar’s first-listed consequence, outside every missense calibration, 2.6% are unscored missense variants and 33.4% are indeterminate (Fig. 4g; Supplementary Table S30).

Yet a missing score is not neutral. Labelled missense variants that REVEL’s and AlphaMissense’s own releases leave unscored are pathogenic far less often than the panel’s, with likelihood ratios of 0.18 (gene-resampled 95% interval 0.11 to 0.27) and 0.25 (0.15 to 0.38; Supplementary Table S27). But dbNSFP’s REVEL column gives 0.58, and outside the 97 genes AlphaMissense’s release leaves wholly unscored its ratio is 0.12 (Supplementary Note S38): the ratios describe what each release omits, not how to classify a variant. Within scope a missing score is rare (AlphaMissense: 9 of 1,220 expert-panel missense classifications of uncertain significance without PP3 or BP4 evidence; Supplementary Note S38), so a variant without any score is mainly one outside the tool’s scope.

What a missense-scope predictor leaves unscored follows consequence class almost entirely, across genes and disease categories alike (Supplementary Note S39). Three gaps are concrete (Supplementary Table S32). First, seven of dbNSFP’s 32 missense-scope predictors score 0.016 to 0.053 of the 1,608 pathogenic initiator-codon variants but 0.436 to 0.649 of the 202 benign ones, most of which lie off the MANE Select start codon (MANE, Matched Annotation from NCBI and EMBL-EBI). Second, CADD, REVEL and AlphaMissense score none of the 1,717 mitochondrial-genome variants. Third, AlphaMissense’s own release scores no ClinVar missense variant in 97 of the 4,298 genes carrying at least ten.

On Smith and Kitzman’s splicing benchmark [58], SpliceAI (whose scores ClinGen’s splicing recommendations use for PP3 and BP4 [59]) and ConSpliceML [60] answer every benchmark variant but only 0.959 and 0.951 of its simulated background (Supplementary Tables S14 and S28; Supplementary Figure S14). Splicing Clinically Applicable Pathogenicity prediction (S-CAP) [61], the precision–recall winner on the benchmark’s MLH1 dataset, is not identified as a winner by AUROC (Supplementary Note S2).

### Evo 2 across nine species: the readout, the comparator and what conservation’s unscored variants cost

Whether Evo 2 outperforms conservation depends on the readout, the conservation score and its unscored variants (Fig. 5). The atlas is a case study in charging for reach, not a language-model benchmark. It measures separation from population variation, not prioritisation among coding candidates: positives are 84.6% to 100% coding, non-human negatives 0.4% to 2.8% (Fig. 5f). Negatives are Ensembl [62] population variants in seven species, common genotype sites of the Pig Genotype-Tissue Expression (PigGTEx) project [63] in pig and ClinVar benign or likely benign records in human, so we report species separately.

**Figure 5.**
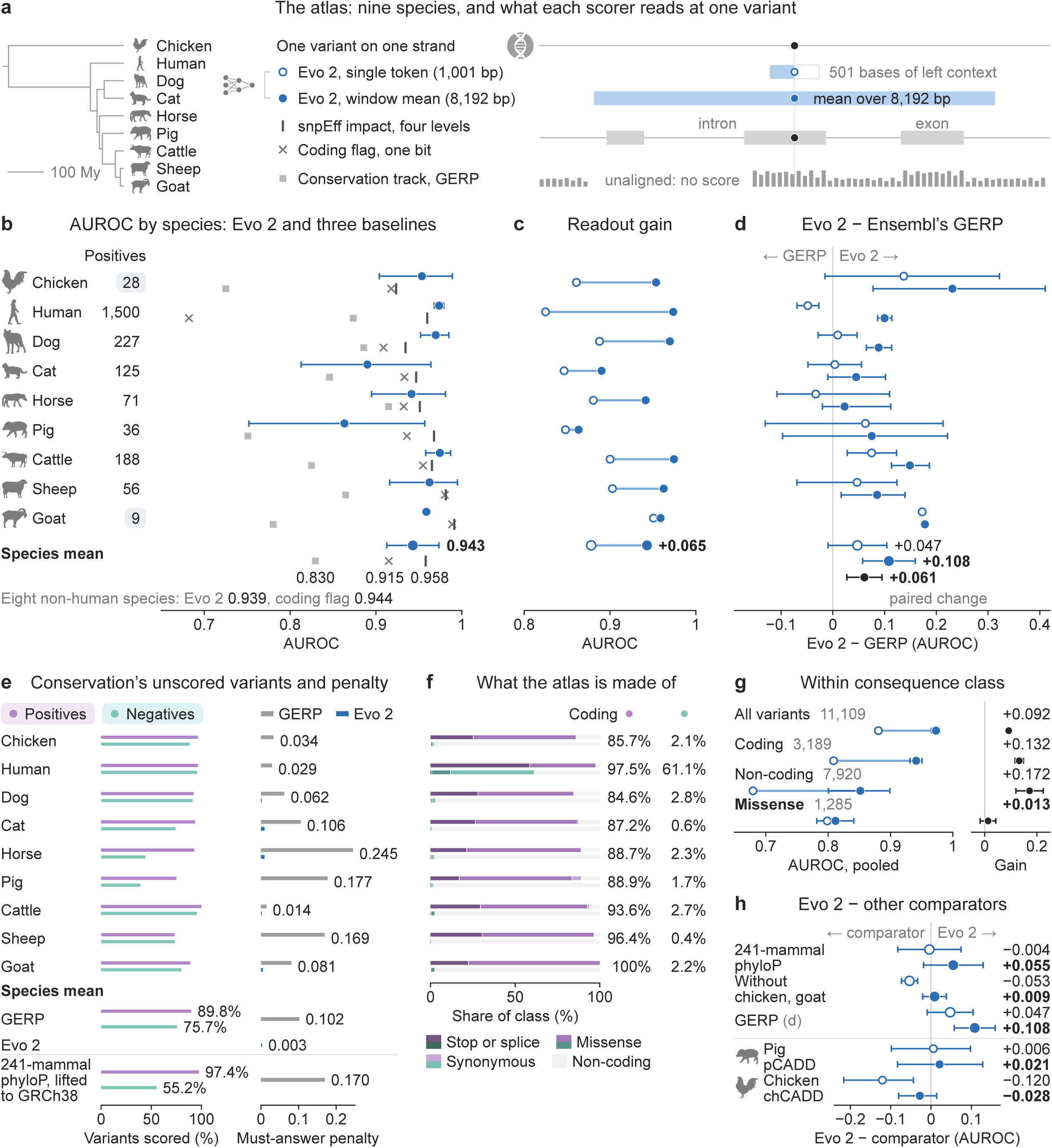
Evo 2 across nine species: what readout and comparator change. Catalogued alleles (positives) against population or, in human, ClinVar benign variants (negatives). (a) Species on a time-scaled tree [64] (My, million years) and what each scorer reads: Evo 2-40B’s single-token (ring) and window-mean (dot) readouts, two annotation baselines and the Genomic Evolutionary Rate Profiling (GERP) track. (b) Area under the receiver operating characteristic curve (AUROC) per species: Evo 2 at 8,192 bp (dot, Table 1’s locus-bootstrap 95% interval; species mean, t interval on eight degrees of freedom), snpEff impact (|), coding flag (×) and GERP where it scores (square); below, species means. Chips, under 30 positives (sizes in Table 1); goat, no interval. (c) Both readouts (dots of b); mean gain. (d) Evo 2 minus GERP on 9,529 variants all three score (locus-bootstrap 95% intervals), means and paired change (t intervals). (e) GERP’s reach on positives and negatives, its must-answer penalty (covered minus must-answer AUROC) beside Evo 2’s; species means below, with the lifted Zoonomia 241-mammal phyloP’s (phyloP-241). (f) Consequence classes of positives (upper bar) and negatives (lower bar), coding first; left column, coding share of positives, right, of negatives. (g) Pooled consequence strata of 11,109 window-scored variants: both readouts and gain (locus-bootstrap 95% intervals). (h) Evo 2 minus phyloP-241 (also without chicken, goat) and GERP (species means of d), pCADD (pig) and chCADD (chicken); intervals as in d.

**Table 1.** Cross-species discrimination by Evo 2-40B against Genomic Evolutionary Rate Profiling (GERP) and two annotation baselines.

| Species | N (positives) | AUROC (8,192-bp window mean) | AUROC (1,001-bp single token, 501 bases upstream) | AUROC (conservation-matched, 1,001-bp single token) | Conservation (GERP) | Coding flag | snpEff impact |
|---|---|---|---|---|---|---|---|
| Goat | 99 (9) | 0.959<br>(no interval) | 0.951<br>(no interval) | — | 0.780 | 0.989 | 0.991 |
| Chicken | 308 (28) | 0.954<br>[0.904, 0.989] | 0.861<br>[0.768, 0.941] | 0.818 | 0.725 | 0.918 | 0.924 |
| Pig | 396 (36) | 0.863<br>[0.752, 0.957] | 0.848<br>[0.731, 0.942] | 0.804 | 0.751 | 0.936 | 0.968 |
| Sheep | 616 (56) | 0.962<br>[0.916, 0.995] | 0.903<br>[0.823, 0.966] | 0.911 | 0.865 | 0.980 | 0.981 |
| Horse | 781 (71) | 0.941<br>[0.895, 0.981] | 0.880<br>[0.818, 0.979] | 0.846 | 0.915 | 0.932 | 0.951 |
| Cat | 1,365 (125) | 0.890<br>[0.813, 0.964] | 0.846<br>[0.784, 0.910] | 0.837 | 0.846 | 0.933 | 0.947 |
| Cattle | 2,068 (188) | 0.974<br>[0.958, 0.987] | 0.900<br>[0.868, 0.929] | 0.851 | 0.825 | 0.955 | 0.965 |
| Dog | 2,497 (227) | 0.970<br>[0.952, 0.985] | 0.888<br>[0.856, 0.918] | 0.846 | 0.886 | 0.909 | 0.935 |
| Human | 3,000 (1,500) | 0.974<br>[0.968, 0.979] | 0.825<br>[0.808, 0.842] | 0.761 | 0.874 | 0.682 | 0.960 |
| **Mean (macro)** | 11,130 (2,240) | **0.943<br>[0.913, 0.973]** | **0.878<br>[0.849, 0.907]** | **0.834** | **0.830** | **0.915** | **0.958** |
| **Mean, eight OMIA species** | 8,130 (740) | **0.939<br>[0.905, 0.973]** | **0.885<br>[0.856, 0.913]** | **0.845** | **0.824** | **0.944** | **0.958** |

notes. N is the panel size, positives in parentheses. The 8,192-bp column uses the 11,109 variants with a window score (Supplementary Note S32). Square brackets hold 95% confidence intervals, not bounds, from a locus-clustered bootstrap per species, each endpoint widened to the variant-level one where that is wider (a rule of unknown coverage). They are likely too narrow where effective loci are few: horse, chicken, sheep and pig (Supplementary Note S12). The conservation-matched means omit goat. Goat's nine positives occupy two loci, so its area under the receiver operating characteristic curve (AUROC) is a descriptive point estimate. Sheep, chicken and pig include positives on chromosomes, or parts of them, that their negatives miss (Supplementary Note S32), and pig's negatives, common genotype sites, confound its row with negative source. Mean rows, in bold, are unweighted species means with species-clustered t intervals, on eight and seven degrees of freedom respectively, targeting the mean over species exchangeable with these and conservative for these alone; their N cells are panel totals. Non-human positives come from Online Mendelian Inheritance in Animals (OMIA). Conservation-matching resamples negatives to the positives' GERP distribution, leaving GERP at chance by construction, and uses only the variants GERP scores (Supplementary Note S13). GERP is computed only where defined, so its column rests on fewer variants than the others (Supplementary Table S1).

Alt text: Evo 2 across nine species. Its AUROC averages 0.943 at the 8,192-bp readout, against 0.958 for snpEff impact. Its margin over GERP grows from +0.047 to +0.108 between readouts, but over 241-mammal phyloP it is +0.055, or +0.009 without chicken and goat, not resolved. GERP’s unscored variants cost it 0.102 AUROC, Evo 2’s 0.003. Positives are 84.6% to 100% coding.

Evo 2 has an unweighted species-mean AUROC of 0.943 at the 8,192-bp window-mean readout (95% CI 0.913 to 0.973) and 0.878 at the 1,001-bp single-token readout (0.849 to 0.907; Table 1; Fig. 5b, c). It outperforms annotation at neither readout: the snpEff [65] impact ordering reads 0.958, ahead of Evo 2 by +0.015 (95% CI −0.020 to +0.049), and a coding flag 0.944 against Evo 2’s 0.939 in the eight OMIA species. Added to impact and GERP in a cross-validated logistic model, Evo 2 raises the model’s species-mean AUROC by +0.008 (+0.002 to +0.014). Across checkpoints, on eight species, it reads 0.896 at 1B, 0.924 at 7B and 0.939 at 40B, though no step isolates model size (Supplementary Note S10). The gain from the single token to the 8,192-bp readout is smallest on missense variants, +0.013 (−0.017 to +0.042; Fig. 5g). On one porcine cis-eQTL panel Evo 2 reads near chance (0.488; Supplementary Note S25).

Over Ensembl’s GERP track, on the 9,529 variants both readouts and GERP score, Evo 2’s species-mean margin is +0.047 (95% CI −0.009 to +0.104) at 1,001 bp and +0.108 (+0.058 to +0.159) at 8,192 bp (Fig. 5d; Supplementary Note S21). With variants and comparator fixed, that is a paired change of +0.061 (+0.027 to +0.095). The readouts differ in left context (501 against 4,096 bases), aggregation and scorer at once, and about half the gain is context and scorer (Supplementary Note S20). GERP scores positives more often than negatives in eight of nine species. Charged for its unscored variants, GERP has a must-answer penalty of 0.102 by the species mean, against 0.003 for Evo 2 (Fig. 5e; Supplementary Note S32). With unscored variants bounded, the 8,192-bp comparison is identified in Evo 2’s favour in five of the eight species where it can be estimated, in sheep only when its positives on chromosomes without negatives are kept (Supplementary Notes S22 and S32).

Against the Zoonomia 241-mammal phyloP [44,45], read where each variant’s base lifts to GRCh38, Evo 2’s species-mean margin at 8,192 bp is +0.055 (−0.019 to +0.129), and +0.009 (−0.021 to +0.038) without chicken, which is not among the 241 mammals, and goat, whose positives occupy two loci; neither is resolved (Fig. 5h). At 1,001 bp it is −0.004 (−0.083 to +0.074), phyloP leading in cat, dog and human, and within consequence class (eight OMIA species, pairs pooled) −0.114 (−0.190 to −0.040) at 1,001 bp and −0.036 (−0.103 to +0.019) at 8,192 bp (Supplementary Note S43; Supplementary Figure S19b). Without chicken and goat, the 1,001-bp margin is −0.053 (−0.074 to −0.033), resolved in phyloP’s favour. The lift reaches 98.9% of non-human positives but 53.3% of negatives, pooled. The gap belongs to the pairwise chains, not the alignment: a 241-way phyloP read on the cattle genome scores 1,921 of the 1,985 cattle variants GERP and both readouts score (Supplementary Table S5). With the lift’s unscored variants bounded, Evo 2 is identified ahead only in cattle (in 71.6% of replicates) and human among the eight species with three or more positive loci (Supplementary Table S38); goat, with two, is identified at its point estimates alone. Species-trained scores, pCADD [66] in pig and chCADD [67] in chicken, are not resolved from Evo 2 at 8,192 bp, though chCADD, at 0.981, leads the single token by 0.120 (Fig. 5h). Protein predictors tested on OMIA variants [68] were not compared.

AlphaMissense [41], read at the aligned human residue [69], and SIFT [1], via Ensembl’s Variant Effect Predictor [70], reach positives far more often than negatives, so AlphaMissense’s lead over Evo 2 is not identified (Supplementary Notes S15 and S16). GPN-Star [30] is resolved neither ahead of Evo 2 nor behind it in chicken or human (Supplementary Note S16). On the BRCA1 saturation-editing panel [71], the 8,192-bp readout reads 0.874 and the single token 0.632 (Supplementary Note S18).

On the consequence-matched arm, within class phyloP’s covered species mean falls from 0.890 to 0.740, keeping 62% of its discrimination above chance, but its must-answer AUROC holds (0.727 against 0.720) and within missense it does not fall (Supplementary Note S44; Supplementary Table S39). For most scores, much of their discrimination thus comes from setting coding positives against non-coding negatives, and Evo 2, not scored on the arm, also falls within class on the atlas’s own variants (Supplementary Table S21).

### The same accounting beyond one panel

In ten published benchmarks returned by one PubMed query, 1,045 of 2,842 stated orderings (36.8%) are not identified once unscored variants are charged, but 2,070 of the orderings (73%) are implied by the sorted figure legends of two benchmarks that each rank about thirty tools [72,73] (Fig. 6a, b). Of the query’s 75 records, 10 state an AUROC ordering and report each predictor’s scored variants (Supplementary Note S24). Per benchmark the share runs from 0.0% to 95.2% (median 20.6%); nine of the ten state an unidentified ordering. If one benchmark’s sorted legend is not a ranking, the count is 773 of 2,516 (30.7%); allowing two-decimal AUROCs their rounding, at most 1,098 (38.6%); on clinical panels only, 1,034 of 2,786 (37.1%). Barbosa and colleagues also weight a threshold metric by each tool’s share scored [72]. Assessed apart, a hand-selected set of eight benchmarks leaves 10.3% unidentified (Supplementary Note S24).

**Figure 6.**
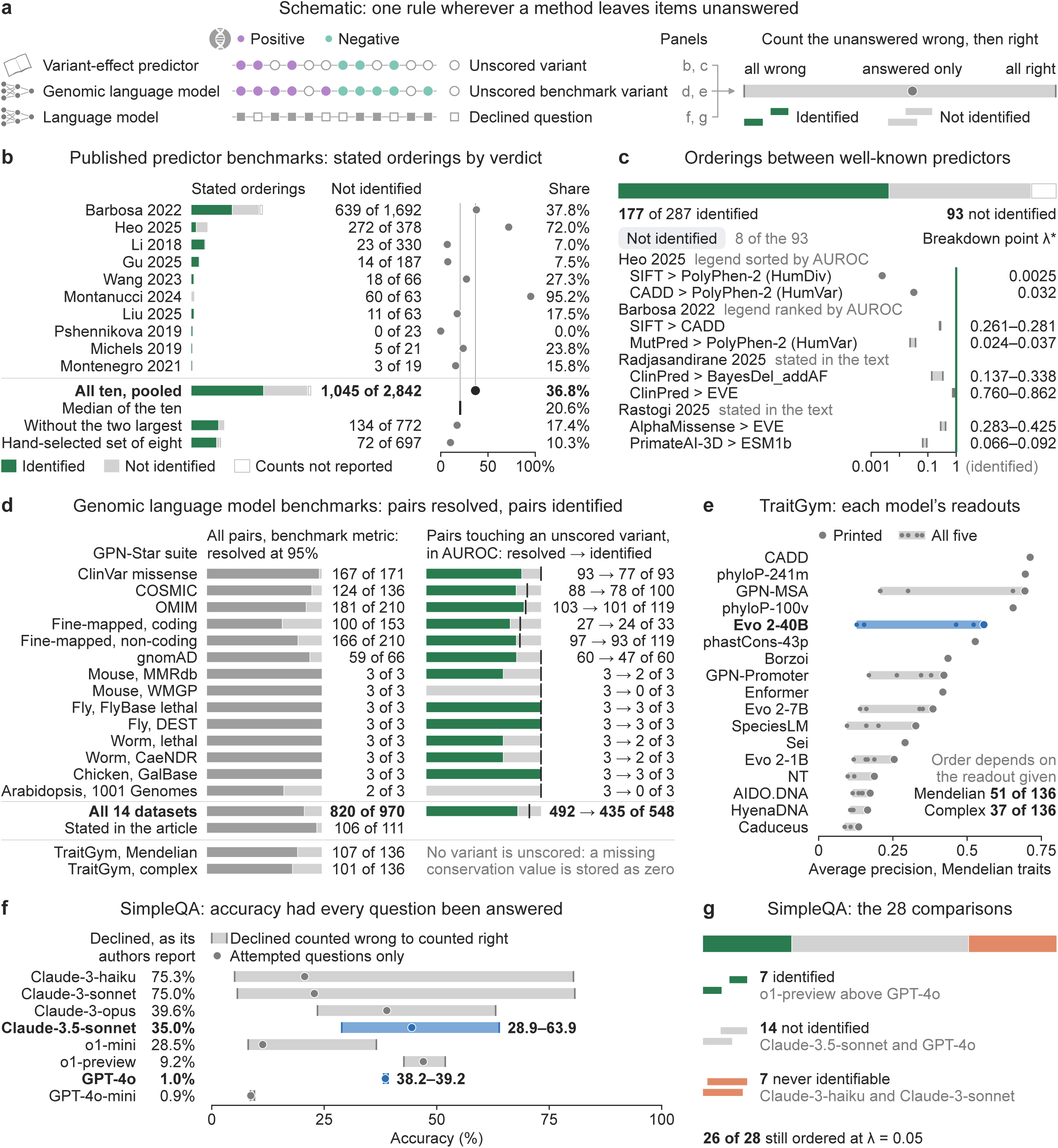
The same accounting beyond one panel. (a) A method leaves items unanswered (open); counting them wrong, then right, bounds its accuracy, and two methods are ordered only when their bounds are disjoint. (b) Ten benchmarks returned by one PubMed query that rank predictors by area under the receiver operating characteristic curve (AUROC) and report variants scored: stated orderings by verdict and share not identified (points; rules, pooled share and median). Below, pooled, median, without the two largest, and a hand-selected set of eight. (c) Stated orderings between thirteen predictor families in 9 of the 16 benchmarks; Table 2’s eight of the 93 not identified, breakdown points λ* (band, over unreported class splits); white, counts not reported (as in b). (d) GPN-Star’s 14 datasets, its article’s stated orderings, and TraitGym: pairs resolved at 95% in the benchmark’s metric (paired difference beyond 1.96 resampling standard errors); among pairs touching an unscored variant, pairs resolved in AUROC before charging (tick) and identified at 95% once charged (green). (e) TraitGym, Mendelian traits: average precision at the printed readout (point), across the five zero-shot readouts of each self-supervised model (band; blue, Evo 2-40B); right, pairs of the 136 whose order a choice of readouts reverses; NT, Nucleotide Transformer; m, v, p, mammals, vertebrates, primates. (f) SimpleQA as reported: bounds on the accuracy of the eight models of the benchmark’s article [48], from a declined question counted wrong (the share correct) to counted right; circle, attempted questions only; blue, the two models whose bounds are printed. (g) Their 28 comparisons by verdict.

**Table 2.** Printed orderings that the bound does not decide.

| **Benchmark (how the ordering is printed)** | **Panel (P / B)** | **Printed higher: AUROC; variants; bound** | **Printed lower: AUROC; variants; bound** | **λ\*** | **Articles printing the pair** | **Not identified, of the pair's stated orderings** |
|---|---|---|---|---|---|---|
| Heo and Kim [73] (legend sorted by AUROC) | 1,951 / 2,638 | SIFT: 0.8686; scored 1,800 of 1,951 P, 2,319 of 2,638 B; 0.704 to 0.893 | PolyPhen-2 (HumDiv): 0.8681; scored 1,652 of 1,951 P, 2,181 of 2,638 B; 0.608 to 0.908 | 0.0025 | 13 | 7 of 16 |
| Heo and Kim [73] (legend sorted by AUROC) | 1,951 / 2,638 | CADD: 0.8971; scored 1,951 of 1,951 P, 2,638 of 2,638 B; 0.897 to 0.897 | PolyPhen-2 (HumVar): 0.8961; scored 1,652 of 1,951 P, 2,181 of 2,638 B; 0.627 to 0.927 | 0.032 | 10 | 5 of 12 |
| Barbosa et al. [72] (legend ranked by AUROC) | 93 / 100 | SIFT: 0.85; 4 of 193 unscored; 0.816 to 0.856 | CADD: 0.84; 1 of 193 unscored; 0.832 to 0.842 | 0.261 to 0.281 | 9 | 2 of 7 |
| Barbosa et al. [72] (legend ranked by AUROC) | 278 / 193 | MutPred: 0.8; 192 of 471 unscored; 0.290 to 0.927 | PolyPhen-2 (HumVar): 0.78; 41 of 471 unscored; 0.665 to 0.812 | 0.024 to 0.037 | 7 | 4 of 6 |
| Radjasandirane et al. [4] (text) | 2,296 / 11,757 | ClinPred: 0.980; 13,947 of 14,053 scored; 0.971 to 0.980 | BayesDel_addAF: 0.977; 14,037 of 14,053 scored; 0.976 to 0.977 | 0.137 to 0.338 | 5 | 1 of 4 |
| Rastogi et al. [7] (text) | 6,103 / 4,353 | AlphaMissense: 0.879; 7 of 10,456 unscored; 0.878 to 0.879 | EVE: 0.838; 3,851 of 10,456 unscored; 0.344 to 0.933 | 0.283 to 0.425 | 3 | 3 of 4 |
| Radjasandirane et al. [4] (text) | 2,296 / 11,757 | ClinPred: 0.980; 13,947 of 14,053 scored; 0.971 to 0.980 | EVE: 0.896; 4,030 of 14,053 scored; 0.132 to 0.985 | 0.760 to 0.862 | 3 | 1 of 4 |
| Rastogi et al. [7] (text) | 6,103 / 4,353 | PrimateAI-3D: 0.859; 492 of 10,456 unscored; 0.790 to 0.870 | ESM1b: 0.852; 274 of 10,456 unscored; 0.814 to 0.859 | 0.066 to 0.092 | 1 | 1 of 1 |

notes. Each row is an ordering that a benchmark states and that the bound leaves unidentified for every class split of the unscored variants its printed counts allow. AUROCs are as printed; "variants" gives the scored counts by class where the benchmark prints them, and the number unscored or scored otherwise; the bound is the sharp bound at the most favourable split. P and B, variants the benchmark labels pathogenic and benign. λ\*, the breakdown point: the least share of each predictor's unranked pathogenic–benign pairs that, resolving arbitrarily, could reverse the order; a range covers the admissible class splits. Rows were chosen by one rule. Pairs of the thirteen well-known predictor families (Supplementary Note S40) were ordered by the number of the 29 articles that print both on one panel, and each pair enters at its stated, unidentified ordering with the lowest λ\*. A benchmark supplies at most two rows: a pair whose lowest instance lies in a benchmark that already has two takes its next instance from another benchmark or is passed over. Orderings whose coverage is printed as a share, not as counts, are excluded. Without the cap, the rule gives ten rows, and all ten rows come from the first two benchmarks (Supplementary Table S35). The two rows of Heo and Kim rest on a legend sorted by AUROC that the article's text does not call a ranking. Because the rule takes the lowest λ\*, near-ties lead: the first row's printed AUROCs differ by 0.0005 and the third's and fourth's by 0.01 and 0.02 at two decimals, within rounding; the last four rows show that printed gaps up to 0.084 can also stay undecided.

Alt text: Seven panels. In ten published benchmarks, 1,045 of 2,842 stated orderings are not identified (36.8%; per-benchmark median 20.6%), two benchmarks supplying most. In the GPN-Star suite 820 of 970 model pairs are resolved, and 435 of 548 are identified at 95% once unscored variants are charged. On SimpleQA, bounds decide 7 of 28 comparisons among eight language models.

Most orderings between well-known predictors are decided. With six further, hand-selected benchmarks, 9 of the 16 state orderings between two of thirteen predictor families, 287 in all: 177 are identified, 93 are not (61 of 253 if a sorted legend is not a ranking), 14 depend on the unreported class split and 3 are not assessable (Fig. 6c; Supplementary Note S40). In CAGI6 [7], AlphaMissense ranked above REVEL is identified (0.879 against 0.869), whereas PrimateAI-3D stated above ESM1b is not (0.859 against 0.852; breakdown point λ* 0.066 to 0.092). Of the 93, 25 have λ* below 0.1 (Supplementary Note S40); Table 2 shows eight, chosen by one rule. Of 997 pairs of tools from different families printed by two or more of the 29 AUROC-printing articles, 350 appear in opposite orders: the bound identifies both orders for 135 (the panel decides), one for 148 and neither for 65; 2 are not assessable (260 of 760 as printed), the latter count taking predictor names exactly as printed (Supplementary Note S40).

The genomic language model benchmarks are mostly decided. In the 14 datasets released with GPN-Star [30], 820 of 970 model pairs are resolved at 95% confidence in the suite’s own metric on jointly scored variants, as are 106 of the 111 orderings its article states (Fig. 6d; Supplementary Note S41). Of the 548 pairs touching a file with unscored variants, 492 are resolved at 95% in AUROC on jointly scored variants and 435 identified at 95% once unscored variants are charged (485 at the point estimates); AlphaMissense’s ClinVar lead over GPN-Star is not. On TraitGym [47], 29 and 35 of the 136 printed orderings (Mendelian and complex traits) are unresolved. Choosing each self-supervised model’s best of its five readouts on test labels inflates average precision by only 0.008 and 0.002 on average, yet the point order of 51 and 37 pairs depends on which readout each model is given (Fig. 6e). For 30 and 31 of these pairs, both orders are resolved.

The rule also applies outside genomics. SimpleQA reports the shares of 4,326 factual questions each language model answers correctly, incorrectly or not at all [48]. Its headline, the share correct, already counts a declined question wrong, the bound’s lower end. Its authors describe a penalty for wrong answers [48], as do Kalai and colleagues [22], and such penalties reorder these models [74]. The bounds ask which model would be more accurate had it answered every question. For the eight models, which decline 0.9% to 75.3% of questions, the bounds identify 7 of the 28 comparisons, each in the order of the share correct. Of the rest, 7 can never be identified and 14 are undecided, though 26 of the 28 stay ordered at λ = 0.05 (Fig. 6f, g; Supplementary Note S42).

## Discussion

This study measures the accounting around a predictor’s accuracy more securely than the skill behind it. An identified ordering holds whatever the unscored variants would have scored; an undecided one is not shown to be wrong, only to need an assumption about what was not scored, and its breakdown point says how strong that assumption must be. For a missense-only predictor, scope-matched accounting is closer than whole-panel accounting to CAFA’s partial evaluation mode [11], which scores a method only on the targets it answers. The must-answer penalty counts against printing REVEL’s covered AUROC beside CADD’s as though both came from one panel, not against the narrow tool (Supplementary Table S8).

Several results go against the story an audit of missing scores might be expected to tell. Most orderings that published benchmarks state between well-known predictors are decided, AlphaMissense above REVEL in CAGI6 among them, and most model pairs of the GPN-Star suite stay decided once unscored variants are charged. By the one measure available, declined variants are seldom measurably harder than scored ones, so nothing measured here would justify narrowing the bound. For a missense-scope predictor, which genes and disease categories go unscored follows almost entirely from consequence class. In the atlas, Evo 2 does not outperform annotation, and its lead, resolved over Ensembl’s conservation track at the window mean, is not resolved over the 241-mammal phyloP. The 97-predictor benchmark [5] is right that its ranking is stable under a least-deleterious fill; a ranking stable overall can still reverse individual orders under the opposite fill, and those orders need not be identified (Supplementary Note S37).

In the clinic, an indeterminate score and a missing one earn the same zero points but are different facts. In the releases read here, a labelled missense variant that REVEL or AlphaMissense leaves unscored is less often pathogenic than the panel’s variants, so a missing score carries information the calibrations do not use. Which release a laboratory reads can also decide whether a whole gene receives computational evidence. We propose no rule for using a missing score as evidence: the likelihood ratios describe what each release omits, on labels that may themselves rest on computational evidence (Supplementary Note S8). Reporting unscored and indeterminate variants apart, a step consistent with the calibration’s handling of EVE [19], would show a laboratory which of the two it holds.

The readout result concerns protocol: changing left context, aggregation and scorer together can change a verdict with variants fixed. Stating the readout makes a result reproducible, though not necessarily right. The comparisons we are aware of score a whole window rather than one position [16,31,34,47], so our single-token result warns about reproducibility and corrects no published verdict (Supplementary Note S21). Zero-shot studies of protein [75] and DNA [76] language models likewise find that the scoring rule changes accuracy, and Evo 2’s scores move with window length and with context that should be biologically neutral [77]. We scored zero-shot, as a model is deployed in a species with no labels of its own; what fine-tuning or probing adds depends on the task [47,78], and on regulatory tasks probing pre-trained representations adds little over models trained on one-hot-encoded sequence [79]. The near-chance cis-eQTL result, shared by per-base GERP (Supplementary Note S25), concerns zero-shot likelihood readouts, not supervised sequence-to-function models [80,81].

Nine limitations bear on the conclusions; Supplementary Note S31 lists further ones. First, Evo 2 was not scored on the consequence-matched arm, which is deposited with its builder, seed and the baselines a model must beat; the checkpoint comparison covers eight of the nine species, omitting human (Supplementary Note S10). Second, two benchmarks supply most of the audited orderings, a legend sorted by accuracy is counted as stating every pair, thirteen full texts were not read, and the frame is one database and one query (Supplementary Note S24 and Table S33). Third, the count of orderings printed in opposite directions depends on how printed predictor names are harmonised to tools (Supplementary Note S40). Fourth, the dbNSFP, CADD and REVEL scores are licensed and not deposited; the reach indicators and builders let others rebuild the panels under their own licences (Supplementary Note S34). Fifth, the confidence sets assume that no gene carries a large share of the variance, which fails on the missense subset: with the panel as the population, the 95% sets of the ten predictors resting on fewest genes covered 89.4% of the time, so the counts at 95% may be slightly high (Supplementary Note S3). Sixth, both corrections we apply in the atlas, charging GERP’s unscored variants and reading at 8,192 bp, move the verdict towards the model under test, as does reach accounting itself, which cannot penalise a model that scores any sequence. Stratifying by consequence class moves it the other way (Supplementary Note S14). Seventh, ClinVar labels may rest partly on the tools being scored, and this could be tested only on expert-panel labels (Supplementary Note S8). Eighth, circularity through training genes is not estimated (Supplementary Note S6). Ninth, OMIA carries no review status, so the atlas’s non-human positives cannot be held to ClinVar’s review standard (Supplementary Note S32).

Evo 2 scored on the consequence-matched arm, with an interval clear of the deep-alignment baseline, would change the atlas verdict. A fine-mapped cis-eQTL panel in another species on which the model’s interval lay above 0.5, its controls matched as in TraitGym [47], would overturn the near-chance regulatory result. Whether a pair is identified follows from reach as read from a given file, not from more data on the scored variants, so a release that scores more variants can decide comparisons this one cannot.

### Potential implications

Extending a recommendation of Harrizi and colleagues [17], we propose that benchmarks of variant effect predictors, and of any method that may decline to answer, report each method’s accuracy with its reach per label class, an out-of-fold baseline that reads no sequence and, for language models, the readout. A predictor’s covered AUROC and pair coverage are enough to recompute its must-answer AUROC and bound, so a published table can be re-read without rescoring and the orderings it states sorted into decided and undecided.

The glmtrust package (Additional file 2) runs the accounting on any panel labelled 0/1 that leaves each declined variant missing. Its audit command reports each scorer’s reach per class, must-answer AUROC and penalty, and a head-to-head comparison on the variants both scorers answer, withheld when their declared readouts disagree. Its reach command counts identified pairs and gives each pair’s breakdown point. Where score values cannot be redistributed [82], deposited reach indicators, as in Additional files 4 and 5, still let others recompute pair coverage.

Developers of a new predictor can score it on the Additional file 4 variants and set its bound beside those of the 49 dbNSFP predictors (Additional file 3). On the missense subset, a predictor with a covered AUROC of 0.98 must score both variants of 98.4% of pathogenic–benign pairs to be identified above AlphaMissense and 99.6% to be identified above REVEL. At 0.95, no reach suffices for either. A new genomic language model can be set against Evo 2 at both readouts and against conservation on the 11,130 atlas variants without rescoring Evo 2. On the consequence-matched arm, where annotation cannot separate same-class negatives from the positives, it can be set against the conservation baselines (Additional file 3).

## Methods

### Genomes, assemblies and annotations

Each species was scored on the assembly its panel was built and verified on (Supplementary Table S24), human on GRCh38; the BRCA1 saturation-editing and splicing assay panels stay on GRCh37, their published build. Consequences were annotated with snpEff 5.2 [65] on canonical transcripts (-canon), against databases built on each assembly with its matched Ensembl gene models (Supplementary Note S32 and Table S24), each variant taking its first, most severe, annotation. A variant is coding when that annotation’s first &-separated term is missense, synonymous, stop-gained, stop-lost, start-lost, frameshift, initiator-codon, splice-donor or splice-acceptor, as 3,191 atlas variants are, 1,286 of them missense; the four-level impact ordering ranks that term’s putative-impact class.

### Variant panels

The eight non-human species have 740 OMIA positives [32,33] (Table 1; Supplementary Note S32). OMIA carries no review status, and its classifications under the Animal Variant Classification Guidelines [83,84] are too sparse for a tier-restricted rerun, so its labels cannot be held to ClinVar’s review standard.

Positives were paired with population negatives at 10:1, ten per positive matched on its trinucleotide context and alternate allele, at sites holding no positive. The negatives were drawn, with no frequency threshold, from each species’ Ensembl variation release [62] (Supplementary Table S24) in seven non-human species and, for pig, from the PigGTEx v0 genotype panel on Sscrofa11.1 [85]. The matching balances mutational context between the classes, since most variant effect predictors and conservation scores track mutation rate [86]. Human, which OMIA excludes, uses ClinVar [35] SNVs on GRCh38 at 1:1 (Supplementary Note S32 and Table S12). The unweighted species mean is primary throughout, since the 1:1 human balance moves every prevalence-sensitive quantity. The conservation-matched control resamples negatives to the positives’ GERP profile in 12 quantile bins at up to 5:1 over 40 replicates, goat excluded (Supplementary Note S13); the consequence control recomputes discrimination within nested consequence strata.

Consequence-matched negatives were drawn for each atlas positive with an 8,192-bp score from its species’ population source: Ensembl variation in seven species, PigGTEx genotype sites in pig, ClinVar benign or likely benign records in human. They were matched on consequence class (the first snpEff term on the canonical transcript), reference trinucleotide and alternate base, exactly or on the opposite strand, with at most five per positive and arm (one in human) and none at a positive’s site. The same-class arm draws negatives of the positive’s own class, the synonymous arm synonymous negatives for a missense, stop-gained, start-lost or stop-lost positive. Candidates enter in the order of a keyed hash (seed 20261111), so the panel follows from the seed and the candidate pool. GERP, the 241-mammal phyloP, GPN-Star, pCADD and chCADD were read on it as on the atlas (Supplementary Note S44).

Human clinical panels. One ClinVar GRCh38 release, of fileDate 2026-07-28, built the ClinVar reach panel (SNVs at one review star or better) and the dbNSFP panel cut from it. The reach panel admits only aggregate classifications of pathogenic or benign, likely categories included (exact strings in Supplementary Note S32). A variant with several molecular-consequence terms takes the one with the lowest Sequence Ontology accession, the order ClinVar lists them in, a fixed rule rather than a severity ranking. The dbNSFP panel keeps 97.3% of the reach panel’s pathogenic but 12.8% of its benign SNVs (Supplementary Note S32). ClinVar classifications change between releases [87], so label age was read from two archived releases (fileDate 2021-01-02 and 2023-01-07; Supplementary Note S5).

Regulatory and TraitGym panels. The regulatory panel sets 5,000 candidate causal pig cis-eQTL variants against 15,000 controls within eGenes (genes with a cis-eQTL), all at a minor-allele frequency of at least 0.05 (Supplementary Note S25). The candidates were fine-mapped in PigGTEx [63] v0 with SuSiE-inf, an extension of the Sum of Single Effects (SuSiE) model [88–90]. The TraitGym readout comparisons use the scores TraitGym distributes for its Mendelian-trait and complex-trait test splits [47], 3,380 variants with 338 causal and 11,400 with 1,140.

### Evo 2 scoring

We used Evo 2 [31] zero-shot at the released 40B weights (evo2_40b; smaller checkpoints and scorers in Supplementary Notes S10 and S19). Evo 2’s StripedHyena 2 architecture builds on the Hyena operator family [91].

The single-token readout, called the 1,001-bp readout after its nominal window, is the next-token log-softmax at the variant position given the 501 reference bases upstream of it. The 8,192-bp readout is the window mean from the scoring code Evo 2’s authors released [31], with the variant at 0-based index 4,096. Upstream positions cancel between the reference and alternate windows, so this readout is the variant’s effect at its own position and the positions downstream, averaged over the window (Supplementary Note S19).

The score is logP(ref) − logP(alt), higher meaning more deleterious, and discrimination is reported on it unless labelled otherwise (some deposited tables store its negative; Supplementary Note S19). What the reference window’s likelihood separates on its own is measured in Supplementary Note S17. Scoring is forward-strand except in the strand analysis of Supplementary Note S29; TraitGym’s Evo 2 readout averages the two strands [47], which adds +0.0031 at 40B. Additional file 3’s 90 golden windows, with their expected scores at both readouts, let a rescoring be checked variant by variant.

### Comparators: predictors, conservation and splicing

Forty-nine published human predictors, nine of them label-free conservation or selection tracks, were taken from the academic branch of dbNSFP release 5.3.1 (README 5.3.1a) [92] and read from its rank-score columns, because the raw columns do not share a direction. Each was reduced to the maximum over transcripts and gene models, so a variant counts as reached if any transcript carries a value; where a score is missing on the canonical transcript, dbNSFP’s maintainers recommend checking its rank score [49]. The current release is 5.4, in which, the maintainers report, a score is dropped only when the transcript it was computed on has been retired, and which keeps 95% of the REVEL values of release 4.9a [49]. On the ClinVar panel, CADD v1.7 [39], REVEL v1.3 [40] (scored if any transcript carries a value) and AlphaMissense [41] (its canonical-transcript file) come from each tool’s own release, joined on GRCh38 position and alleles.

The cross-species baseline is GERP [42,43], the per-base conservation score Ensembl provides for all nine assemblies, read from the per-species Ensembl Compara bigWig track built on each panel’s assembly (releases 104 to 114; Supplementary Table S24). Alignment depth differs across species, and not only for chicken, the one non-mammal (Supplementary Note S21), so each species’ GERP is a different estimator. The Zoonomia 241-mammal phyloP [44,45] was read from the University of California, Santa Cruz (UCSC) GRCh38 track, non-human variants at the position their base lifts to through the UCSC chain of their assembly [93] (GenArk hub chains [94] for cat, sheep and goat). A variant without a lifted value is unscored, and phyloP’s bound is reported. pCADD [66] and chCADD [67] were read on the pig and chicken assemblies without a lift (Supplementary Note S43).

Scores for SpliceAI [3], Pangolin [95], Super Quick Information-content Random-forest Learning of Splice variants (SQUIRLS) [96], Splicing-based Analysis of Variants (SPANR) [97], Hexamer Additive Linear (HAL) [98], Modular Modeling of Splicing (MMSplice) [99], S-CAP [61] and ConSpliceML [60], on the benchmark variants and the simulated background, are taken from the supplementary tables of Smith and Kitzman [58]. Further baselines and comparators are specified in Supplementary Notes S9, S20, S32 and S33.

### Reach, penalty and partial identification

A variant is reached when the method returns a finite score.

Call A_cov_ the covered AUROC, and n and k the panel and reached counts per class. The must-answer AUROC is then (A_cov_·k□·k□ + ½(n□n□ − k□k□)) / (n□n□) = 0.5 + (A_cov_ − 0.5)ρ, where ρ = r□r□= k□k□/(n□n□), r being class reach.

The target is the AUROC θ that a predictor would attain on the whole panel had it scored every variant. With nothing assumed about the declined variants, θ lies in the sharp bound [ρA_cov_, ρA_cov_ + (1 − ρ)] [24]. The bound’s lower end places every declined positive below every negative and every declined negative above every positive; its upper end does the reverse. Keeping declined variants in the accounting follows the intention-to-diagnose approach of diagnostic-accuracy studies, in which non-evaluable results stay in the accuracy table [100]. A comparison is identified, or decided, when the two predictors’ bounds are disjoint. The bound and the rule hold for any metric averaging a contribution between 0 and 1 over items or pairs, ρ being the share of its weight answered and A the covered value (Supplementary Note S36). Since the bounds use no relation between two predictors’ missing answers, disjoint bounds are necessary as well as sufficient where their ends are attained, as for item averages and AUROC. Predictor 1 is then identified above predictor 2 exactly when ρ□A□ + ρ□(1 − A□) > 1, which requires ρ□ + ρ□ > 1. Because the endpoints are themselves estimates, we also report per-comparison confidence sets of the Imbens–Manski type [50,51] and family-wise counts by Holm’s step-down procedure [101] (Supplementary Note S3). The sets cover a difference with the stated probability if the two ends of its bound are asymptotically normal across genes, which we assume and do not prove. A comparison declared identified at 95% has then passed a test, at the 5% level, of the hypothesis that the two bounds overlap (Supplementary Note S36). Under the contamination model [25], θ lies in [A_cov_ − (1 − ρ)λA_cov_, A_cov_ + (1 − ρ)λ(1 − A_cov_)]. With predictor 1 the one whose covered AUROC is higher, the two intervals meet at λ* = (A□ − A□)/((1 − ρ□)A□ + (1 − ρ□)(1 − A□)), the breakdown point [54]: below it the order holds under every resolution, and it exceeds one, the sharp bounds never meeting, exactly when the comparison is identified (Supplementary Note S7).

For scope matching, each predictor’s scope is read from its documentation (Supplementary Table S25) as the whole panel, coding variants (canonical splice sites included) or missense variants, membership following ClinVar’s consequence term, and each pair is compared on the narrower of its two scopes. Alternative readings are sensitivity analyses (Supplementary Note S4). We set three conventions against the bound (Supplementary Note S37). The first fills each unscored variant with the predictor’s lowest dbNSFP rank score on the panel (least deleterious) or its highest. The second compares on the variants both predictors score, a difference being resolved when its 95% percentile interval over 2,000 gene resamples excludes zero. The third gives every declined variant one common score, a pair keeping one order when every such score, from below the lowest scored value to above the highest, orders it the same way (Supplementary Note S5). Whether declined variants are harder was measured with the eleven predictors that score more than 0.99 of the missense subset (Supplementary Note S37). A calibrated level’s likelihood ratio is the share of pathogenic variants at that level over the share of benign ones, set against the odds each strength targets [102] (Supplementary Note S8).

Published benchmarks came from one PubMed query for comparisons of variant predictors on clinical labels, fixed before it was run on 11 November 2026 and screened on title, abstract and open full text (Supplementary Note S24 and Table S33). A stated ordering counted as identified only when the two bounds, from the pair coverage each article’s counts allow, are disjoint for every class split of the unscored variants those counts admit. A second, hand-selected set of articles was read the same way, reported apart and joined to the first (Supplementary Notes S24 and S40). On both TraitGym splits, each of its 17 models was read at the readout its leaderboard prints and each of its ten self-supervised models also at all five zero-shot readouts TraitGym distributes (Supplementary Note S41), and a pair’s order was taken as resolved when a difference cleared zero by 1.96 standard errors over 2,000 resamples of whole chromosomes. Supplementary Note S27 asks the same question of the 34 score files TraitGym distributes for the Mendelian split, resampling the causal variants’ loci as well. In the benchmarks released with GPN-Star [30] and TraitGym [47], no model was rescored. Each GPN-Star file’s orientation was read without labels, from its stored range and its rank correlation with the files beside it. A pair was resolved when its difference in the benchmark’s own metric cleared zero by the same rule as on TraitGym, over 2,000 paired resamples drawn as the article draws them (Supplementary Note S41). For SimpleQA [48], the shares of questions eight language models answered correctly (c) and did not attempt (d) were taken from the article’s third table. Accuracy had a model answered every question lies in the sharp bound from c to c + d, the share attempted taking the place of pair coverage (Supplementary Note S42).

### Metrics, tests and intervals

Discrimination is AUROC unless otherwise stated, computed with scikit-learn’s roc_auc_score [103]; calibration and the conformal layers are specified in Supplementary Note S28.

Multiplicity is controlled in the Westfall–Young max-T families [104] of Supplementary Note S26, in the per-species locus-bootstrap tests of Evo 2 against GERP at each readout (over the eight estimable species, by Benjamini–Hochberg [105] at q ≤ 0.05), and by Holm’s procedure in the identification comparisons, the head-to-head comparisons on shared variants and the disease-category tests. Every other family is reported as estimates with intervals and carries no headline claim (Supplementary Notes S3, S21 and S23).

A mean over species carries a t interval on one degree of freedom fewer than the species it averages (eight for all nine), the across-cluster t construction of Ibragimov and Müller [106], because a nine-cluster percentile interval undercovers at any number of resamples. Precision-weighted pools over species are random-effects meta-analyses on locus-resampled standard errors (Supplementary Notes S14 and S23). On gene-resampled analyses, point estimates, bounds and identification counts describe the finite panel, and intervals the population of genes it samples. Where a paired bootstrap is impractical, we use DeLong’s method [107,108], whose variant-level interval is not taken as evidence of a real difference. Summed within genes, its placements give the gene-clustered variance behind the identification confidence sets (Supplementary Note S3). Supplementary Note S3 defines the effective number of genes behind an endpoint. Table 1’s per-species intervals use B = 20,000 resamples (Supplementary Note S12). Additional file 3’s README lists each estimand’s resampling unit, replicate count and seed. A script in Additional file 3 reproduces every printed endpoint, the variant-level ones to within 0.002.

We tested equivalence by two one-sided tests at a margin δ = 0.02, set after we saw the data; no equivalence is declared at that margin, so no conclusion rests on it (Supplementary Notes S10 and S22).

### Circularity and unscored variants

Of the two kinds of circularity [14], testing on training variants is checked in part on labels newer than the predictors’ stated ClinVar training data (Supplementary Note S5). Testing on training genes is not estimated, though within-gene AUROCs exclude the signal it exploits (Supplementary Note S6). A third kind, label circularity, in which a ClinVar label rests on the tools being scored, is tested on the ClinGen Evidence Repository [46]. A label depends on computational evidence when its American College of Medical Genetics and Genomics/Association for Molecular Pathology (ACMG/AMP) points [102] meet its class threshold only with PP3 or BP4, and each tool’s AUROC is compared with and without such labels (Supplementary Note S8). PP3 and BP4 evidence is assigned on missense variants from the calibrated intervals of Pejaver and colleagues [18] for REVEL, CADD and phyloP and of Bergquist and colleagues [19] for AlphaMissense.

Variants of uncertain significance are the reach panel release’s single-nucleotide records so classified, at one review star or better. A variant earns zero points when ClinVar’s first-listed consequence is not missense, when the release read carries no score for it or when its score is indeterminate, and the three are counted apart (Supplementary Note S38). AlphaMissense scores, given to four decimals, count as BP4 at or below 0.169 and PP3 at or above 0.792, and those in between are indeterminate. Where the canonical-transcript release holds several rows for one substitution, the last is read. AlphaMissense’s isoform release is read, by GRCh38 position and alleles, only for whether it holds a row at a substitution (Supplementary Note S38 and Table S31). Possible substitutions in the 84 genes of the ACMG secondary-findings list, version 3.3 [57], are enumerated from the MANE Select coding sequence [109], on which expert-panel classifications are also placed by position. The likelihood ratio of a score’s unscored level is defined in the same way on missense variants, with a percentile interval over 2,000 gene resamples. Unscored variants (Supplementary Note S39) are analysed by gene, with Genome Aggregation Database (gnomAD) v4.1 constraint (LOEUF, the loss-of-function observed/expected upper bound fraction [110]), HUGO Gene Nomenclature Committee (HGNC) gene groups [111] and MANE v1.5. They are also analysed by the

16 top-level Medical Subject Headings (MeSH) disease categories, mapped through MedGen from ClinVar’s conditions. A category’s expected share gives each pathogenic variant the panel-wide unscored share of its consequence class, and tests are Holm-corrected.

### Software and reproduction

Scoring ran on cloud graphics processing units (GPUs) in containers started from NVIDIA’s BioNeMo framework image 2.6.2 (recipe in Additional file 2; Supplementary Note S19). Twenty-seven scripts in Additional file 3 check 1,749 printed values, 1,500 of them from the deposit alone and 249 with scores the deposit withholds (Supplementary Notes S34 and S35).

The analysis code in Additional file 2 used NumPy 2.5.1 [112], SciPy 1.18.0 and scikit-learn 1.9.0 [103]. Conservation queries used pyBigWig, and the phyloP cross-check pybigtools 0.3.0; coordinate lifts used pyliftover 0.4.1, and allele frequencies PLINK2 [113]. Cattle phyloP scores came from Zenodo record 13332541, which accompanies Zhao and colleagues [114].

## Supporting information

Additional_file_3_Recompute_tables_and_scripts

## Abbreviations

ACMG/AMP: American College of Medical Genetics and Genomics/Association for Molecular Pathology
AHIDA: Anstee Hub for Inherited Diseases in Animals
AUROC: area under the receiver operating characteristic curve
BP4: ACMG/AMP code for benign computational evidence, applied at a calibrated strength
CADD: Combined Annotation Dependent Depletion
CAFA: Critical Assessment of Functional Annotation
CAGI6: sixth Critical Assessment of Genome Interpretation
CC0: Creative Commons Zero
CC BY: Creative Commons Attribution
CC BY-NC: Creative Commons Attribution-NonCommercial
CC BY-NC-ND: Creative Commons Attribution-NonCommercial-NoDerivatives
CC BY-NC-SA: Creative Commons Attribution-NonCommercial-ShareAlike
CI: confidence interval
ClinGen: Clinical Genome Resource
dbNSFP: database for nonsynonymous single-nucleotide polymorphisms’ functional predictions
eQTL: expression quantitative trait locus
ESM: Evolutionary Scale Modeling
ESM1b: Evolutionary Scale Modeling 1b
EVE: evolutionary model of variant effect
FarmGTEx: Farm Animal Genotype-Tissue Expression
GERP: Genomic Evolutionary Rate Profiling
gnomAD: Genome Aggregation Database
GPN-Star: Genomic Pretrained Network with Species Tree and Alignment Representations
GPU: graphics processing unit
HAL: Hexamer Additive Linear
HGNC: HUGO Gene Nomenclature Committee
JSON: JavaScript Object Notation
LOEUF: loss-of-function observed/expected upper bound fraction
MANE: Matched Annotation from NCBI and EMBL-EBI
MeSH: Medical Subject Headings
MMSplice: Modular Modeling of Splicing
My: million years
OMIA: Online Mendelian Inheritance in Animals
PigGTEx: Pig Genotype-Tissue Expression
PolyPhen-2: Polymorphism Phenotyping v2
PP3: ACMG/AMP code for pathogenic computational evidence, applied at a calibrated strength
REVEL: Rare Exome Variant Ensemble Learner
S-CAP: Splicing Clinically Applicable Pathogenicity prediction
SIFT: Sorting Intolerant From Tolerant
SNV: single-nucleotide variant
SPANR: Splicing-based Analysis of Variants
SQUIRLS: Super Quick Information-content Random-forest Learning of Splice variants
SuSiE: Sum of Single Effects
UCSC: University of California, Santa Cruz
UTR: untranslated region.

## Additional Files

Additional files 2 and 3 are zip archives, which the submission system would unpack, so they are provided, byte-identical, through the Zenodo record cited under Data Availability.

### Additional file 1

File name: 12_Additional_file_1_Supplementary_v1.1.docx File format: Microsoft Word document (.docx)

Title of data: Supplementary Notes, Tables and Figures

Description of data: Supplementary Notes S1 to S44, Tables S1 to S39 and Figures S1 to S20: the extended Methods and provenance, the identification and per-gene accounting, calibrated evidence and label circularity, the atlas’s controls, readout analyses and negative results, the audit of published benchmarks and the limitations in full.

### Additional file 2

File name: 13_Additional_file_2_Code.zip

File format: ZIP archive (.zip) of Python and shell source, JavaScript Object Notation (JSON), Markdown and Apache Parquet files, among others; Parquet readers at https://parquet.apache.org/

Title of data: Analysis code, recompute layer and the glmtrust package

Description of data: The analysis code (src/, tools/ and analyses/), the recompute-layer JSON files and glmtrust 0.1.1, which implements the reach accounting with breakdown points, the sequence-blind baselines and the calibration, conformal and selective-prediction layer, with its reproduction benchmarks.

### Additional file 3

File name: 15_Additional_file_3_Recompute_tables_and_scripts.zip

File format: ZIP archive (.zip) of Apache Parquet, comma-separated values, tab-separated values, JSON, Markdown, Python, shell and plain-text files

Title of data: Reduced per-variant tables and recompute scripts

Description of data: Analysis-ready per-variant tables, tables of breakdown points, per-gene bounds and values audited from published benchmarks, each with a README data dictionary; 27 checking scripts that print each expected value beside its recomputation; and, for Figures 1 to 6 and S8 to S20, the code that draws each and the files it reads (Additional file 2 draws the others; Supplementary Notes S34 and S35).

### Additional file 4

File name:

16_Additional_file_4_dbNSFP_reach_panel_coordinates_and_reach_only.parquet File format: Apache Parquet (.parquet); Parquet readers at https://parquet.apache.org/

Title of data: Per-variant reach matrix for 49 dbNSFP predictors Description of data: The 328,328-variant panel of ClinVar substitutions that dbNSFP annotates as nonsynonymous or splice-site, on GRCh38: coordinates, labels, metadata and one boolean reach indicator per predictor, true where the predictor returned a value and false where it declined. The score values are withheld for licensing reasons (see Data Availability); the file is released under Creative Commons Zero (CC0) 1.0 with the written permission of dbNSFP’s maintainers.

### Additional file 5

File name: 17_Additional_file_5_ClinVar_reach_panel_conservation_and_reach.parquet File format: Apache Parquet (.parquet); Parquet readers at https://parquet.apache.org/

Title of data: Per-variant conservation values and reach matrix for CADD, REVEL, AlphaMissense, phyloP and phastCons

Description of data: All 1,484,836 ClinVar single-nucleotide variants on GRCh38 classified pathogenic or benign (likely categories included), before the review-status filter: coordinates, labels, review status, consequence, the phyloP and phastCons values and one boolean reach indicator per scorer. The CADD, REVEL and AlphaMissense values are withheld (see Data Availability); the reach indicators are released under CC0 1.0 on the licence reading set out in LICENSING.md (Additional file 3), and the phyloP and phastCons values are products of UCSC. Applying the one-star review-status filter cuts from it the 1,434,335-variant panel this article reports.

### Additional file 6

File name: 18_Additional_file_6_splicing_deployment_background.parquet

File format: Apache Parquet (.parquet); Parquet readers at https://parquet.apache.org/

Title of data: Simulated 500,000-variant deployment background with eight splice-predictor scores

Description of data: The eight splice-predictor scores of Smith and Kitzman [58], redistributed under Creative Commons Attribution (CC BY) 4.0 with attribution, on hg19 (GRCh37) coordinates with each variant’s transcript, reference and alternate allele, and one boolean reach indicator per predictor.

## Declarations

### Ethics approval and consent to participate

Not applicable. The study used only secondary data, public except as stated under Data Availability, and generated no new data from humans or animals.

### Consent for publication

Not applicable.

### Availability of Source Code and Requirements

Project name: glmtrust 0.1.1

Project home page: https://github.com/asediya/evo2-reliability-atlas (glmtrust in its glmtrust/ directory; Additional file 2 is the identical review copy)

Operating system(s): Platform independent; Additional file 3’s run_all.sh needs a POSIX shell

Programming language: Python (3.9 or later for glmtrust; the analyses ran under 3.12.10 and Additional file 3’s scripts were checked under 3.13.13)

Other requirements: glmtrust needs NumPy 1.21 or later, SciPy 1.7 or later and scikit-learn 1.0 or later, with polars optional; the analysis layer adds pandas 3.0.3, polars 1.42.1 and PyArrow 24.0.0, among other packages listed in the requirements file of Additional file 2. Scoring with Evo 2 also needs a GPU host with evo2 and vortex, which the container recipe in Additional file 2 provides; re-analysing the deposited scores does not. Seven of Additional file 3’s scripts check part of their values against scores the deposit withholds: a dbNSFP release, the CADD, REVEL and AlphaMissense releases or the CADD scores TraitGym distributes, each obtained under the reader’s own licence, or pCADD and chCADD scores, whose hosts state no licence.

### License: MIT

RRID: SCR_029089

Any restrictions to use by non-academics: None for the code (MIT licence). The dbNSFP release, under Creative Commons Attribution-NonCommercial-NoDerivatives (CC BY-NC-ND) 4.0, and several predictor score files carry non-commercial terms, as do the PigGTEx-derived values in Additional files 2 and 3, under Creative Commons Attribution-NonCommercial (CC BY-NC) 4.0 (Data Availability).

### Data Availability

The data sets supporting the results of this article are available in the Zenodo repository, https://doi.org/10.5281/zenodo.22046811 [36], and as Additional files 2 to 6. The reduced per-variant tables from which this article’s values can be recomputed are Additional files 3 to 6, and the analysis code and recompute layer are Additional file 2. Files the authors generated are released under CC0 1.0 and the code under the MIT licence; third-party values carried in the deposits stay under their sources’ terms, which LICENSING.md and LICENSES.tsv in Additional file 3 give file by file. Additional file 4 holds dbNSFP-derived presence flags and no score value, and is released under CC01.0 with the written permission of dbNSFP’s maintainers [92]. Values carried under terms other than CC0 1.0, CC BY 4.0 or unrestricted use are PigGTEx’s fine-mapping probabilities, eQTL coordinates and pig variant sites (CC BY-NC 4.0) [63]; GPN-Star scores (Apache License 2.0) [30]; TraitGym scores (MIT licence) [47]; TimeTree 5 divergence times (TimeTree’s terms) [64]; the 40 SimpleQA values (quoted from an article under arXiv’s non-exclusive licence 1.0) [48]; and OMIA identifiers and phenotype names (attributed to OMIA, whose records remain under the University of Sydney’s copyright) [32,33]. The cattle variant sites and allele frequencies computed from the Farm Animal Genotype-Tissue Expression (FarmGTEx) imputed-genotype release [115], which records no licence, are released under CC BY 4.0 with attribution, a reading for the journal’s curators to confirm.

All data analysed are public secondary datasets, with one exception stated below; Methods and Supplementary Table S24 give the releases. They are OMIA [32,33]; Ensembl assemblies, gene models, orthologues, variation, GERP tracks and SIFT [62], with its Variant Effect Predictor [70]; ClinVar [35]; the ClinGen Evidence Repository [46] and Criteria Specification Registry; MedGen (file release 2026-11-10); dbNSFP 5.3.1a [92]; the CADD [39], REVEL [40] and AlphaMissense [41] releases, with UniProt’s ID mapping [116]; pCADD [66] and chCADD [67]; UCSC phyloP and phastCons, cattle phyloP [114] and the Zoonomia 241-mammal phyloP [44,45], with UCSC and GenArk chain and sequence files [93,94]; PigGTEx v0 [63]; the benchmark datasets released with GPN-Star [30] and TraitGym [47], with TraitGym’s leaderboard; the SimpleQA results table [48]; ProteinGym [6]; the EVEE shard [117]; Smith and Kitzman’s splicing data [58]; BRCA1 and LDLR function scores [71,118] from MaveDB [119]; MANE v1.5 [109]; HGNC’s complete set and gene groups [111]; gnomAD v4.1 constraint [110]; the ACMG secondary-findings list, version 3.3 [57]; TimeTree 5 divergence times [64]; little brown bat reads (BioProject PRJNA624023; Supplementary Note S30); Protein Data Bank entries 1JM7 and 1T29; and values printed in the published benchmark articles of Supplementary Notes S24 and S40.

Of the 7,394 samples in the FarmGTEx release [115] behind the label-free cattle panel, 160 carry no public accession (Additional file 2, reports/cattle_sample_table.tsv). The deposit carries every per-variant score and label analysed except licensed score values, withheld as set out below; ESM-2 [120] scores of ProteinGym mutants; the EVEE shard’s frozen scores; and sets recorded only as AUROCs or summaries (Supplementary Note S34 names each). The scoring windows are regenerated from the public assemblies by the window builders in Additional file 2. Two pieces of code are not deposited. One is the step that drew the atlas’s negatives from the Ensembl variation files, specified in Methods; Additional file 2’s pig builder (src/ccs/build_pig_real_neg.py) applies the same matching to PigGTEx genotypes. The other is the code of the two 40B passes behind the strand subpanel and the readout decomposition, whose scores and summary are deposited (Supplementary Notes S20 and S29). In the deposited atlas every pair of reference and alternate alleles carries ten negatives per positive in each non-human species but cat, whose 125 positives have 1,240 (9.92:1); how that shortfall arose, any seed of the Ensembl draws and the Hugging Face revisions of the Evo 2 checkpoints were not recorded.

Licensed score values are withheld. The academic branch of dbNSFP, which includes release 5.3.1a, is distributed under CC BY-NC-ND 4.0 [121], and several of its predictors carry their own non-redistribution terms, so Additional file 4 omits its score values. Additional file 5 likewise omits CADD values (under non-commercial terms), REVEL values (under the Open Database License 1.0 in its Zenodo record, though its download page states non-commercial use) and, so that the three rebuild and verify alike, AlphaMissense values. TraitGym’s column of CADD scores is withheld too, as are pCADD and chCADD scores, whose hosts state no licence; of AlphaMissense’s isoform release, under Creative Commons Attribution-NonCommercial-ShareAlike (CC BY-NC-SA) 4.0, only counts are deposited. Additional files 2 and 3 supply the builders that rebuild both panels from sources obtained under the reader’s own licence, and regenerate_full_panels.py (Additional file 3) checks a licensed rebuild against the deposited reach indicators and value digests. The CC0 release of Additional file 5’s reach indicators rests on the reading of the source licences that LICENSING.md sets out for the journal’s curators to confirm; should the reading not be accepted for REVEL, REVEL’s reach column will be released under the Open Database License 1.0 instead.

### Disclosure of use of AI-assisted tools including generative AI

Claude (Anthropic) [122] was used as an assistant for text editing and analysis code; all outputs were checked by the authors, who take full responsibility for the content. The models were Claude Opus and Claude Fable, used by Varunkumar Asediya from March to November 2026 [122].

### Authors’ contributions

V.A.: conceptualization, methodology, software, formal analysis, investigation, data curation, validation, visualization, writing – original draft.

U.GK.: formal analysis (supporting), writing – review and editing.

H.N.K.: conceptualization, resources, project administration, supervision, writing – review and editing.

C.S.P.: conceptualization, resources, project administration, supervision, writing – review and editing.

## Acknowledgements

We thank the depositors of the bat and cattle resequencing data and the authors of ESM-2 [120]. OMIA data were made available by software support from the Sydney Informatics Hub, funded from the Ronald Bruce Anstee bequest to the Sydney School of Veterinary Science for the Anstee Hub for Inherited Diseases in Animals (AHIDA). The species silhouettes, recoloured and resized, are from PhyloPic: chicken by Edwin Price, dog by Margot Michaud, pig by Gabriela Palomo-Munoz and cattle by Ivan Iofrida under CC BY 4.0, the other five in the public domain; Additional file 2 lists their contributors and licences (assets/silhouettes/credits.json).

## References

1. Kumar P, Henikoff S, Ng PC. Predicting the effects of coding non-synonymous variants on protein function using the SIFT algorithm. Nat Protoc. 2009;4(7):1073–1081. doi:10.1038/nprot.2009.86.

2. Adzhubei IA, Schmidt S, Peshkin L, et al. A method and server for predicting damaging missense mutations. Nat Methods. 2010;7(4):248–249. doi:10.1038/nmeth0410-248.

3. Jaganathan K, Kyriazopoulou Panagiotopoulou S, McRae JF, et al. Predicting splicing from primary sequence with deep learning. Cell. 2019;176(3):535–548.e24. doi:10.1016/j.cell.2018.12.015.

4. Radjasandirane R, Diharce J, Gelly JC, et al. Assessment of variant effect predictors unveils variants difficulty as a critical performance indicator. bioRxiv. 2024. doi:10.1101/2024.07.08.602580 (version 2; published as Insights for variant clinical interpretation based on a benchmark of 65 variant effect predictors, Genomics 2025;117(3):111036, doi:10.1016/j.ygeno.2025.111036).

5. Livesey BJ, Marsh JA. Variant effect predictor correlation with functional assays is reflective of clinical classification performance. Genome Biol. 2025;26(1):104. doi:10.1186/s13059-025-03575-w.

6. Notin P, Kollasch AW, Ritter D, et al. ProteinGym: large-scale benchmarks for protein fitness prediction and design. Adv Neural Inf Process Syst. 2023;36:64331–64379. doi:10.52202/075280-2810.

7. Rastogi R, Chung R, Li S, et al. Critical assessment of missense variant effect predictors on disease-relevant variant data. Hum Genet. 2025;144(2-3):281–293. doi:10.1007/s00439-025-02732-2.

8. dbNSFP Team. ClinVar benchmark for pathogenicity prediction scores in dbNSFP [blog post]. dbNSFP. 6 March 2026. https://www.dbnsfp.org/blog/clinvar-benchmark-for-pathogenicity-prediction-scores/ (accessed 12 November 2026).

9. Brandes N, Goldman G, Wang CH, et al. Genome-wide prediction of disease variant effects with a deep protein language model. Nat Genet. 2023;55(9):1512–1522. doi:10.1038/s41588-023-01465-0.

10. Radivojac P, Clark WT, Oron TR, et al. A large-scale evaluation of computational protein function prediction. Nat Methods. 2013;10(3):221–227. doi:10.1038/nmeth.2340.

11. Jiang Y, Oron TR, Clark WT, et al. An expanded evaluation of protein function prediction methods shows an improvement in accuracy. Genome Biol. 2016;17(1):184. doi:10.1186/s13059-016-1037-6.

12. Bossuyt PM, Reitsma JB, Bruns DE, et al. STARD 2015: an updated list of essential items for reporting diagnostic accuracy studies. BMJ. 2015;351:h5527. doi:10.1136/bmj.h5527.

13. Shinkins B, Thompson M, Mallett S, et al. Diagnostic accuracy studies: how to report and analyse inconclusive test results. BMJ. 2013;346:f2778. doi:10.1136/bmj.f2778.

14. Grimm DG, Azencott CA, Aicheler F, et al. The evaluation of tools used to predict the impact of missense variants is hindered by two types of circularity. Hum Mutat. 2015;36(5):513–523. doi:10.1002/humu.22768.

15. Livesey BJ, Marsh JA. Using deep mutational scanning to benchmark variant effect predictors and identify disease mutations. Mol Syst Biol. 2020;16(7):e9380. doi:10.15252/msb.20199380.

16. Lu B, Liu X, Lin PY, et al. Genomic heterogeneity inflates the performance of variant pathogenicity predictions. Nat Commun. 2026. doi:10.1038/s41467-026-78623-5.

17. Harrizi S, Nait Irahal I, Mostafa K, et al. Gene identity, not variant effect, dominates ClinVar benchmarks of missense pathogenicity predictors. bioRxiv. 2026. doi:10.64898/2026.08.21.746180.

18. Pejaver V, Byrne AB, Feng BJ, et al. Calibration of computational tools for missense variant pathogenicity classification and ClinGen recommendations for PP3/BP4 criteria. Am J Hum Genet. 2022;109(12):2163–2177. doi:10.1016/j.ajhg.2022.10.013.

19. Bergquist T, Stenton SL, Nadeau EAW, et al. Calibration of additional computational tools expands ClinGen recommendation options for variant classification with PP3/BP4 criteria. Genet Med. 2025;27(6):101402. doi:10.1016/j.gim.2025.101402.

20. Frazer J, Notin P, Dias M, et al. Disease variant prediction with deep generative models of evolutionary data. Nature. 2021;599(7883):91–95. doi:10.1038/s41586-021-04043-8.

21. Choe YJ, Gangrade A, Ramdas A. Counterfactually comparing abstaining classifiers. Adv Neural Inf Process Syst. 2023;36:28281-28293. doi:10.52202/075280-1229.

22. Kalai AT, Nachum O, Vempala SS, et al. Evaluating large language models for accuracy incentivizes hallucinations. Nature. 2026;653(8116):1047–1051. doi:10.1038/s41586-026-10549-w.

23. Fernández-Barrios R, Pastor-López I, González-Santocildes A, et al. When disagreement labels do not identify F1 rankings. arXiv. 2026. doi:10.48550/arXiv.2611.01712.

24. Manski CF. Partial identification of probability distributions. New York: Springer; 2003. doi:10.1007/b97478.

25. Horowitz JL, Manski CF. Identification and robustness with contaminated and corrupted data. Econometrica. 1995;63(2):281–302. doi:10.2307/2951627.

26. Li Y, Engelhard MM. Extending ROC analysis to uncertainty-aware risk prediction with an interval-based AUC (iAUC). arXiv. 2026. doi:10.48550/arXiv.2602.04775.

27. Nguyen E, Poli M, Durrant MG, et al. Sequence modeling and design from molecular to genome scale with Evo. Science. 2024;386(6723):eado9336. doi:10.1126/science.ado9336.

28. Benegas G, Batra SS, Song YS. DNA language models are powerful predictors of genome-wide variant effects. Proc Natl Acad Sci U S A. 2023;120(44):e2311219120. doi:10.1073/pnas.2311219120.

29. Benegas G, Albors C, Aw AJ, et al. A DNA language model based on multispecies alignment predicts the effects of genome-wide variants. Nat Biotechnol. 2025;43(12):1960–1965. doi:10.1038/s41587-024-02511-w.

30. Ye C, Benegas G, Albors C, et al. Predicting genome-wide functional constraints with GPN-Star. Nature. 2026;659(8139):141–150. doi:10.1038/s41586-026-11005-5.

31. Brixi G, Durrant MG, Ku J, et al. Genome modelling and design across all domains of life with Evo 2. Nature. 2026;652(8112):1349–1361. doi:10.1038/s41586-026-10176-5.

32. Tammen I, Mather M, Leeb T, et al. Online Mendelian Inheritance in Animals (OMIA): a genetic resource for vertebrate animals. Mamm Genome. 2024;35(4):556–564. doi:10.1007/s00335-024-10059-y.

33. Nicholas FW, Tammen I, Sydney Informatics Hub. Online Mendelian Inheritance in Animals (OMIA) [dataset]. 1995. doi:10.25910/2amr-pv70.

34. Jiang J, Tiezzi F, Maltecca C. Cross-species functional variant classification using the Evo 2 genomic language model: implications for farm animal genetics and breeding. Research Square. 2025. doi:10.21203/rs.3.rs-7579108/v1.

35. Landrum MJ, Lee JM, Benson M, et al. ClinVar: improving access to variant interpretations and supporting evidence. Nucleic Acids Res. 2018;46(D1):D1062–D1067. doi:10.1093/nar/gkx1153.

36. Asediya V, GK U, Kadarmideen HN, et al. Unscored variants leave many comparisons of variant effect predictors undecided: evidence from ClinVar, published benchmarks and genomic language models: code and data. Version 0.1.1. Zenodo. 2026. doi:10.5281/zenodo.22046811.

37. Pollard KS, Hubisz MJ, Rosenbloom KR, et al. Detection of nonneutral substitution rates on mammalian phylogenies. Genome Res. 2010;20(1):110–121. doi:10.1101/gr.097857.109.

38. Siepel A, Bejerano G, Pedersen JS, et al. Evolutionarily conserved elements in vertebrate, insect, worm, and yeast genomes. Genome Res. 2005;15(8):1034–1050. doi:10.1101/gr.3715005.

39. Schubach M, Maass T, Nazaretyan L, et al. CADD v1.7: using protein language models, regulatory CNNs and other nucleotide-level scores to improve genome-wide variant predictions. Nucleic Acids Res. 2024;52(D1):D1143–D1154. doi:10.1093/nar/gkad989.

40. Ioannidis NM, Rothstein JH, Pejaver V, et al. REVEL: an ensemble method for predicting the pathogenicity of rare missense variants. Am J Hum Genet. 2016;99(4):877–885. doi:10.1016/j.ajhg.2016.08.016.

41. Cheng J, Novati G, Pan J, et al. Accurate proteome-wide missense variant effect prediction with AlphaMissense. Science. 2023;381(6664):eadg7492. doi:10.1126/science.adg7492.

42. Cooper GM, Stone EA, Asimenos G, et al. Distribution and intensity of constraint in mammalian genomic sequence. Genome Res. 2005;15(7):901–913. doi:10.1101/gr.3577405.

43. Davydov EV, Goode DL, Sirota M, et al. Identifying a high fraction of the human genome to be under selective constraint using GERP++. PLoS Comput Biol. 2010;6(12):e1001025. doi:10.1371/journal.pcbi.1001025.

44. Zoonomia Consortium. A comparative genomics multitool for scientific discovery and conservation. Nature. 2020;587(7833):240–245. doi:10.1038/s41586-020-2876-6.

45. Christmas MJ, Kaplow IM, Genereux DP, et al. Evolutionary constraint and innovation across hundreds of placental mammals. Science. 2023;380(6643):eabn3943. doi:10.1126/science.abn3943.

46. Preston CG, Wright MW, Madhavrao R, et al. ClinGen Variant Curation Interface: a variant classification platform for the application of evidence criteria from ACMG/AMP guidelines. Genome Med. 2022;14(1):6. doi:10.1186/s13073-021-01004-8.

47. Benegas G, Eraslan G, Song YS. Benchmarking DNA sequence models for causal regulatory variant prediction in human genetics. bioRxiv. 2025. doi:10.1101/2025.02.11.637758.

48. Wei J, Nguyen K, Chung HW, et al. Measuring short-form factuality in large language models. arXiv. 2024. doi:10.48550/arXiv.2411.04368.

49. dbNSFP Team. Interpreting “missing” prediction scores across dbNSFP versions [blog post]. dbNSFP. 19 August 2026. https://www.dbnsfp.org/blog/missing-prediction-scores-after-upgrade/ (accessed 12 November 2026).

50. Imbens GW, Manski CF. Confidence intervals for partially identified parameters. Econometrica. 2004;72(6):1845–1857. doi:10.1111/j.1468-0262.2004.00555.x.

51. Stoye J. More on confidence intervals for partially identified parameters. Econometrica. 2009;77(4):1299–1315. doi:10.3982/ECTA7347.

52. Critical Assessment of Genome Interpretation Consortium. CAGI, the Critical Assessment of Genome Interpretation, establishes progress and prospects for computational genetic variant interpretation methods. Genome Biol. 2024;25(1):53. doi:10.1186/s13059-023-03113-6.

53. Tejura M, Fayer S, McEwen AE, et al. Calibration of variant effect predictors on genome-wide data masks heterogeneous performance across genes. Am J Hum Genet. 2024;111(9):2031–2043. doi:10.1016/j.ajhg.2024.07.018.

54. Masten MA, Poirier A. Inference on breakdown frontiers. Quant Econ. 2020;11(1):41–111. doi:10.3982/QE1288.

55. Richards S, Aziz N, Bale S, et al. Standards and guidelines for the interpretation of sequence variants: a joint consensus recommendation of the American College of Medical Genetics and Genomics and the Association for Molecular Pathology. Genet Med. 2015;17(5):405–424. doi:10.1038/gim.2015.30.

56. Wilcox EH, Sarmady M, Wulf B, et al. Evaluating the impact of in silico predictors on clinical variant classification. Genet Med. 2022;24(4):924–930. doi:10.1016/j.gim.2021.11.018.

57. Lee K, Abul-Husn NS, Amendola LM, et al. ACMG SF v3.3 list for reporting of secondary findings in clinical exome and genome sequencing: a policy statement of the American College of Medical Genetics and Genomics (ACMG). Genet Med. 2025;27(8):101454. doi:10.1016/j.gim.2025.101454.

58. Smith C, Kitzman JO. Benchmarking splice variant prediction algorithms using massively parallel splicing assays. Genome Biol. 2023;24(1):294. doi:10.1186/s13059-023-03144-z.

59. Walker LC, de la Hoya M, Wiggins GAR, et al. Using the ACMG/AMP framework to capture evidence related to predicted and observed impact on splicing: recommendations from the ClinGen SVI Splicing Subgroup. Am J Hum Genet. 2023;110(7):1046–1067. doi:10.1016/j.ajhg.2023.06.002.

60. Cormier MJ, Pedersen BS, Bayrak-Toydemir P, et al. Combining genetic constraint with predictions of alternative splicing to prioritize deleterious splicing in rare disease studies. BMC Bioinformatics. 2022;23(1):482. doi:10.1186/s12859-022-05041-x.

61. Jagadeesh KA, Paggi JM, Ye JS, et al. S-CAP extends pathogenicity prediction to genetic variants that affect RNA splicing. Nat Genet. 2019;51(4):755–763. doi:10.1038/s41588-019-0348-4.

62. Martin FJ, Amode MR, Aneja A, et al. Ensembl 2023. Nucleic Acids Res. 2023;51(D1):D933–D941. doi:10.1093/nar/gkac958.

63. Teng J, Gao Y, Yin H, et al. A compendium of genetic regulatory effects across pig tissues. Nat Genet. 2024;56(1):112–123. doi:10.1038/s41588-023-01585-7.

64. Kumar S, Suleski M, Craig JM, et al. TimeTree 5: an expanded resource for species divergence times. Mol Biol Evol. 2022;39(8):msac174. doi:10.1093/molbev/msac174.

65. Cingolani P, Platts A, Wang LL, et al. A program for annotating and predicting the effects of single nucleotide polymorphisms, SnpEff: SNPs in the genome of Drosophila melanogaster strain w1118; iso-2; iso-3. Fly (Austin). 2012;6(2):80-92. doi:10.4161/fly.19695.

66. Groß C, Derks M, Megens HJ, et al. pCADD: SNV prioritisation in *Sus scrofa*. Genet Sel Evol. 2020;52(1):4. doi:10.1186/s12711-020-0528-9.

67. Groß C, Bortoluzzi C, de Ridder D, et al. Prioritizing sequence variants in conserved non-coding elements in the chicken genome using chCADD. PLoS Genet. 2020;16(9):e1009027. doi:10.1371/journal.pgen.1009027.

68. Reeb J, Hecht M, Mahlich Y, et al. Predicted molecular effects of sequence variants link to system level of disease. PLoS Comput Biol. 2016;12(8):e1005047. doi:10.1371/journal.pcbi.1005047.

69. Hu H, Xie W. Human genomic tools to assess missense variant pathogenicity in domestic mammals using a mock-variant strategy. Open Vet J. 2025;15(10):5183–5191. doi:10.5455/OVJ.2025.v15.i10.35.

70. McLaren W, Gil L, Hunt SE, et al. The Ensembl Variant Effect Predictor. Genome Biol. 2016;17(1):122. doi:10.1186/s13059-016-0974-4.

71. Findlay GM, Daza RM, Martin B, et al. Accurate classification of BRCA1 variants with saturation genome editing. Nature. 2018;562(7726):217–222. doi:10.1038/s41586-018-0461-z.

72. Barbosa P, Ribeiro M, Carmo-Fonseca M, et al. Clinical significance of genetic variation in hypertrophic cardiomyopathy: comparison of computational tools to prioritize missense variants. Front Cardiovasc Med. 2022;9:975478. doi:10.3389/fcvm.2022.975478.

73. Heo JY, Kim JH. Assessing the performance of 28 pathogenicity prediction methods on rare single nucleotide variants in coding regions. BMC Genomics. 2025;26(1):641. doi:10.1186/s12864-025-11787-4.

74. Ravikumar S. The missing “I don’t know”: why three reasoning-reliability findings converge on calibrated abstention. In: Proceedings of the 15th International Joint Conference on Natural Language Processing and the 5th Conference of the Asia-Pacific Chapter of the Association for Computational Linguistics (Volume 2: Short Papers). Association for Computational Linguistics; 2026. p. 513–524. doi:10.18653/2026.aacl-short.34.

75. Meier J, Rao R, Verkuil R, et al. Language models enable zero-shot prediction of the effects of mutations on protein function. Adv Neural Inf Process Syst. 2021;34:29287–29303.

76. Alfisi I, Ciapi F, Baragli M, et al. Benchmarking DNA foundation models for zero-shot variant effect prediction shows the importance of context, training, and architecture. Genome Biol. 2026;27(1):354. doi:10.1186/s13059-026-04238-0.

77. Mathur V, Sachidanandam R. Benchmarking DNA foundation models: biological blind spots in Evo2 variant-effect prediction. bioRxiv. 2026. doi:10.64898/2026.03.10.710786.

78. Dalla-Torre H, Gonzalez L, Mendoza-Revilla J, et al. Nucleotide Transformer: building and evaluating robust foundation models for human genomics. Nat Methods. 2025;22(2):287–297. doi:10.1038/s41592-024-02523-z.

79. Tang Z, Somia N, Yu Y, et al. Evaluating the representational power of pre-trained DNA language models for regulatory genomics. Genome Biol. 2025;26(1):203. doi:10.1186/s13059-025-03674-8.

80. Nguyen DT, Knutsen TM, Sandve SR, et al. Sequence-based chromatin activity modeling and regulatory impact prediction of genetic variants in farmed animals using deep learning. NAR Genom Bioinform. 2025;7(4):lqaf139. doi:10.1093/nargab/lqaf139.

81. Avsec Ž, Latysheva N, Cheng J, et al. Advancing regulatory variant effect prediction with AlphaGenome. Nature. 2026;649(8099):1206–1218. doi:10.1038/s41586-025-10014-0.

82. Livesey BJ, Badonyi M, Dias M, et al. Guidelines for releasing a variant effect predictor. Genome Biol. 2025;26(1):97. doi:10.1186/s13059-025-03572-z.

83. Boeykens F, Abitbol M, Anderson H, et al. Development and validation of animal variant classification guidelines to objectively evaluate genetic variant pathogenicity in domestic animals. Front Vet Sci. 2024;11:1497817. doi:10.3389/fvets.2024.1497817.

84. Desmet L, Ferrari C, Abitbol M, et al. The Animal Variant Classification Guidelines v2: an update with new criteria and improved clarifications. Anim Genet. 2026;57(5):e70197. doi:10.1002/age.70197.

85. Warr A, Affara N, Aken B, et al. An improved pig reference genome sequence to enable pig genetics and genomics research. GigaScience. 2020;9(6):giaa051. doi:10.1093/gigascience/giaa051.

86. Loay H, Kar P, Koch E, et al. Mutation rate heterogeneity biases variant effect prediction and reveals genuine mutational robustness. Am J Hum Genet. 2026;113(10):2042–2054. doi:10.1016/j.ajhg.2026.08.011.

87. Harrison SM, Rehm HL. Is ‘likely pathogenic’ really 90% likely? Reclassification data in ClinVar. Genome Med. 2019;11(1):72. doi:10.1186/s13073-019-0688-9.

88. Cui R, Elzur RA, Kanai M, et al. Improving fine-mapping by modeling infinitesimal effects. Nat Genet. 2024;56(1):162–169. doi:10.1038/s41588-023-01597-3.

89. Wang G, Sarkar A, Carbonetto P, et al. A simple new approach to variable selection in regression, with application to genetic fine mapping. J R Stat Soc Series B Stat Methodol. 2020;82(5):1273–1300. doi:10.1111/rssb.12388.

90. Zou Y, Carbonetto P, Wang G, et al. Fine-mapping from summary data with the “Sum of Single Effects” model. PLoS Genet. 2022;18(7):e1010299. doi:10.1371/journal.pgen.1010299.

91. Poli M, Massaroli S, Nguyen E, et al. Hyena Hierarchy: towards larger convolutional language models. In: Proceedings of the 40th International Conference on Machine Learning (ICML). Proc Mach Learn Res. 2023;202:28043-28078.

92. Liu X, Li C, Mou C, et al. dbNSFP v4: a comprehensive database of transcript-specific functional predictions and annotations for human nonsynonymous and splice-site SNVs. Genome Med. 2020;12(1):103. doi:10.1186/s13073-020-00803-9.

93. Perez G, Barber GP, Benet-Pages A, et al. The UCSC Genome Browser database: 2025 update. Nucleic Acids Res. 2025;53(D1):D1243–D1249. doi:10.1093/nar/gkae974.

94. Clawson H, Lee BT, Raney BJ, et al. GenArk: towards a million UCSC genome browsers. Genome Biol. 2023;24(1):217. doi:10.1186/s13059-023-03057-x.

95. Zeng T, Li YI. Predicting RNA splicing from DNA sequence using Pangolin. Genome Biol. 2022;23(1):103. doi:10.1186/s13059-022-02664-4.

96. Danis D, Jacobsen JOB, Carmody LC, et al. Interpretable prioritization of splice variants in diagnostic next-generation sequencing. Am J Hum Genet. 2021;108(9):1564–1577. doi:10.1016/j.ajhg.2021.06.014.

97. Xiong HY, Alipanahi B, Lee LJ, et al. The human splicing code reveals new insights into the genetic determinants of disease. Science. 2015;347(6218):1254806. doi:10.1126/science.1254806.

98. Rosenberg AB, Patwardhan RP, Shendure J, et al. Learning the sequence determinants of alternative splicing from millions of random sequences. Cell. 2015;163(3):698–711. doi:10.1016/j.cell.2015.09.054.

99. Cheng J, Nguyen TYD, Cygan KJ, et al. MMSplice: modular modeling improves the predictions of genetic variant effects on splicing. Genome Biol. 2019;20(1):48. doi:10.1186/s13059-019-1653-z.

100. Schuetz GM, Schlattmann P, Dewey M. Use of 3×2 tables with an intention to diagnose approach to assess clinical performance of diagnostic tests: meta-analytical evaluation of coronary CT angiography studies. BMJ. 2012;345:e6717. doi:10.1136/bmj.e6717.

101. Holm S. A simple sequentially rejective multiple test procedure. Scand J Stat. 1979;6(2):65–70.

102. Tavtigian SV, Harrison SM, Boucher KM, et al. Fitting a naturally scaled point system to the ACMG/AMP variant classification guidelines. Hum Mutat. 2020;41(10):1734–1737. doi:10.1002/humu.24088.

103. Pedregosa F, Varoquaux G, Gramfort A, et al. Scikit-learn: machine learning in Python. J Mach Learn Res. 2011;12:2825–2830.

104. Westfall PH, Young SS. Resampling-based multiple testing: examples and methods for p-value adjustment. New York: Wiley; 1993. ISBN 978-0-471-55761-6.

105. Benjamini Y, Hochberg Y. Controlling the false discovery rate: a practical and powerful approach to multiple testing. J R Stat Soc Series B Methodol. 1995;57(1):289–300. doi:10.1111/j.2517-6161.1995.tb02031.x.

106. Ibragimov R, Müller UK. t-Statistic based correlation and heterogeneity robust inference. J Bus Econ Stat. 2010;28(4):453–468. doi:10.1198/jbes.2009.08046.

107. DeLong ER, DeLong DM, Clarke-Pearson DL. Comparing the areas under two or more correlated receiver operating characteristic curves: a nonparametric approach. Biometrics. 1988;44(3):837–845. doi:10.2307/2531595.

108. Sun X, Xu W. Fast implementation of DeLong’s algorithm for comparing the areas under correlated receiver operating characteristic curves. IEEE Signal Process Lett. 2014;21(11):1389–1393. doi:10.1109/LSP.2014.2337313.

109. Morales J, Pujar S, Loveland JE, et al. A joint NCBI and EMBL-EBI transcript set for clinical genomics and research. Nature. 2022;604(7905):310–315. doi:10.1038/s41586-022-04558-8.

110. Karczewski KJ, Francioli LC, Tiao G, et al. The mutational constraint spectrum quantified from variation in 141,456 humans. Nature. 2020;581(7809):434–443. doi:10.1038/s41586-020-2308-7.

111. Seal RL, Braschi B, Gray K, et al. Genenames.org: the HGNC resources in 2023. Nucleic Acids Res. 2023;51(D1):D1003–D1009. doi:10.1093/nar/gkac888.

112. Harris CR, Millman KJ, van der Walt SJ, et al. Array programming with NumPy. Nature. 2020;585(7825):357–362. doi:10.1038/s41586-020-2649-2.

113. Chang CC, Chow CC, Tellier LC, et al. Second-generation PLINK: rising to the challenge of larger and richer datasets. GigaScience. 2015;4:7. doi:10.1186/s13742-015-0047-8.

114. Zhao R, Owen R, Marr M, et al. The potential of AI models to identify regulatory variants underlying cattle traits. bioRxiv. 2025. doi:10.1101/2024.08.01.606140.

115. Liu S, Gao Y, Canela-Xandri O, et al. A multi-tissue atlas of regulatory variants in cattle. Nat Genet. 2022;54(9):1438–1447. doi:10.1038/s41588-022-01153-5.

116. UniProt Consortium. UniProt: the Universal Protein Knowledgebase in 2025. Nucleic Acids Res. 2025;53(D1):D609–D617. doi:10.1093/nar/gkae1010.

117. Pearce MT, Dooms T, Yamamoto R, et al. EVEE: interpretable variant effect prediction from genomic foundation model embeddings. Zenodo. 2026. doi:10.5281/zenodo.19701997.

118. Tabet DR, Coté AG, Lancaster MC, et al. The functional landscape of coding variation in the familial hypercholesterolemia gene LDLR. Science. 2026;391(6787):eady7186. doi:10.1126/science.ady7186.

119. Esposito D, Weile J, Shendure J, et al. MaveDB: an open-source platform to distribute and interpret data from multiplexed assays of variant effect. Genome Biol. 2019;20(1):223. doi:10.1186/s13059-019-1845-6.

120. Lin Z, Akin H, Rao R, et al. Evolutionary-scale prediction of atomic-level protein structure with a language model. Science. 2023;379(6637):1123–1130. doi:10.1126/science.ade2574.

121. Genos Bioinformatics LLC. dbNSFP license [web page]. dbNSFP. 2026. https://www.dbnsfp.org/license (accessed 12 November 2026).

122. Anthropic. Claude (Claude Opus and Claude Fable) [Large language model]. Used by Varunkumar Asediya, March–November 2026.

