## Additional_file_1 for "Unscored variants leave many comparisons of variant effect predictors undecided: evidence from ClinVar, published benchmarks and genomic language models"

### Additional file 1: Supplementary Information

Reference numbering follows the main manuscript.

Notes S2, S3, S9, S19, S20, S22, S28, S32 and S33 hold extended Methods.

### Contents

**Supplementary Notes**

Note S1. Relation to prior cross-species work, and the overlap we assume rather than claim

Note S2. Deployment reach of eight splice predictors, the ClinVar consequence panel and gene-structured missingness

Note S3. dbNSFP panel: gene-resampled intervals, confidence sets for the bounds and family-wise counts

Note S4. Judging each comparison inside the predictors' declared scopes, and covered AUROCs printed side by side

Note S5. Identification: completions, transcript join, within-gene, rare-variant and label-age accounting, and the missingness indicator

Note S6. Comparing predictors within gene, what that changes, and the accounting inside each gene

Note S7. How far each comparison survives: breakdown points and a dominance map

Note S8. Calibrated clinical evidence, label circularity and functional labels

Note S9. The fitted components against the DOME recommendations

Note S10. What Evo 2 adds to annotation and conservation, and the capacity ladder

Note S11. The 8,192-bp readout: AUPRC, the cross-species share of ranked pairs, and the readout effect

Note S12. Fixed false-positive operating points, and Table 1's intervals: locus clustering and the macro row's t interval

Note S13. The conservation-matched control, and the residual it leaves

Note S14. What the atlas measures: composition controls, within-class pairs and the missense stratum by species

Note S15. A human missense predictor carried to the atlas through the orthologue

Note S16. GPN-Star, a second genomic language model, on the chicken and human atlas variants, and SIFT's reach on the atlas

Note S17. The reference-likelihood confound across three panels

Note S18. The BRCA1 positive control, and the start-codon failure mode it exposes

Note S19. Scoring provenance: checkpoints, scorers, window extraction, reference checks and score orientation

Note S20. Extended Methods: comparators and statistical machinery

Note S21. Evo 2 against conservation at both readouts: per-species margins and scope

Note S22. Resampling units, permutation schemes, seeds, equivalence testing and the fragility of atlas verdicts

Note S23. The comparison is not measured on the same variants: reach, its cause and identification

Note S24. Orderings stated in published benchmarks: a systematic search

Note S25. The pig cis-eQTL panel and its near-chance result

Note S26. At the readouts tested, competence stops at a variant class, not at a genomic region

Note S27. Readouts that reorder models on an independent benchmark

Note S28. Calibration, abstention and conformal prediction across species

Note S29. Strand consistency at the 1B and 40B checkpoints

Note S30. The label-free cattle analysis conditioned on conservation, and the cross-clade bat pilot

Note S31. Further limitations

Note S32. Panel provenance: sources, construction and exclusions of every panel

Note S33. Specialist and supervised baselines, the embedding probe and a proximity control

Note S34. The regeneration scope and reproducibility status of the deposited archives

Note S35. Where each reported value is recomputed

Note S36. A general statement of the bound, the decision rule and the breakdown point

Note S37. What the conventions for unscored variants assume: least-deleterious fill, shared variants and the difficulty of declined variants

Note S38. Variants of uncertain significance: unscored, indeterminate and outside scope

Note S39. Which genes, variant classes and disease categories go unscored

Note S40. Named orderings and orderings printed in opposite directions

Note S41. A census of public genomic language model benchmarks

Note S42. The accounting on a language-model factuality benchmark

Note S43. Stronger conservation comparators: 241-mammal phyloP, pCADD and chCADD

Note S44. The consequence-matched arm

**Supplementary Figures**

**Supplementary Tables**

**References**

### Supplementary Notes

#### Note S1. Relation to prior cross-species work, and the overlap we assume rather than claim

Prior work [34] scored 721 OMIA SNVs in our eight non-human species with Evo 2-7B, by zero-shot log-likelihood in 8,192-bp windows, our headline readout, and showed that the separation largely tracks variant type. It reports an AUROC of 0.934 against variant-type-blind controls and 0.717 against variant-type-matched controls, and an area under the precision–recall curve (AUPRC) of 0.832 blind falling to 0.166 matched. With Evo 2-40B we read 0.939 by the species mean over the same eight species (Table 1) and 0.750 on the 533 non-human atlas missense variants it scores (Note S15). Checkpoint- and species-matched, our Evo 2-7B scores at 8,192 bp read 0.937 pooled and 0.924 by the species mean, against their 0.934 and 0.909. The matched figures, 0.750 against 0.717, come from designs that define matching differently and use different checkpoints, so their difference is not evidence of a model discrepancy. That study reports Evo 2 without a conservation comparator, so a comparator's silence does not arise there. In that study, a probe trained on other species' labels adds nothing to Evo 2's zero-shot score against unmatched population controls and 0.02 to 0.05 AUROC by the species mean against consequence-matched ones, depending on whether cat is counted [34]. To its result the atlas adds the 40B checkpoint, a conservation comparator charged for its silence, the readout contrast on the same variants, a human ClinVar arm and deposited per-variant scores and labels. Their positives, controls and per-variant scores are not public, so the comparison above uses their published summary figures. Given a common source database, OMIA, and an identical species set, the variant-level overlap between their 721 and our 740 positives is likely substantial, and we assume so rather than claim independence.

#### Note S2. Deployment reach of eight splice predictors, the ClinVar consequence panel and gene-structured missingness

**Source and provenance.** Five of the splicing panel's six arms [58] are saturation-mutagenesis assays. The sixth is 296 MLH1 variants, 7.6% of the panel, which Smith and Kitzman curated from clinical variant databases and literature reports rather than assaying de novo. The six datasets carrying a binary label (Table S13) give 3,912 variants, of which 1,060 (27.1%) are disruptive, 82 of them MLH1 essential-splice calls admitted by rule. The 1,386-variant BRCA1 arm is the intronic and synonymous slice of the saturation-genome-editing set of Findlay et al. [71] that supplies this paper's positive control, on the same build. No independence between the two is claimed. Nothing is fitted on either. Dropping the BRCA1 arm, which Evo 2 reads best, moves its n-weighted mean deficit against SpliceAI (Table S13) from −0.2626 to −0.3092: the overlap narrows the deficit. With eight negatives among 246 variants, the Essential Splice class (Table S20) is set against the other four classes only descriptively; that contrast is not tested. Smith and Kitzman's 500,000 background SNVs lie in the internal protein-coding exons, and their 100-bp flanks, of 14,577 genes; they carry no labels and serve only for reach, and our call rates on them reproduce the published ones within 0.054 percentage points [58].

**The ClinVar consequence panel.** Drawn from the reach panel's ClinVar release without a review-status filter, it holds 3,714 variants across 16 cells, balanced by consequence and label with at most 250 per cell. Its regulatory consequence classes carry 54 to 250 positives, against the 18 available in the OMIA consequence gradient of Note S26.

**Composition control on the reach comparison.** The benchmark and the background do not share a variant-class mix, and reach depends on class, so a raw comparison confounds curation with composition. We standardised the background to the benchmark's class mix by reweighting per-class reach (Tables S14 and S28). Standardising moves the benchmark-to-background gap by 0.0002 for SpliceAI, ConSpliceML and SQUIRLS (SQUIRLS 0.0666 raw, 0.0664 standardised), so composition explains almost none of it for these three. For S-CAP the raw gap of 0.251 falls to 0.175, so composition accounts for about 30% and the remainder survives.

Standardised reach on the background differs from benchmark reach in both directions (Tables S14 and S28; Figure S14a). SQUIRLS scores all benchmark negatives but only 0.899 of positives, a class gap of −0.101 in an apparently complete panel. For SpliceAI, ConSpliceML and SQUIRLS the missingness is organised by gene: the annotation a run consults, not the variants a user brings, decides which genes it answers (Figure S14b).

With the unscored variants bounded at their extremes, HAL's AUROC on the whole benchmark lies in [0.135, 0.947] and S-CAP's bound also contains 0.5, so neither is identified above chance, although their covered AUROCs, 0.717 and 0.650, lie above it (Figure S14d). Pooled over the benchmark, the reach of HAL and S-CAP differs by only +0.034 and −0.040 between disruptive and non-disruptive variants, but within a dataset it can differ sharply, as S-CAP's does on MLH1.

Smith and Kitzman's per-dataset comparison credits the tool with the top precision–recall area, and any tool within its 95% interval, each on the variants it answers [58]. On their MLH1 dataset S-CAP wins at 0.988 on the 138 variants it scores, 84.1% of them disruptive against 54.1% of the dataset. On those same variants four tools read higher: Pangolin at 0.999, SpliceAI and ConSpliceML at 0.998 and MMSplice at 0.995. S-CAP's must-answer AUROC there is 0.551, the lowest of the eight, and its bound [0.109, 0.992] overlaps every other tool's, so by AUROC its printed win is not identified. HAL's win on RON exon 11 is not identified either (bound [0.274, 0.891]), though HAL leads on the 378 of 598 variants it scores. Smith and Kitzman's headline, that SpliceAI and Pangolin perform best overall, survives: they remain the most frequent winners under every convention, although their lead over HAL and S-CAP is not identified in most datasets.

##### *Gene-structured missingness and the union-ensemble limit*

Among the 3,029 background genes carrying at least 50 variants, SpliceAI's per-gene reach is 1.000 at both the tenth and the ninetieth percentile and zero for 100 genes (Figure S14b). SQUIRLS has 296 zero-reach genes and ConSpliceML 78. On the same 3,029 genes, S-CAP's silence is organised by neither gene nor variant class, which explain, respectively, 0.017 and 0.151 of the variance in its answered-or-declined indicator (Figure S14c).

Of the 500,000 variants, 7.9% are scored by all eight predictors and the mean variant is scored by 6.63 of them (Figure S14a). For essential-splice, intron-near-junction and proximal-intron variants the proportion scored by all eight is zero.

Pairwise Jaccard overlap between zero-reach gene sets is at most 0.134 (SpliceAI against ConSpliceML). Across the eight, 432 genes are invisible to at least one predictor and none is invisible to every predictor that has holes at all. A union across the five predictors that have zero-reach genes (Figure S14b) would therefore repair gene-level reach almost completely.

The individual tools do not implement that union, and combining them would require a prespecified aggregation rule and renewed evaluation of class-dependent selection.

#### Note S3. dbNSFP panel: gene-resampled intervals, confidence sets for the bounds and family-wise counts

Variants within a gene share sequence context and annotation history, so intervals on this panel resample genes. In DeLong's method [107,108] an unreachable variant has a placement value of exactly one half, so the same covariance gives a closed-form interval on a must-answer difference. The same placements give confidence sets for the bounds. Each bound endpoint is a Mann–Whitney statistic in which a pair touching an unscored variant counts 0 in the lower bound and 1 in the upper, so summing its placements within genes gives a gene-clustered variance for every endpoint and every difference between two endpoints. A comparison counts as identified at 95% when its facing endpoints are more than c standard errors apart, c being chosen to cover a partially identified difference with 95% probability [50,51]; it lies between 1.645 and 1.960, reaching at most 1.958 over the identified comparisons. The sets are per-comparison, not simultaneous over the 1,176 comparisons. They keep 74 of the 83 and 376 of the 429 missense-subset comparisons; treating variants as independent would keep all 83 and 418. The nine that drop out all clear the bar by less than 0.012 AUROC, and for the bound endpoints the gene-clustered standard errors are a median 6.2 times the variant-level ones. The sets assume that no single gene carries a non-vanishing share of the variance. On the missense subset that fails for the reach component of the endpoints: over the 48 predictors that leave a variant unscored, the variance of pair coverage across genes rests on a median of 7.8 effective genes ((Σψ²)²/Σψ⁴, the inverse Herfindahl index of the genes' shares of the clustered variance, ψ being a gene's summed influence, from Additional file 4's reach indicators), 41 of the 48 below 20, and TTN carries 82% of AlphaMissense's, DNAH11 46% of REVEL's and FBN1 62% of each VARITY score's. Cluster-robust intervals undercover in that regime; how far the sets undercover here, scores included, is measured at the end of this Note.

Holm's step-down procedure on the p-values dual to these sets, which holds the family-wise error rate over the 1,176 comparisons at 5%, keeps 67 of the 83 and 316 of the 429, and a step-down max-T bootstrap keeps 68 and 323. Of the 83, 16 involve bStatistic, a regional background-selection statistic that reads near chance (covered AUROC 0.5491), and all 16 survive each standard; without it 83 falls to 67, 74 to 58 at 95% and 67 to 51 under Holm's procedure, and on the missense subset 429 falls to 382, 376 to 330 and 316 to 271. In 8 of the 47 identified comparisons between a conservation track and another predictor, the other predictor is unsupervised.

How many genes carry a confidence set was measured by the effective number of genes defined above, applied to each endpoint's clustered variance. Lower ends rest on a median of 24.7 effective genes on the whole panel (quartiles 20.8 to 27.0) and 14.5 on the missense subset (6.7 to 29.2), where 21 of the 49 rest on fewer than 10 and AlphaMissense's on 1.5, 80% in TTN. Removing FBN1, TTN, DNAH11, BRCA2 and SCN1A, the five genes carrying most endpoint variance, and recomputing bounds and sets drops 2 of the 74 and 7 of the 377 that stay identified at 95% inside declared scopes (Note S4). With the panel as the population, over 500 gene resamples, the one-sided 95% sets covered 95.1% of endpoints on the whole panel and 93.3% on the missense subset, 89.4% for the ten predictors resting on fewest genes, and one comparison the panel does not identify, AlphaMissense above ESM1b (bounds overlapping by 0.0008), was declared identified in 23% of resamples.

#### Note S4. Judging each comparison inside the predictors' declared scopes, and covered AUROCs printed side by side

Scopes, read from each predictor's documentation (Table S25, Scope column), nest: the whole panel for 14 predictors, coding substitutions for three and missense substitutions for the other 32. On the panel's 202,643 missense substitutions, median reach on pathogenic variants rises from 0.36 to 0.98 and the median must-answer penalty falls from 0.284 to 0.035. Note S3's confidence sets are recomputed on each pair's variants. Of the 431 identified on the narrower scope, 377 stay identified with 95% confidence; 316 of the 431 stay identified under Holm's procedure and 322 under the max-T bootstrap; all 49 predictors appear in an identified pair, against only 23 of the 49 on the whole panel. The 431 are verdicts on different panels and need not be transitive. Of the 1,176 comparisons, 745 remain undecided on the narrower scope, only 129 of them within 0.02 AUROC of being decided.

Two readings bracket the 431. MetaSVM and MetaLR declare nonsynonymous SNVs and MetaRNN rare ones, though it scores them all; read as every nonsynonymous consequence, the three leave 393 identified. Judged only on the 188,926 missense variants whose ClinVar record gives no frequency of 0.01 or more in any of its three cohorts, the five predictors Table S25 scopes to rare missense variants, M-CAP and the four VARITY scores, leave 435. Of the 431, 181 join two predictors neither of which is a conservation track, 52 of them with neither listing ClinVar among its training data, and 115 of the 231 comparisons among the 22 predictors trained without clinical labels or allele-frequency inputs (Note S7) are identified. On the missense subset, 180 of the 429 identified comparisons join two predictors neither of which is a conservation track, and in 131 of those at least one lists ClinVar among its training data, so its covered AUROC can carry training overlap (Figure 3d; Tables S25 and S26).

Scope matching removes the cost of a declared scope, not that of class-dependent silence inside it: 18 of the 49 reach the classes unequally by at least 0.05 inside their scope, most strongly fathmm-XF_coding, which declines pathogenic variants more often (−0.224).

Of the 58 undecided whole-panel comparisons whose side-by-side order reverses on the variants both score, 22 do so under Holm's correction with the 1,093 undecided comparisons as the family and only the reversals tested, each reversal's two-sided p-value (normal, on the standard deviation of its 2,000 gene-resampled differences) being held to 0.05/1,093, 0.05/1,092 and so on, and 25 under Holm's correction over all 1,176, the family of Additional file 3's dbnsfp_convention_pairs.csv; that run uses the same p-values but ranks every comparison's shared-variant difference, so the step-down passes the many decisive differences that do not reverse and reaches laxer thresholds before it stops. On the missense subset 6 of the 747 reverse, none resolved. On the whole panel, AlphaMissense trails CADD_raw by 0.007 side by side but leads by +0.025 (+0.021 to +0.028) on the 205,373 variants both score.

#### Note S5. Identification: completions, transcript join, within-gene, rare-variant and label-age accounting, and the missingness indicator

Completions under a stated rule. Over completions giving every declined variant one common score, REVEL's ClinVar AUROC runs from 0.414 to 0.693, AlphaMissense's from 0.402 to 0.677 and CADD's from 0.990 to 0.991; phyloP and phastCons keep one value. Under such rules nine of the ten ClinVar pairs are ordered, every one except REVEL against AlphaMissense, compared with the three identified without a rule.

The transcript join. On ClinVar's own transcript, 52,617 of the 253,041 variants REVEL's release scores are not missense. AlphaMissense's canonical-transcript file leaves 11,196 ClinVar missense variants unscored, 1.7% of pathogenic and 7.0% of benign ones. Restricted to the 204,191 variants ClinVar calls missense, REVEL's bound is [0.9450, 0.9733] and AlphaMissense's [0.8804, 0.9664], against 0.9346 for CADD, which scores essentially all of them, 0.8872 for phyloP and 0.8159 for phastCons. On the 202,643 missense substitutions of the dbNSFP panel, REVEL's own release scores 8,248 that dbNSFP leaves without a REVEL value, 1,852 pathogenic and 6,396 benign, against 2 the other way; AlphaMissense's release scores 1,982 that dbNSFP leaves unscored and leaves 8,227 unscored that dbNSFP scores. On the whole dbNSFP panel the GERP++_RS and GERP_92_mammals tracks differ by 0.0383 in covered AUROC while their reach runs from 0.6723 to 0.9961. Reach on the 49-predictor panel is therefore reach as dbNSFP distributes it, an upper bound on what a laboratory reading one clinical transcript from dbNSFP would see, though not on what a tool's own release reaches. The source also moves Note S7's breakdown points: on the missense subset REVEL's lead over AlphaMissense breaks at λ = 0.397 with each tool's own release against 0.106 with dbNSFP's columns, and on the whole panel AlphaMissense's own release leads REVEL's in covered AUROC, 0.9631 against 0.9540, reversing dbNSFP's order. With the tools' own releases REVEL's lead over CADD on the missense subset is identified (λ* = 1.88), though it breaks at λ = 0.423 with dbNSFP's columns, while AlphaMissense's lead over CADD, identified with dbNSFP's columns (λ* = 1.12), breaks at λ = 0.398 with the tools' own releases.

Head-to-head differences with genes resampled. On the ClinVar panel, variants sharing any gene symbol, from ClinVar's own gene annotation or the dbNSFP gene, form one cluster. REVEL leads AlphaMissense by +0.0096 [0.0065, 0.0128] on 192,168 variants, and on the variants ClinVar calls missense REVEL leads CADD by +0.0392 [0.0349, 0.0434], AlphaMissense leads CADD by +0.0318 [0.0280, 0.0356] and REVEL leads AlphaMissense by +0.0096 [0.0066, 0.0128]. These intervals are several times wider than variant-level DeLong intervals on the same differences, and every sign survives them. Restricted to missense variants, REVEL's lead over CADD is four times its whole-panel +0.0091, because its values on variants ClinVar calls non-missense dilute it. On the whole ClinVar panel CADD leads REVEL by 0.034 on the sets each covers and by 0.465 once each must answer, a gap that prices REVEL's declared scope rather than its errors (Figure 4e; Figure S16b). Three of the ten pairs are identified, exactly the three among CADD, phyloP and phastCons, and seven of the ten on the 204,191 variants ClinVar calls missense.

Within gene. Pairs formed within a gene and pooled by pair count identify 544 comparisons on the missense subset against the pooled 429, and 110 on the whole panel against 83. On the missense subset 400 are identified both ways, and one shared comparison reverses; at the point estimates 115 of the 1,176 pairs reverse sign. Within-gene pairs are 0.03% of the missense subset's pathogenic–benign pairs, so genes rich in both classes dominate them.

Rare variants. A missense variant counts as rare when ClinVar's Exome Aggregation Consortium (ExAC), Exome Sequencing Project (ESP) and 1000 Genomes allele frequencies give it none or a largest value below 0.001 (0.01 as a sensitivity analysis). In those frequencies, 88% of pathogenic and 46% of benign missense variants have none, so "rare" mostly means absent, and the check cannot reach frequency-assigned benign labels or predictors that take frequency as input. Restricted to the 165,933 variants below 0.001, 414 are identified against 429, 389 of them in common and none reversed; below 0.01 the count stays at 429. Review status matters as well: one star holds 72.3% of the benign missense labels but 58.8% of the pathogenic ones. At two stars or more, where allele frequency alone separates the missense classes at 0.959 on two-star and 0.861 on higher-star labels, against 0.677 on one-star ones, 566 missense-subset comparisons are identified rather than 429 (Figure 3c).

Label age. ClinVar's release of fileDate 2021-01-02 postdates every ClinVar snapshot the 49 predictors' stated training data name, though four of the 15 trained or tuned on ClinVar name none (LIST-S2, MPC, MVP and gMVP, missense predictors of Table S25); a label that release does not carry with the same class cannot have entered the ClinVar training data of any predictor that dates its snapshot. The archived releases are read by the same classification rule as the reach panel (Methods). Thirteen of the 49 also list the Human Gene Mutation Database (HGMD) among their training data, and a ClinVar label newer than the release can repeat an older HGMD entry. On the dbNSFP panel 232,563 labels are newer, 144,157 of them on the missense subset; 220,167 were absent from that release, 12,297 carried another classification and 99 the opposite one. On the newer labels 77 whole-panel comparisons are identified, all among the 83, and 396 on the missense subset, 388 of them among the 429; none reverses. The six whole-panel comparisons that drop out are near ties whose bounds only just separate on all labels. Because the newer labels are 70.8% of the panel, the stronger test sets them against the disjoint older ones. On the 95,765 labels the 2021 release already carried with the same class, 94 comparisons are identified, 71 of which are also identified on the newer labels and one reversed there, and 535 on the missense subset, 384 of them also identified and two reversed, every reversal between two conservation tracks. The covered AUROC of the 15 predictors that used ClinVar labels in training or tuning rises by a median of 0.006 on the newer labels, against 0.030 for the 31 that did not use them (Table S26); REVEL, AlphaMissense and popEVE, which used ClinVar only for testing, calibration or evaluation, are in neither group. The sequence-blind gene prior, which no training can reach, reads 0.041 higher on the newer labels, so that contrast mixes training overlap with composition. Reweighting the newer labels, class by class, to the older labels' mix of consequence group, review status and decile of the out-of-fold gene prior removes the composition these capture: the gene prior's gain falls from 0.041 to 0.001, which checks the balance, since its decile is a stratum. The median shift, which carries no interval, becomes −0.005 for the ClinVar-trained predictors against +0.004 for the 31 others. Against the release of fileDate 2023-01-07, 182,622 labels are newer: 81 identified, 80 among the 83, one reversed (DANN against phyloP100way_vertebrate), and 392 on the missense subset, 375 among the 429, none reversed. On the ClinVar panel the 1,136,238 labels newer than the 2021 release leave the three whole-panel orderings and the seven on missense variants unchanged, with REVEL's and AlphaMissense's bounds at [0.047, 0.998] and [0.030, 0.999].

The missingness indicator. For both missense-only scores the indicator of whether a value was produced separates the classes better than the values do under the must-answer rule on the whole ClinVar panel (Table S8, missingness alone against must-answer). The indicator's midrank AUROC is 0.5 + (r₊ − r₋)/2, r₊ and r₋ being the reach on pathogenic and benign variants (Methods), exactly one half for a scorer that answers everything. Set against the must-answer AUROC, even read orientation-free, the indicator wins whenever |r₊ − r₋| > r₊r₋, whatever the covered AUROC, because the one-half rule discards what the indicator knows. Whether the values add anything is tested by the lexicographic completion, which ranks by the indicator and then by value and beats the indicator by exactly ρ(A_cov_ − 0.5). Its gene-resampled interval excludes zero for every one of the 49 predictors, on the whole dbNSFP panel and on its missense subset (Table S9). For REVEL and AlphaMissense the missingness follows declared scope and the transcript join described above, so the indicator is substantially a missense indicator, and stratifying by consequence removes most of it.

#### Note S6. Comparing predictors within gene, what that changes, and the accounting inside each gene

Pooled and within-gene comparisons can disagree: on the dbNSFP panel's missense subset AlphaMissense against REVEL reads −0.0100 pooled and +0.0101 within the 3,143 genes carrying both classes, so the sign flips. On the human ClinVar missense panel the same difference, AlphaMissense minus REVEL, moves from −0.0096 pooled (REVEL's lead in Note S5) to +0.0124 within gene and the must-answer difference from −0.0357 to −0.0044; these figures are indicative (Note S22).

Resampling genes rather than variants gives [−0.0426, −0.0302] for that pair's pooled must-answer difference on the human ClinVar missense panel, 5.9 times wider than the [−0.0368, −0.0347] of the closed-form calculation that treats variants as independent.

We report no overall ranking of the 49 predictors; the bound orders only the pairs it identifies (Figure 3b, g, Note S7 and Figure S11). Every interval on the dbNSFP panel is gene-clustered on a canonicalised gene key; an uncanonicalised key would fragment genes and narrow the intervals (Additional file 2: src/ccs/build_dbnsfp_panel.py). The gene-balanced drop is, for each predictor, its pooled AUROC minus the unweighted mean of its within-gene AUROCs over the genes carrying at least five variants of each class. Of the whole panel's 15,606 genes, 2,418 carry at least five of each class, as do 1,203 of the missense subset's 15,389; the number entering a predictor's mean varies with the genes it scores. On the missense subset the drop is a median of 0.027 across the 49 predictors, each averaged over up to 1,203 genes. The drop's two terms differ in variant set and aggregation, so it is not a bias in a fixed estimand; on the whole panel it is negative for three predictors. It carries no interval.

The gene-balanced drop does not measure gene-level circularity. The 13,188 genes the within-gene term never sees hold 28.3% of the panel's variants and contribute nothing to the drop. The nine label-free conservation tracks, which never saw ClinVar's gene labels and so cannot be circular, carry a larger median drop on the whole panel than the other 40, 0.0438 against 0.0280. The drop thus confounds circularity with the between-gene component of constraint signal, which is larger for the label-free tracks than for the other 40.

Overlap between a predictor's training variants and these labels is not corrected here. Two contrasts can be measured, only the second bearing on variant-level leakage: BayesDel with its allele-frequency feature leads its frequency-free version by +0.0072 [+0.0067, +0.0077], and VARITY_R and VARITY_ER lead their leave-one-out versions by +0.0017 and +0.0005 ([+0.0016, +0.0019] and [+0.0004, +0.0006]), genes resampled.

##### *The accounting inside each gene*

The per-gene map repeats the bound inside each of the 132 genes with at least 30 missense variants of each class, on that gene's missense variants alone. In each, a predictor is dominated when another's lower bound exceeds its upper bound.

Within a gene, bounds are narrow, so a median of 987 of the 1,176 comparisons are decided at the point estimates; a median of one predictor is undominated per gene, and 39 of the 49 in at least one.

Such leads rarely survive sampling: with each gene's pathogenic and benign variants resampled separately, a sole undominated predictor keeps its lower bound above every other predictor's upper bound in at least 97.5% of draws in only 2 of the 87 such genes, BayesDel_addAF in COL7A1 and ClinPred in HNF1A.

#### Note S7. How far each comparison survives: breakdown points and a dominance map

The breakdown point of a comparison [54], the least λ at which the two predictors' intervals (Methods) meet, is λ* = (A_hi_ − A_lo_)/((1 − ρ_hi_)A_hi_ + (1 − ρ_lo_)(1 − A_lo_)), hi marking the higher covered AUROC. The contamination model sets one λ for both predictors and acts on pathogenic–benign pairs, not variants, so λ* needs only the two covered AUROCs and the two pair coverages.

With genes resampled as in Note S3, the one-sided 95% lower confidence limit of each point estimate λ*, 0 when at least one draw in twenty reverses the pair, is 0.011 for REVEL against AlphaMissense on the whole panel and 0.076 on the missense subset. At these lower limits, 829 and 622 of the 1,176 whole-panel comparisons remain ordered at λ = 0.05 and 0.1, against the 866 and 664 the main text gives at the point estimates.

The 22 predictors trained without clinical labels or allele-frequency inputs are ranked on labels none of them trained on. Four of them, AlphaMissense, PrimateAI, popEVE and MisFit_D, learned from population data, on which many benign labels also rest, and clinical labels reach four others, CADD_raw, DANN and the two Eigen scores, indirectly through the PolyPhen-2 scores among their features. On the missense subset seven are undominated: AlphaMissense, ESM1b, popEVE, MisFit_D, the two Eigen scores and MutationAssessor. Of these, popEVE, Eigen-PC and MutationAssessor are undominated only because their declines widen their bounds. Under the gene-clustered 95% sets of Note S3, PROVEAN and CADD_raw drop out of those dominated on the missense subset (Figure 3g), and Holm's procedure over the 1,176 comparisons keeps the rest. Circularity through PP3 and BP4 can act on these comparisons wherever a ClinVar assertion used a scored tool (Note S8).

#### Note S8. Calibrated clinical evidence, label circularity and functional labels

A score that cannot be calibrated in a species without labels offers discrimination alone, not calibrated ACMG/AMP evidence [18,55].

We applied ClinGen's calibrated intervals to the ClinVar panel's 204,191 missense variants (Table S22). Of the four tools, phyloP is the weakest at clinical thresholds: REVEL leaves 14.6% of pathogenic missense variants indeterminate or unscored, AlphaMissense 24.9%, CADD 17.5% and phyloP 33.9%, almost all indeterminate rather than unscored (Figure 4f; Table S22). Evidence opposite to the ClinVar label is rarer: across the four tools BP4 reaches 3.4% to 7.7% of pathogenic missense variants and PP3 2.7% to 8.3% of benign ones (Figure S13; Table S22). A calibration sets odds, not an error rate, so it expects some such evidence. Evidence yield has been measured per proband in rare disease [123]; these figures give it by ClinVar class. The CADD yields assume that intervals calibrated on an earlier release transfer to v1.7 scores, and REVEL, read as its maximum over transcripts, gives an upper bound on its PP3 yield and a lower one on its BP4 yield. BP4 also presumes no predicted effect on splicing [55], which these counts do not check, so the BP4 shares may include pathogenic variants that act through splicing.

Label circularity. The ClinGen Evidence Repository publishes each expert-panel classification with the ACMG/AMP criteria it applied [46], under Creative Commons Zero (CC0) 1.0 with attribution to ClinGen. Of its 13,293 classifications, keeping one per GRCh38 single-nucleotide variant leaves 7,479 labelled variants. Applied criteria were converted to Tavtigian et al.'s naturally scaled ACMG/AMP points [102], benign ones negative: supporting 1, moderate 2, strong 4 and very strong 8, with strength read from a criterion's suffix where given. A classification is pathogenic at 10 or more points, likely pathogenic at 6 to 9, likely benign at −1 to −6 and benign at −7 or less [102], with BA1 stand-alone [55]. On the 7,479 deposited labels the points reproduce the stated class in 7,274. Of the 7,479 labels, 4,324 applied PP3 or BP4, and 1,085 reach their class with those points but not without them; a label whose applied criteria miss its class even with those points does not rest on them.

All 7,226 expert-panel variants in the ClinVar panel, 4,168 of them missense, keep their Repository class in ClinVar. Removing the missense labels that depend on PP3 or BP4 lowers all four tools' AUROCs alike, AlphaMissense's included (Table S23), so the drop reflects which labels are easy to call, not reuse of one tool's scores. We do not compare against the subset of classifications that never applied PP3 or BP4: which classifications apply computational evidence is itself selected, and that subset differs in gene and consequence composition.

A ClinVar assertion may rest on PP3 or BP4 evidence, which changed 15% of the expert-curated missense classifications it was applied to [56]. In the ClinVar panel, 603 of 3,024 pathogenic and 36 of 1,144 benign expert-panel missense labels depend on that evidence. Removing them lowers each tool's AUROC by 0.004 to 0.008 at the point estimates, with upper 95% limits of 0.008 to 0.015 (Table S23). Removing the 560 of the 1,144 benign expert-panel missense labels that reach their class only through BA1, BS1 or BS2 raises each tool's AUROC, by +0.013 to +0.034, every interval excluding zero with genes resampled, and each by more than removing the labels that depend on PP3 or BP4 lowers it (Table S23); phyloP's gain, +0.034 (95% CI +0.014 to +0.055), is four times its 0.008 fall. The check reaches only expert-panel labels, 2.0% of the panel's missense variants.

Each expert panel's specification in ClinGen's Criteria Specification Registry names the tools it reads for PP3 and BP4, gene by gene, and 39 of the 45 panels in the Evidence Repository read REVEL for missense PP3, alone or with others. Matched to the current specification for their gene, 3,631 of the 4,168 expert-panel missense variants in the ClinVar panel come from panels whose specification names REVEL, as do 584 of the 639 missense labels that depend on those points. Among those 3,631, removing the dependent labels lowers REVEL's AUROC by 0.006 (0.003 to 0.010) and AlphaMissense's by 0.005 (0.002 to 0.009). The difference, +0.002 (−0.002 to +0.005) on the same gene draws, does not separate the tool whose scores the panels read from one they almost never name; as the two tools' scores are correlated, it excludes a REVEL-specific effect larger than about 0.005 rather than any effect. Two specifications name AlphaMissense beside REVEL, and a classification made under an earlier version of its specification may have followed other rules.

ClinGen's calibrations set the PP3 and BP4 intervals from ACMG/AMP likelihood-ratio thresholds of 2.406, 5.790, 33.53 and 1,124 from supporting to very strong at a prior of 0.0441 [18]. A likelihood ratio averaged over a level (Table S27) is a weaker test than the calibration's own at the level's inner threshold, where its evidence is weakest.

Each level is held to point-system odds of 2.08, 4.33, 18.7 and 350 from supporting to very strong at a prior of 0.1, reciprocals for BP4 [102]. Read as likelihood ratios, each averaged over its level, every PP3 and BP4 level of the four tools meets, at its point estimate, the odds its strength targets, and does so in each label-age and review-status stratum taken separately, including labels added since the 2023 release (Table S27). This is a consistency check, not a validation: a ClinVar label may itself rest on PP3 or BP4 evidence from these tools, the most recently added labels are the stratum most exposed to it, and likelihood ratios on such labels cannot separate a valid calibration from labels that applied it. Read in the 1% of a tool's scored missense variants nearest each level's inner threshold, every level meets its target except CADD's BP4 supporting level, which reads 0.56 (0.47 to 0.65) against 0.48, an interval reaching the target but lying wholly above 0.42, the reciprocal of the calibration's threshold of 2.406. A declined variant, taken as its own level, carries a ratio below one (Table S27), since REVEL and AlphaMissense decline benign missense variants more often than pathogenic ones. A decline nonetheless carries no evidence: no calibration assigns it any, and the ratio reflects which variants each release leaves out, not a rule a laboratory could carry to other variants.

The release that built the reach panel holds 2,097,787 single-nucleotide missense variants of uncertain significance at one review star or better, the variants a laboratory most needs evidence for. AlphaMissense's own release, read by GRCh38 position, declines 2.7% of them, and its calibrated intervals [19] give PP3 evidence to 14.3% and BP4 evidence to 47.4%, leaving 38.2% indeterminate or unscored (Table S30), against 24.9% of pathogenic labelled missense variants (Table S22). The unscored and the indeterminate among them are counted apart in Note S38, with the variants of uncertain significance that no missense calibration reaches.

A label that no computational evidence can enter checks the tools' ClinVar results for concordance but, on BRCA1, cannot test circularity through PP3: BRCA1's expert panel reads BayesDel for PP3, and 55 of its 65 BRCA1 single-nucleotide classifications in the Evidence Repository apply PS3 or BS3, the ACMG/AMP functional-evidence codes. The GRCh37-to-GRCh38 offset that places the BRCA1 saturation-editing panel [71] on ClinVar's build, checked against alleles (Additional file 3), and the SIFT query in cattle (Note S16), whose ARS-UCD1.2 coordinates were carried unchanged to ARS-UCD2.0 (GCA_002263795.4) after the two sequences matched at every cattle reference base, are the only cross-assembly projections in any comparison. On the 308 substitutions ClinVar labels at one star or better, the functional class, loss of function against functional or intermediate, and ClinVar's agree on 300.

AlphaMissense, REVEL and CADD read 0.944, 0.926 and 0.902 on the functional labels and 0.954, 0.938 and 0.931 on ClinVar's, and REVEL minus AlphaMissense on the functional labels is −0.018 (−0.048 to +0.011) with residues resampled. The check covers one gene, whose ClinVar classifications can themselves draw on functional assays.

On all 2,077 missense substitutions of the saturation-editing panel, AlphaMissense, read from its own release by protein substitution, separates loss-of-function variants at 0.889 against Evo 2's 0.822 at 8,192 bp, +0.067 (+0.034 to +0.099) with residues resampled. On the 2,803 substitutions the canonical transcript annotates, Evo 2 reads 0.849 at 8,192 bp; the 1,090 unannotated ones are not a certifiable non-coding set (Figure S6).

LDLR allows the test of circularity through PP3 that BRCA1 does not. Its Familial Hypercholesterolemia expert panel reads REVEL for PP3 and BP4, and 33 of its 233 LDLR single-nucleotide classifications in the Evidence Repository reach their class only with those points. Tabet et al. [118] measured low-density lipoprotein uptake for nearly every LDLR substitution. They proposed a score of 0.04 or below as the threshold for a functional mutation, set on the clinical diagnoses of UK Biobank carriers rather than on ClinVar; that class is the functional label here. Of the 721 missense variants the assay scores and ClinVar labels at one star or better, 685 pathogenic and 36 benign, all 146 at or below the threshold are ClinVar-pathogenic, as are 539 of the 575 above it. AlphaMissense, REVEL and CADD read 0.706, 0.616 and 0.572 on the functional labels and 0.971, 0.972 and 0.968 on ClinVar's: REVEL minus AlphaMissense moves from −0.090 (−0.148 to −0.032) to +0.001 (−0.022 to +0.020), residues resampled, a shift of +0.091 (+0.028 to +0.149) on the same draws. Reuse of REVEL's evidence would move the difference that way, but removing the 33 labels that record the reuse lowers REVEL's AUROC on ClinVar's labels by only 0.002 (0.000 to 0.005) more than AlphaMissense's. The rest reflects what the functional labels ask, loss of uptake against partial or normal uptake, among variants ClinVar mostly calls pathogenic. On LDLR the two labels measure different things, and the test cannot separate reuse the Evidence Repository does not record from that difference.

#### Note S9. The fitted components against the DOME recommendations

The article fits few models, and none is its subject: Evo 2 and every published predictor are scored as released. The fitted components are the dbNSFP panel's sequence-blind baselines, the cross-validated logistic models of Note S10, the calibration and selective-prediction layers (Note S28), the supervised floor (a convolutional network and a k-mer classifier) and the embedding probe of Note S33. For these, following the Data, Optimization, Model and Evaluation (DOME) recommendations for supervised learning in biology [124]:

Data. Every fitted component uses deposited panels (Additional files 2 to 6), whose sizes and class counts are stated where each is used, except the supervised floor's 500-bp sequence windows, which are not deposited. The gene and consequence baselines are fitted out of fold, in five label-stratified folds over eight seeds; calibration transfer holds each species out of its own map; the supervised floor is evaluated leave-one-species-out and by 5-fold cross-validation within species.

Optimisation. The gene baseline is the gene's pathogenic rate shrunk towards the training folds' rate with an untuned pseudo-count of one; the gene-and-consequence baseline shrinks each gene–consequence pair's rate the same way towards its consequence's rate, itself shrunk towards the folds' rate. Across ten estimators of the pair's rate, the gene-and-consequence baseline reads 0.959 to 0.975, above CADD's must-answer AUROC of 0.968 in seven of them and above every other predictor's in all ten, and the bound rule identifies it above 13 to 23 of the 49. On each predictor's own covered variants, the gene rate exceeds the covered AUROC of 18 of the 49. Out of fold, the gene rate reads 0.880 on the whole panel, above the must-answer AUROC of 45 of the 49 and the upper bound of 10. On the missense subset, allele frequency alone reads 0.734 and the gene-and-frequency baseline 0.956, above the covered AUROC of 33 of the 49, all 18 that learned from neither clinical nor population labels among them; the 16 above it learned from clinical or population labels. The missense subset's gene-and-frequency baseline is an unpenalised logistic regression on the logit gene rate, the base-10 logarithm of the largest ExAC, ESP or 1000 Genomes frequency ClinVar carries (floored at one in a million) and a frequency-present flag. Within the training folds each row takes its gene rate out of fold, so no row's rate has seen its own label. Folds are not held out by gene, since the gene rate would then carry no information and the combined rule would reduce to consequence class alone.

On the dbNSFP panel, with rates estimated out of fold, consequence class alone reads 0.837, and gene and consequence together 0.974. Under the primary estimator the combined rule lies above the must-answer values of all 49 predictors (not under every estimator of the rates) but is identified above only 23 of them, as many as under any estimator. On the missense subset the gene alone reads 0.930, close to the 0.921 reported on ClinVar missense variants [17]. MutationTaster2021 goes further, labelling ClinVar-pathogenic variants deleterious by lookup [125]; it enters 4 of the 83 identified comparisons and 7 of the 429. On the ClinVar panel, consequence class alone reads above phyloP's 0.922 (Figure 4c). Under the ACMG/AMP criteria [55], a null variant in a gene where loss of function causes disease is itself very strong evidence of pathogenicity (PVS1), and a synonymous variant predicted not to alter splicing, at a base not highly conserved, is evidence of benignity (BP7). The pathogenic shares of the consequence classes thus partly reflect how such labels are assigned. An accuracy on a panel of this composition is therefore largely a statement about the panel, though a predictor that has learned gene identity has learned something real.

Model. The glmtrust package implements the baselines and the calibration and abstention layers.

Evaluation. Discrimination is AUROC, calibration error is reported under twelve binning rules (Note S28; Additional file 2: reports/tables.md, Table D7), and every comparison is charged for what it declines.

#### Note S10. What Evo 2 adds to annotation and conservation, and the capacity ladder

A cross-validated logistic model adding the Evo 2 feature at the 1,001-bp single-token readout to per-base GERP raised AUROC over GERP alone by a pooled increment (ΔFM) of +0.0202 (95% CI [+0.0132, +0.0279], variant-level bootstrap; n = 9,532 variants across nine species, 5-fold cross-validation, within-species standardised), a modest contribution whose interval treats variants as independent. The two scores are only weakly correlated (median per-species Spearman +0.31), so the increment is not wholly redundant with per-base GERP.

Three summaries of the increment over conservation answer different questions. The pooled ΔFM of +0.0202 above folds all nine species into one cross-validated model. The n-weighted mean of the per-species ΔFM is +0.054 [+0.039, +0.078], a nine-cluster percentile interval, which undercovers (Note S20). The unweighted species mean, the paper's primary aggregation, is +0.071 [+0.029, +0.113] (t on eight degrees of freedom; Figure S3c) at the 1,001-bp readout and +0.108 [+0.051, +0.166] at 8,192 bp. The per-species values Figure S3c draws, positive in all nine species from +0.024 in pig to +0.180 in goat, carry no deposited interval or p-value, so no per-species significance statement is available.

The same construction measures what Evo 2 adds to annotation. At 1,001 bp Evo 2 adds, by the species mean, +0.027 [+0.013, +0.041] over the four-level snpEff impact ordering, on all 11,130 variants, and +0.003 [−0.001, +0.008] over impact and GERP, on the 9,529 variants both readouts and GERP score. At 8,192 bp the two increments are +0.032 [+0.016, +0.048] and +0.008 [+0.002, +0.014], the second positive in eight of nine species; with each species predicted by a model fitted on the other eight, they are +0.024 [+0.013, +0.036] and +0.005 [+0.003, +0.008]. Once annotation and conservation are known, the likelihood adds little on this panel; conservation here is GERP, and the increment over the 241-mammal phyloP was not computed.

On the capacity ladder all three checkpoints are scored identically on the same 11,109-variant panel, so the 40B rung is Table 1's 8,192-bp column. The atlas's human 1B scores are a scoring fault and are excluded, so the ladder is reported on the eight OMIA species. A second 1B implementation, run on 327 of the same windows (Note S29; Additional file 3), agrees with the atlas's 1B scores within each of the other eight species (Spearman 0.82 to 0.97) but not in human (0.25), and reads 0.945 on the 40 human windows against 0.650 for the atlas's 1B. Across the atlas, human positives score a median 0.00019 at 1B against 0.0031 at 7B (0.0011 to 0.0023 at 1B in the other eight species), and their 1B and 40B scores rank-correlate at 0.06 (0.33 to 0.92 among the other species' positives). Pooled over the 287 non-human windows, the second implementation reads 0.896 and the atlas's 1B 0.870, so the 1B rung is known only to about 0.03. At 8,192 bp the eight-species means (evo2_1b_base, evo2_7b and evo2_40b; Table S2) give a 1B-to-40B gain of +0.043. The 1B checkpoint was trained at an 8,192-token context [31], while 7B and 40B are context-extended and differ in training tokens, so no step isolates model size. The 1B-to-7B step is positive in all eight species (exact two-sided sign test, p = 0.008), runs from +0.007 (horse) to +0.061 (sheep), and its locus-resampled 95% interval excludes zero in cattle, dog and sheep; the mean step is +0.028 [+0.014, +0.042]. The 7B-to-40B step is positive in seven of the nine species, but over the eight OMIA species it is +0.015 [−0.005, +0.035] and is not resolved. To keep the readout fixed, the splicing and ClinVar consequence panels were rescored at 1B and 7B with the 40B single-token scorer on identical windows and offsets, so only the checkpoint changes.

**The capacity ladder at 1B, 7B and 40B, on the splicing and ClinVar consequence panels**

| **panel** | **1B** | **7B** | **40B** |
| --- | --- | --- | --- |
| Splicing, n = 3,912 | 0.5351 [0.5133, 0.5576] | 0.5304 [0.5091, 0.5529] | 0.5959 [0.5766, 0.6165] |
| ClinVar, n = 3,714 | 0.6029 [0.5848, 0.6220] | 0.6364 [0.6191, 0.6544] | 0.6857 [0.6682, 0.7033] |

Every rung scores the same variants, so the sevenfold step from 1B to 7B is paired: −0.0047 [−0.0214, +0.0121] on the splicing panel and +0.0334 [+0.0198, +0.0469] on the ClinVar consequence panel.

On this ladder the 1B and 7B rungs, evo2_1b_base and evo2_7b_base, are 8k-trained base checkpoints, while the 40B rung, evo2_40b, is context-extended, so any contrast reaching the 40B column spans context extension as well as capacity. Only the 1B-to-7B step is quoted as a checkpoint contrast, and its two checkpoints also differ in training data [31].

#### Note S11. The 8,192-bp readout: AUPRC, the cross-species share of ranked pairs, and the readout effect

At the 8,192-bp readout Evo 2-40B reads 0.943 by the species mean and 0.973 pooled (95% CI [0.968, 0.978], variant-level bootstrap; n = 11,109). Its macro-mean AUPRC is 6.3 times the no-skill base rate (Table S7, with per-species and pooled values), so the discrimination is not a base-rate artefact at this readout. The pooled figure is given only beside the species mean: of the 19,846,664 positive–negative pairs it ranks, 83.0% are cross-species, so it largely measures whether one species' positives separate from another species' population variants, which is not this paper's question. The unweighted species mean is therefore our primary summary throughout: species are the unit of generalisation, and pooling allows the largest panels to dominate while letting per-species miscalibration cancel. On the same variants the single-token readout gave 0.878 by the species mean and 0.881 pooled. Switching readouts is worth +0.065 AUROC by the species mean (95% CI [+0.032, +0.097], t on eight degrees of freedom) and +0.092 pooled (95% CI [+0.083, +0.101], paired variant-level bootstrap; n = 11,109; per-species values in Additional file 2, reports/tables.md, Table D1). The effect ranges from +0.009 in goat, which has no interval, to +0.149 in human; the species-mean interval excludes zero, and its lower limit, +0.032, is less than half the pooled point estimate.

#### Note S12. Fixed false-positive operating points, and Table 1's intervals: locus clustering and the macro row's t interval

The operating point a prioritisation would use is the threshold at a fixed false-positive rate: the lowest score that at most that share of a species' negatives reach. At 1%, the 8,192-bp readout recovers 0.532 [0.371, 0.812] of cattle's positives, 0.543 [0.468, 0.659] of human's, 0.714 [0.548, 0.917] of chicken's, 0.746 [0.316, 0.899] of horse's, 0.750 [0.559, 0.875] of pig's, 0.811 [0.688, 0.920] of cat's, 0.821 [0.640, 0.950] of sheep's and 0.859 [0.787, 0.906] of dog's. Goat, with 89 negatives, has no 1% operating point. At a prior of one causal variant in 1,000 candidates, cattle's recall of 0.532 at that rate means a positive predictive value near 0.05. A consequence filter keeping snpEff's HIGH and MODERATE first terms flags 0.2% to 2.2% of negatives in the eight OMIA species, 1 to 9 per species but 38 of cattle's 1,879 and 25 of dog's 2,269. At those rates it recovers a mean 0.898 of positives, against 0.738 for the 8,192-bp readout (mean difference +0.160, 95% CI −0.002 to +0.322, t on seven degrees of freedom). With loci resampled and no correction over the eight species, the difference excludes zero only in sheep, +0.339 [+0.075, +0.906], and narrowly in cat, +0.139 [+0.009, +0.395], and with 1 to 38 flagged negatives per species these operating points are coarse.

Positives are catalogued disease- or trait-associated alleles and cluster at the locus scale; the trinucleotide-matched negatives do not. Counting a locus as a 100-kb bin, the positives of the nine panels occupy 2 (goat), 20 (chicken), 28 (pig), 38 (sheep), 36 (horse), 83 (cat), 147 (cattle), 187 (dog) and 944 (human) loci. Kish's effective cluster count is lower still where one locus dominates: 1.2 in goat, where eight of nine positives lie within 570 bp, and 6.7 in horse, where 26 of 71 positives share one 100-kb bin. Negatives are effectively unclustered, at most two or three per bin in seven species. Re-binning at 1 Mb lowers the large effective counts (human's by 38%) but leaves the small ones almost unchanged (goat 1.2, horse 6.6, pig 22.3; chicken and sheep the same), so the caveat below on small counts does not depend on the bin width. Besides Table 1's locus-clustered bootstrap (20,000 replicates), two other estimators are deposited: a variant-level percentile bootstrap and a cluster-robust DeLong interval (sandwich variance over loci, Student's t on G − 1 degrees of freedom, G the species' positive-bearing loci). The three agree within 1.07× in cattle, dog and human, where the effective cluster count is large, and elsewhere diverge without tracking it: sheep agrees as closely and pig within 1.08×, cat's printed interval widens most, and horse's 8,192-bp locus interval is narrower than the variant-level one. Cat widens by 1.51× at 8,192 bp, to [0.813, 0.964], the largest single correction in the table. Where the effective count is small (horse's 6.7, chicken's 14.0, sheep's 21.5 and pig's 23.1), no resampling estimator here is reliable, and those intervals are likely too narrow. Goat and chicken carry fewer than 30 positives, so their point estimates should not be ranked against the others.

Species-level intervals and verdicts are given only where positives occupy at least ten loci, a threshold set after the data were seen that only goat fails. Goat is not estimable; Table D1 in Additional file 2's reports/tables.md prints its unclustered interval for completeness only. The cluster-robust interval, the only estimator that makes the shortage visible, rests on one degree of freedom and spans [0.771, 1.000] at 1,001 bp and [0.758, 1.000] at 8,192 bp. Such spans say only that goat's discrimination is unquantified.

Table 1's GERP and conservation-matched columns carry no intervals by design: each is computed on the variants GERP scores, a different set from the Evo 2 columns in each species, so a paired interval across them would not be meaningful.

##### *The macro row's species-clustered t interval*

For the Mean (macro) row, a percentile bootstrap reads 0.048 and 0.046 wide against 0.061 and 0.058 for the t interval at the 8,192-bp and 1,001-bp readouts, widths taken before rounding, narrower for the reason Note S20 gives. Without goat the 8,192-bp macro mean is 0.941.

#### Note S13. The conservation-matched control, and the residual it leaves

Because positives preferentially occupy conserved sequence whereas population negatives do not, part of the atlas's separation could reflect a conservation confound rather than a signal beyond conservation. With negatives resampled to the positives' GERP profile (Methods), discrimination persisted: the matched mean over the eight species with a matched value is 0.834, against 0.869 unmatched, a drop of 0.034 on the unrounded means. On the GERP-scorable subset both columns are computed on, the drop reaches 0.064 in human, 0.761 against 0.825. Table 1's two Mean cells, 0.878 and 0.834, must not be subtracted: one averages nine species, the other eight, as goat has no matched value. The persistence is consistent with a signal beyond Ensembl's GERP rather than solely a conserved-versus-intergenic confound (within consequence class Evo 2 is not ahead of the 241-mammal phyloP; Note S43). Two caveats belong with that reading. First, sheep appears to gain from the control, rising from 0.903 in Table 1 to 0.911 matched, which removing a confound should not do. The matched value uses only the 451 of 616 sheep variants GERP can score, on which the unmatched AUROC is 0.912, so matching in fact costs 0.001. The apparent gain is the easier scorable subset, not the control. Second, the matched pool comes from GERP's class-biased scorable subset (Note S23, Table S1). This control therefore cannot fully escape the defect it is testing for, and its per-species values should be read as within-subset contrasts, not as panel-wide accuracies. Run at both readouts on the variants with an 8,192-bp score, the same control lowers the eight-species mean by 0.031 at 8,192 bp and 0.034 at 1,001 bp, so the readout gain survives it.

#### Note S14. What the atlas measures: composition controls, within-class pairs and the missense stratum by species

Atlas negatives are matched exactly on trinucleotide context and alternate allele, but not on GC content (below), genomic region, gene density, coding status or mappability; frequency is checked only in cattle (Note S32). This construction, catalogued positives against population negatives, measures how well catalogued positives separate from population negatives, not pathogenicity discrimination within a matched sequence class. The pooled missense stratum is the closest estimate of discrimination within a matched sequence class. Some 41% of the pairs it ranks are within-species. The missense stratum is small per species: only cattle (155, 36) and human (752, 167), variants and negatives in parentheses, hold enough negatives to be read as estimates on their own, and dog's 24 enter only the precision-weighted pool below. Local GC content runs higher in positives than in negatives in the eight non-human species and lower in human, a mean gap of about 0.07 over the nine. Read in each species' own direction, which biases the estimate upward, GC content alone separates the classes at a macro AUROC of 0.67 (101-bp windows; Additional file 2: reports/atlas_controls.json). The consequence control resamples nothing, recomputing discrimination within nested consequence strata (all annotated, non-coding, coding, and missense within coding).

In seven species the negatives come from Ensembl with no frequency threshold and include some deleterious alleles, which depress the measured AUROC; in pig they are common genotype sites of PigGTEx, which may inflate it. Trait rather than disease alleles make up 203 of the 737 scored non-human positives, and every headline atlas value includes them (Table S10). As expected of alleles selected in domestication, they sit at less constrained sites: at missense positives the median phyloP-241 is 5.54 for trait and 7.41 for disease alleles. Within consequence class, Evo 2's 8,192-bp margin over GERP in the eight OMIA species is +0.121 (+0.021 to +0.239) pooled by pair count, mostly dog's and cattle's pairs, but +0.045 (−0.113 to +0.204), not resolved, weighted by precision across species (the seven with at least 100 pairs); in human it is +0.012 (−0.030 to +0.056), loci resampled. On the 1,285 missense variants with a window score, mostly over cross-species pairs, the pooled AUROC is 0.812 at 8,192 bp against 0.799 at 1,001 bp, +0.013 (−0.017 to +0.042); Table S21 gives the within-class missense pairs.

Per species within this stratum, which unlike Table S21 does not require a GERP score, cattle's missense AUROC reads 0.744 at 1,001 bp and 0.672 at 8,192 bp, and human's 0.832 and 0.865, neither change resolved: −0.072 (−0.177 to +0.034) and +0.033 (−0.002 to +0.066), loci resampled. Human holds 92.3% of the stratum's within-species pairs, and weighted by precision across the species with at least 20 missense negatives, cattle, dog and human, the change is +0.003 (−0.130 to +0.136).

Table S21 pools its within-class pairs over species by pair count, and the eight OMIA species' pairs are mostly dog's (48.9%), cattle's (32.2%) and cat's (13.5%). Per species, Evo 2's within-class margin over GERP at 8,192 bp is +0.168 in dog and +0.138 in cattle, and runs from −0.146 in chicken to +0.644 in pig (160 pairs). Weighted by precision across the seven species with at least 100 pairs (random effects, between-species variance by restricted maximum likelihood, Hartung–Knapp–Sidik–Jonkman interval with Röver's modification), the readout gain is +0.065 (−0.006 to +0.135), against the pair-weighted +0.095 of Table S21.

#### Note S15. A human missense predictor carried to the atlas through the orthologue

Hu and Xie, scoring OMIA missense alleles at the aligned human residue, reported AUROCs of 0.55 to 0.61 against OMIA's non-deleterious alleles [69]; the atlas's population negatives are a different contrast, so its values are not comparable with theirs.

Each non-human atlas variant whose snpEff consequence includes missense_variant enters: 534 animal missense variants (455 positive, 79 negative) in 416 genes. We implemented the transfer on the gene models of the Ensembl release that annotated each species (Table S24), reading each variant's protein change on the canonical transcript, or else the longest coding sequence, with its reference base checked there, and taking the human orthologue from Ensembl Compara of that release, one-to-one where one exists. The candidate human proteins are those AlphaMissense [41] scores, found through UniProt's ID mapping [116] and read as AlphaMissense's own sequences. The animal protein is aligned to each by global alignment with affine gap costs [126] under the Blocks Substitution Matrix 62 (BLOSUM62) [127], and the candidate of highest identity is kept. A variant transfers when its reference residue aligns to an identical human residue, and takes AlphaMissense's prediction for that human position and the animal alternate residue.

Transfer reaches 401 of 455 (88.1%) positives and 42 of 79 (53.2%) negatives (Wilson intervals 84.8% to 90.8% and 42.3% to 63.8%), a gap of +0.350 (+0.239 to +0.454) with loci resampled (Figure S8a). The aligned human residue differs for 46 (26 positives), the release's Compara names no human orthologue for 28 (19 positives), the residue faces a gap in the alignment for 16 (8 positives), and one positive has no protein change on its release's gene models. That positive is a DMD allele that OMIA gives on the gene's minus strand, where it is the nonsense change p.Gln1617* (Note S32). The 28 include the eight cat CMAH blood-group alleles: CMAH was inactivated in the human lineage after the *Homo*–*Pan* divergence [128], so no human predictor can score them.

AlphaMissense reads 0.783 (0.710 to 0.851) on the 443 variants that transfer (Figure S8c, d). Three-quarters of the pooled pairs on these variants cross species; weighted by pairs within species, its lead over Evo 2 is +0.102 (+0.012 to +0.196). At a pair coverage of 0.469 its must-answer AUROC is 0.633 (0.589 to 0.677). Evo 2 reads 0.750 (0.692 to 0.806) on 533 of the 534 and GERP 0.684 on the 497 it scores, both inside AlphaMissense's sharp bound of [0.367, 0.899], so neither comparison is identified.

The rule's silence is class-dependent: population variants sit more often where the human residue differs or has no counterpart, so an accuracy quoted on the transferred variants alone describes a different variant set from Evo 2's. The comparison has limits of its own: the 79 negatives are few and mostly in cattle and dog, some positives are trait rather than disease alleles (Table S10), and the scores are human predictions read at an aligned position, not predictions for the animal protein.

#### Note S16. GPN-Star, a second genomic language model, on the chicken and human atlas variants, and SIFT's reach on the atlas

GPN-Star's released scores cover two atlas species, chicken on galGal6, the atlas's GRCg6a, and human on GRCh38 [30]. The score is the release's calibrated log-likelihood ratio (llr_calibrated) of each atlas variant's alternate allele, from the chicken model (gg6) and the human model (gpn-star-hg38-m447-200m). Both were negated, so that higher reads as more deleterious.

At 8,192 bp Evo 2 reads 0.954 in chicken and 0.974 in human, and GPN-Star 0.931 and 0.969, GPN-Star minus Evo 2 differences of −0.022 (−0.096 to +0.054) and −0.005 (−0.012 to +0.002) with positive-bearing 100-kb loci resampled, so neither model is resolved ahead of the other in either species. Where GERP scores, GPN-Star leads it by +0.200 (+0.077 to +0.356) and +0.095 (+0.082 to +0.109).

SIFT [1] was read from Ensembl's Variant Effect Predictor [70], release 116 (sheep, release 110), for every atlas variant Ensembl can map (tables/atlas_sift.README.md in Additional file 3). Ensembl's REST API serves neither CanFam3.1 nor a mapping from it, so the 2,497 dog variants are not mapped, and three cat positives are not mapped because Ensembl's genome lacks their reference base, the three the atlas's own reference check drops (Note S32).

SIFT scores amino-acid substitutions only, and in human it answers 43.5% of positives and 16.1% of negatives. With pair coverage below 0.02 in each of the seven non-human species, neither Evo 2 nor GERP is identified against it, a verdict on SIFT's scope rather than its errors.

A variant counts as scored when any transcript carries a SIFT score, and takes the lowest score over those transcripts. Read with lower scores as more deleterious, SIFT's covered AUROC is 0.828 (0.795 to 0.858) in human and 0.657 (0.556 to 0.757) in cattle, the two species with more than ten scored negatives, with positive-bearing 100-kb loci resampled; in the other six it rests on one to eight negatives. In human, where it answers far more negatives, its bound is [0.058, 0.988], which contains Evo 2's 0.974 and GERP's [0.805, 0.884]. On the variants SIFT scores, Evo 2 at 8,192 bp reads 0.903 in human and 0.652 in cattle, leading SIFT in human by +0.076 (+0.043 to +0.109) and unresolved against it in cattle, −0.005 (−0.123 to +0.112).

#### Note S17. The reference-likelihood confound across three panels

A variant delta is the difference between two window likelihoods, so it would inherit any separation the reference window's likelihood alone carries. The animal-panel rows below are measured on the 327-variant strand subpanel of Note S29, whose deposited scores keep the reference likelihood at all three aggregations: the 8,192-bp window mean, its central 1,001 bp and the single token at the variant.

**Discrimination from the reference window's mean log-likelihood alone, MLL(reference), on three panels**

| **panel** | **readout** | **MLL(reference) alone** |
| --- | --- | --- |
| nine-species animal subpanel (327 variants) | 8,192 bp, 40B | 0.5713 |
| nine-species animal subpanel (327 variants) | central 1,001 bp of the 8,192-bp window, 40B | 0.6908 [0.634, 0.749] |
| human ClinVar consequence panel | 1,001 bp, 1B and 7B | 0.4878 [0.4693, 0.5064] at 1B; 0.4610 [0.4426, 0.4802] at 7B |
| splicing benchmark | 1,001 bp, 1B and 7B | 0.5122 [0.4921, 0.5326] at 1B; 0.5736 [0.5509, 0.5966] at 7B |

The confound is real on the animal panel and modest, and it is checkpoint-dependent on both human panels, in opposite directions. At 1B the reference likelihood alone is indistinguishable from chance on each of them. At 7B the ClinVar quantity falls below chance (table above), so the reference window is mildly anti-predictive there. The animal panel's central kilobase reads higher, the largest of the window readings and 0.682 even on same-species pairs, at the window-mean aggregation, which carries about half the readout gain (Table S6). On the 389-variant strand subset of Note S29, scored at Evo 2-1B in 1,002-bp windows, the reference likelihood alone reads 0.5820 [0.5266, 0.6370], and discrimination rises across its quartiles from 0.7005 to 0.8792. The spread, 0.1787 against an asymptotic standard error of 0.069, is carried by the lowest quartile alone. Discrimination is therefore weakest where the reference window is least likely under the model. The single-token figures, 0.877 for the reference base alone and 0.899 for the full delta, are read from the 8,192-bp windows, with 4,096 bases of left context, not from the 1,001-bp arm's own pass. Within species the two statistics rank variants as closely as across them, and on same-species pairs the full delta reads 0.906 against 0.882 for the reference, although 88.9% of the pooled pairs cross species. The strata are too small for per-species intervals. Within the 8,192-bp window delta, the single-token term has a median of 53% of the whole delta's magnitude and exceeds it in a fifth of variants. The window delta ranks variants less like the reference base alone (|Spearman| 0.81 against 0.94), and its downstream terms alone read 0.865.

#### Note S18. The BRCA1 positive control, and the start-codon failure mode it exposes

On the BRCA1 control [71], Evo 2-40B at 8,192 bp reads 0.874 [0.858, 0.889], a site-clustered interval no wider than the Hanley–McNeil interval under full independence, [0.858, 0.890]. On the 2,077 missense substitutions alone it reads 0.822 [0.797, 0.846], and read at the single token, on 1,001-bp windows centred on the variant, the whole panel gives 0.632. Per-residue predicted deleteriousness tracks the measured phenotype at Spearman r_s_ = +0.52 [+0.43, +0.59] (n = 326 residues), and holds within each of the two assayed domains, RING and the BRCA1 C-terminal (BRCT) domain (Figure S6c). The nearest external reference, the Arc Institute notebook for this protocol, prints a zero-shot BRCA1 AUROC of 0.73 for the 1B checkpoint on the loss-of-function versus functional and intermediate split of all 3,893 substitutions. It is a notebook output, not a figure in [31], so it sets a scale, and our 40B result is not a matched reproduction of it. The control also exposes a readout-specific failure: start-codon loss at residue 1 is missed by the window-mean readout here, although Evo 2's authors report large likelihood changes at start codons on average [31], so the miss may be specific to this locus or window. The control validates the 8,192-bp readout, not the single-token one.

#### Note S19. Scoring provenance: checkpoints, scorers, window extraction, reference checks and score orientation

The archived scoring container installed evo2 0.6.0 (Additional file 2, container/). The 8,192-bp mean log-likelihoods come from a scorer that loads each checkpoint whole on NVIDIA H200 GPUs (evo2_1b_base, evo2_7b and evo2_40b) and calls evo2's score_sequences. The 1,001-bp single-token 40B scores of the atlas and of the splicing and ClinVar consequence panels, and the 1B and 7B rungs of Note S10's splicing and ClinVar ladder (evo2_1b_base and the 8k-trained evo2_7b_base), come from a scorer that streams the model's blocks through the GPU. On 3,705 atlas variants of seven species, also scored through NVIDIA's hosted Evo 2-40B service at the same left context, the streamed scores agree with the hosted ones at Pearson r 0.9996 to 0.9998 per species (Spearman 0.997 to 0.9998). The six per-species AUROCs that both classes allow differ by at most 0.0095, in chicken; the hosted scores are not deposited. The 1B and 7B window sums of the splicing and ClinVar consequence panels, and the strand subpanel's 1B rung (Note S29), call BioNeMo's predict_evo2 in 8-bit floating point with a fallback to bfloat16; which of the two ran, and the revision of the converted checkpoint, were not recorded. No Hugging Face revision of any Evo 2 checkpoint was recorded, nor the compute precision of the 8,192-bp scorer; the streamed single-token scorer runs the 40B checkpoint's blocks with their 8-bit floating-point layers.

The single-token readout (Methods) takes its 501 upstream bases from a 1,002-bp window; the model is autoregressive, so the 500 downstream bases are never read. The 8,192-bp readout (Methods) averages the log-probability of every token after the first, given those before it. The upstream positions are identical in the reference and alternate windows and cancel up to numerical noise between passes, so this window delta is the variant's effect at its own position and the 4,095 positions downstream, summed and divided by the window's 8,191 terms.

Unlike the atlas's single-token readout, the splicing, ClinVar consequence and BRCA1 short-window panels use centred windows of exactly 1,001 bp (500 + variant + 500), the variant at index 500 after 500 bases of left context. As scored, every single-token window is effectively its left context (the eQTL window ablation's 2,048-bp rung supplies 1,024 bases of it), so the window-length comparisons of Note S25 are a left-context ladder, a weaker claim than "window size" suggests. The ref_ok build-match gate checked reference alleles base for base against the source assembly, and a deposited check finds all 11,109 atlas windows at 8,192 bp concordant (Additional file 2: reports/atlas_ref_ok.json). That gate is enforced at panel construction for the 8,192-bp panels and the pig negative sets, and the splicing and ClinVar consequence panels drop any window whose base at the variant offset disagrees with the reference allele. For the other 1,001-bp panels, the atlas's among them, it is a recorded column, not a filter, and the BRCA1 short-window builder sets it unconditionally.

The 1,001-bp single-token readout covers the full nine-species panels and carries the reach, calibration, abstention and selection analyses (Tables S16 to S18; Notes S28 and S30) and the eQTL single-token values. The BRCA1 control validates only the 8,192-bp readout (Note S18), so where an 8,192-bp version exists, as for the trust layer (calibration, abstention and conformal prediction; Table S19), that version is the one to read.

**Case and sign.** Both the mean-log-likelihood and the single-token scorers upper-case every window before tokenising, so soft-masked repeats are read as ordinary bases. The atlas table's evo2_meanll_8192 and the BRCA1 table's evo2_meanll_delta are mean_LL(alt) − mean_LL(ref) and must be negated to give logP(ref) − logP(alt), while the eQTL table's evo2_40b_score is already oriented to deleteriousness and is used as stored; negating the wrong one gives 1 − AUROC. The analysis helpers that flip an AUROC below 0.5 upward cannot reveal a score stored with the wrong sign, and they read a near-chance AUROC upward. They never fire on the GERP and phyloP columns: unflipped, GERP's AUROC exceeds 0.5 in all nine species. Additional file 3's recompute scripts orient each atlas score by a fixed sign and reproduce the atlas values, and the eQTL AUROCs of Note S25, several below 0.5, are printed unflipped.

#### Note S20. Extended Methods: comparators and statistical machinery

Two model comparators were run alongside. Nucleotide Transformer v2 500M multispecies [78] was scored as a masked-marginal reference-versus-alternate log-likelihood ratio at the token covering the variant, on the same windows as the single-token Evo 2 readout. It is a masked rather than an autoregressive model, so it sits outside the scope of our claim about autoregressive likelihood readouts. ESM-2 [120] (esm2_t33_650M_UR50D) was run on the protein-coding subset only, as a masked-marginal ratio between wild-type and mutant residue; its reach is in Note S33. The masked-marginal readout is the one Meier et al. [75] establish for zero-shot variant-effect prediction. Neither is a like-for-like competitor: both bound what the atlas comparison means and do not rank models.

To split the readout effect, two terms are formed on a 2,599-variant subset from four quantities, three read from one further 40B pass over the 8,192-bp windows and the fourth from the 1,001-bp pass. The left-context-and-scorer term holds aggregation fixed at a single position and compares the 8,192-bp pass with the 1,001-bp pass (pooled 0.9168 against 0.8792, +0.038); the aggregation term holds context fixed at 8,192 bp and compares the full-window mean with that same single-token delta (pooled 0.9542 against 0.9168, +0.037). By the species mean the aggregation term contributes +0.031 of the subset's +0.067 readout gain, and pooled +0.037 of +0.075 (Table S6), so about half the gain is left context and scorer, not aggregation. The split is incomplete, not path-dependent: one cell of the two-by-two is missing (below). The central-1,001-bp mean of the same pass conditions each term on at least 3,596 upstream bases, so the step to it from the 1,001-bp pass changes left context, scorer and aggregation together (+0.071 on the macro scale; pooled, +0.078), and the step from it to the full window (−0.004; pooled, −0.003) changes only the span averaged. At about 4 kb of left context, averaging over the variant and the 500 bases downstream adds +0.035 on the macro scale. The fourth cell of a two-by-two, a window mean at 501 bases of conditioning context, is not in the design, so how the aggregation gain depends on context is unknown.

Every nine-species mean carries a t interval, because a nominal 95% percentile interval on a mean of nine covers only P(|t₈| < 1.96√(8/9)) ≈ 0.90, about five percentage points short.

The selection analyses used Spearman rank correlation between minor-allele frequency and predicted deleteriousness, expected negative and monotone towards rare alleles. Minor-allele frequency is folded, min(AF, 1 − AF), while the score is fixed reference-to-alternate and never polarised against an outgroup, so where the alternate allele is the major one the two describe different alleles. This dilutes the correlation rather than creating it, so the magnitude should not be interpreted. The AUROC routine, scikit-learn's roc_auc_score, uses midranks for ties, and two independent deposited routines agree with it on every value reported here.

#### Note S21. Evo 2 against conservation at both readouts: per-species margins and scope

At the 1,001-bp single-token readout, Evo 2 and GERP are close: pooled over the 9,532 variants both score (Note S22), Evo 2 scores 0.8805 against GERP's 0.8780; Figure 5d draws the comparison species by species (Table S4), and Figure S7 the within-species percentiles behind it. Per species, on locus-resampled one-sided p-values over the eight species that can be estimated, Evo 2 beat GERP with Benjamini–Hochberg significance in cattle alone (+0.075) and trailed it in human (−0.048 [−0.069, −0.027], loci resampled, uncorrected); goat's +0.173 rests on positives in one locus and has no locus-level p-value.

##### *The 8,192-bp readout: per-species margins, phylogeny and a windowed comparator*

Per species, the advantage of Evo 2 over GERP was tested at both readouts by the two-sample locus-clustered bootstrap and, at 8,192 bp, by one-sided DeLong tests [107] that treat the co-scorable variants as independent and are reported for comparison only. DeLong intervals for the paired Evo 2 − GERP difference on the co-scorable variants, in descending order of margin, are chicken +0.231 [+0.083, +0.380], goat +0.178 [+0.066, +0.290], cattle +0.149 [+0.113, +0.185], human +0.100 [+0.086, +0.114], dog +0.089 [+0.063, +0.114], sheep +0.086 [+0.022, +0.149], pig +0.075 [−0.087, +0.238], cat +0.045 [−0.002, +0.092] and horse +0.023 [−0.032, +0.077]. Six of the nine exclude zero, and seven of nine survive Benjamini–Hochberg on the one-sided test. With positives resampled by 100-kb locus, the intervals of cattle, chicken, dog, human and sheep exclude zero and survive Benjamini–Hochberg, and cat's drops out at a one-sided p of 0.051. The lead's locus-resampled interval therefore excludes zero in five of the eight species where it can be estimated; those of chicken and sheep rest on only 14.0 and 21.5 effective loci, where such intervals are likely too narrow (Note S12). Without chicken, whose GERP track is not the same estimator as the mammals', the macro difference becomes +0.093 (Evo 2 0.937 against GERP 0.844), still favouring the model in all eight remaining species. Alignment depth differs across species, and not only for chicken: at protein-altering positives other than trait alleles the median GERP is 2.18 in human against 0.34 to 0.50 in cattle, cat, dog and sheep, while at the same positives the 241-mammal phyloP's medians run from 6.9 to 7.8 in those five species. By phyloP-241, then, the positives are similarly constrained, and the non-human GERP tracks are the weaker estimators, for reasons these data do not show. Human, which trails conservation by 0.048 at the single-token readout, leads it by 0.100 here, and the cause is the readout, not the panel: GERP is the same in both comparisons (Table S4).

Nine tips will not support a phylogenetic mixed model, and seven of the nine are non-human mammals sharing a single divergence from human, a TimeTree 5 [64] median of about 90 million years, so we cannot give the macro margin a phylogenetically corrected standard error. Averaged within the five taxonomic orders first, the macro advantage over conservation is +0.109 (+0.012 to +0.205), within 0.0002 of its all-nine value of +0.108 but with an interval twice as wide; it is positive in all five orders, and all eight one-representative-per-order choices fall between +0.095 and +0.124.

Without goat, whose nine positives occupy two loci, the 1,001-bp margin, the 8,192-bp margin and the paired change, with t intervals over species, are +0.032 (−0.019 to +0.083), +0.100 (+0.045 to +0.154) and +0.068 (+0.033 to +0.103). With species as the sampling unit, each interval targets the mean over species exchangeable with those analysed and is conservative, in expectation, for the mean over those alone. Resampling loci within each species gives the interval for the mean over these nine fixed species, which inherits the per-species locus intervals' optimism where effective loci are few. The margin is then +0.047 (+0.014 to +0.082) at 1,001 bp and +0.108 (+0.079 to +0.139) at 8,192 bp, and the paired change +0.061 (+0.044 to +0.078); the 1,001-bp interval excludes zero only through goat. Goat's eight co-scorable positives sit in one locus, so its fixed-set replicates resample negatives alone. Without goat, the 1,001-bp margin's fixed-set interval is +0.032 (−0.004 to +0.070).

A windowed comparator. Evo 2's readouts read a window and GERP a single base, so the margins above could reflect the window alone. Averaging GERP uniformly over each variant's own 1,001-bp and 8,192-bp scoring windows, read from the same Ensembl tracks (Table S24), tests that, though these windows are symmetric and the readouts are not (Methods). Against the windowed averages Evo 2 leads by +0.292 (+0.188 to +0.396) at 1,001 bp and +0.409 (+0.288 to +0.529) at 8,192 bp, with 95% t intervals on eight degrees of freedom here and below. Evo 2 at 8,192 bp also leads a cross-validated logistic model on per-base GERP and both window averages, built as in Note S10, by +0.091 (+0.036 to +0.146) on the 9,529 variants with a window value. At 8,192 bp the window averages read below 0.5 in five species (chicken 0.371, horse 0.410, pig 0.465, cat 0.471 and dog 0.484), so they describe the tracks around these alleles and are not the comparator. Regional conservation is a weaker baseline here than the base itself, so against the window averages the per-base margins are the conservative ones. Per base, GERP reads 0.831 by the species mean, against 0.586 and 0.530 averaged over the 1,001-bp and 8,192-bp windows (Table S4 and Figure S18b). The cross-validated model above reads 0.848, against Evo 2's 0.939 on the same 9,529 variants, so the 8,192-bp margin does not come from setting a window readout against a single base.

The readout dependence does not show that one readout is universally standard or that a named published result is overturned, and it does not remove residual composition, clustering or reach uncertainty.

#### Note S22. Resampling units, permutation schemes, seeds, equivalence testing and the fragility of atlas verdicts

Additional file 3's README lists each estimand's resampling unit, replicate count and seed. Where a permutation p-value is reported it is two-sided and family-wise, P(maxfamily |Δpermuted| ≥ |Δobserved|): the share of permutations whose largest absolute difference across the family reaches the observed absolute difference. The ClinVar rungs permute labels within a rung with rung sizes fixed; the splicing classes hold labels fixed and swap the two predictors' within-class rank scores at each variant, the null being that the two predictors are exchangeable on a given variant. Ranks are used there because swapping raw scores on incommensurable scales, an Evo 2 log-likelihood ratio against a bounded probability, inflates the null.

The ClinVar calibration and reach-panel intervals rest on 200 to 300 resamples and are indicative. The calibration arm's headline, calibrated error orders of magnitude below raw error, is not evidence about the scores: a pure-noise score shows the same collapse through the same isotonic pipeline, and a base-rate constant reaches zero calibrated error. It measures the calibrator, and its interval endpoints are imprecise as well. We report the species-clustered percentile intervals because they are still wider and more conservative than the variant-level alternative, not because nine clusters is adequate. A nine-cluster percentile interval undercovers (Note S20), so these intervals are too narrow, erring towards declaring a difference, and the equivalence-test p-values (Additional file 2: reports/tables.md, Table D5) err towards declaring equivalence.

##### *Equivalence testing: the margin, the estimand and the resampling unit*

Where we report the absence of a difference that carries a conclusion, we bound it with an interval or test equivalence by two one-sided tests at α = 0.05, declared only if the 90% interval lies wholly inside ±δ. We know of no published source that sets a meaningful margin for an AUROC difference between variant effect predictors, so we set δ = 0.02 after the data were seen, from the pooled increment Evo 2 adds beyond conservation, and apply it to every contrast (Note S10).

That increment is ΔFM = +0.0202, so the criterion is self-binding: we cannot declare a difference negligible that is larger than an increment we ourselves report as Evo 2's contribution beyond conservation. Sensitivity at δ = 0.01 and 0.05 is in the same Table D5.

The equivalence test is run on the pooled Evo 2 − GERP difference over the 9,532 co-scorable variants, which is +0.0024. The p = 0.28 of the two one-sided tests (TOST) in Table D5 belongs to this pooled difference.

With that estimand fixed, a variant-level bootstrap gives a 90% interval of [−0.009, +0.014] and would declare equivalence; a species-clustered bootstrap gives [−0.024, +0.056] and does not. With species as the unit of generalisation (Note S11), the species-clustered interval applies, so equivalence is not established. The increment from which δ is set has a larger species-level estimand as well (the +0.054 n-weighted species mean; Note S10), but the choice does not carry the verdict: the species-clustered equivalence interval reaches +0.056 and so exceeds both candidate values of δ (0.020 and 0.054). We use the pooled ΔFM because it is the only internally derived scale available, not because it shares the between-species estimand of the test it bounds. On the species-mean difference, the primary estimand, the 1,001-bp margin over GERP on the 9,532 co-scorable variants is +0.0485, with a 90% t interval on eight degrees of freedom of [+0.003, +0.094]: it is not equivalent to zero at δ = 0.02 (TOST p = 0.86), nor at 0.01 (p = 0.92) or 0.05 (p = 0.48), and it is one-sided superior at p = 0.041. Without the three cat positives whose 1,001-bp score is not that of the catalogued substitution (Note S32), the margin is +0.0474 with a 90% interval of [+0.002, +0.093], and the one-sided p is 0.045. The one-sided test and the 9,532-variant set belong to this equivalence analysis and carry no claim of their own.

##### *Fragility of the atlas identification verdicts under locus resampling*

The verdict at the point estimates, identification in Evo 2's favour at 8,192 bp in six of nine species, holds in 100.0% of locus-resampled replicates in cattle, human and dog, in 99.8% in chicken and in 93.5% in sheep, whose margin over the bound runs from −0.010 to +0.076 between the 2.5th and 97.5th percentiles; goat, on two loci, is not estimable (Note S12). These verdicts charge Evo 2 only for its 21 unscored variants at 8,192 bp, a lower bound on its declines (Note S32). Of the eight species where it can be estimated, the same verdict holds at 1,001 bp in two of them (cattle and chicken).

#### Note S23. The comparison is not measured on the same variants: reach, its cause and identification

GERP reaches the two classes unequally, beyond sampling error in four of the nine species (Table S1), so a per-species conservation AUROC can rest on a class-skewed subset of the panel it is quoted for. We establish the skew within our own nine panels and do not survey the published literature.

The must-answer premise sits beside a method that itself declines to answer, so the two kinds of silence must be distinguished. Conservation's silence is involuntary, usually unreported and class-shaped. The selective abstention layer (Note S28) declines by design: its silence is chosen, its answered fraction is fixed in advance, and the answered set carries a measured error rate. It is not class-symmetric either: refusing each species' least-confident 15% retains 0.33 to 0.82 of positives against 0.86 to 0.90 of negatives. That is a mean class gap of 0.331 against conservation's 0.141, and ours is the larger of the two in eight of nine species. The must-answer penalty on that arm is 0.158 by the species mean, against the 0.102 we charge GERP. At equal reach the must-answer penalty charges a better-discriminating score more (Methods). We read conservation's as a cost and ours as a design choice not because our refusal falls evenly on the classes but because a stated refusal rate is part of a decision procedure, while an undeclared one silently changes the panel a published accuracy refers to.

Evo 2's per-species margins over GERP, pooled with Note S14's random-effects method and leaving out goat, give +0.092 [+0.052, +0.132] (I² 63%) at 8,192 bp and +0.016 [−0.032, +0.064] (I² 77%) at 1,001 bp. If declined variants are no easier than scored ones, GERP's bound narrows to [ρAcov, Acov], which Evo 2's 8,192-bp lower bound clears in all nine species, by as little as 0.007 in horse (0.922 against 0.915).

The interval that matters for a reader in a tenth species is the prediction interval. On the locus-resampled pool it is [+0.001, +0.183] at 8,192 bp, which assumes a further species exchangeable with these eight and includes zero when any one of five of them is left out; its lower limit clears zero by less than its Monte Carlo standard error. At 1,001 bp it is [−0.100, +0.132].

##### *The unresolved cause of class-dependent missingness*

**The controls below do not establish why the missingness is class-dependent: a cruder coding/non-coding stratification finds the class gap surviving in four of the eight species in which it is defined (pig, horse, cat and cattle; goat has no stratum holding five variants of each class), but a consequence-matched analysis finds a positive class gap separable from zero in none of the five testable species. We privilege the matched arm because the unmatched one confounds class with consequence.** The obvious explanation is that positives are overwhelmingly coding (84.6% to 100% across all nine panels) while negatives outside human are sampled from population variation, so the class gap could be a coding-versus-non-coding artefact rather than a property of the conservation track. We tested it with a consequence-matched risk difference: Mantel–Haenszel-type stratum weights [129] over consequence categories (each variant's full annotation) holding at least five variants of each class, a stratified Miettinen–Nurminen score interval [130,131] and a score test of homogeneity across categories. Horse falls from +0.489 marginal to +0.161 [−0.055, +0.514], cat moves from +0.194 to +0.239 [−0.153, +0.276] on a single intronic category in which all six positives are reached against 418 of 549 negatives, cattle reads +0.032 [−0.016, +0.112], and human, the only well-powered stratum, is a clean null at −0.003 [−0.028, +0.029]. Dog's +0.009 turns to −0.120 [−0.228, −0.037], but its categories disagree (score test of homogeneity 34.4 on 4 degrees of freedom) and the reversal comes from its intronic category, a property of that category rather than a pooled effect. Within class, the GERP-missing variants do not differ from the GERP-scorable ones in Evo 2 score (negative-class rank-biserial, pooled across species, +0.012 [−0.021, +0.046], p = 0.47; positive class +0.057), so, pooled, the unscorable variants are not a subpopulation the model itself distinguishes. For the negative class the species mean of the same statistic is −0.107, negative in eight of nine species, and the intervals exclude zero in cat (−0.166 [−0.242, −0.085]) and dog (−0.118 [−0.212, −0.022]), so in those two species the negative-class statistic is not null. A runs test cannot settle the cause either. On the GERP no-call pattern of the atlas negatives it gives z = −1.28 in pig and −1.83 in horse, at overall no-call rates of 57.6% and 51.5%, close to unclustered, against −7.31 in cattle, −10.86 in dog and −13.87 in human at rates of 4% to 9%. But adjacent negatives lie a median 2.42 Mb apart in pig and 2.43 Mb in horse, far beyond the lengths of alignment gaps, so the test has no power at their scale. Three things follow. First, reach as we use it is a property of the deposited annotation for a species, not an intrinsic property of the conservation method, so the deployment claim is about the tracks a user can actually download. Second, the control is close to uninformative rather than reassuring: only five of nine panels have a matchable stratum at all, and "not separable from zero" here reflects power, not evidence of absence. Third, the cause decides generalisation: if the mechanism is a coding/non-coding artefact, the deployment penalty restates "positives are coding" rather than being an independent property of conservation scoring, and we cannot exclude that. The marginal gap is real and its deployment cost is real; its cause is not resolved by these data.

#### Note S24. Orderings stated in published benchmarks: a systematic search

One PubMed query returned 75 records; Table S33 gives their flow to the 17 that print per-predictor AUROCs on clinically labelled variants. Ten of the 17 state an AUROC ordering while reporting scored counts or coverage [72,73,132–139]; seven do not (Table S33). A stated ordering is identified when the leader's lower bound clears the other's upper bound for every class split the printed counts admit, not identified when it clears for none, and dependent on the split otherwise. In the ten, 1,702 of 2,842 stated orderings are identified and 1,045 are not (36.8%); 92, all in Barbosa et al., depend on the split and 3 are not assessable (Table S34). Printed AUROCs are taken as exact; allowing each two-decimal AUROC its rounding, 52 of Barbosa et al.'s 961 identified orderings and 1 of Montenegro et al.'s 16 are not identified for every split; taking each benchmark's share not identified, on clinical panels only, Liu et al. contributing their ClinVar panel's 7, the per-benchmark median is 19.8%. The main text gives the pooled counts under both readings.

Barbosa et al. rank 36, 34 and 34 tools on three panels and Heo and Kim 28 methods on one, supplying 2,070 of the 2,842 orderings (73%) and 911 of the 1,045 unidentified ones (87%); without them the count is 134 of 772 (17.4%). Heo and Kim's 378 orderings rest on a legend sorted by AUROC: their text states 52, all identified, which gives 773 of 2,516 (30.7%); Figure S12c shows this reading. Per benchmark, nine of the ten state an unidentified ordering, six under the strictest reading of three of them (Table S34). The extraction was checked against the sources (Table S33).

Wang et al. [132] rank FATHMM-XF first at 0.8033 on non-coding ClinVar variants, ahead of FATHMM-MKL at 0.7954 and ReMM at 0.7848, both of which score every variant. FATHMM-XF scores 446 of 515 pathogenic and 1,688 of 1,850 benign variants, the one split its printed metrics allow, so charged for the rest it could lie anywhere from 0.63 to 0.84, and neither ordering is identified.

A second set, selected by hand, is reported apart (the hand-selected set of Figure 6b; Figure S12a, b). Of its 28 screened articles, 16 print AUROCs [4,7,132,133,140–151]. Of these, eight rank predictors by AUROC while reporting how many variants each predictor scored. In seven of them, the bound leaves at least one stated ordering unidentified: 23 of 330 in Li et al. [133], 18 of 66 in Wang et al. [132], 3 of 20 in Rastogi et al. [7], 7 of 192 in Radjasandirane et al. [4], 14 of 44 in Barbosa et al. [141] (not the Barbosa et al. [72] of the systematic set), 3 of 29 in Riepe et al. [145] and 4 of 12 in Strauch et al. [143]. Across the eight, the bound does not identify 72 of the 697 stated orderings (10.3%); it identifies 589, and 22 are not assessable. Where the printed values leave the class split open, 14 stated orderings, 1 in Rastogi et al. and 13 in Radjasandirane et al., depend on it. With the six benchmarks that only this set holds, the ten make the 16 of Note S40; joined, an article in both sets counts once.

#### Note S25. The pig cis-eQTL panel and its near-chance result

Three controls constrain, but do not isolate, the source of the near-chance result on the pig cis-eQTL panel (AUROC 0.488). First, the three released checkpoints do not close the gap: from Evo 2-1B to Evo 2-40B, the eight-species atlas macro gains +0.043 (Note S10), while the eQTL effect size |AUROC − 0.5| on the 2,000-variant subpanel at 8,192 bp falls from 0.008 to 0.002, intervals overlapping (Table S2). Second, context alone was not the dominant factor on the same 2,000-variant subpanel: at a fixed single-token readout, widening the window from the 1,002-bp window of the 1,001-bp readout (Note S19) to 2,048 or 4,096 bp moves AUROC from 0.485 to 0.487 or 0.484, while switching to mean log-likelihood at a fixed 1,002-bp window moves it to 0.496; both effects are small (Additional file 2: logs/eqtl_ablation.md prints each as one minus these values, oriented above one half). Third, all tested sequence-based scorers were near chance on that 2,000-variant subpanel, which is not matched within eGene: Evo 2-40B, Nucleotide Transformer v2 (NT-500M) [78] and three GERP flavours (Figure S2d). Distance to the transcription start site, a predictor using no sequence and, with the association statistic |z| excluded as circular, the only non-circular positive control on this panel, separates candidate causal variants from their within-eGene controls at AUROC 0.651 (95% CI [0.641, 0.660]; n = 19,341). The labels therefore carry some recoverable signal, which bounds the comparison without settling it: that a trivial positional baseline reads this panel at 0.651 and a 40-billion-parameter sequence model at 0.488 is informative about the model. But 0.651 is modest, so a capability gap cannot be cleanly separated from labels that carry little signal; neither reading is dismissed, and we do not claim the model ought to have matched a positional baseline. Reference [31] reports Evo 2-40B, scored zero-shot on the DART-eval regulatory tasks, at AUROC 0.58 for chromatin-accessibility quantitative trait loci (caQTLs) and 0.66 for DNase-I-sensitivity QTLs, above Nucleotide Transformer (0.52, 0.61) and below the sequence-to-function model ChromBPNet (0.77, 0.89). We therefore report the eQTL result as a null at the single-token readout on this panel, possibly specific to it, not as a demonstrated failure of the model.

On the 20,000-variant panel Evo 2's deviation from chance, at AUROC 0.488, is statistically *resolvable:* a permutation at 20,000 resamples with labels permuted within eGene returns p = 0.00025 against a null mean of 0.4991 (standard deviation 0.0034). The deviation from chance, |AUROC − 0.5|, is 0.012, against 0.378 for the whole atlas at the same 1,001-bp single-token readout (nine-species mean 0.878, Table 1). Because the confounds left uncontrolled below, the choice of controls by PIP and the classes' GC content, repeat context and mappability, could each move an AUROC by more than 0.012, the magnitude claim is what survives, near chance and more than an order of magnitude below the atlas's, not the sign of the deviation or its nominal precision.

##### *Scope across checkpoints and architectures, and the confounds*

Evo 2-40B reads the atlas's non-coding disease and trait alleles at 0.852 at the 8,192-bp readout (Figure 5g), so, with its own paper's caQTL and DNase QTL readings above, the eQTL null is not blindness to non-coding sequence in general. The evidence is one autoregressive family at its three released checkpoints plus one masked-language comparator (NT-500M in Figure S2d), not a survey of a model class.

The control set is not neutral: selecting controls at a posterior inclusion probability (PIP) of 0.001 or less within the same eGene selects variants that fine-mapping confidently excluded, which are systematically those in low linkage disequilibrium with the candidate causal variant or in regions carrying no association signal. The panel is not matched on allele frequency, and Evo 2 scores correlate with frequency by our own label-free analysis, but the two classes do not differ in frequency. In the PigGTEx genotype files (Science Data Bank, doi:10.57760/sciencedb.08089), which keep only sites with a minor-allele frequency of at least 0.05 in each tissue, candidate causal variants and controls have indistinguishable minor-allele-frequency distributions (median 0.241 against 0.243; Mann–Whitney p = 0.98), and frequency alone reads AUROC 0.500 (95% CI [0.487, 0.512]; eGenes resampled). With each candidate causal variant matched to up to three controls within 0.005 in frequency (4,836 of the 5,000 matched), Evo 2 reads 0.487 (95% CI [0.478, 0.497]). Of the 5,000, 2,217 (44.3%) lie in eGenes that hold no control; restricted to the 1,430 eGenes that hold both classes, Evo 2 reads 0.478, and 0.475 on pairs within an eGene. Matched on both frequency and distance to the transcription start site (3,839 of the 4,844 with both values), Evo 2 reads 0.487 (95% CI [0.476, 0.497]), while distance alone falls to 0.500 on the matched set (Additional file 2: reports/tables.md, Table D6). Nor is the panel matched on GC content, repeat context or mappability.

#### Note S26. At the readouts tested, competence stops at a variant class, not at a genomic region

Table S20 sets Evo 2-1B, summed over a 1,001-bp window, against purpose-built splice predictors by variant class: it reads 0.853 on essential-splice variants but 0.471 to 0.721 on the other four classes, below SpliceAI in each. Its pooled DeLong difference against SpliceAI (p = 1.65 × 10⁻⁴⁸) treats 3,912 variants in six genes as independent and is not relied on; by assay, Evo 2 trails SpliceAI in six of six assays (Table S13). At this checkpoint and readout, essential-splice variants, a class with eight negatives (Note S2), read higher than the other four classes, a contrast described, not tested. The pattern fails at 40B's single-token readout; whether it holds at 40B and 8,192 bp is open, since on the BRCA1 assay that readout nearly matches the specialists.

The ClinVar consequence panel argues against the simplest explanation, that regulatory sequence is harder: read by Evo 2-40B at the single token, as the eQTL panel is, its 3′ and 5′ untranslated-region (UTR) variants give 0.649 and 0.692 against the eQTL panel's 0.488. On human complex-trait variants Evo 2-40B reads 0.516 in the orientation TraitGym's workflow declares for a log-likelihood ratio (0.484 as distributed), within 0.02 of chance either way. Conservation (phastCons-43p) reads 0.617 on the same panel, so the near-chance reading concerns what a zero-shot likelihood readout recovers, not whether the panel is learnable. Location does not separate these cases; one difference between them is that ClinVar's UTR entries are rare alleles of large effect and the eQTL and complex-trait entries are common variants that modulate expression. The complex-trait entries are TraitGym's test split [47], 11,400 variants with 1,140 causal, read from the scores TraitGym distributes, not rescored here.

The ClinVar panel resolves the consequence gradient that the 18-variant OMIA regulatory rung below tests only weakly: all eight of its classes are resolved above chance under Westfall–Young family-wise control, each by more than its minimum detectable difference; the panel's intervals and permutations take variants, not genes, as their unit. At Evo 2-1B, summed over a 1,001-bp window (Table S15), discrimination runs from 0.850 on missense to 0.650 on 5′ UTR variants.

The gradient runs unevenly (Table S15): synonymous variants read 0.790, above nonsense at 0.746, and ClinVar's pathogenic synonymous entries are enriched for splice-disrupting alleles, so a decay from nonsense to regulatory does not survive this panel.

##### *The OMIA consequence gradient and its lowest interval*

Scored against each species' own trinucleotide-matched negatives, with per-species AUROCs combined as a positive-count-weighted mean, the positives of the seven species whose OMIA records carry a consequence annotation (not human or cattle) show an ordered gradient by molecular consequence in Evo 2-40B discrimination at the 1,001-bp single-token readout. Per-consequence AUROC fell from nonsense (0.954, n = 112, 95% CI [0.922, 0.981]) to missense (0.896, n = 324, [0.869, 0.922]), splicing (0.772, n = 70, [0.713, 0.833]) and regulatory variants (0.579, n = 18, [0.504, 0.652]), with variant-level bootstrap intervals stratified by species, which ignore locus clustering and are likely too narrow. With the negatives fixed across classes, the gradient is not a property of the null, though GERP, run through the same estimator on the variants it reaches, also reads lowest on the regulatory class (0.720; Table S11).

#### Note S27. Readouts that reorder models on an independent benchmark

For its Mendelian test set we read 34 of the zero-shot score files TraitGym distributes, readouts of 17 models [47], each oriented as its workflow declares. They are the log-likelihood ratio and its absolute value for each genomic language model, a precomputed embedding Euclidean distance or inner product for five of them, and one readout each for Enformer, Borzoi, CADD, Sei and three conservation tracks. TraitGym also distributes, as prediction files, the inner product, cosine distance and Euclidean distance between the two alleles' embeddings for all ten language models; we did not read these. Its metric is average precision within each chromosome, weighted by the chromosome's share of variants; our values equal its leaderboard's for 12 of the 17 models, and for the other five (AIDO.DNA, Caduceus, Evo 2-1B, Nucleotide Transformer and SpeciesLM) the leaderboard prints one of these embedding readouts, so those five are compared on the 34 score files only. By the same metric Evo 2-40B reads 0.557 against 0.527 for phastCons-43p, 0.655 for phyloP-100v and 0.696 for phyloP-241m. By AUROC, Evo 2-40B's log-likelihood ratio reads 0.8495 in the orientation the workflow declares (0.1505 as distributed), against 0.8594 for phastCons-43p.

Across every pairing of one readout of each model, 26 of the 136 model pairs change order, all among the 115 in which either model offers a choice of readout. In 7 of them both orders are resolved, uncorrected for the pairings compared, with standard errors from 2,000 resamples of the 19 chromosomes. Each of the 7 involves the Genomic Pretrained Network with Multiple-Sequence Alignment (GPN-MSA), and in each the pairings that put GPN-MSA behind read it by Euclidean distance or inner product. For example, Borzoi leads GPN-MSA read by Euclidean distance by 0.229 but trails GPN-MSA read by its log-likelihood ratio by 0.259. The count is 5 with a Bonferroni correction over the pairings within each pair, 10 with percentile intervals in place of standard errors, and 9 when the causal variants' loci are resampled, each matched group drawn with the causal variant it was matched to. The causal variants chain into 107 clusters within 10 kb, and the 9 pairs resolved both ways are the same with the chaining widened to 100 kb or 1 Mb.

#### Note S28. Calibration, abstention and conformal prediction across species

A log-likelihood ratio has no probability scale. This Note's arms are at the single-token readout, which the BRCA1 control does not validate (Note S18); Table S19 rebuilds the trust layer at 8,192 bp. Min-max rescaled within each species, a constructed baseline, Evo 2 scores are miscalibrated (expected calibration error, ECE, 0.318, species mean, 1,001-bp readout), and leave-one-species-out isotonic transfer [152] appears to repair them, to 0.053 (Figure S4b; Table S16); Figure S1 shows the transferred two-parameter map species by species.

Transferring a Platt map [153] across species is, by construction, the same operation as fitting one global two-parameter sigmoid on the pooled donor species. Whether a flexible isotonic calibrator beats those two global parameters is not resolved: of the four estimators bootstrap-tested, three cannot distinguish them and the fourth (equal-mass 15) favours transfer, so any advantage is bounded above by about 0.014 ECE, with no equivalence margin set on that scale. The null is not a sample-size artefact (Table S16), and the beta family [154] does not rescue it. A calibrated probability therefore carries no species-specific warrant in an unlabelled target; an in-species oracle calibrated on target labels is roughly 3.4-fold better on macro ECE (0.0157 against 0.0533), but only in sample, which does not establish prospective performance.

Refusing each species' own least-confident 15% of calls withholds 1,673 of 11,130 variants and removes 316 of 893 errors (35.4%; Table S17; Additional file 2: reports/tables.md, Tables D10 and D12), a 2.35-fold enrichment over random refusal with a macro mean lift of 3.42 (Table S17).

Macro selective error on the isotonic posterior falls from 0.058 at full coverage to 0.038 at 85% coverage and then flattens, while a random-refusal control stays between 0.051 and 0.059 (Figure S4c; Additional file 2: reports/tables.md, Table D2c), but over the whole curve the ordering is much weaker: on the Platt posterior, pooled over species, the risk–coverage area is 0.0784 against 0.0802 for random ordering, and the ordering is worse than random in two of nine species (Additional file 2: analyses/results/risk_coverage.json).

The 85% knee is in-sample, located on the panel that reports its benefit, but re-selected on eight species and re-measured on the ninth it returns 85% coverage for all nine. Human contributes 61% of the errors while carrying one-ninth of the macro weight, and within human the error-detection ordering is at chance. The aggregate lift is real: a within-species permutation preserving each panel's size and error count places it outside all 10,000 null draws (p < 0.0001; Additional file 2: reports/tables.md, Table D11). Human's showing tracks its 1:1 composition, not its labels: down-sampled to a 10:1 composition, a composition control rather than a prevalence-corrected re-estimate [155,156], its refusal lift rises from 1.71 to 3.19 and its error-detection AUROC from 0.474 to 0.708 (Table S17; Additional file 2: reports/human_prevalence_check.json).

A class-conditional (Mondrian) conformal layer [157,158], calibrated on the six label-rich species, reaches its nominal 90% positive-class coverage at the 1,001-bp readout in two of the three held-out species by abstaining on 50.5% and 62.7% of their calls (Figure S4e; Additional file 2: reports/tables.md, Table D2d); the third, pig, at 0.889 (32 of 36; exact 95% interval 0.739 to 0.969), is not an established shortfall. At 8,192 bp the same layer gives no variant a two-label set but returns an empty set, which glmtrust's decision policy counts as an abstention, for 368 of the 11,109 variants (3.3%). Its class-conditional guarantee holds pooled over the six calibration species (positive-class coverage 0.922) but not species by species, and it does not transfer: coverage falls to 0.444 in goat, 0.714 in chicken and 0.694 in pig, each exact 95% interval below the nominal 0.90. These intervals treat positives that share loci as independent, so all three may be too narrow, and goat's, on two loci, is descriptive only. Exchangeability fails by construction across species [159]. Conformal risk control, a threshold λ set on the label-rich species to hold the positive-class false-negative rate at α and then transported, holds strictly, at an achieved rate of at most 0.05, in five of nine species when at most 5% of positives may be missed (seven within the 0.02 tolerance of Additional file 2: logs/crc_certificate.md), goat and chicken of the three held out among them, and misses in pig, at 0.111, while human holds only by calling 3% of variants negative. That threshold λ is set on the same positives the isotonic map was fitted on, with no separate calibration split, so its finite-sample correction is nominal and gives no guarantee.

At the default threshold, before any abstention, the model misses about two of every five catalogued positives in the eight non-human species by the species mean, one in three weighted by positives (Table S18), and abstention suppresses false alarms far better than it recovers missed positives (Additional file 2: reports/tables.md, Table D12). We do not claim that abstention detects the variants the model reads near chance, such as those of the eQTL panel.

##### *The cost matrix behind the always-abstain result*

Under an asymmetric cost matrix [156] charging 10 for a missed positive, 1 for a false alarm, 0.5 for an abstention and nothing for a correct call, always abstaining cost 0.500 per variant pooled and beat every learned decision cell, including the best Evo 2-based rule (0.536). That verdict is aggregation-specific, because expected cost is prevalence-sensitive: by the unweighted species mean, the paper's primary aggregation, the same Evo 2 policies read 0.447 (raw threshold), 0.505 (reject option) and 0.487 (Mondrian conformal) against the 0.500 floor. The conformal cell depends on its calibration split: over split seeds 0 to 99 its median cost by the species mean is 0.489, below the floor in 97, and the split behind the 0.536 and 0.487 above is the most favourable of the 100. The always-abstain result is specific to pricing a miss at ten false alarms and to the single-token readout: at 8,192 bp, with donors holding at least 50 positives, a reject option and a conformal rule each cost 0.141 per variant pooled, against 0.500 for always abstaining, and on the deposited posterior, whose donors are every other species, the reject option costs 0.144 against 0.146 for a cost-aware threshold that never abstains, a small gain.

##### *The trust layer rebuilt at the 8,192-bp readout*

At the 8,192-bp readout the layer improves on every axis (Table S19). The arm the package ships is exposed to ties at the refusal boundary: its isotonic posterior takes 218 distinct values, 597 variants sit on the boundary, and permuting the row order moves the pooled lift between 3.91 and 4.26. Like for like, isotonic at both readouts (Additional file 2: reports/tables.md, Table D9), errors fall from 861 to 415, so 32 of the 478 fewer errors come from the calibrator rather than the readout. Two 1,001-bp conclusions change at this readout: that calibration transfer buys little over one fitted sigmoid, and that abstention does not beat always abstaining when costs are pooled. With donors holding at least 50 positives, transfer lowers macro ECE from 0.0285 for an unpenalised pooled two-parameter sigmoid to 0.0211 (difference −0.0074 [−0.0133, −0.0040]); all four bootstrap-tested binning estimators favour transfer, which is better in six of nine species by calibration error, and admitting every other species as a donor gives 0.0209 against 0.0279.

Specification. Raw Evo 2 scores were mapped to probabilities by isotonic regression [152], leaving one species out, with target scores outside the donor score range clipped to that range. At 1,001 bp a donor needs at least 50 positives, admitting six species, so goat, chicken and pig never donate; at 8,192 bp every other species donates, as in the packaged tool, except in the results stated with donors holding at least 50 positives, which keep the 1,001-bp donors. The uncalibrated ECE and the raw-threshold cost policy instead rescale each species' score to 0 to 1 by its own minimum and maximum; neither is a calibration.

ECE is assessed on the isotonic map and per-variant selective prediction on a two-parameter Platt posterior [153] fitted on the same pool, because the isotonic posterior's 248 distinct values across 11,130 variants leave 99.63% of them in tie blocks, so the 15% boundary would be set by sort order, not confidence. Prevalence-corrected probabilities use Elkan–Saerens prior correction [155,156], mapping donor-pool prevalence to the held-out panel's own construction ratio, which is not a deployment prevalence: the target is 0.091 in seven of the nine rows (cat 0.092, human 0.500), the 10:1 ratio the panels were built at. The correction lowers macro ECE to about 0.022 over the eight non-human species and 0.033 over all nine (Additional file 2: reports/tables.md, Table D8), though not uniformly: chicken worsens (0.020 to 0.025) and human is unchanged (0.117 to 0.118).

Conformal prediction used split-conformal inference [157,158] in a class-conditional (Mondrian) construction, calibrated on one half of a stratified 50/50 split of the pooled label-rich species and applied unchanged to zero-label targets. The class quantile uses α = 0.10 within each class, the score being the probability given to the wrong class; a prediction set holding both labels is an abstention and one holding neither is empty.

The selective predictor ranks calls by the confidence |2p − 1| and withholds each species' least-confident 15%; an error is a call on the wrong side of 0.5.

#### Note S29. Strand consistency at the 1B and 40B checkpoints

A genome has no preferred strand, so scoring a window or its reverse complement is a free protocol choice.

To hold panel and window length fixed, the 1B model was scored on the 40B run's 327-variant, 8,192-bp panel in both orientations (Additional file 3). The two rungs were scored by different implementations (below), so capacity is not the only variable. The 327-variant subpanel is 51.7% positive and is stratified at about 40 variants per species; pig contributes 20, all of them positives.

**Strand agreement and the strand-choice shift at 1B and 40B**

|  | **1B** | **40B** |
| --- | --- | --- |
| Pearson, forward against reverse complement | 0.7732 | 0.7123 |
| Spearman, forward against reverse complement | 0.8837 | 0.8992 |
| strand-choice AUROC shift | 0.0081 | 0.0079 |
| as a percentage of +0.096, the random-effects pool of the nine per-species DeLong 8,192-bp margins (Additional file 2: analyses/results/meta_8192.json) | 8.4% | 8.2% |
| gain from strand-averaging | +0.0073 | +0.0031 |
| forward-strand AUROC | 0.8985 | 0.9238 |

The 1B column cannot separate strand from implementation: its two strands agree (Spearman 0.8837) about as well as two 1B implementations agree on one strand (0.82 to 0.97 within species; Note S10), so only the 40B column measures strand, and there the shift is about 8% of the 8,192-bp margin. Strand-averaging improves discrimination at both scales. The table is the full-window readout, the only one the two rungs share: the 1B rung was scored by the window-sum scorer. On the same 327 windows the 40B run also gives the mean over the central 1,001 bases, where the strand-choice shift is 0.0046. The AUROC shift is small at both window averages and unresolved at the full window (below), but per variant the two strands' scores agree only as tabulated above; an architecture that is reverse-complement equivariant by construction [160] cannot differ between strands at all. The reach, calibration, abstention, eQTL, selection and 1,001-bp margin arms are scored on the forward strand at the single-token readout, so they carry a strand sensitivity measured here at the window readouts and not at their own.

Two deposited runs at the 1,002-bp window mean, on a 389-variant subset at Evo 2-1B and at Evo 2-7B, favour the reverse complement instead. That subset is not an atlas subset, and its per-variant scores are not deposited. On it forward minus reverse complement reads −0.0111 at 1B and −0.0121 at 7B, and forward minus the strand average −0.0183 and −0.0282. Resampling the 327 variants, the 40B strand-choice shift at the full-window readout is +0.008 (−0.007 to +0.024), of the order of the run-to-run floor. Two 40B passes over the same windows differ by at most 0.000125 per variant and move per-species AUROC by up to 0.012 in goat, the smallest stratum with both classes.

#### Note S30. The label-free cattle analysis conditioned on conservation, and the cross-clade bat pilot

The label-free cattle panel is exploratory (Note S32), and it keeps a frequency–deleteriousness relationship within four of five conservation quintiles. Conditioning the cattle gradient on conservation gives, without a single label, a pattern consistent with the competence boundary seen on the eQTL panel, a weaker claim than recovering the eQTL-panel null (Figure S3b). With the variants partitioned into phyloP quintiles, on the 27,234 of the 28,006 variants phyloP reaches, rarer variants still score as more deleterious at the same phyloP, so the gradient is not merely a conservation artefact. Its strength orders with conservation (Figure S3b), from a Spearman correlation of −0.018 [−0.044, +0.011] in the least-conserved quintile to −0.118 [−0.143, −0.093] in the most conserved (bootstrap intervals over 500 resamples; Additional file 2: reports/fig6_free.json, selection_conditioned.strata), though adjacent quintiles are not separable, so we claim no monotone trend across them, and its confidence interval crosses zero in the least-conserved quintile, where conservation itself carries little signal. Label-free here means free of curated pathogenicity calls, not of annotation: phyloP is alignment-derived. The least-conserved quintile is mostly neutral sequence, where no gradient may simply be correct, and phyloP quintiles proxy constraint, not regulatory status, so we read the flattening as a limit on where the score is informative, not a demonstrated regulatory failure. Alignment-based constraint may also under-mark fast-turnover regulatory elements. The label-free and eQTL arms are not independent: both are Evo 2 scores, with the boundary drawn by phyloP in one and by SuSiE PIP in the other. Nor is the label-free arm a reproduction of the eQTL result in a second species.

A wild bat (*Myotis lucifugus)* with no curated variant resource shows a frequency–deleteriousness relationship of the same sign (r_s_ = −0.125 [−0.138, −0.113], over 23,888 variants that take only 32 distinct minor-allele frequencies; Figure S3d). Its ten libraries, four of them DNA pools, set an allele-count floor (no frequency below 0.05 is observable), so it is not a replication of the cattle result.

#### Note S31. Further limitations

(1) Panel size is confounded with alignment quality in these data, and the supervised baselines are point estimates without paired intervals. (2) Negatives are not matched on breed or discovery cohort, so every atlas number keeps a residual composition signal (Note S14). (3) Label quality was audited on the human panel only, and the non-human positives were not stratified by evidence tier. (4) Evo 2 was pretrained on sequence without variant-effect labels [31]; whether its corpus holds other genomes of these species, which would show it population negatives' alleles more often than catalogued disease alleles, was not checked.

#### Note S32. Panel provenance: sources, construction and exclusions of every panel

**Cross-species disease atlas.** The eight OMIA species contribute 740 curated positives; candidate records were set aside when they failed the reference-allele build gate, were not single-nucleotide variants or had no symmetric scoring window at a contig end. The atlas therefore keeps only variants Evo 2 could score with a full window: its reach at 1,001 bp is 1 by construction, and its 21 declines at 8,192 bp are a lower bound. OMIA is literature-curated, and many of its entries rest on a single small study. Of the 1,495 pathogenic or likely pathogenic human positives, 1,357 carry at least one review star, a filter the ClinVar panels apply and the atlas does not. The five with a mixed or uncertain classification are kept because the panel is deposited as drawn; at 0.3% of the class they cannot move its AUROC by more than 0.004. Four of the reference assemblies are superseded in Ensembl: chicken GRCg6a, sheep Oar_rambouillet_v1.0, cattle ARS-UCD1.2 and dog CanFam3.1. We used the builds matching the OMIA and Ensembl records, so no atlas variant is lifted between builds of its own species. In two cat positives the identifier carries an accession other than that of the chromosome OMIA records, D4's for a DMD allele on X (NC_058380.1_27099390_G_A) and one outside the cat assembly for a PAX3 allele on C1 (NC_051841.1_205787310_G_A); each was scored, annotated and read for conservation at the chromosome OMIA records (Additional file 3: tables/atlas_variant_positions.parquet). The assemblies of Table S24 are cited as cattle [161], pig [85], sheep [162], goat [163], horse [164], chicken [165], dog [166,167] and cat [168]. Two citations are approximate: [162] describes ARS-UI_Ramb_v2.0, the successor of the sheep build used, and [165] the original chicken genome, not GRCg6a.

**Human ClinVar benchmark.** The trust layer was tested on frozen scores, CADD, REVEL and AlphaMissense, on a random 150,000-variant subsample of about 1.6 million ClinVar variants, of which each score answers 19,193 to 30,993, with isotonic calibration fitted under 4-fold cross-validation and a coding/non-coding split. The benchmark file is the EVEE Zenodo shard [117].

**Independent regulatory panel (PigGTEx cis-eQTLs).** The panel draws on pig cis-eQTLs fine-mapped by PigGTEx across 32 tissues, with the PIP as the candidate causal label. The two PIP thresholds apply to the maximum PIP across the 32 tissues for each variant–eGene pair, so the classes are not symmetric: a positive needs 0.9 in any one tissue, admitting winner's-curse false positives, while a control must stay at or below 0.001 in every tissue. Contamination of the positive class pushes discrimination towards chance, which is the direction of our own headline null, so this construction works in favour of the result we report. Ranking on the unsigned magnitude |score| instead of the signed score gives AUROC 0.496, also near chance. A separate 2,000-variant subpanel, the first 1,000 candidate causal variants and the first 1,000 controls in variant-identifier order, carries the 8,192-bp readout for Figure S2's matched-readout contrast (Note S25). An identifier begins with its chromosome, so the subpanel is not representative: its candidate causal variants lie on chromosomes 10 to 14 and its controls on chromosomes 10 to 12, and 102 of its 626 eGenes hold both classes.

**Label-free selection panels.** The cattle panel was stratified-sampled across five minor-allele-frequency bins to 28,006 windows (Table S3), with purifying selection as the label-free ground truth: deleterious alleles are held at low frequency, which is the relationship the site-frequency spectrum encodes [169–171]. Its frequencies come from the FarmGTEx cattle imputed-genotype release [115] and are computed over samples, some repeated tissues of one animal, rather than individuals, so they are least accurate in the rarest bin and the analysis is exploratory. The windows lie on chromosomes 1 to 22 only, so cattle's autosomes 23 to 29 and its X chromosome are not represented. The panel is not trinucleotide- or mutational-context-matched, and rare variants differ from common ones in mutational spectrum, so a context effect alone could produce a frequency–deleteriousness gradient. Yet across full trinucleotide context, the 96 pyrimidine-referenced classes, each holding at least 43 variants, the Spearman r_s between frequency and deleteriousness_ is negative in 91 of the 96, with an n-weighted mean of −0.102 against the pooled −0.099. Stratifying rather than matching, this control shows that context composition does not produce the gradient, not that context plays no part in the score. Genotyping error, most frequent among the rarest alleles, is not excluded, as no genotype-quality filter is applied, and imputation error, also largest for the rarest alleles, is a further untested alternative. The bat pilot of Note S30 used reads of wild little brown bats (*Myotis lucifugus;* BioProject PRJNA624023) mapped to Myoluc2.0 (GCA_000147115.1), an assembly of the Broad Institute's mammalian sequencing effort [172].

The BRCA1 saturation-genome-editing function scores are those of Findlay et al. [71], as redistributed with the Arc Institute Evo 2 reference notebook together with the GRCh37.p13 chromosome 17 sequence.

**Human clinical panels.** The ClinVar consequence panel has no review-status filter; excluding its 353 unstarred variants (9.5%) moves the single-token 1B-to-7B capacity step from +0.033 to +0.036. The dbNSFP cut moves the reach panel's class ratio from 1:7.40 to 1:0.98. Membership is by presence in dbNSFP, but the consequence column and its missense subset carry ClinVar's own first term, so 16,549 rows (5.0%) show a term outside dbNSFP's scope, 99.2% of them benign; without them consequence class alone reads 0.819 rather than 0.837. ClinVar's gene annotation and allele frequencies, used for gene resampling and the rare-variant sensitivity, come from a later weekly ClinVar release, which lacks 81 of the reach panel's variants; those take no ClinVar gene. The reach panel admits a variant only when ClinVar's aggregate classification is exactly Pathogenic, Likely_pathogenic or Pathogenic/Likely_pathogenic, or Benign, Likely_benign or Benign/Likely_benign.

Table S24 gives each species' assembly, GERP track, gene models, snpEff database and source of negatives; the snpEff annotation covers every variant of all nine panels, positives at the coordinates of their OMIA table. Negatives miss chromosomes carrying positives in three species, reaching 12 of sheep's 27 chromosomes (1, 2 and 10 to 19; its pool ends within chromosome 2, as a draw taken in lexical order of chromosome names and cut short would), 19 of chicken's 34 and only pig's 18 autosomes. In all, 32 sheep, 13 chicken and 2 pig positives lie on chromosomes without negatives, and 4 more sheep positives lie on chromosome 2 beyond its last negative, at 45.78 Mb, where the sheep pool ends.

Which species carry which claims. At 8,192 bp the atlas makes three claims per species for Evo 2 against GERP: a locus-resampled margin excluding zero (Note S21), identification with unscored variants bounded, and a verdict holding in at least 95% of replicates (Note S22). Among the eight OMIA species, only cattle and dog support all three on negatives reaching every chromosome and over a hundred positive-bearing loci each; human does too, against ClinVar's benign records. Chicken supports all three, though 20 of its 28 positives are trait alleles. Sheep supports the first two, and still does without its 36 unreached positives (its verdict holding in 89.8% of replicates), but not without only the 32 on chromosomes carrying no negatives: its margin is then +0.065 (−0.053 to +0.154), its verdict holding in 67.2%. Goat's nine positives occupy two loci, so none of the three can be estimated. Pig, horse and cat support none: they are not identified at the point estimate, and are identified in only 7.8%, 5.6% and 25.4% of replicates. The grading is the same without the 51 positives the negatives do not reach, and without all 51 Evo 2's species-mean margin over GERP at 8,192 bp is +0.114 (+0.058 to +0.171). With loci resampled and no correction across the nine species, the snpEff impact ordering minus Evo 2 at 8,192 bp is resolved in three species: pig, +0.104 (+0.008 to +0.217), where annotation leads; dog, −0.035 (−0.066 to −0.007); and human, −0.014 (−0.020 to −0.008), where Evo 2 leads. Without pig, whose row confounds species with negative source, the species-mean difference falls from +0.015 to +0.004 (−0.023 to +0.031).

Annotation of the positives. Of the 594 positives OMIA curates as coding, 27 are non-coding by snpEff's first term, 16 of them in dog. Counting the 27 as coding, each with the impact rank of its curated class, raises the coding flag's mean over the eight OMIA species from 0.944 to 0.955 and the impact ordering's over the nine from 0.958 to 0.968. Negatives carry no curation and are left as snpEff calls them, so this bounds the effect of the positives' annotation errors rather than giving an error-free reading, and those errors work against the annotation baselines, not for them.

Frequency of the negatives. The atlas negatives, unmatched on frequency, could inflate discrimination without any signal of deleteriousness, since disease alleles are overwhelmingly rare and Evo 2's score correlates with allele frequency (Note S30). We tested this in cattle, the one species whose negatives span rare and common frequencies in a population panel available here (FarmGTEx), by restricting the negatives to progressively rarer strata. Against negatives carrying any frequency, discrimination is 0.915 at 1,001 bp and 0.984 at 8,192 bp; restricted to a minor-allele frequency ≤ 0.005, it is 0.911 and 0.984, well inside the resampling intervals. Discrimination is flat across frequency, so the atlas result is not a frequency effect in the species we can check. Only 723 of cattle's 1,880 negatives and 51 of its 188 positives carry a frequency in that panel, because curated disease alleles are largely absent from a population sample, which is why the control is run this way round.

##### *Panel completion at 8,192 bp and its 21 exclusions*

Of the 21 excluded variants, 18 lie within 4,096 bases of a contig end, where no full 8,192-bp window exists. Horse loses 15 of the 21, cat 3, and goat, cattle and dog one each. Charged under the must-answer rule, the 21 cost Evo 2 0.009 of AUROC in cat and in horse, and 0.003 by the nine-species mean. Cat's three, all positives, each carry a reference allele that disagrees with the assembly base, two matching it only on the opposite strand and one with reference and alternate swapped, so the reference check dropped them. Read on the gene's minus strand, the two are the catalogued alleles MYBPC3 p.Arg819Trp and DMD p.Gln1617* (G>A on the assembly's strand). The 1,001-bp readout, which takes the reference base from the assembly, scored G>T at both and no change at the swapped one (a delta of exactly 0), and their snpEff classes are those of the substitutions scored, synonymous and missense, not missense and stop gained. All 21 stay in the 1,001-bp analyses, where cat reads 0.846 with them and 0.848 without; that readout's reference check is recorded but not enforced, so its reference-allele failures are confined to these 21 variants, 0.19% of the panel. Every readout gain printed is computed on variants both readouts score; Table 1 and Figures 5b and 5c draw each readout on its own panel, and their species means agree with the matched ones to the printed digits.

#### Note S33. Specialist and supervised baselines, the embedding probe and a proximity control

The two supervised baselines are a floor, not competitors, both trained on the sequence of a 500-bp window, 250 bases upstream of the variant and 249 downstream. The 1D convolutional network passes the stacked one-hot reference and alternate windows (8 channels) through two Conv1d blocks of 64 channels with kernel width 9 to a global max-pool and a dropout-0.4 head. The second is gradient boosting (HistGradientBoostingClassifier) on 4-mer features, the normalised reference k-mer spectrum concatenated with the alternate-minus-reference difference. Both run over eight species, goat's 98 variants being too few to fit, trained leave-one-species-out on seven and by 5-fold stratified cross-validation within species.

The embedding probe reads the blocks.24.mlp.l3 hidden state of evo2_1b_base at the variant, for the reference and alternate windows, and fits a logistic regression (C = 0.5) under GroupKFold by gene. On a 10,000-variant balanced subset of the pig cis-eQTL panel (Note S25), the alternate-minus-reference difference reaches a gene-grouped AUROC of 0.504 [0.493, 0.516] and reference, alternate and difference together 0.519 [0.508, 0.531]; we report both. Both intervals resample variants rather than eGenes, so they may be narrower than the panel's dependence structure warrants. The concatenated set's interval excludes 0.5, so the probe points to a detectable effect of negligible size rather than a null.

##### *Specialist baselines, supervised results and controls*

ESM-2 [120] scored variants in six of the nine species but enough of both classes only in human, the one panel tabulated (Additional file 2: reports/tables.md, Table D4), where it reads above Evo 2 and GERP. The human ESM-2 panel is the 912 variants carrying an ESM-2 score, 652 of them positive (71.5%), because ESM-2 scores only protein-coding variants, which are enriched for positives. It is therefore not comparable to the atlas panels and we do not pool it. The headline AUROCs are not on one variant set: ESM-2's 0.887 and Evo 2's 0.838 are on all 912, GERP's 0.860 on the 881 it scores, where Evo 2 reads 0.837. The matched GERP margin, +0.023 [−0.011, +0.058], does not exclude zero, and charged for the 31 variants it cannot call, GERP reads 0.835; ESM-2 − Evo 2, +0.049 [+0.019, +0.079], is on all 912 variants, not the 752 annotated as missense. Supervised models trained on the labels themselves fare worse (Additional file 2: reports/tables.md, Table D4): zero-shot Evo 2 beats both in all eight species however they are trained. The network trained without its own species falls to 0.482 in human, below chance. The k-mer classifier performs comparably to the network in most species and beats it in four of eight, by +0.090 in human, itself evidence that neither extracts much beyond composition. In four of the eight species (pig, sheep, cat and dog) the leave-one-species-out network outperforms the one trained with that species' labels, which points to a specification or regularisation failure alongside data starvation. We therefore claim only that these labels are too few to train a competitive supervised predictor by this route, not that data volume is the sole limit. As a further panel-level control, 118 of 8,890 negatives (1.3%) fall within 1 kb of a same-chromosome positive, but the macro AUROC is unchanged (0.878 → 0.878) after excluding all 656 negatives within 50 kb, so proximity between classes is not inflating the result.

Run unchanged over 644,387 ProteinGym [6] single mutants scored with ESM-2's smaller esm2_t30_150M_UR50D checkpoint (197 held-out assays), the isotonic transfer of Note S28 moves median expected calibration error only from 0.141 to 0.139, beating no calibration on 116 of 197 assays. Through the Mondrian conformal pipeline of Note S28, a weaker Nucleotide Transformer backbone preserved overall coverage, pooled over the two classes (species means of 0.981 against Evo 2's 0.970 on the three held-out species), at a higher cost in abstention (a mean of 75% of held-out variants against 60%). One pair of backbones does not make abstention rate a proxy for model quality.

#### Note S34. The regeneration scope and reproducibility status of the deposited archives

Figures S1, S2 and S4 and the deposit's own report tables regenerate from Additional file 2 alone. Figures S5 and S7 regenerate from Additional file 2 with tables from Additional file 3 (Additional file 2: FIGURES.md). Figures 1 to 6 and S8 to S20 regenerate from Additional file 3's own scripts, tables and Additional files 4 to 6; six of its prep scripts need the licensed score values, and the inputs of Figures S19 and S20 are written from the withheld pCADD and chCADD values. The builders of Figures S3 and S6 also read the primary score and label tree, which is not deposited, so run from the deposit they stop with an error naming the missing input.

Of the 27 checking scripts in Additional file 3, seven need, for part of their values, scores the deposit withholds, licensed ones or pCADD and chCADD; the others run from the deposited tables and reproduce every published value they cover. Every reach-based number, including the sequence-blind baselines and all atlas values but those recorded only as summaries and the pCADD and chCADD values of Notes S43 and S44, regenerates from Additional files 2 to 6 alone. The deposit withholds pCADD and chCADD scores because neither host states a licence; provenance/build_atlas_species_cadd.py and build_matched_comparators.py rebuild them from the hosts, and recompute_atlas_comparators.py and recompute_atlas_matched.py check them against stored digests. The atlas values recorded only as summaries are the convolutional-baseline AUROCs of Figure S5b and the k-mer AUROCs of Table D4 (Additional file 2: reports/tables.md), Nucleotide Transformer v2's AUROCs and the chicken and cattle phyloP AUROCs of Table S5. The 83, 429 and 431 identified comparisons regenerate from two summary tables of Additional file 3 alone, tables/dbnsfp_breakdown_points.parquet and the coding-scope table tables/dbnsfp_coding_scope.parquet, though the covered AUROCs in them need the licensed scores to recompute. The breakdown points of Note S7 regenerate the same way, but not their gene-resampled lower limits; the comparisons' confidence sets and Holm and max-T counts need the licensed scores. Note S19's account of which panels dropped reference mismatches needs the 1,001-bp window sequences, which are not deposited; the deposited tables are consistent with it but do not test it. The pig eQTL allele frequencies of Note S25 also lie outside the deposit; Additional file 2 rebuilds them from PigGTEx genotypes. The per-variant scores of the eQTL window ablation (Note S25) are not deposited either, only its summary, nor are the 3,705 hosted Evo 2-40B scores of Note S19, the per-variant 1B and 7B scores of Note S29's 389-variant reverse-complement subset, the ESM-2 scores of Note S33's ProteinGym mutants or the EVEE shard's frozen scores.

#### Note S35. Where each reported value is recomputed

The arm table of Additional file 3's README maps each analysis arm to its reduced per-variant tables and the recompute script that checks its published values, and Figure S10 maps each variant panel and the published predictor benchmarks from source to the figures they feed.

#### Note S36. A general statement of the bound, the decision rule and the breakdown point

Setting. A metric averages a contribution between 0 and 1 over units (items or pairs of items). A predictor answers a unit when it responds on every item in it; ρ is the share of the metric's weight it answers and A its covered value, the average contribution there. A completion gives every declined item a response; nothing is assumed about which is true. For AUROC the units are the pathogenic–benign pairs, ρ the pair coverage and A the covered AUROC. The must-answer penalty, A minus the must-answer AUROC (which counts each unanswered pair as a tie), is (A − 0.5)(1 − ρ), positive whenever A exceeds one half and ρ < 1.

Proposition 1 (sharp bound). Under every completion the full-coverage value lies in [ρA, ρA + 1 − ρ]. Both ends are attained for item averages and, when a declined variant could score beyond every scored one, for AUROC. For other pair averages the interval holds but its ends need not be attained: with three subjects failing in turn, risk scores of 1 and 2 for the first and third, and the second declined, the concordance index attains only 0, 1/6 and 1/3 of an interval from 0 to 2/3. The F-score and the precision–recall area are not averages; their extremes are not functions of ρ and A.

Proposition 2 (orderings). Where both ends are attained, predictor 1 exceeds predictor 2 under every completion exactly when ρ₁A₁ > ρ₂A₂ + 1 − ρ₂; elsewhere the condition is sufficient. An identified ordering agrees with the covered values and holds under any penalty on wrong answers. If ρ₁ + ρ₂ ≤ 1 no covered values order the pair; if ρ₁ + ρ₂ > 1 some do. A bound with ρ ≤ ½ contains one half. On the dbNSFP panel, all 465 pairs of the 31 predictors with ρ < ½ are among the 536 comparisons that are never identifiable.

Proposition 3 (joint identification). What one predictor would have answered does not restrict another, unless one is a known function of the other, so every pair of attainable values is jointly attainable, and where the ends are attained disjoint bounds are necessary as well as sufficient. As a corollary, an identified ordering always has a positive head-to-head difference on the units both answer. Accordingly, none of the 83 identified comparisons reverses, and none of the 429 on the missense subset, when the side-by-side order of the covered AUROCs is set against that difference: a property of the rule, not a finding.

Proposition 4 (breakdown point). If a share λ of the unanswered weight resolves arbitrarily and the rest like the answered weight, the value lies in [A − (1 − ρ)λA, A + (1 − ρ)λ(1 − A)]. An ordering then holds exactly while λ < λ* = g/(g + o), with g the gap between the covered values and o the trailer's upper end minus the leader's lower end (the formula of Note S7); λ* exceeds one exactly when the ordering is identified.

Inference. The confidence sets of Note S3 assume that the ends of a bound are asymptotically normal across genes (Methods). A simulation in Additional file 3 (theory/THEORY.md, section 6) with 400 clusters of unequal size covered each end in at least 94.8% of 2,000 samples in each of three designs. None of its clusters carried most of the variance, as one gene does on the missense subset (Note S3), so it does not verify coverage on this panel.

What is textbook. The bound is Manski's [24], the contamination model Horowitz and Manski's [25], the confidence set of the Imbens–Manski type [50,51] and the breakdown point an established concept [54]; for untied scores the AUROC ends are those of the interval-based area under the curve [26]. Comparing two such bounds is textbook as well: with nothing linking two parameters their joint identified set is a rectangle, and the never-identifiable case extends the observation that a no-assumption bound on an average treatment effect contains zero [24]. Choe and colleagues identify an abstaining classifier's counterfactual score when abstention is stochastic and the evaluation data are independent of training [21]; here abstention is arbitrary, and an ordering can be identified although neither score is. With unknown labels, a joint analysis can be sharper than two marginal bounds [23]; with unknown predictions it cannot.

Proofs, and a script that checks each statement by enumeration on small benchmarks and recomputes the identification counts from the deposited breakdown points, are in Additional file 3 (theory/THEORY.md and scripts/verify_theory.py).

#### Note S37. What the conventions for unscored variants assume: least-deleterious fill, shared variants and the difficulty of declined variants

A fill (Methods) gives every unscored variant one score, so it orders every pair of predictors. On the missense subset the least-deleterious fill, with which the 97-predictor benchmark checks its ranking [5], orders all 1,176 comparisons and resolves 957, of which the bound identifies 427 (Table S29). Under the most-deleterious fill the same five predictors lead.

With both scores as dbNSFP distributes them, REVEL minus AlphaMissense is +0.0100 (+0.0068 to +0.0129) on the 189,301 variants both score and −0.0178 (−0.0294 to −0.0076) under the least-deleterious fill; the bound does not identify the pair (Note S7). By Note S36 an identified ordering holds under every fill and on shared variants, as the computation confirms.

On the missense subset, which lies inside every declared scope, a target is a predictor that declines at least 30 pathogenic and 30 benign variants (42 of the 49), and a reference one that scores more than 0.99 of the subset: CADD, DANN, ESM1b, the two BayesDel scores and six conservation tracks. A target's difference is the reference's AUROC on the variants it declines minus that on the variants it scores, averaged over references, with genes resampled. Declined variants are harder for 3 targets, easier for 7 and indistinguishable for 32; the median difference is +0.0098 and the intervals have a median width of 0.048. The direction depends on the reference: conservation tracks read declined variants as harder for 4 targets and easier for 10, the BayesDel scores as harder for 12 of 38 and easier for 1. A sequence-blind gene prior separates them worse for 19 targets, so gene composition is not controlled. The pathogenic share is a median 0.163 among declined against 0.293 among scored variants. On the 58,169 missense variants all 49 score, 28.7% of the subset, 41 predictors read higher than on their own scored variants, the pattern the sixth Critical Assessment of Genome Interpretation (CAGI6) reported [7]; a higher AUROC on that selected set does not show that a predictor's declined variants are harder. A target's difference measures other predictors, not the declining one, so it shows neither that declined variants are harder nor that they resemble scored ones, and it leaves the bound unchanged.

#### Note S38. Variants of uncertain significance: unscored, indeterminate and outside scope

We counted the three, outside scope, unscored and indeterminate, apart on ClinVar's variants of uncertain significance in the release that built the reach panel (Table S30; Methods). AlphaMissense's canonical-transcript release [41] leaves 57,494 missense variants of uncertain significance unscored. Of the 57,494, 39,742 lie in 292 genes, each with at least 20 missense variants of uncertain significance, in which the release scores none of them.

The possible missense substitutions of the 84 genes in which the American College of Medical Genetics and Genomics recommends reporting secondary findings (ACMG SF v3.3) [57] were enumerated from their MANE Select [109] coding sequences (Table S31). Table S31 shows that which AlphaMissense file a laboratory reads decides whether CALM1, CALM2 and WT1 receive any evidence from it.

Among the 204,191 labelled missense variants of the ClinVar panel, those REVEL leaves unscored are 6.7% pathogenic and AlphaMissense's 9.1%, against 28.6% overall; the likelihood ratios, with gene-resampled 95% intervals, are in Table S27 and the main text. ClinGen's second calibration [19] could not recommend EVE [20] because its predictions left about half of the benign calibration variants unscored, in genes with a skewed ratio of pathogenic to benign variants. The unscored level's likelihood ratio belongs to the file read: a missing value in dbNSFP's REVEL column gives 0.585 on the dbNSFP panel's 202,643 missense substitutions. It also mixes two mechanisms: 645 of AlphaMissense's 1,019 unscored pathogenic missense variants and 130 of REVEL's 252 lie in genes the release leaves wholly unscored (Table S32), and outside those genes the ratios are 0.117 and 0.101.

A separate export of the ClinGen Evidence Repository [46], with 13,717 classifications against the 13,293 of Note S8, holds 4,118 expert-panel classifications of uncertain significance, of which 1,843 carry no met PP3 or BP4 code. Placed by position, 1,220 of these are single-nucleotide variants that are missense on MANE Select, 325 are other single-nucleotide variants and 298 are not single-nucleotide substitutions. AlphaMissense has no score for 9 of the 1,220 and is indeterminate for 518; CADD scores all and is indeterminate for 363. REVEL, which the panels behind 994 of the 1,220 specify, was not read on variants of uncertain significance, and CADD only in the 84 genes and on these classifications. The expert panels are not a sample of clinical laboratories.

#### Note S39. Which genes, variant classes and disease categories go unscored

Unscored means that the source carries no value: dbNSFP 5.3.1a for the 32 predictors that declare a missense scope, and each tool's own release on the ClinVar panel. Annotation sources, the 16 disease categories and the tests are described in Methods.

Consequence class. Over the 2,277 genes of the dbNSFP panel with at least 10 pathogenic variants, the 32 predictors score a median 0.274 of a gene's pathogenic variants, and the missense share of those variants accounts for that value (r² = 0.967; Figure S17), as expected of predictors that score missense variants only. On the whole panel X-linked genes are scored more completely than autosomal ones (0.4946 against 0.3622), and 10 of 246 gene groups differ from other genes, each scored more completely (tubulins 0.953 and ion channels 0.610, against 0.372 over all genes).

Disease categories. No disease category departs from what its consequence classes predict by as much as 0.01. REVEL's own release leaves 0.569 to 0.729 of a category's pathogenic variants unscored. Across the 16 disease categories, the share of pathogenic variants REVEL leaves unscored follows the share expected from each category's consequence classes (r² = 0.994, the squared Pearson correlation over the 16 categories), and no category's gap exceeds 0.009; 0 of 16 gaps differ from zero after correction. For AlphaMissense's own release the largest gap is 0.006 and 1 of 16 differs (immune system diseases, 0.596 against 0.601 expected). Both tools score missense variants only, so the agreement is largely built in. Categories overlap, and 18,716 of the 170,680 pathogenic variants (11.0%) fall in none.

Inside missense scope. Over the 31 predictors with at least 20 unscored pathogenic missense variants, gene explains a median 0.845 of the variance in which such variants are declined (0.059 expected if declines were spread at random). dbNSFP's REVEL column carries no value in 137 of the 4,254 genes with at least 10 missense variants, and REVEL's own release scores more than 0.9 of the gene in 95 of the 137. The unscored share of a gene's pathogenic missense variants is not associated with gene constraint (LOEUF [110]; Spearman's rs = +0.015, 95% CI −0.034 to +0.065) or with X-linkage (X minus autosomes, −0.0126, −0.0288 to +0.0054), and 0 of 197 gene groups differ.

Variant classes. Table S32 gives the initiator-codon and mitochondrial-genome variants, and the whole genes, that widely used scores leave unscored. For 30 of the 32 predictors only dbNSFP's column was read, so their whole-gene gaps may belong to dbNSFP's copy rather than to the tool, as most of REVEL's do (above) and some of AlphaMissense's (Table S32).

#### Note S40. Named orderings and orderings printed in opposite directions

Printed predictor names were harmonised to tools and then to families, a family merging the tools of one predictor (PolyPhen-2 merging its HumDiv and HumVar scores, SIFT merging SIFT and SIFT4G), and thirteen families were fixed before any verdict was tabulated: AlphaMissense, REVEL, CADD, ESM1b, EVE, PrimateAI-3D, MetaRNN, BayesDel, VARITY, ClinPred, MutPred, SIFT and PolyPhen-2. The two sets of Note S24 hold 29 articles that print AUROCs and 16 assessable benchmarks, in which 287 stated orderings join two of the families: 177 are identified, 93 are not, 14 depend on the unreported class split and 3 are not assessable. Of the 93, 68 lie in Barbosa et al. [72] and Heo and Kim [73], and 32 come from the latter's sorted legend (61 of 253 without it). Their breakdown points (Note S7) have a median of 0.215, and 25 lie below 0.1. In 88 of the 93, one predictor leaves more than a tenth of the pathogenic–benign pairs unranked.

The best-known orderings are mostly decided. The rank column of CAGI6 [7], counted apart from the 287, places AlphaMissense above REVEL (0.879 against 0.869; 7 and 69 of 10,456 variants unscored), which is identified (λ* 2.865 to 4.017), as is AlphaMissense above ESM1b in all three assessable articles that print it. What stays undecided mostly involves a predictor that leaves many variants unscored: all 8 assessed instances for AlphaMissense and EVE, all 11 for CADD and EVE and 8 of 10 for REVEL and VARITY are not identified. Table S35 lists the ten orderings that the rule of Table 2 yields without its cap.

A pair of tools from different families counts as printed in opposite orders when two or more articles each print both on one panel and their orders differ, tied AUROCs set aside; the main text gives the counts, with names harmonised and exactly as printed (Analyses, The same accounting beyond one panel).

#### Note S41. A census of public genomic language model benchmarks

We applied the accounting to the per-variant scores that two benchmarks release; no model was rescored. The suite released with GPN-Star [30] holds 132 score files on 14 datasets in six species, 970 model pairs (Table S36), and its article evaluates each dataset on the variants every compared model scores. On that complete case, by the suite's own metric (average precision or a tail odds ratio) and resampling, 820 pairs are resolved at 95% confidence and 735 under Holm's procedure; with whole chromosomes resampled, 772 and 609. Of the 111 orderings the article's text states, 106 are resolved as stated and none is reversed. Its text also orders two classes of model: on OMIM, sequence-to-function models (Enformer, Borzoi and AlphaGenome) read below alignment-based genomic language models and conservation scores, and on complex traits above Evo 2-40B. Expanded to every released member of each class, its two statements about classes of model give 30 pairs, 26 resolved as stated and 1 the other way (AlphaGenome above PhyloP-P243 on OMIM, by +0.077 in AUPRC).

Fifty-five files leave variants unscored, 26 with reach that differs between the label classes; charged for them, no verdict on the pairs they touch reverses (Table S36). AlphaMissense leaves 2,381 of 50,164 ClinVar missense variants unscored (4.7%), so its whole-panel AUROC lies between 0.866 and 0.959, a range that contains GPN-Star's 0.925. The suite's dataset cards do not state the files' orientation, and the files do not share one. Read without labels, from each file's stored range and its rank correlation with the conservation and GPN-Star files beside it, 29 of the 132 run opposite to the GPN-Star file in their folder. With the orientations so read, the best file on every dataset is one the article's text places ahead of each file it compares it with, and never behind; on ClinVar it is AlphaMissense, which the text places above GPN-Star.

TraitGym [47] distributes five zero-shot readouts for each of ten self-supervised models and prints each model's best. Over all its models at the printed readouts, 107 and 101 of 136 pairs are resolved for Mendelian and complex traits, whole chromosomes resampled, so 29 and 35 printed orderings are unresolved. Under Holm's procedure 84 and 88 are resolved. TraitGym stores a missing conservation value as zero, at most 0.9% of a track; charging those values instead changes no AUROC verdict. Choosing each model's readout on the other chromosomes lowers average precision by a mean of 0.008 and 0.002 and reorders 2 pairs in each class. The choice of readout matters more: the order of 51 and 37 pairs depends on which of the five readouts each model is given, both orders being resolved in 30 and 31 (18 and 20 with a Bonferroni correction). Note S27 asks the same question of the 34 readouts it uses and counts 26 pairs, 7 with both orders resolved, which the census reproduces.

#### Note S42. The accounting on a language-model factuality benchmark

SimpleQA [48] grades each answer to a short factual question as correct (a share c), not attempted (d) or incorrect (w). The third table of its article gives the three shares of eight language models on the 4,326 questions, so the benchmark reports how often each model declines (Table S37). Covered accuracy is c/(c + w); the bound is that of Methods. The bounds identify 7 of the 28 pairwise comparisons, each involving a model that declines under a tenth of the questions; 7 can never be identified, the two widths summing to at least one, and 14 are undecided. Binomial confidence sets of the Imbens–Manski type [50,51] keep 6 of the 7 with 95% confidence, and 6 or 5 under Holm's procedure as models' outcomes are taken as independent across questions or as arbitrarily dependent. Under the contamination model of Note S7, 26 of the 28 comparisons stay ordered at λ = 0.05, and the 21 not identified break at a median λ of 0.295. Assuming declined questions would be answered no more accurately than attempted ones, 18 of the 28 are identified.

Three cautions keep the reading fair. First, an open question has no chance level of one half: the one-half value exceeds covered accuracy for all eight models, so the bound's lower end, a declined question counted wrong, is the analogue of the must-answer AUROC. Second, the benchmark's own three summary metrics (percentage correct, correct given attempted and their F-score) disagree on 7 of the 28 pairs, none of which the bounds identify. A penalty for wrong answers, which the benchmark's authors describe [48], reorders these eight models [74]. The bounds show which orderings need no assumption. Third, one cell of the source table cannot be as printed: GPT-4o's correct given attempted reads 38.0, below its share correct of 38.2. The three shares give 38.6, and the accounting uses only the shares.

#### Note S43. Stronger conservation comparators: 241-mammal phyloP, pCADD and chCADD

GERP's alignment depth differs across species (Note S21), so three stronger comparators were read at the atlas variants. The 241-mammal phyloP of the Zoonomia Cactus alignment [44,45] (phyloP-241) is one track for all nine species, read in place for human and elsewhere at the GRCh38 position each variant's base lifts to through the UCSC chain of its own assembly (Methods). It scores all 3,000 human variants and, pooled over the eight non-human species, 732 of 740 positives (98.9%) but 3,940 of 7,390 negatives (53.3%), or 97.4% and 55.2% by the nine-species mean. A base whose chain reaches no human position is unscored, so the scored variants are those the pairwise chains align; chicken is not among the 241 mammals, and its 55 values are those of the human-orthologous mammalian column. Charged for the variants the lift leaves unscored, phyloP-241 loses 0.170 by the species mean, against GERP's 0.102 (Table S38). That cost belongs to the lift, not the alignment, as a 241-way phyloP read on the cattle genome itself shows (Table S5).

On the 7,023 variants that GERP and both Evo 2 readouts also score, phyloP-241 leads GERP by +0.058 (+0.004 to +0.113). Evo 2's margins over phyloP-241, by species and by species mean, are in Table S38; with loci resampled the 8,192-bp margin excludes zero in chicken, cattle and human, and in no species in phyloP-241's favour. With every unscored variant bounded at both extremes, Evo 2 is identified ahead of phyloP-241 at 8,192 bp in cattle and human, and in goat at its point estimates, each verdict resting on phyloP-241 as the lift reaches it (Table S38); at 1,001 bp the comparison is identified only in human, in phyloP-241's favour.

Within consequence class, pooled as in Table S21, Evo 2's margin in the eight OMIA species is −0.114 (−0.190 to −0.040) at 1,001 bp and −0.036 (−0.103 to +0.019) at 8,192 bp, and in human −0.054 (−0.092 to −0.014) and −0.015 (−0.055 to +0.024): phyloP-241 leads at 1,001 bp, and at 8,192 bp neither is resolved ahead.

pCADD [66] and chCADD [67] score every pig and chicken variant on the atlas's own assemblies and so are charged nothing. Evo 2 minus pCADD is +0.006 (−0.098 to +0.098) at 1,001 bp and +0.021 (−0.083 to +0.128) at 8,192 bp. chCADD reads 0.981 (0.960 to 0.997), and Evo 2 minus chCADD is −0.120 (−0.216 to −0.044) and −0.028 (−0.081 to +0.014). Both read consequence annotation: pCADD minus the impact ordering is −0.125 (−0.224 to −0.044) in pig, chCADD minus it +0.058 (+0.011 to +0.118) in chicken. Neither host states a licence, so pCADD and chCADD scores are not deposited; Additional file 3's builder regenerates them.

UCSC has no database for the cat, sheep and goat assemblies, so they lift through the GenArk hub chains of the same assemblies, the other five species through database chains; without those three species the mean margins are +0.046 (−0.045 to +0.137) at 8,192 bp and −0.024 (−0.104 to +0.056) at 1,001 bp (k = 6, n = 6,349). Every value here is recomputed by scripts/recompute_atlas_comparators.py in Additional file 3.

#### Note S44. The consequence-matched arm

The atlas matches its negatives to its positives on trinucleotide context alone, so its positives are mostly coding and its non-human negatives mostly not (Figure 5f). The consequence-matched arm gives each positive population variants of its own consequence class and context from its species' own source (the same-class arm) and, for a protein-changing positive, synonymous ones (the synonymous arm; Methods). On the same-class arm the snpEff impact ordering reads near chance (Table S39), so the matching removes what annotation alone separates, as it must by construction; the near-chance reading checks the matching.

Evo 2 is not scored on this panel, and no Evo 2 score exists for any matched negative. The panel is deposited with its builder and seed in Additional file 3, and the baselines below are what a model scored on it has to beat.

Within consequence class both conservation scores drop on the same positives, phyloP-241 by +0.152 (+0.087 to +0.216) and GERP by +0.174 (+0.067 to +0.281) by the species mean (Table S39; Figure S20). Here and below a drop is the arm's AUROC subtracted from the atlas AUROC of the arm's own positives against the original negatives, so it is not the difference between Table S39's columns, whose atlas column takes every atlas positive. phyloP-241 reaches 89.8% of the matched negatives by the species mean, so its must-answer AUROC stays close to its covered one. Against synonymous negatives neither score's drop excludes zero: +0.009 (−0.003 to +0.020) and −0.003 (−0.053 to +0.047). What the atlas measured for conservation was therefore in large part, though not wholly, the protein-changing contrast, which the same-class arm removes and the synonymous arm keeps (Table S39). Gene is not matched, so gene-level constraint remains in these values. Stop gained reads well below missense on the arm for phyloP-241, and below it for GERP, whose intervals overlap (Table S39). Site conservation cannot see what makes a nonsense allele damaging (loss-of-function intolerance, nonsense-mediated decay, exon position), and the low stop-gained value is consistent with that.

GPN-Star falls within class in chicken and human (Table S39). chCADD falls in chicken from 0.981 to 0.549 (0.423 to 0.692), the pattern of a score that reads annotation, while pCADD in pig holds at 0.838 (0.723 to 0.926) against 0.842. Under the other pairing rules phyloP-241's mean runs from 0.719 to 0.741 (same-class arm, species mean: every pair of the arm, every served positive of the class, each positive against its own negatives, or exact-context matches only). Classes are the canonical transcript's; 111 of the 1,135 human negatives classed synonymous change the protein on another transcript. Pig's negatives are common genotype sites, as in the atlas. Human's ClinVar benign records may rest partly on BP4 evidence from conservation, phyloP among the calibrated tools [18], so human's conservation rows may be partly circular. Every value is recomputed by scripts/recompute_atlas_matched.py in Additional file 3.

### Supplementary Figures

#### Figure S1. Reliability of the transferred calibration, species by species

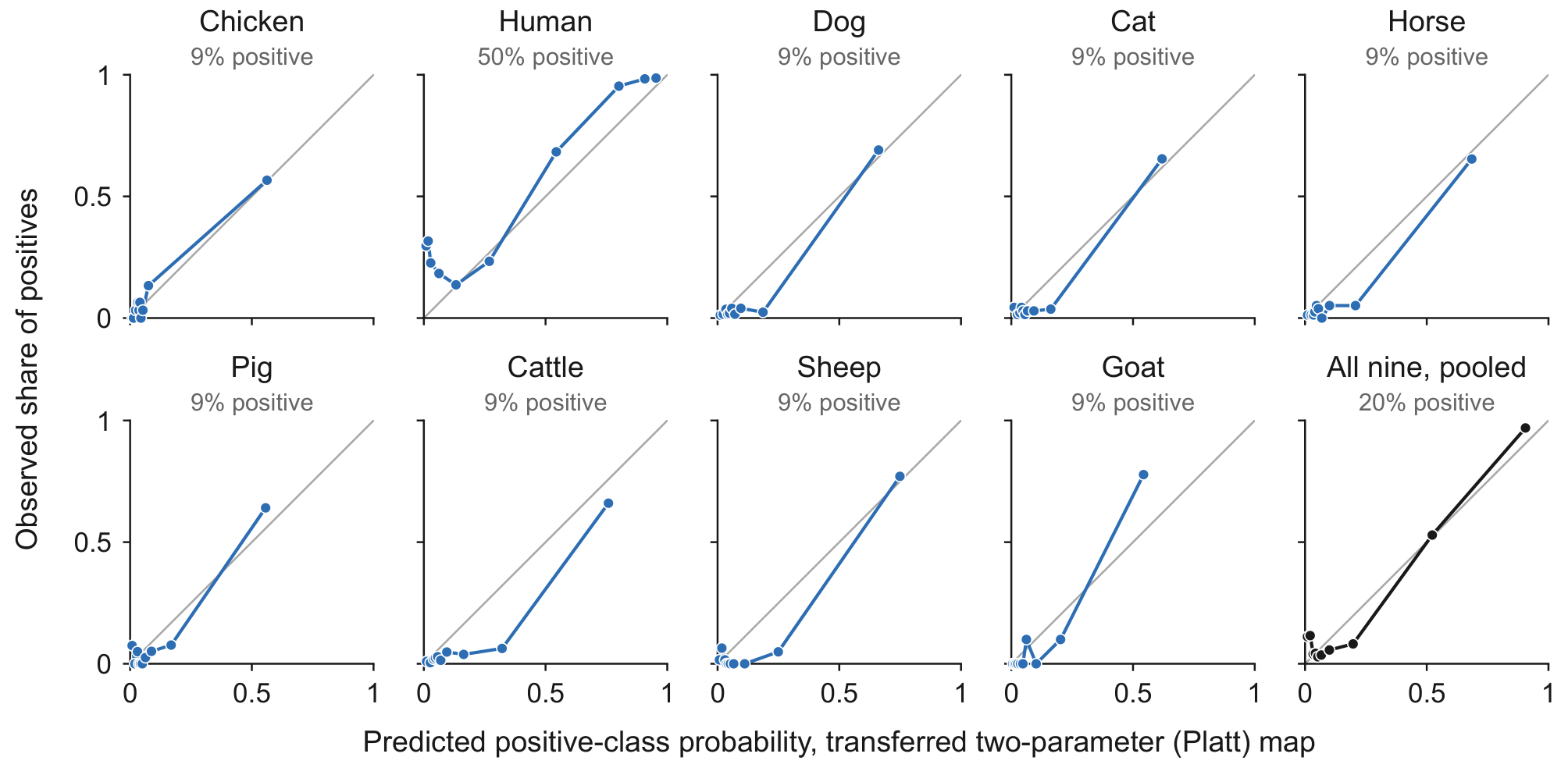

Evo 2-40B at the 1,001-bp single-token readout, 11,130 variants (per-species n in Table 1), under the two-parameter Platt map transferred leave-one-species-out. One panel per species and one pooled, each headed by its share of positives, with ten equal-count bins plotting the observed share of positives against mean predicted probability, against the diagonal of perfect calibration.

Alt text: Ten small reliability diagrams, nine species and one pooled, each plotting the observed share of positives against predicted probability in ten equal-count bins over a grey diagonal. The eight species built at one positive to ten negatives have most bins near zero and, chicken apart, mid-range bins below the diagonal; human, built 1:1, has its lowest bins far above it, at about 0.3.

#### Figure S2. Evo 2 reads near chance on one porcine cis-eQTL panel

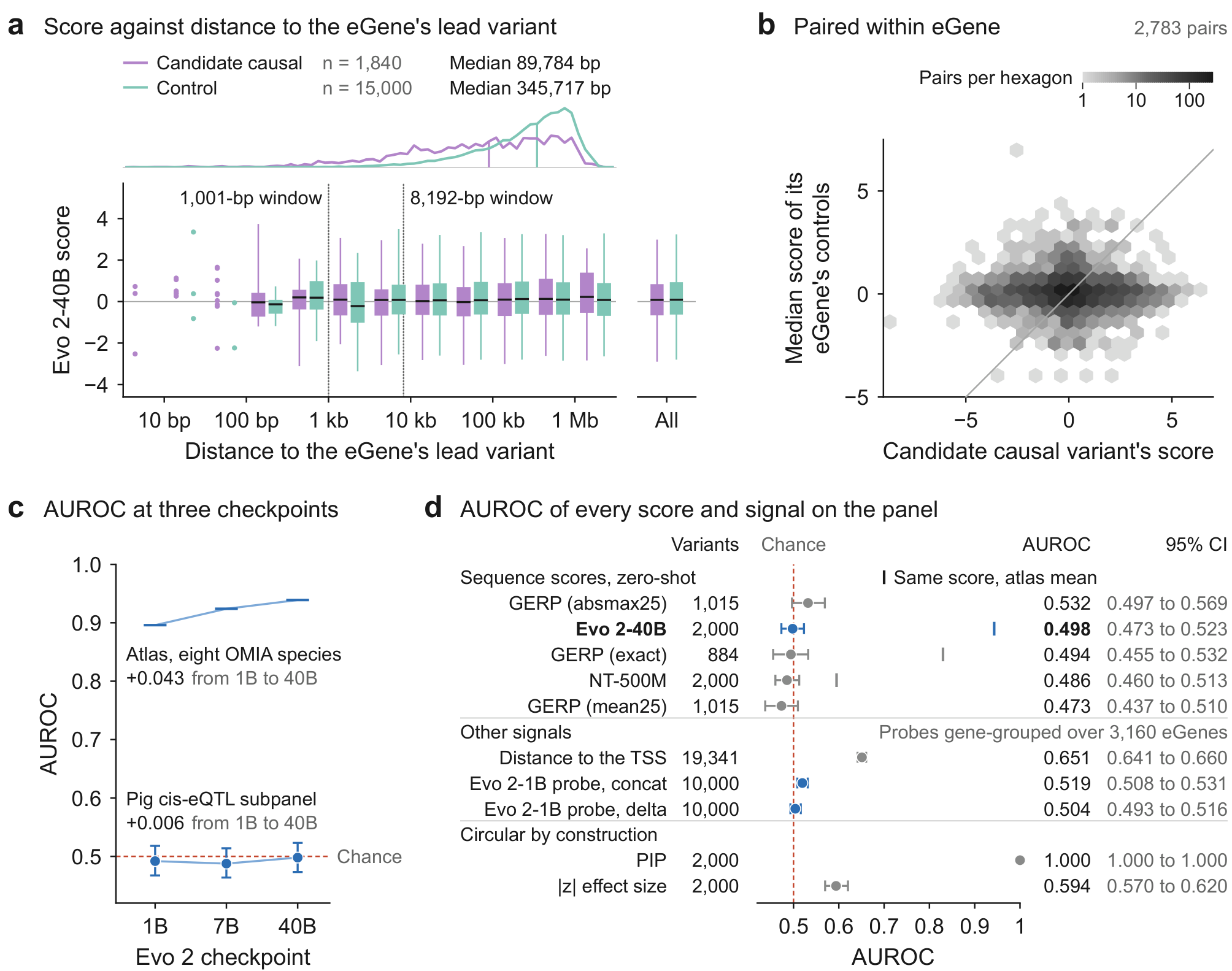

Evo 2-40B zero-shot on candidate causal pig cis-eQTL variants (PigGTEx [63,88]) against within-eGene controls, scores oriented so that higher is more deleterious; a and b at the 1,001-bp single token, c and d's Evo 2-40B row at the 8,192-bp window mean (2,000-variant subpanel). (a) Distance from the eGene's lead variant (leads not drawn): class densities with medians, and score per half-decade and overall (bar, median; box, interquartile range; whisker, 5th to 95th percentile; points, under ten variants); dotted lines, the lengths of the two readout windows. (b) Each candidate causal against its eGene's control median; 2,217 of the 5,000 have no control. (c) AUROC (area under the receiver operating characteristic curve) at the three released checkpoints with 95% intervals (the 40B point is d's Evo 2-40B row; here and in d, percentile bootstraps over variants, not eGenes); dashes, the species mean over the eight OMIA species (at 40B, Figure 5b's value; human's 1B scores excluded as a scoring fault, Note S10). (d) AUROC, 95% interval and variants scored for every scorer and signal; ticks, the same score's mean over all nine atlas species (Evo 2-40B's and GERP's are Figure 5b's species means, 0.943 and 0.830; NT-500M's rests on its atlas AUROCs, recorded as summaries, Note S34), whereas c's dashes are means over the eight OMIA species. GERP is read at the variant's base (exact) or as the mean or largest absolute value within 25 bp either side; NT-500M, Nucleotide Transformer v2; probes, logistic regressions, cross-validated with eGenes as groups, on Evo 2-1B embeddings of the alternate-minus-reference difference (delta) or of reference, alternate and difference together (concat), Note S33; |z|, the largest absolute PigGTEx z value across tissues; PIP, posterior inclusion probability; TSS, transcription start site.

Alt text: Evo 2 and candidate causal pig cis-eQTL variants. a: score boxes of the two classes coincide; candidate causals sit nearer the lead. b: a band straddling the diagonal. c: from 1B to 40B the eQTL AUROC moves +0.006, the species mean +0.043. d: all five scores span chance, Evo 2-40B at 0.498; distance to the transcription start site reads 0.651.

#### Figure S3. Score is associated with allele frequency without curated labels, and its limits

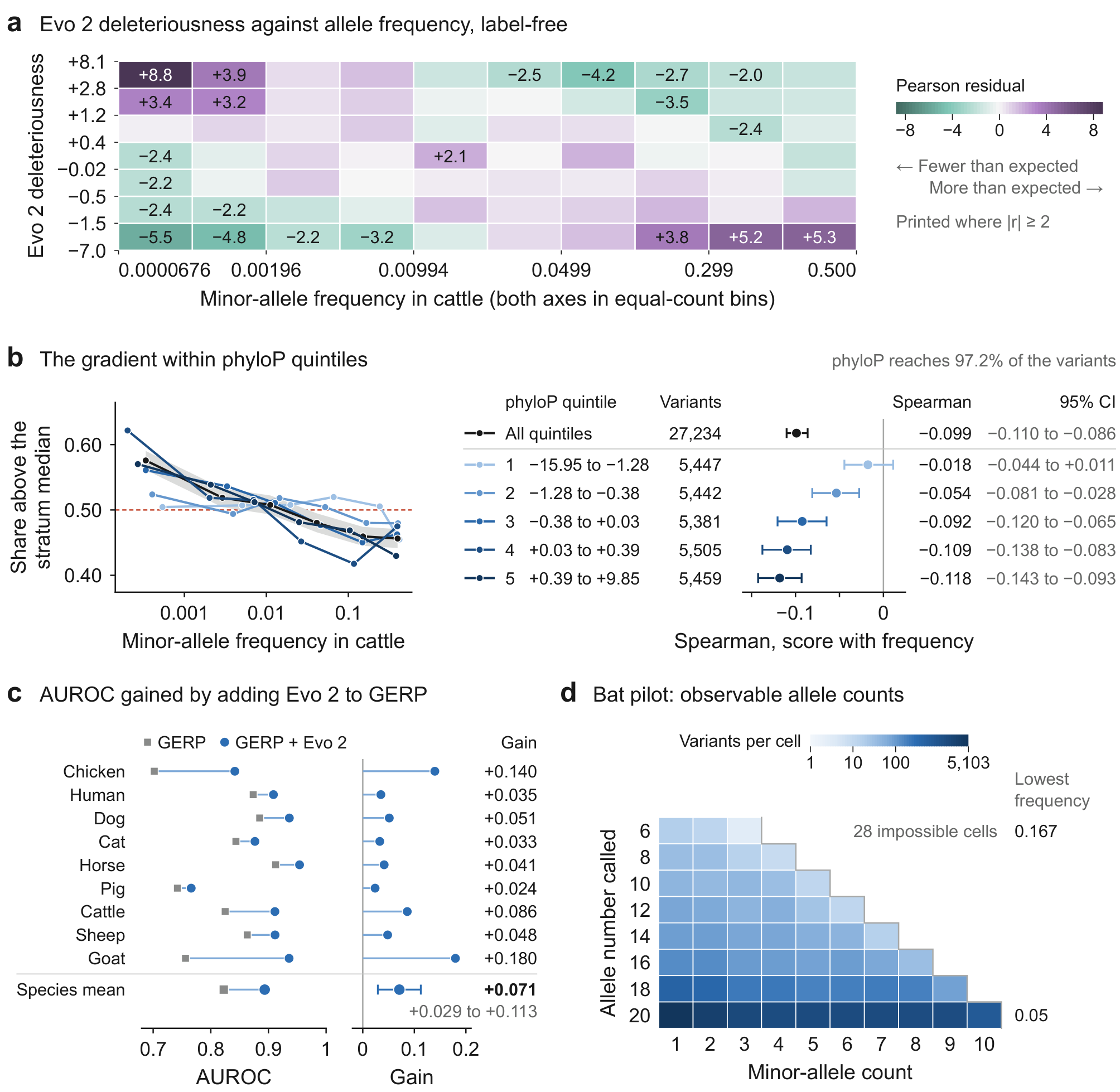

All panels: 1,001-bp single token. (a) Pearson standardised residuals of Evo 2 deleteriousness against minor-allele frequency (MAF), equal-count bins, 28,006 cattle variants. (b) Left, share above each phyloP quintile's median against MAF, darker with conservation; black, all quintiles with their Wilson 95% band; dashed, the null of 0.5. Right, for each line, its variants and the Spearman correlation of deleteriousness with MAF with a 95% interval (percentile bootstrap over variants), on the 27,234 variants phyloP reaches. (c) Per species, AUROC of per-base GERP alone and with Evo 2, and the gain (the one figure panel using disease labels, from the nine labelled panels), each from five-fold cross-validated logistic models fitted within species on the variants that GERP and the single-token readout both score (9,532 in all; Note S10), not on Figure 5b's variants (the GERP column is Figure 5b's quantity on a different variant set), with the species mean and t interval on eight degrees of freedom. (d) Wild-bat pilot, ten *Myotis lucifugus* libraries, 23,888 variants, by allele number called and minor-allele count; cells beyond the step line are impossible, and a row's lowest frequency is one over its allele number.

Alt text: Evo 2 and allele frequency. a: residual grid, 28,006 cattle variants. b: by rising phyloP quintile, the Spearman correlation of deleteriousness with frequency runs from −0.018 in the lowest, the only quintile spanning zero, to −0.118. c: Evo 2's lift over GERP is positive in all nine species, mean +0.071. d: 23,888 bat variants; 28 cells are impossible.

#### Figure S4. Flexible calibration shows no robust transfer gain; confidence has limited utility

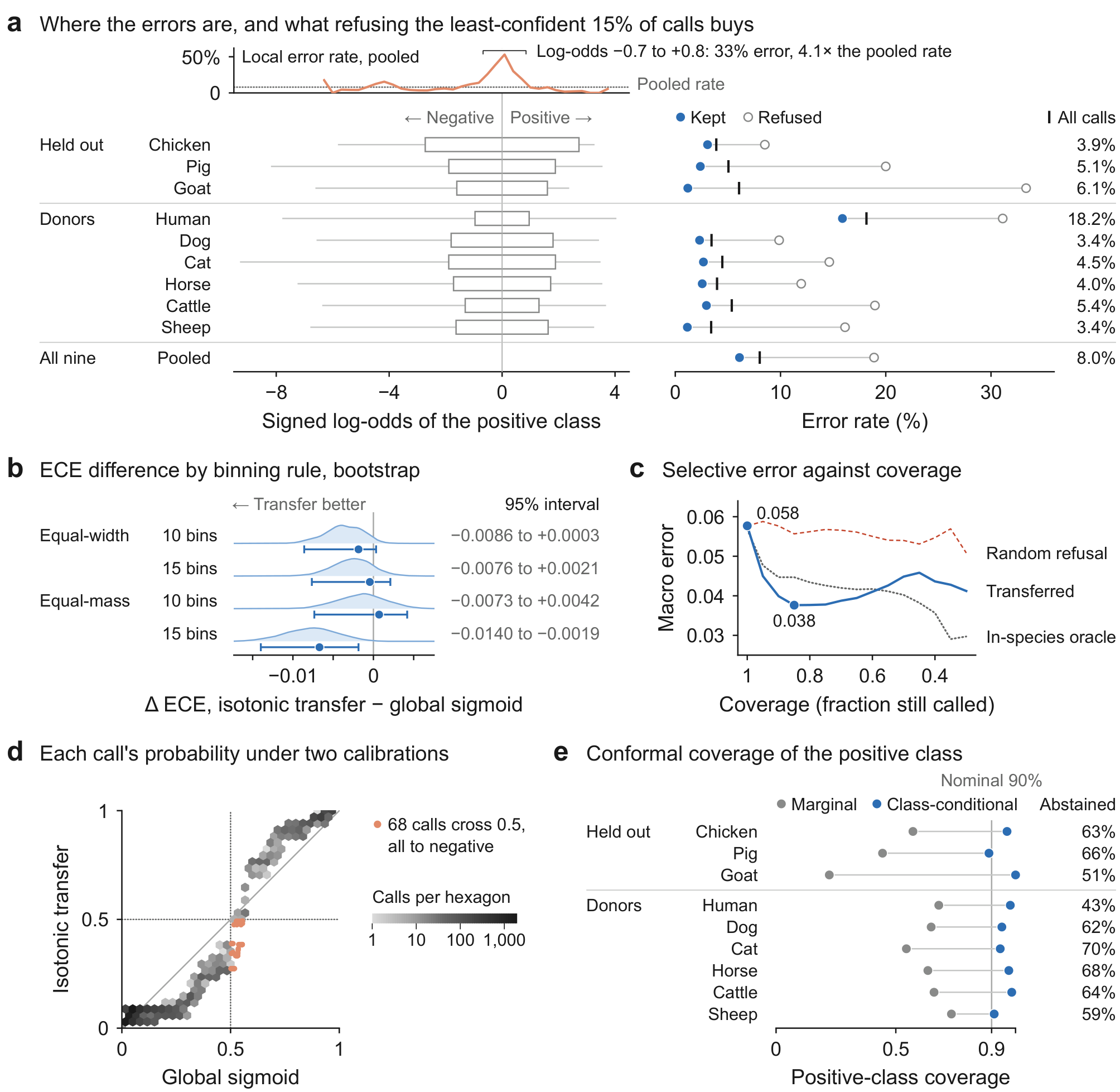

Evo 2-40B, 11,130 variants, 1,001-bp single token; held-out species never donate a calibration map. (a) Local error by species and Platt log-odds bin (lift over random refusal per species in Table S17): top, the pooled rate per bin (ten or more calls), bracketed above twice the pooled rate; rows, each species' range of calls, its least-confident 15% (open box) and its error rate among kept, refused and all calls. (b) Bootstrap macro expected calibration error (ECE) difference, isotonic transfer minus global sigmoid, by binning rule: distribution, 95% interval and full-sample estimate. (c) Macro selective error against coverage, with random refusal and an in-species oracle. (d) Transferred against sigmoid probability, per call, drawn from the per-variant isotonic reconstruction deposited with Additional file 2 (macro ECE 0.050), while Table S16 prints its own fit (0.053). (e) Positive-class coverage of marginal and Mondrian (class-conditional) conformal predictors against the nominal 90%, with the latter's abstention. Orange marks the pooled error rate per bin in a (dotted, the overall pooled rate) and, in d, the 68 calls that cross the decision boundary.

Alt text: Calibration transfer and selective prediction. a: local error by species and bin, peaking at the decision boundary. b: four ECE-difference distributions peak below zero. c: the transferred arm stays below random refusal, lowest at 0.85 coverage. d: 68 calls cross the decision boundary. e: marginal coverage falls below 90%; class-conditional coverage nears it at 43% to 70% abstention.

#### Figure S5. Evo 2 where GERP is silent, and a supervised baseline

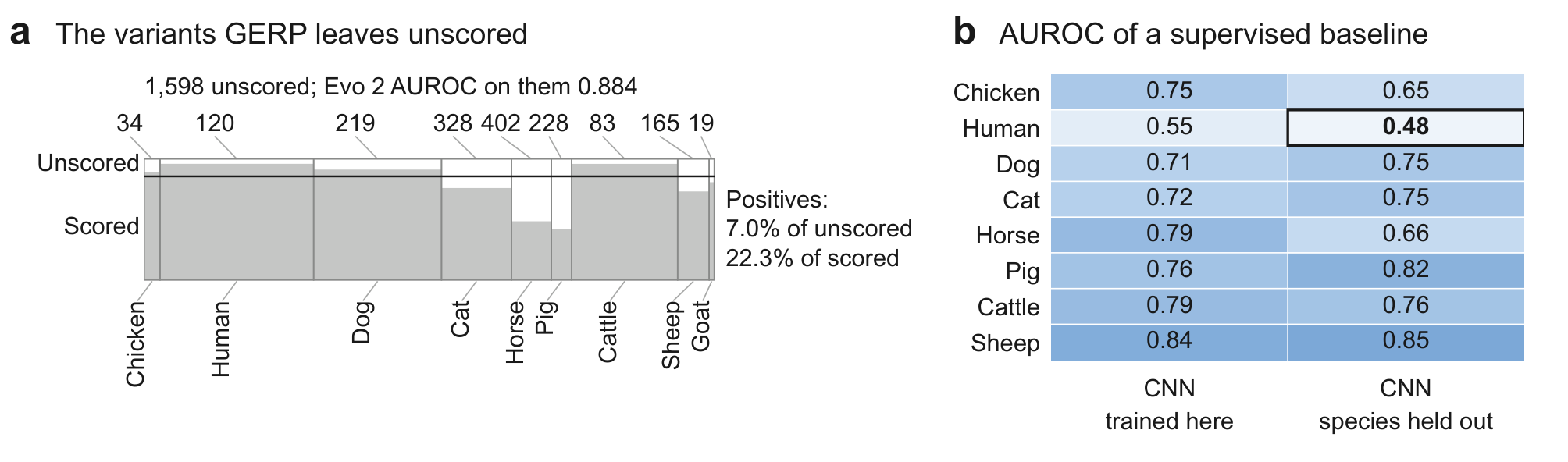

AUROCs at the 1,001-bp single token. (a) Extends Figure 5e, which draws GERP's reach per species (values in Table S1), with each species' count of unscored variants and Evo 2's AUROC on them: one area-true block, a column per species as wide as its share of the 11,130 variants, open where GERP gives no score, its unscored count above; the rule marks the pooled share, and at right is the share of positives among the unscored and among the scored. Evo 2 separates all 1,598 at AUROC 0.884 pooled over species, 90% of its ranked pairs cross-species. (b) AUROC of a convolutional neural network (CNN) trained within species and with it held out (goat, too few variants to fit, has no row); one value, below chance, is outlined. Evo 2's zero-shot AUROC in each of these species, higher than both, is in Figure 5c (1,001-bp readout).

Alt text: Two panels. a: the variants GERP leaves unscored in nine species as one block, by species; Evo 2 separates them at AUROC 0.884. b: a network trained within species reads 0.55 in human to 0.84 in sheep; held out, it falls to 0.48 in human.

#### Figure S6. BRCA1 saturation genome editing scored zero-shot, in sequence and in structure

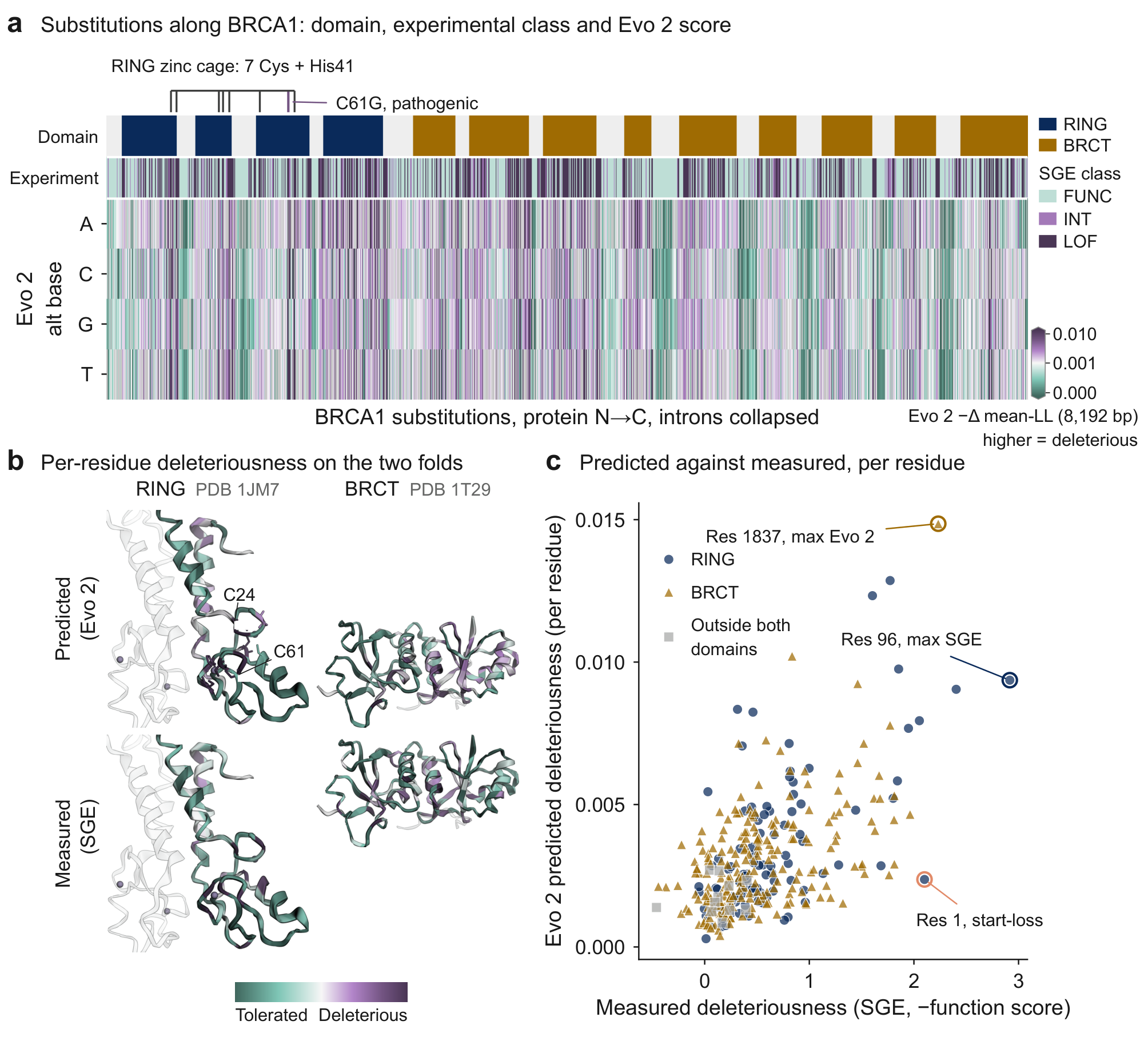

Saturation genome editing (SGE) of BRCA1 [71], 3,893 substitutions, scored zero-shot by Evo 2-40B as the negated 8,192-bp mean log-likelihood change, without labels. (a) Substitutions N→C with RING ligands, domains, each site's experimental class, the worst of its substitutions (FUNC, functional; INT, intermediate; LOF, loss of function), and the quilt, on a two-slope linear colour scale centred on the functional median (+0.00097), not zero; grey, no substitution. By their worst substitution all eight ligands are LOF, and seven are in Evo 2's top decile of residues. (b) Per-residue mean deleteriousness on the RING (Protein Data Bank 1JM7; grey ribbon, BARD1, and spheres, zinc ions, not scored) and BRCA1 C-terminal (BRCT, 1T29) folds, predicted above measured, each map coloured over its own 5th to 95th percentiles, so colours compare within a map, not across maps. (c) Per-residue predicted against measured, by domain: Spearman correlation +0.56 in RING (n = 101) and +0.50 in BRCT (n = 214); the correlation over all residues, and the AUROC over variants, LOF (823) against FUNC and INT (3,070), are in Note S18.

Alt text: Three panels on BRCA1 saturation genome editing. a: a quilt of substitutions ordered along the protein under domain and experimental-class strips. b: the RING and BRCT structures coloured by predicted and by measured deleteriousness. c: per-residue predicted against measured deleteriousness by domain, with the maximum-Evo 2 and maximum-SGE residues and the start-codon miss labelled.

#### Figure S7. Evo 2 against GERP at the single-token readout: within-species percentiles

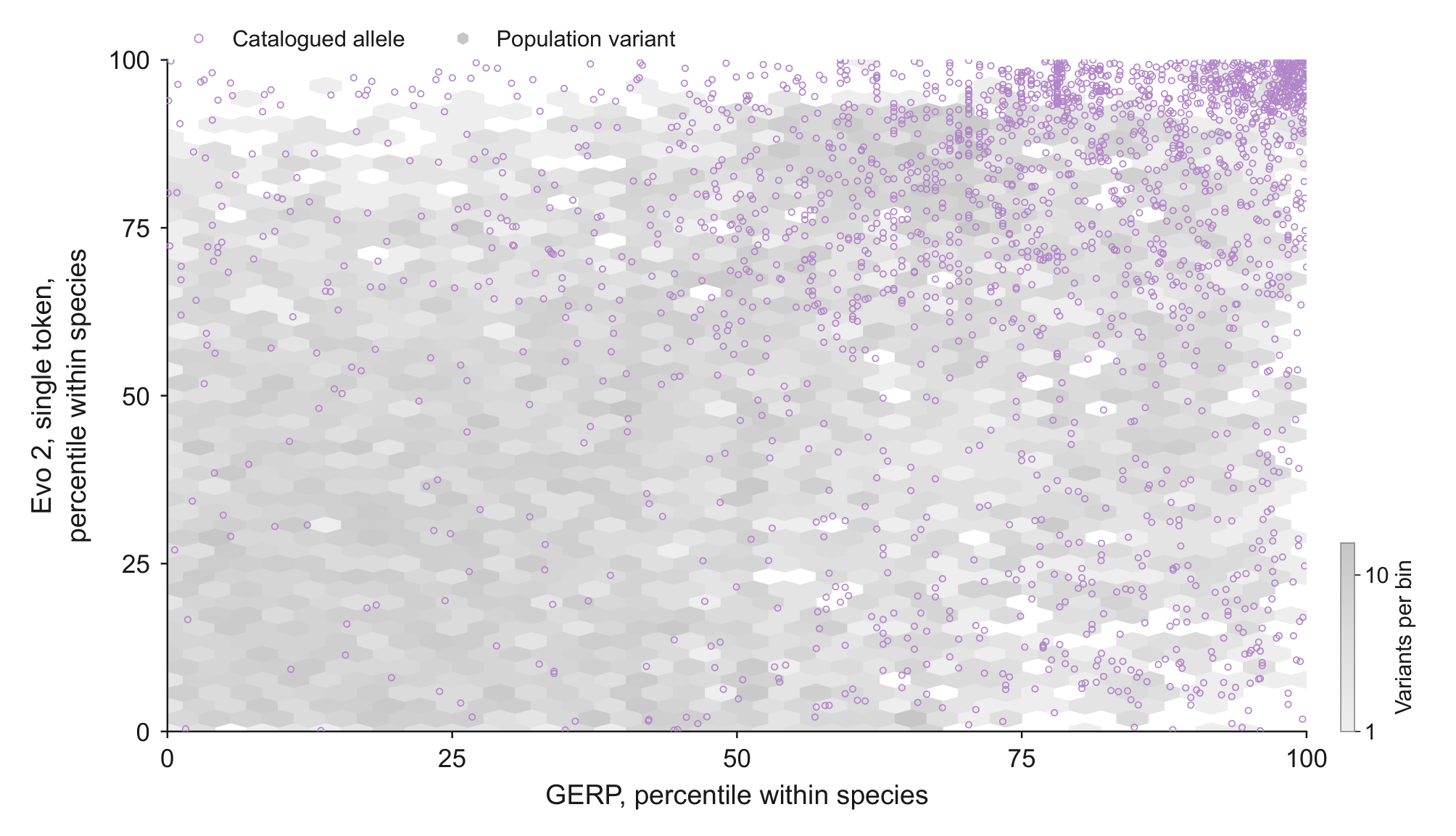

Within-species percentiles of Evo 2-40B at the 1,001-bp single token against GERP [42,43] on the 9,529 variants both readouts and GERP score; catalogued alleles (circles) over population variants (hexagons). Figure 5d gives the per-species margins.

Alt text: Hexbin of population variants with catalogued alleles overlaid: Evo 2's within-species percentile against GERP's; catalogued alleles crowd the upper right.

#### Figure S8. Atlas missense variants scored by AlphaMissense at the aligned human residue

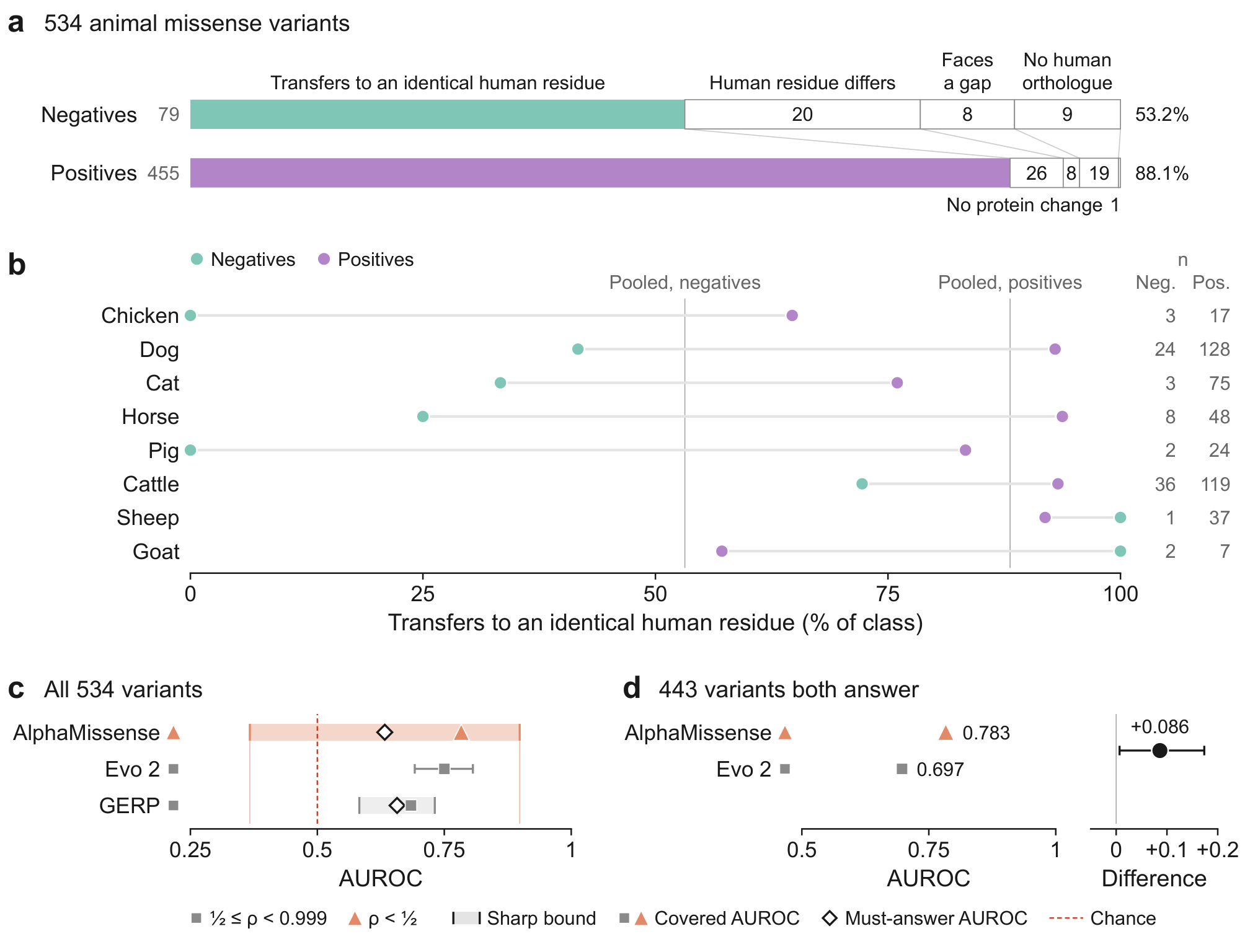

The atlas's 534 non-human missense variants (455 positives, 79 negatives), each as the human substitution at the aligned orthologous residue. (a) Per class, the share transferring to an identical human residue (filled) and the variants declined, by reason (open boxes, with counts). (b) The same share per species, class sizes at right; vertical lines, the pooled shares of a. (c) AUROC on the 534 variants, the covered values on the variants each score answers and the must-answer value on all 534: AlphaMissense's covered (triangle) and must-answer (diamond) AUROC and sharp bound (band, its ends carried down as thin lines) at pair coverage ρ = 0.469; below, Evo 2 with its 95% interval and GERP with its bound; dashed line, chance. (d) The 443 variants both score: AlphaMissense against Evo 2, with their paired difference and 95% interval (locus bootstrap, 2,000 replicates; likewise Evo 2's in c).

Alt text: 534 animal missense variants carried to human orthologues. a: 401 of 455 positives transfer against 42 of 79 negatives. b: by species. c: at pair coverage 0.469, AlphaMissense's AUROC bound, 0.367 to 0.899, contains Evo 2's 0.750 and GERP's 0.684. d: on the 443 both score, AlphaMissense leads, 0.783 against 0.697, by 0.086 (95% interval 0.007 to 0.174).

#### Figure S9. The pair space of each of the 49 dbNSFP predictors

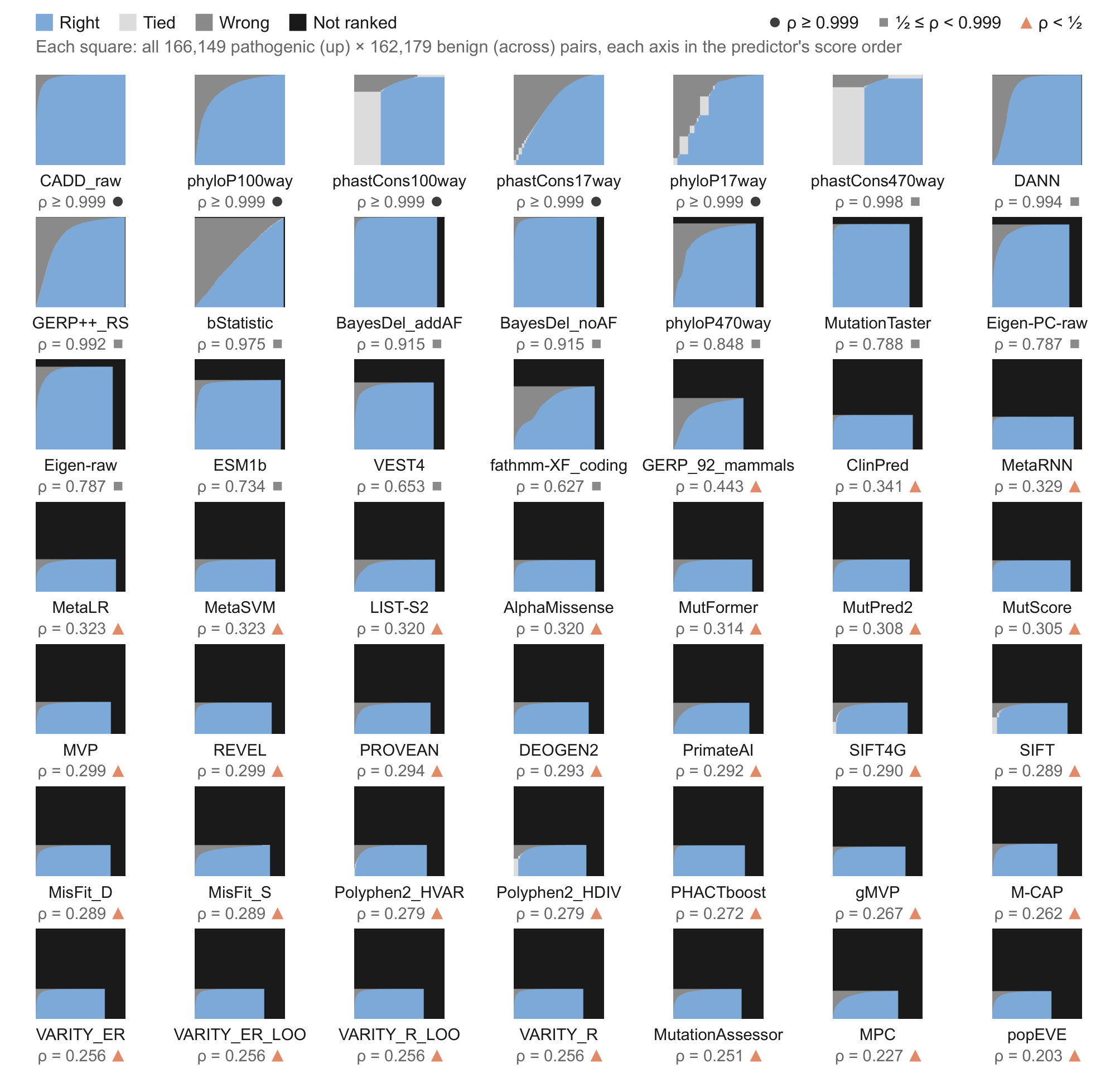

Figure 2b gives each predictor's pair coverage ρ, and Figure 2c the bounds these squares encode; new here, the split of every pair into right, tied, wrong and not ranked. Each square holds every pathogenic–benign pair of the dbNSFP panel of Figure S15, 166,149 pathogenic by 162,179 benign substitutions, one per predictor in that figure's order: pathogenic up, benign across, in score order, declined last. Pairs a predictor ranks fill the lower-left rectangle, of area its pair coverage ρ (printed, with its regime glyph); inside it, the covered receiver operating characteristic (ROC) curve parts pairs ranked right (blue) from wrong (mid grey), ties light grey. Black pairs are not ranked: right plus half the ties gives the bound's lower end, the black share its width.

Alt text: A seven-by-seven plate of squares, one per dbNSFP predictor in Figure S15's order, each blue at its lower left where the predictor ranks pairs and black elsewhere. The first 18 are blue over most of their area; the other 31 are black but for a bottom band, ranking only 20% to 44% of the pairs.

#### Figure S10. The study map: each panel of the study from its source to the figures it feeds

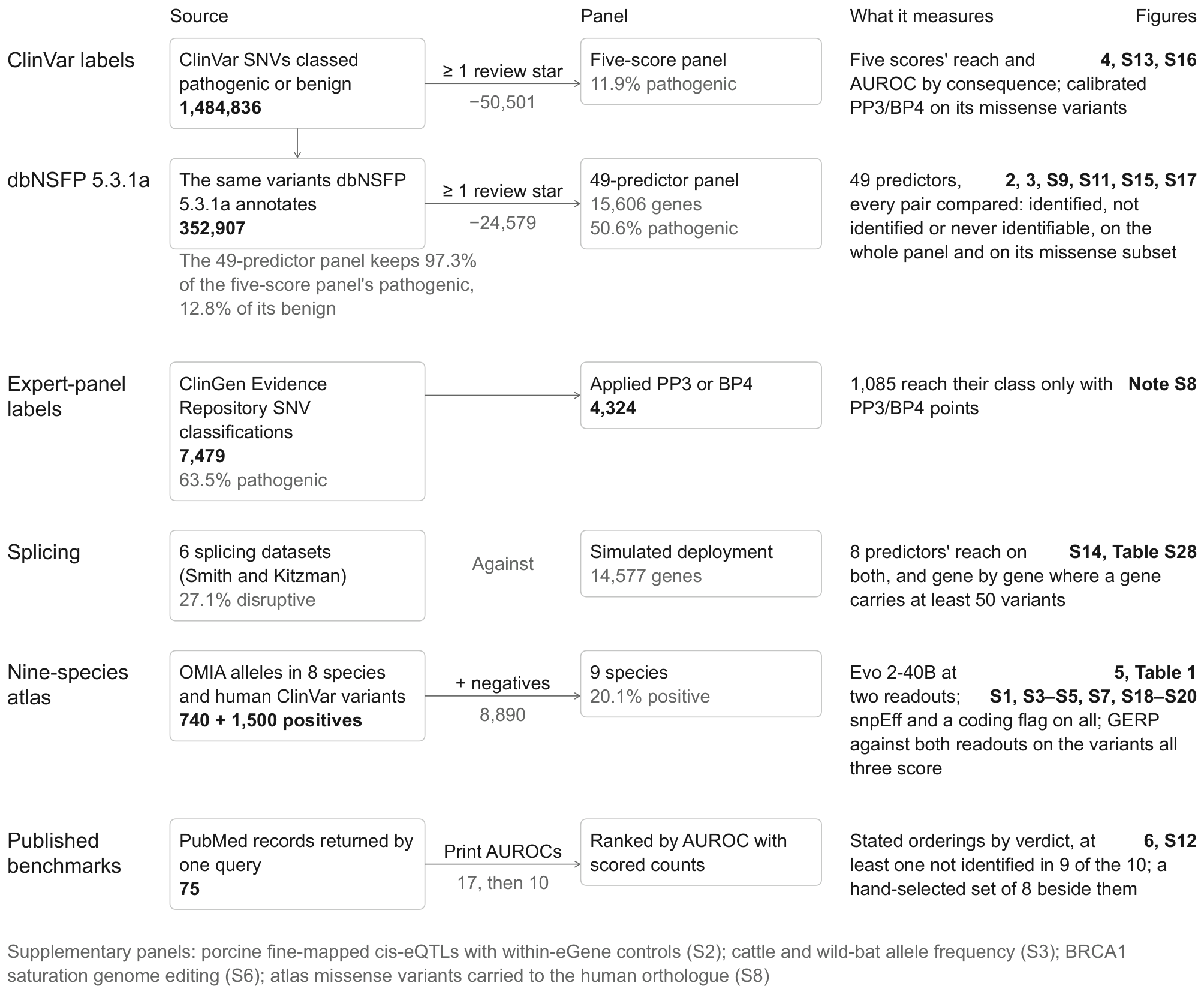

Extends Figure 1d, which gives each panel's size and share of positives, with the source of each variant panel and of the published predictor benchmarks, the cut that makes it and the figures it feeds; the genomic language model benchmarks and SimpleQA of Figures 6d to 6g have no row. Each row is one panel: its source, the cut that makes the analysed panel, the panel, what it measures (AUROC, area under the receiver operating characteristic curve) and the figures, tables or Notes that report it. Panel sizes are on Figure 1d (where a cut is shown, the size is the source less the cut; the atlas adds its negatives to its positives). The published-benchmarks row is the systematic search of Table S33 and Figure 6b; the hand-selected set is in Figure S12a, b (eQTL, expression quantitative trait locus; GERP, Genomic Evolutionary Rate Profiling; OMIA, Online Mendelian Inheritance in Animals; PP3 and BP4, computational pathogenic and benign evidence; SNV, single-nucleotide variant).

Alt text: A six-row flow chart from each source to its panel and the figures that report it: ClinVar's five-score panel; the 49-predictor dbNSFP panel; ClinGen classifications with PP3 or BP4 applied; splicing variants against a simulated deployment; the nine-species atlas; and ten benchmarks from 75 PubMed records.

#### Figure S11. Every rank each of the 49 dbNSFP predictors can hold

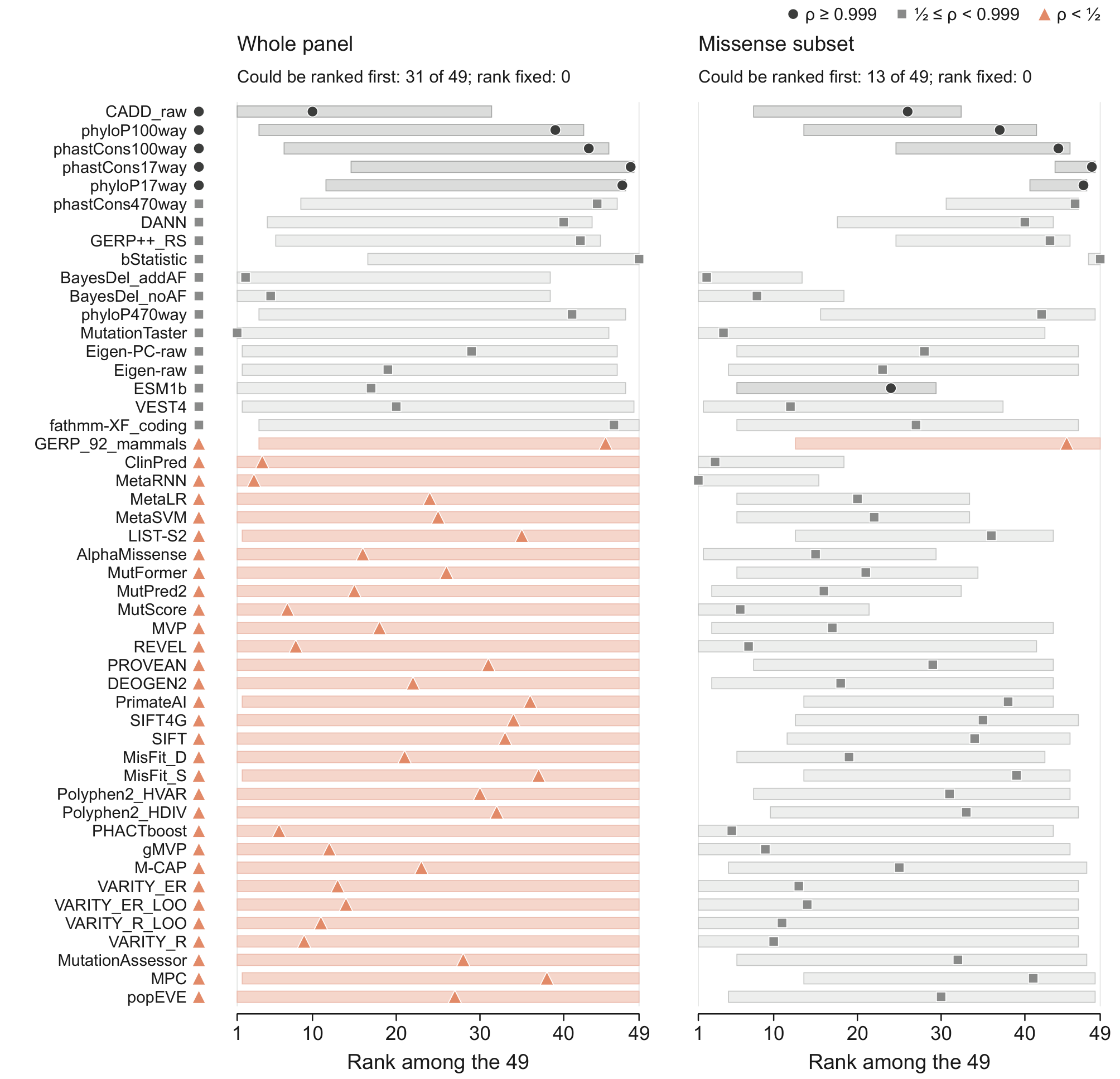

For each predictor of Figure S15, in its order, the band spans every rank among the 49 (1, the highest, at left) that the sharp bounds allow (Figure 2c); the marker is the rank its covered AUROC would give, which the bound does not identify. Left, the whole panel; right, its 202,643 missense substitutions. Marker shape and band tint give the predictor's pair-coverage regime on that panel (key); the glyph beside each name, its regime on the whole panel. Of the 49 predictors, 31 could be ranked first on the whole panel and 13 on the missense subset; no rank is fixed on either, and the median rank set spans 49 and 34 ranks.

Alt text: Two columns of bands, one row per dbNSFP predictor in Figure S15's order, each spanning the ranks the predictor could hold, a marker at its covered-AUROC rank. On the whole panel 31 predictors could rank first and none is fixed; on the missense subset 13 could rank first and the median band spans 34 ranks.

#### Figure S12. Published benchmarks one by one: a hand-selected set, and the ten under other readings (Note S24)

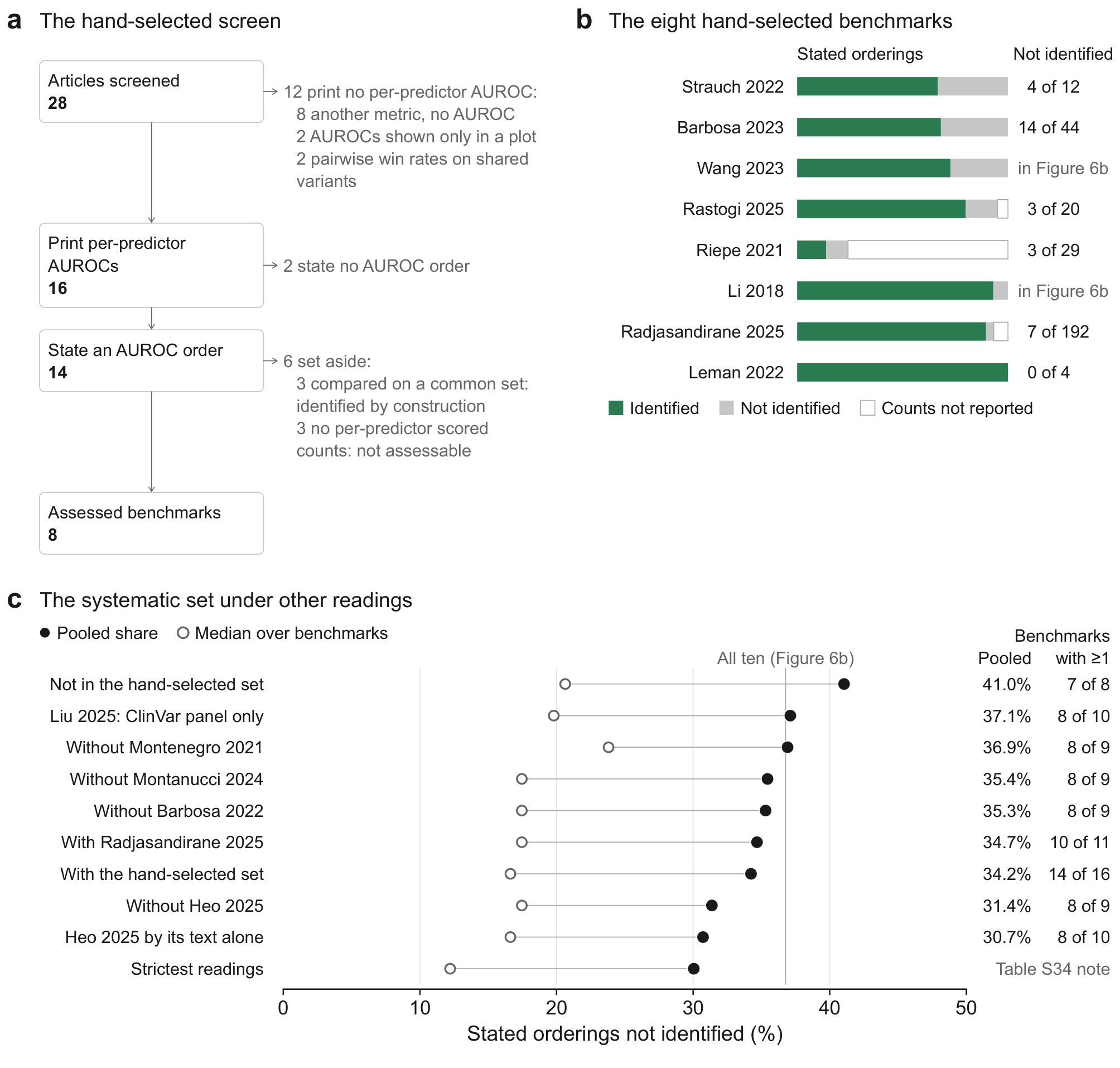

Extends Figure 6b, which gives the systematic set's stated orderings by verdict: new here, the hand-selected set screened and assessed benchmark by benchmark, and the systematic set under other readings. (a) The screen of the hand-selected set: the 28 articles screened, the 16 that print a per-predictor AUROC, the 14 that state an AUROC order and the eight assessed, with what each step sets aside. (b) Each hand-selected benchmark's stated AUROC orderings in Figure 6b's colours (each bar, its benchmark's orderings as 100%): identified, not identified, or counts not reported, with the number not identified at right (Li 2018 and Wang 2023 are also in the systematic set, Figure 6b). (c) The systematic set (Table S33) under other readings: the share of stated orderings the bound does not identify, pooled (filled) and as the median over benchmarks (open), without the two benchmarks it shares with the hand-selected set (Li 2018 and Wang 2023), with single benchmarks left out (among them the two largest, Barbosa et al. [72] and Heo and Kim [73], left out together in Figure 6b) or added (one or all of the hand-selected set of a and b), Liu et al. [135] read on their ClinVar panel only, the sorted legend of Heo and Kim read as stating only what its text states, and the strictest readings (Table S34 note); vertical line, all ten (Figure 6b); right, the pooled share and the benchmarks holding at least one ordering not identified.

Alt text: Three panels. a, the hand-selected screen from 28 articles to 8 assessed benchmarks. b, one bar per assessed benchmark. c, the systematic set's share not identified under ten alternative readings, from 30.7% to 41.0% where printed, against the all-ten line.

#### Figure S13. Calibrated clinical evidence along the distributions of four scores on ClinVar

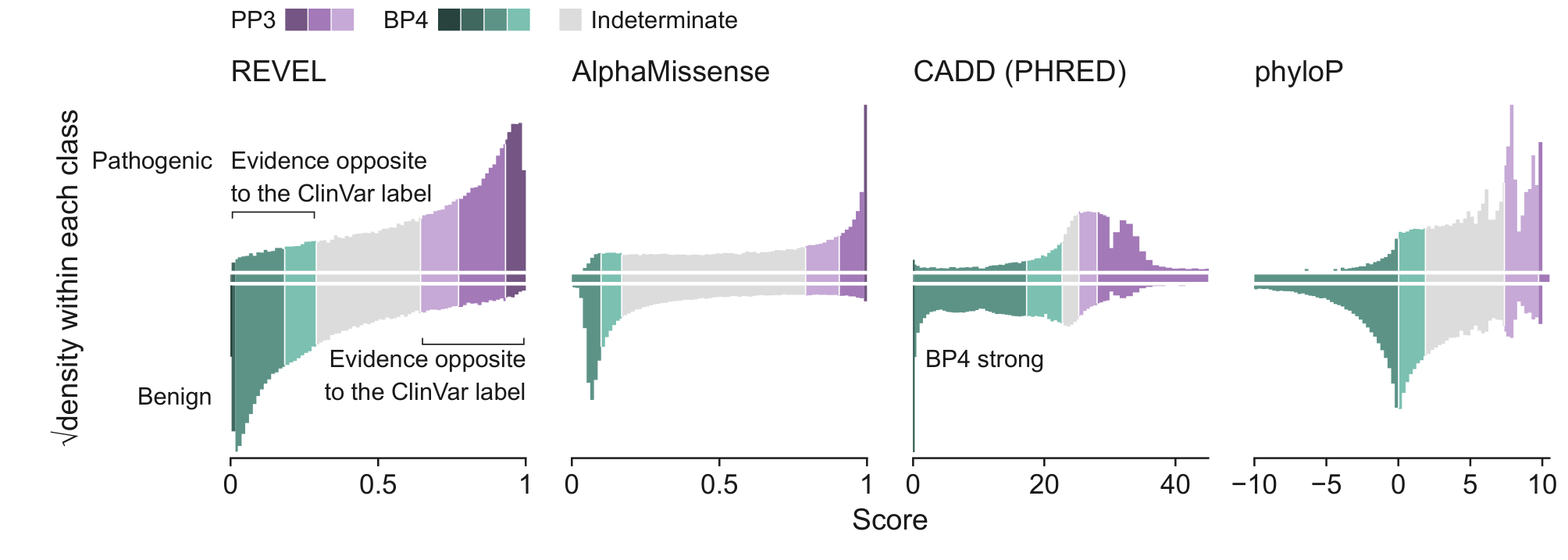

Figure 4f gives the share of each class at each strength of evidence (values in Table S22); here, where on each score those shares lie. Pathogenic (PP3) and benign (BP4) computational evidence at the strengths calibrated for the Rare Exome Variant Ensemble Learner (REVEL), Combined Annotation Dependent Depletion (CADD) and phyloP [18] and for AlphaMissense [19], on the ClinVar panel's missense variants at one review star or better, the only variants the calibration covers. Each score's distribution among the pathogenic variants (above the axis) and the benign ones (below; class sizes on Figure 4f), on a square-root density scale, coloured by the evidence its calibrated intervals give: PP3 purple, BP4 teal, darker when stronger (supporting, moderate, strong and, for BP4, very strong), indeterminate grey; the strip along each axis marks the intervals. CADD is PHRED-scaled; brackets on REVEL mark the tails given evidence opposite to the ClinVar label, and one bin too narrow to show its colour is named. Variants a score leaves unscored (Table S22) are not drawn.

Alt text: Four score histograms, pathogenic above and benign below, coloured by calibrated PP3 and BP4 strength, for REVEL, AlphaMissense, CADD and phyloP on ClinVar missense variants. Most pathogenic variants fall in PP3 bands, most benign ones in BP4 bands, but the tails give BP4 to 3.4% to 7.7% of pathogenic and PP3 to 2.7% to 8.3% of benign variants.

#### Figure S14. Reach of eight splice predictors on a benchmark and on a deployment background

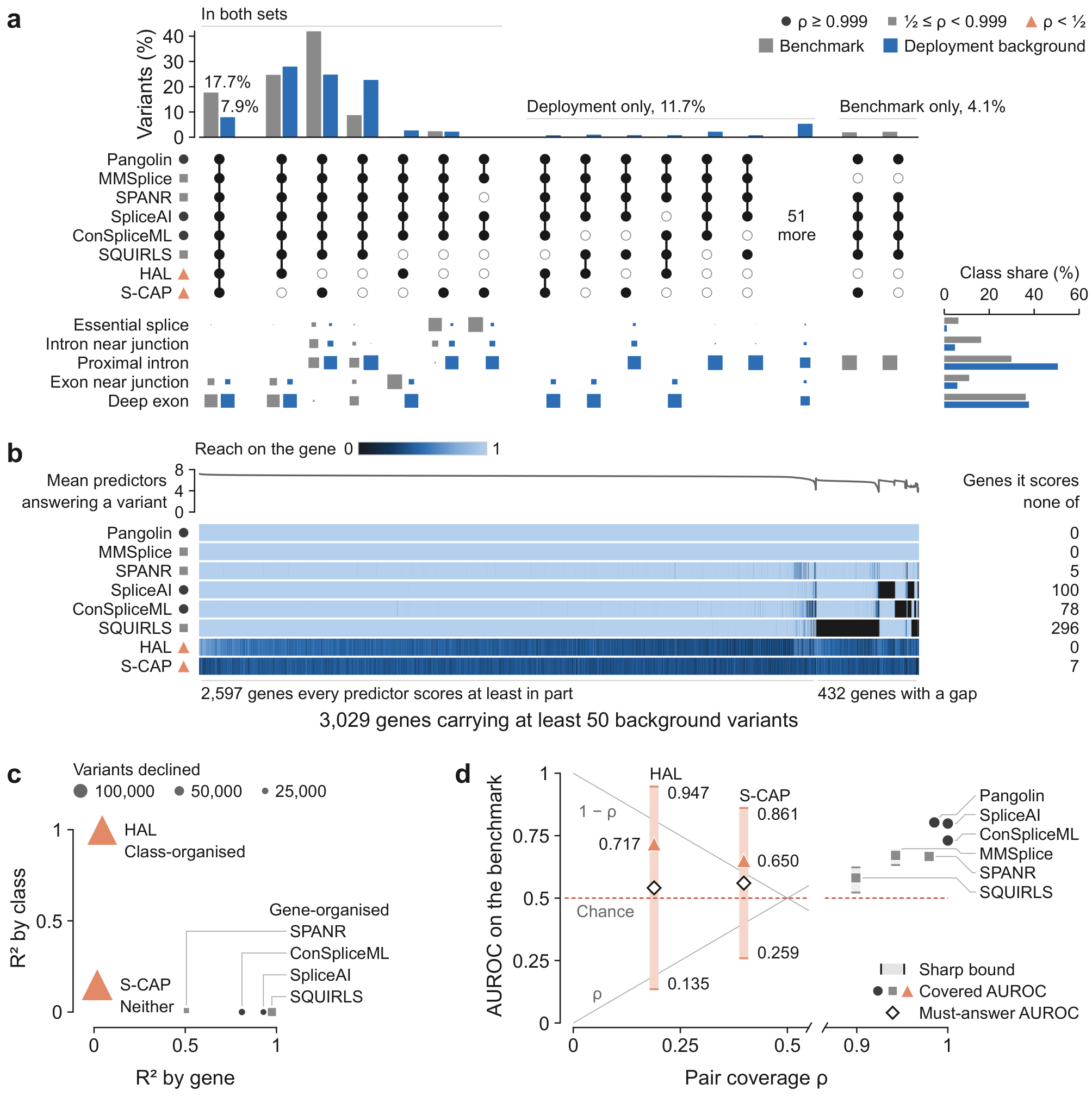

Pangolin [95], SpliceAI, ConSpliceML, Splicing-based Analysis of Variants (SPANR), Modular Modeling of Splicing (MMSplice), Super Quick Information-content Random-forest Learning of Splice variants (SQUIRLS), Splicing Clinically Applicable Pathogenicity prediction (S-CAP) and the Hexamer Additive Linear (HAL) model, on Smith and Kitzman's six-dataset benchmark [58], 3,912 variants, and their 500,000-variant simulated background; the glyph beside each name gives its pair-coverage regime on the benchmark. (a) Answer signatures, the set of predictors scoring a variant (filled dot, scores; open, declines), as shares of benchmark (grey) and background (blue); squares, each signature's variants by class (area, the share of the signature's variants in that class), and at right each class's share of each set. (b) Each predictor's reach on the 3,029 background genes with at least 50 variants, from light (every variant scored) to black (none). (c) Variance in each declining predictor's answering explained by gene and by variant class; marker area, variants declined. (d) Sharp bound (band), covered (marker) and must-answer (diamond) AUROC against pair coverage ρ; dashed, chance; thin lines, the highest lower end (AUROC = ρ) and the lowest upper end (1 − ρ) a bound can have.

Alt text: Eight splice predictors on a 3,912-variant benchmark and a 500,000-variant background. All eight answer 17.7% of the benchmark but 7.9% of the background. Of 3,029 well-covered genes, SQUIRLS scores none in 296, SpliceAI in 100, ConSpliceML in 78, S-CAP in 7, SPANR in 5. HAL's and S-CAP's bounds span chance; the other six lie between 0.522 and 0.803.

#### Figure S15. Where 49 published predictors decline on the 328,328-variant dbNSFP panel

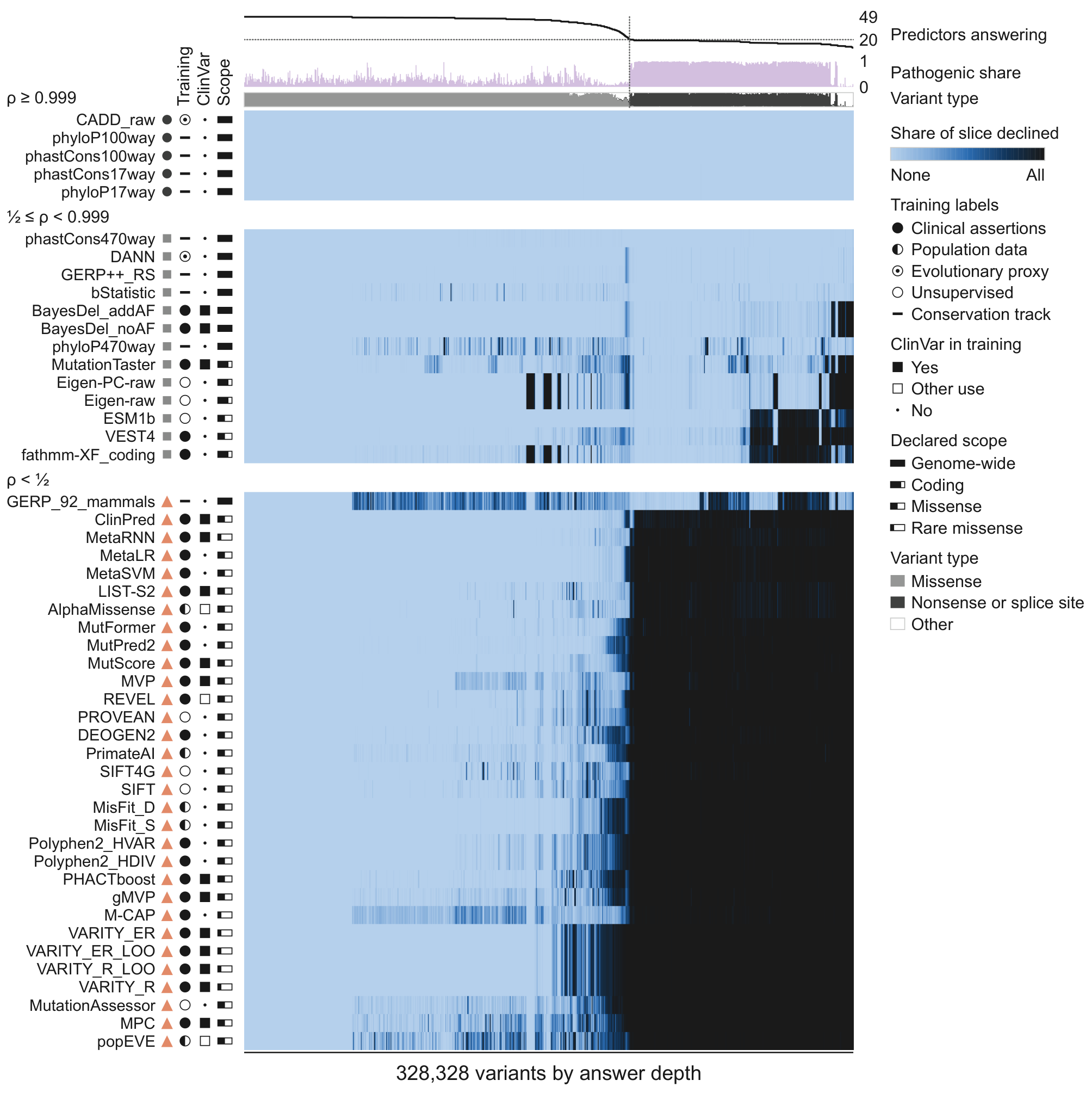

Figure 2b gives each predictor's share of pathogenic and of benign variants scored; here, where in the panel its declines fall. Each row is a predictor, in Figure 2's order, grouped by pair coverage ρ (heading and glyph), with its training labels, use of ClinVar in training and declared scope. The raster orders the variants into 700 slices by how many of the 49 answer them (answer depth) and shades the share of each slice the predictor declines (light, none; dark, all). Above, per slice: the number of predictors answering (dotted, 20 of 49), the pathogenic share and the variant types.

Alt text: Raster of 49 predictors across 328,328 ClinVar variants ordered by how many predictors answer each. Conservation tracks other than GERP_92_mammals answer almost everything; 30 missense predictors decline nearly all variants on the right, where at most 20 of 49 answer and most pathogenic variants, nonsense or splice-site, lie.

#### Figure S16. Five widely used scores on 1,434,335 ClinVar single-nucleotide variants, class by class

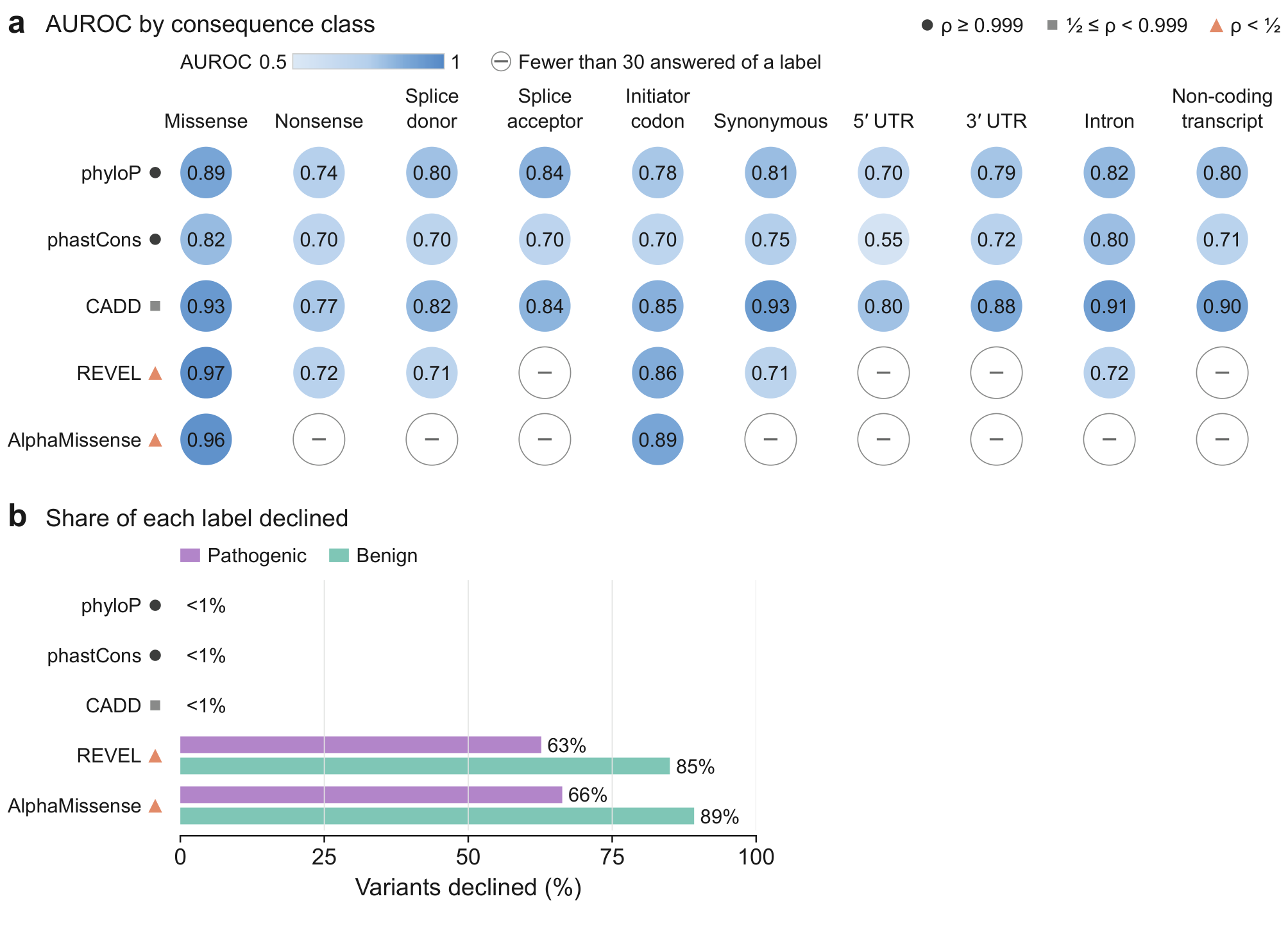

Extends Figure 4b and d, which give the share scored per class and a within-class AUROC summary: new here, each score's AUROC in every class and the share of each label it declines. (a) For each consequence class (UTR, untranslated region; sizes, pathogenic shares and the share scored per class in Figure 4b), each score's area under the receiver operating characteristic curve (AUROC) on the variants of that class it answers, shaded and printed; dash, fewer than 30 answered variants of a label. Glyphs, pair coverage ρ (REVEL 0.056, AlphaMissense 0.036). (b) Shares of pathogenic and benign variants each score declines; phyloP, phastCons and CADD decline under 1% of either. Figure 4c gives the bounds.

Alt text: Five scores by ten consequence classes. phyloP, phastCons and CADD have an AUROC in every class, 0.55 to 0.93; REVEL has one in six classes and AlphaMissense in two, 0.97 and 0.96 on missense. REVEL and AlphaMissense decline 63% and 66% of pathogenic and 85% and 89% of benign variants; the other three under 1%.

#### Figure S17. Which genes, variant classes and disease categories missense-scope predictors leave unscored

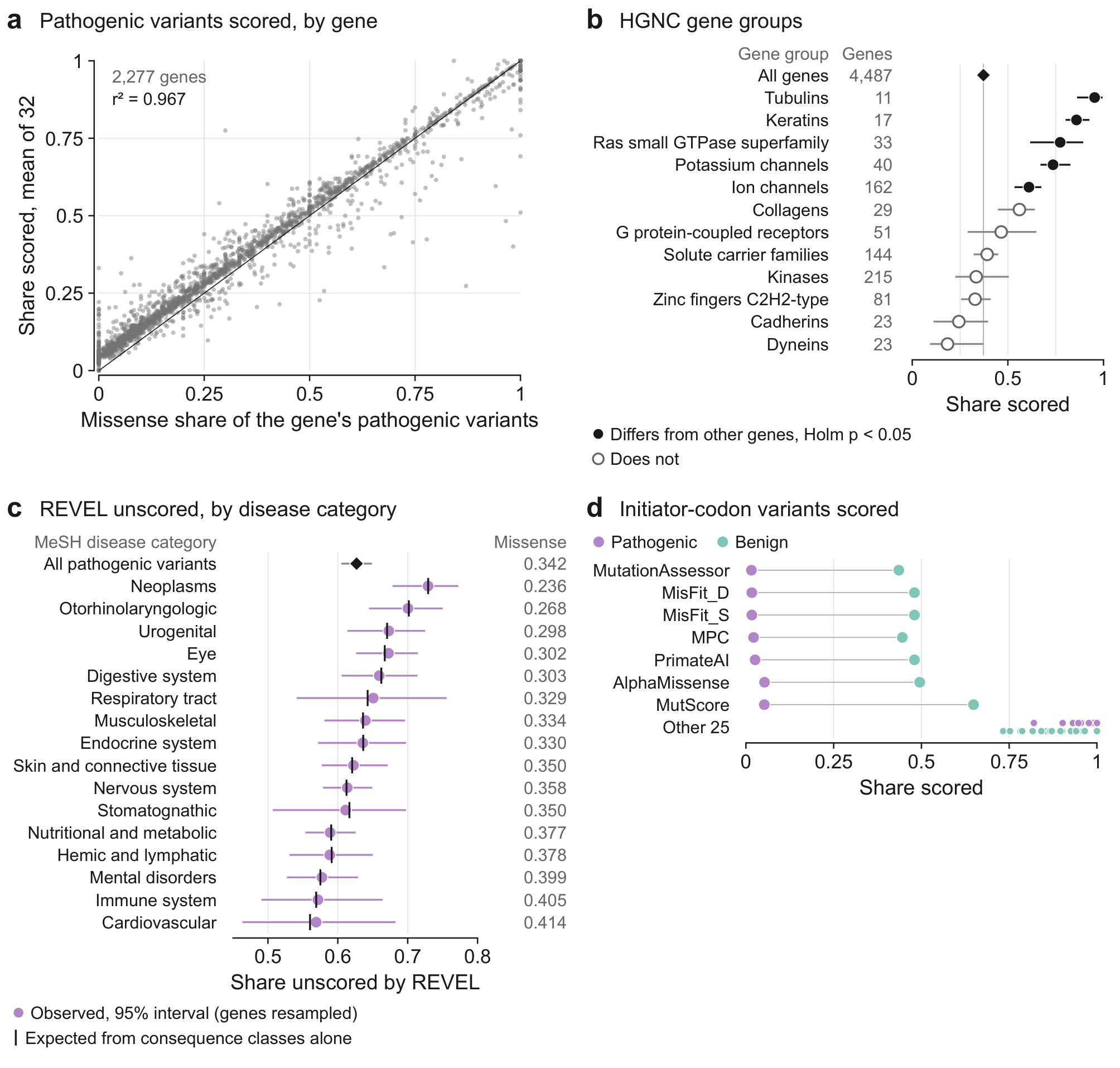

Extends Figure 2b, which gives each predictor's shares of pathogenic and benign variants scored: new here, where the declines of missense-scope predictors fall by gene, gene group, disease category and variant class. (a) For the 2,277 genes with at least ten pathogenic variants on the dbNSFP panel, the share scored (mean over 32 missense-scope predictors) against the share missense; line, equality. (b) Twelve HUGO Gene Nomenclature Committee (HGNC) gene groups, five of the ten that differ from other genes (the other five are parent groups or subgroups of these) and seven well-known families that do not: share scored; bar, the 95% interval of its difference from all other genes (genes resampled), drawn about the share; filled, Holm-adjusted permutation p < 0.05 among 246 groups; diamond, all 4,487 genes with a pathogenic variant. (c) Share of pathogenic ClinVar variants REVEL's own release leaves unscored, by Medical Subject Headings (MeSH) disease category, against the share its consequence classes alone predict; right, missense share. (d) Share of pathogenic and benign initiator-codon variants each predictor scores (counts and values in Table S32).

Alt text: Across 2,277 genes the share of pathogenic variants scored tracks the share that are missense (r² = 0.967). Tubulins, keratins, Ras GTPases, potassium channels and ion channels are scored more fully than other genes. REVEL's unscored share by disease category matches consequence-class expectation. Seven predictors score at most 5.3% of pathogenic initiator-codon variants.

#### Figure S18. Evo 2's scores and GERP's windows across the nine-species atlas

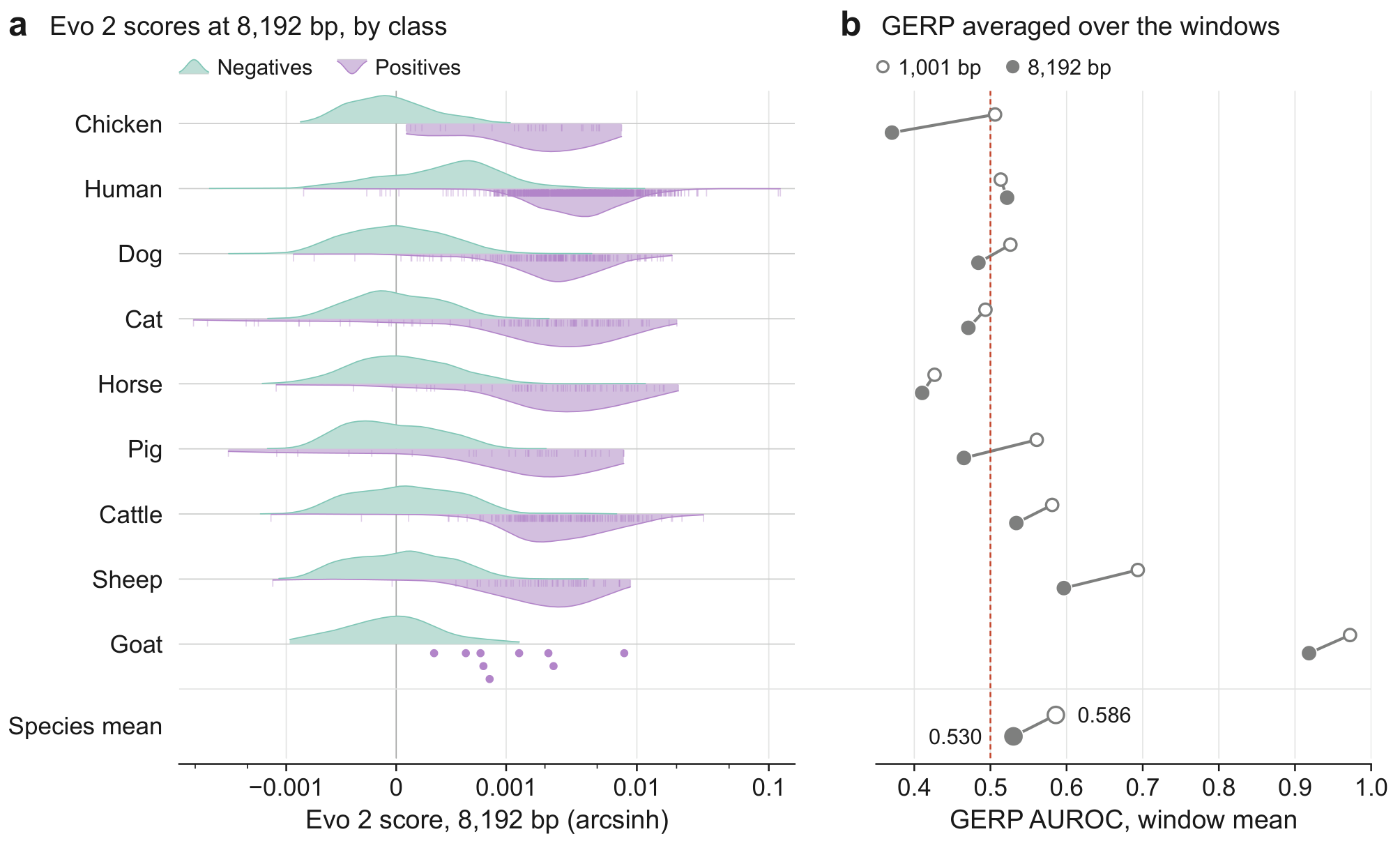

Extends Figure 5b, which gives each species' AUROC for Evo 2 and for GERP per base: new here, the score distributions behind Evo 2's AUROCs, and GERP averaged over Evo 2's two scoring windows. Species in the order of Figure 5. (a) Evo 2-40B's 8,192-bp window-mean score for negatives (above each line) and positives (below; ticks, each positive) on an arcsinh axis, linear within about ±0.0003; kernel densities scaled to their own peak, points where a class has fewer than ten variants; vertical line, zero (AUROCs in Table 1). (b) AUROC of Ensembl's GERP on the 9,529 variants both readouts and GERP score, averaged over the 1,001-bp (open) and 8,192-bp (filled) scoring windows (per-base values in Figure 5b and, on these variants, Table S4); dashed, chance; bottom row, nine-species means.

Alt text: Evo 2's 8,192-bp scores separate positives from negatives in all nine species. Averaged over the 1,001-bp or 8,192-bp window, GERP reads mean AUROCs of 0.586 and 0.530; the 8,192-bp average is below GERP's per-base value (Figure 5b; Table S4) in eight of nine species.

#### Figure S19. Evo 2 against the Zoonomia 241-mammal phyloP

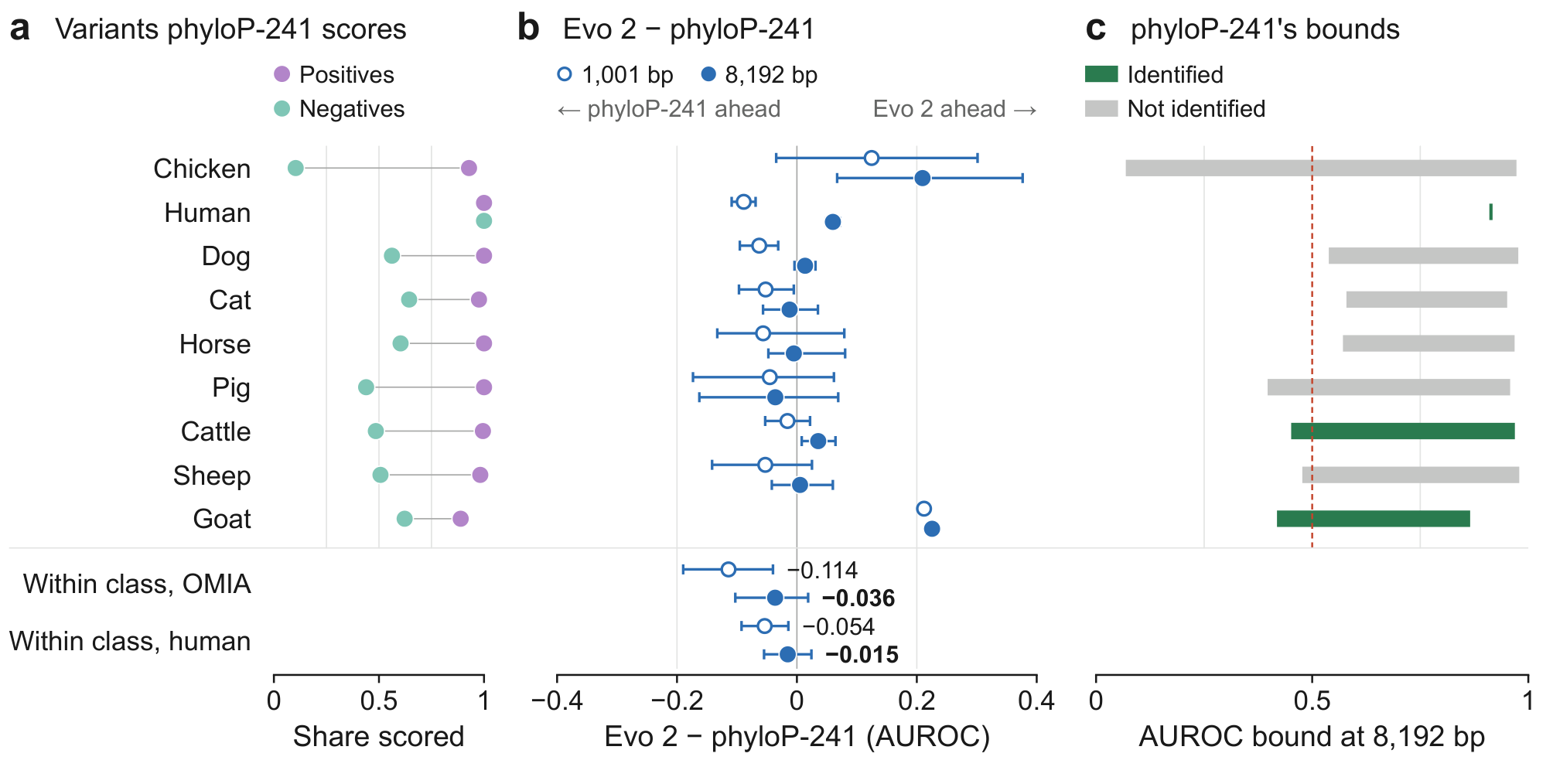

Extends Figure 5h, which gives Evo 2's species-mean margins over phyloP-241: new here, phyloP-241's reach, the margin and phyloP-241's sharp bound, species by species, in the order of Figure 5. (a) Zoonomia 241-mammal phyloP (phyloP-241), each variant lifted to GRCh38 through its assembly's chain: shares of positives and negatives scored (counts and the must-answer penalty in Table S38). (b) Evo 2 minus phyloP-241 on the 7,669 variants both readouts and phyloP-241 score, at 1,001 bp (open) and 8,192 bp (filled), with locus-bootstrap 95% intervals (goat, two positive loci: none; values in Table S38; species means in Figure 5h); below, the margin within consequence class, likewise. (c) phyloP-241's sharp bound at 8,192 bp, every unscored variant at both extremes, coloured in Figure 1's colours by the verdict when read against Evo 2's value in Figure 5b (replicate shares in Table S38); dashed, chance.

Alt text: phyloP-241 scores nearly all positives but about half the negatives in most species. Evo 2's margin over it varies by species and is negative within consequence class at 1,001 bp. At 8,192 bp the comparison is identified in cattle, human and, at point estimates, goat.

#### Figure S20. The consequence-matched arm of the nine-species atlas

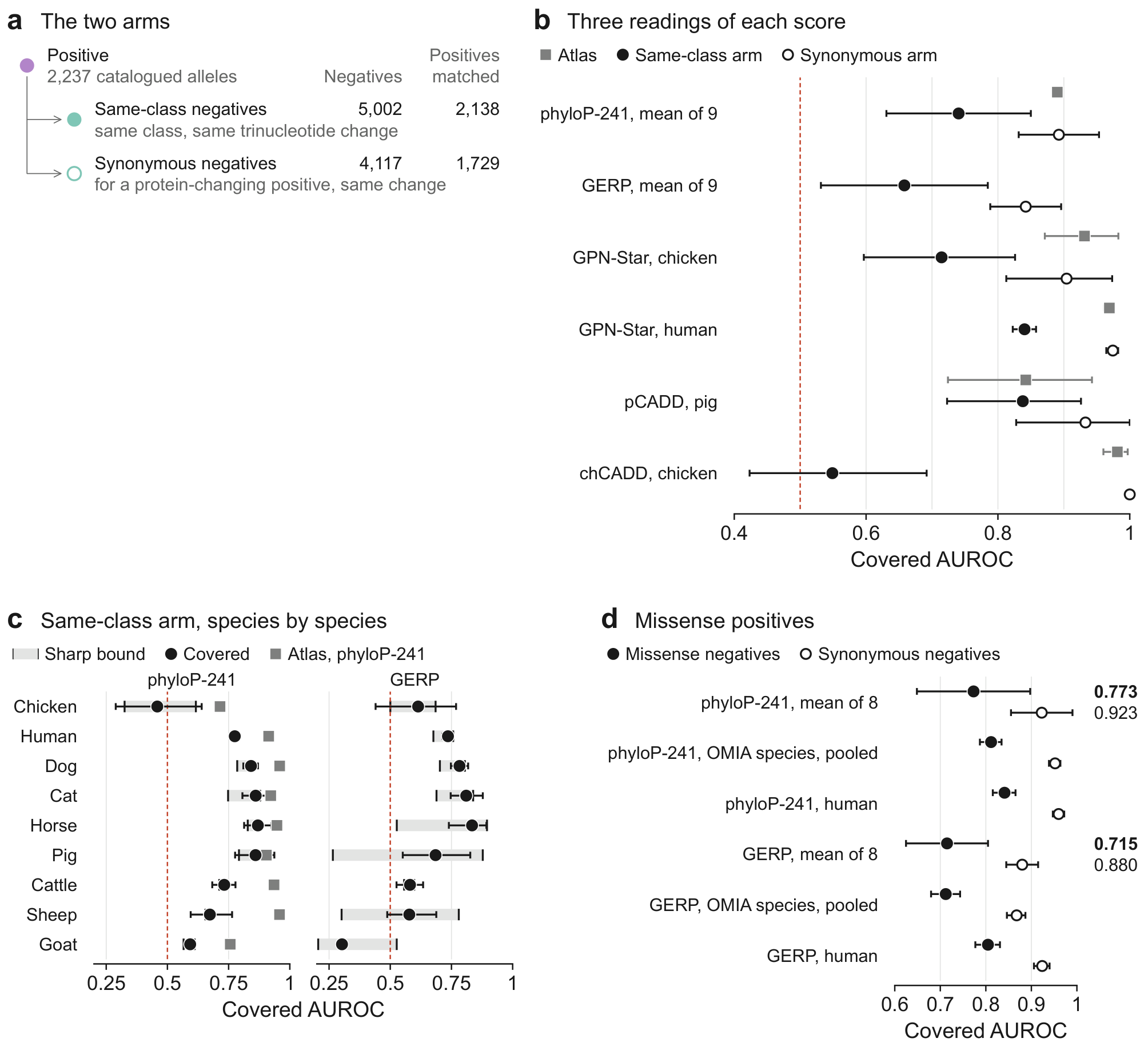

Extends Figures 5b and S19 to negatives matched on consequence class; Evo 2 is not scored. (a) Each of 2,237 catalogued alleles (positives; the atlas's 2,240 positives less three cat alleles, Note S32) faces population variants of its own consequence class and trinucleotide change (same-class arm) and, where it changes the protein, synonymous variants with that change (synonymous arm), up to five per positive and arm (one in human); the same-class arm's snpEff impact ordering is in Table S39's note. (b) Covered area under the receiver operating characteristic curve (AUROC) with the atlas's original negatives (square; GERP's in Figure 5b), on the same-class arm (filled) and on the synonymous arm (open): Zoonomia 241-mammal phyloP (phyloP-241) and GERP by the nine-species mean (95% t intervals), single-species scores (GPN-Star in chicken and human, pCADD, chCADD) with 95% locus-bootstrap intervals. (c) phyloP-241 and GERP on the same-class arm, species by species: sharp bound (band), covered AUROC (95% interval; goat, none) and phyloP-241's atlas value (square). (d) Missense positives against missense and against synonymous negatives: mean of eight species (goat, too few missense positives, has no cell), OMIA species pooled, human. Dashed, chance in b and c. Values for b and the pooled and human rows of d: Table S39.

Alt text: Matched on consequence class, conservation discriminates less: phyloP-241's species-mean covered AUROC falls from 0.890 on the atlas to 0.740 and GERP's from 0.830 (Figure 5b) to 0.658, while against synonymous negatives they read 0.892 and 0.842. chCADD falls from 0.981 to 0.549; pCADD holds at 0.838. Evo 2 is not scored on this panel.

### Supplementary Tables

**Table S1. Reach: what the conservation baseline, GERP, can be run on, and what its silence costs**

| **Species** | **n** | **positive scorable** | **negative scorable** | **class gap** | **must-answer penalty** |
| --- | --- | --- | --- | --- | --- |
| Goat | 99 | 0.889 | 0.800 | +0.089 | 0.081 |
| Chicken | 308 | 0.964 | 0.882 | +0.082 | 0.034 |
| Pig | 396 | 0.750 | 0.392 | +0.358 | 0.177 |
| Sheep | 616 | 0.732 | 0.732 | +0.000 | 0.169 |
| Horse | 781 | 0.930 | 0.441 | +0.489 | 0.245 |
| Cat | 1,365 | 0.936 | 0.742 | +0.194 | 0.106 |
| Cattle | 2,068 | 1.000 | 0.956 | +0.044 | 0.014 |
| Dog | 2,497 | 0.921 | 0.911 | +0.009 | 0.062 |
| Human | 3,000 | 0.963 | 0.957 | +0.007 | 0.029 |

Under the paper's rule the penalty is 0.1019 by the unweighted species mean and 0.2448 in horse. A no-call given the score's own median gives 0.0235 and 0.0100; drawn from its distribution, in expectation, 0.0400 and 0.0645. The class gap's Newcombe hybrid-score 95% interval excludes zero only in pig [+0.190, +0.481], horse [+0.397, +0.543], cat [+0.132, +0.234] and cattle [+0.022, +0.054], and a locus-resampled interval in the same four. In the other five species the interval is chicken [−0.063, +0.134], human [−0.007, +0.021], dog [−0.035, +0.040], sheep [−0.133, +0.105] and goat [−0.242, +0.220]. The must-answer penalty is the covered minus the must-answer AUROC. AUROC, area under the receiver operating characteristic curve; GERP, Genomic Evolutionary Rate Profiling.

Table S2. Atlas and eQTL discrimination at the three released checkpoints

| **Model** | **atlas macro AUROC, eight OMIA species (n = 8,109)** | **eQTL AUROC** | **95% CI** | **n (eQTL)** |
| --- | --- | --- | --- | --- |
| Evo 2-1B | 0.896 | 0.492 | [0.467, 0.518] | 2,000 |
| Evo 2-7B | 0.924 | 0.488 | [0.464, 0.514] | 2,000 |
| Evo 2-40B | 0.939 | 0.498 | [0.473, 0.523] | 2,000 |

Both AUROC columns are read at the 8,192-bp window-mean readout; the atlas macro is over the eight OMIA species, as on the capacity ladder of Note S10. The eQTL columns are on the 2,000-variant subpanel of Note S32 (1,000 candidate causal variants and 1,000 controls), not on the 20,000-variant panel of Note S25, where Evo 2-40B reads 0.488; intervals are percentile bootstraps over variants. AUROC, area under the receiver operating characteristic curve; CI, confidence interval; eQTL, expression quantitative trait locus; OMIA, Online Mendelian Inheritance in Animals.

**Table S3. An exploratory label-free check against the site-frequency spectrum**

| **MAF bin** | **n** | **mean MAF** | **mean deleteriousness** |
| --- | --- | --- | --- |
| ultra-rare | 5,607 | 0.00062 | 0.772 |
| rare | 5,624 | 0.00505 | 0.511 |
| low | 5,581 | 0.02436 | 0.319 |
| common | 5,596 | 0.12139 | 0.201 |
| major | 5,598 | 0.38986 | 0.178 |

Cattle (ARS-UCD1.2), 28,006 variants: mean Evo 2-40B deleteriousness at the 1,001-bp single-token readout, by minor-allele frequency (MAF) bin; the bins span MAF 0.00007 to 0.0020 (ultra-rare), 0.0020 to 0.0099 (rare), 0.010 to 0.050 (low), 0.050 to 0.299 (common) and 0.300 to 0.500 (major). Deleteriousness is logP(ref) − logP(alt).

**Table S4. Evo 2 versus GERP is readout-dependent (co-scorable variants)**

| **Species** | **n (co-scorable)** | **Evo 2 (1,001 bp)** | **Evo 2 (8,192 bp)** | **GERP** | **Δ (8,192 − GERP)** |
| --- | --- | --- | --- | --- | --- |
| Goat | 80 | 0.953 | 0.958 | 0.780 | +0.178 |
| Chicken | 274 | 0.862 | 0.956 | 0.725 | +0.231 |
| Pig | 168 | 0.814 | 0.826 | 0.751 | +0.075 |
| Sheep | 451 | 0.912 | 0.950 | 0.865 | +0.086 |
| Horse | 379 | 0.882 | 0.938 | 0.915 | +0.023 |
| Cat | 1,034 | 0.860 | 0.902 | 0.856 | +0.045 |
| Cattle | 1,985 | 0.900 | 0.974 | 0.825 | +0.149 |
| Dog | 2,278 | 0.895 | 0.974 | 0.886 | +0.089 |
| Human | 2,880 | 0.825 | 0.974 | 0.874 | +0.100 |
| **Macro** |  | 0.878 | 0.939 | 0.831 | +0.108 |

Per-species DeLong intervals (Note S21) exclude zero in six of the nine, so the nine of nine here is a sign count over point estimates. GERP, Genomic Evolutionary Rate Profiling.

**Table S5. Evo 2 against phyloP and GERP where a phyloP track is available**

| **Species** | **n (matched)** | **GERP** | **phyloP** | **phyloP − GERP** | **Evo 2 (8,192 bp)** | **Evo 2 (8,192 bp) − phyloP** | **Evo 2 (1,001 bp)** | **Evo 2 (1,001 bp) − phyloP** |
| --- | --- | --- | --- | --- | --- | --- | --- | --- |
| Chicken | 274 | 0.725 | 0.793 | +0.068 | 0.956 | +0.163 | 0.862 | +0.069 |
| Cattle | 1,921 | 0.828 | 0.931 | +0.103 | 0.978 | +0.047 | 0.909 | −0.022 |
| Human | 2,880 | 0.874 | 0.925 | +0.051 | 0.974 | +0.049 | 0.825 | −0.100 |

phyloP tracks: University of California, Santa Cruz (UCSC) phyloP77way (chicken, galGal6) and phyloP100way (human, hg38); for cattle (bosTau9), Zenodo record 13332541 (Roslin), over a Cactus 241-way alignment and a different species set. n (matched) counts the variants of Table S4 that the phyloP track also scores: all of them in chicken and human, and 1,921 of 1,985 in cattle (177 of 188 positives). GERP, Genomic Evolutionary Rate Profiling.

**Table S6. The readout gain split into a left-context-and-scorer term and an aggregation term**

| **Species** | **single token, 1,001 bp** | **single token, 8,192 bp** | **window mean, central 1,001 bp** | **window mean, 8,192 bp** |
| --- | --- | --- | --- | --- |
| Goat | 0.950 | 0.951 | 0.948 | 0.955 |
| Chicken | 0.863 | 0.916 | 0.952 | 0.947 |
| Pig | 0.843 | 0.866 | 0.878 | 0.861 |
| Sheep | 0.903 | 0.957 | 0.965 | 0.961 |
| Horse | 0.884 | 0.899 | 0.957 | 0.945 |
| Cat | 0.840 | 0.877 | 0.895 | 0.887 |
| Cattle | 0.893 | 0.931 | 0.969 | 0.973 |
| Dog | 0.890 | 0.929 | 0.979 | 0.979 |
| Human | 0.814 | 0.875 | 0.974 | 0.972 |

Terms as in Note S20, on its 2,599-variant subset; the single-token columns read about 0.5 kb and about 4 kb of left context, through different scorers.

**Table S6 (continued)**

|  | **left context and scorer term** | **aggregation term** | **total** |
| --- | --- | --- | --- |
| species mean (± SE) | +0.0359 ± 0.0066 | +0.0309 ± 0.0107 | +0.0668 ± 0.0149 |
| pooled, bootstrap mean (95% CI) | +0.0377 [+0.0286, +0.0469] | +0.0374 [+0.0267, +0.0483] | +0.0751 [+0.0611, +0.0898] |

SE, standard error; CI, confidence interval.

**Table S7. Precision–recall (AUPRC) at the 8,192-bp readout**

| **Species** | **base rate** | **AUPRC** | **AUPRC / base rate** |
| --- | --- | --- | --- |
| Goat | 0.092 | 0.735 | 8.0× |
| Chicken | 0.091 | 0.844 | 9.3× |
| Pig | 0.091 | 0.809 | 8.9× |
| Sheep | 0.091 | 0.886 | 9.7× |
| Horse | 0.093 | 0.822 | 8.9× |
| Cat | 0.090 | 0.849 | 9.5× |
| Cattle | 0.091 | 0.862 | 9.5× |
| Dog | 0.091 | 0.911 | 10.0× |
| Human | 0.500 | 0.971 | 1.9× |
| **Macro** | 0.137 | 0.854 | 6.3× (ratio of the means) |
| **Pooled** | 0.201 | 0.940 |  |

Evo 2-40B, 11,109 window-scored variants. AUPRC is average precision, the area under the precision–recall curve; the base rate is the no-skill baseline. The macro ratio divides that row's AUPRC by its base rate, on unrounded values; the mean of the nine per-species ratios is 8.4×.

**Table S8. Reach and must-answer AUROC for five human scores on ClinVar**

| **Score** | **reach (all)** | **reach (pos.)** | **reach (neg.)** | **covered** | **must-answer** | **penalty** | **missingness alone** |
| --- | --- | --- | --- | --- | --- | --- | --- |
| phastCons | 0.99997 | 0.99999 | 0.99996 | 0.8443 | 0.8443 | 0.00002 | 0.5000 |
| phyloP | 0.99997 | 0.99999 | 0.99996 | 0.9217 | 0.9217 | 0.00002 | 0.5000 |
| CADD | 0.99879 | 0.99935 | 0.99872 | 0.9914 | 0.9904 | 0.00095 | 0.5003 |
| REVEL | 0.17642 | 0.37313 | 0.14985 | 0.9575 | 0.5256 | 0.43192 | 0.6116 |
| AlphaMissense | 0.13493 | 0.33665 | 0.10768 | 0.9631 | 0.5168 | 0.44635 | 0.6145 |

Values are on the 1,434,335 ClinVar single-nucleotide variants at one review star or better (Additional file 5). Missingness alone is the AUROC of the indicator of whether a value was returned, discarding the values. Pair coverage ρ is the product of the two label reaches (phyloP and phastCons > 0.999, CADD 0.998; REVEL and AlphaMissense in Figure S16); phastCons ties 80% of its pathogenic variants at its maximum score, where 18% of its benign variants also sit, and 36% of its benign variants at its minimum. AUROC, area under the receiver operating characteristic curve; CADD, Combined Annotation Dependent Depletion; REVEL, Rare Exome Variant Ensemble Learner.

**Table S9. Reach and its class dependence for 49 dbNSFP predictors**

| **Reach band** | **n predictors** | **median reach** | **median covered** | **median must-answer** | **median penalty** |
| --- | --- | --- | --- | --- | --- |
| reach >99.9% | 5 | 1.0000 | 0.7866 | 0.7866 | 0.0000 |
| reach 70 to 99.9% | 13 | 0.9211 | 0.9218 | 0.8312 | 0.0486 |
| reach <70% | 31 | 0.5885 | 0.9375 | 0.6226 | 0.3149 |
| missense: reach >99.9% | 6 | 1.0000 | 0.8507 | 0.8507 | 0.0000 |
| missense: reach 70 to 99.9% | 43 | 0.9362 | 0.9410 | 0.8706 | 0.0479 |
| missense: reach <70% | 0 | – | – | – | – |

Class-gap intervals resample genes (2,000 draws). On the missense subset, Mendelian Clinically Applicable Pathogenicity (M-CAP) reaches the two classes most unequally, by +0.195. The two missingness rows count the 48 predictors that decline at least one variant. AUROC, area under the receiver operating characteristic curve; dbNSFP, database for nonsynonymous single-nucleotide polymorphisms' functional predictions.

**Table S9 (continued)**

| **Quantity** | **value** |
| --- | --- |
| class gap with a gene-clustered interval excluding zero | 44 of 49 |
| missingness alone outscoring must-answer, orientation-free | 30 of 48 |
| ... the same test taking the sign as given | 0 of 48 |
| predictors with a negative class gap, for which the signed test cannot fire | 34 of 49 |
| missense: class gap with a gene-clustered interval excluding zero | 39 of 49 |
| missense: ... and a gap of at least 0.05 in magnitude | 14 of 49 |
| missense: pairwise comparisons identified | 429 of 1,176 |
| missense: ... of which neither side is a conservation track | 180 of 429 |
| pairs the bound could identify for some covered AUROCs, both at least one half | 506 of 1,176 (1,174 missense) |
| comparisons ordered if unscored variants are no easier than scored | 131 of 1,176 (488 missense) |
| values adding to reach: lexicographic gain, gene-clustered interval excluding zero | 49 of 49 (49 of 49 missense) |

**Table S10. OMIA trait alleles and the effect of removing them**

| **Species** | **n** | **positives** | **trait positives** | **AUROC (full panel)** | **AUROC (disease only)** | **Δ (disease only − full panel)** |
| --- | --- | --- | --- | --- | --- | --- |
| Goat | 98 | 9 | 8 | 0.959 | – | – |
| Chicken | 308 | 28 | 20 | 0.954 | 1.000 | +0.0464 |
| Pig | 396 | 36 | 13 | 0.863 | 0.952 | +0.0888 |
| Sheep | 616 | 56 | 27 | 0.962 | 0.990 | +0.0276 |
| Horse | 766 | 71 | 44 | 0.941 | 0.949 | +0.0079 |
| Cat | 1,362 | 122 | 38 | 0.890 | 0.971 | +0.0805 |
| Cattle | 2,067 | 188 | 28 | 0.974 | 0.982 | +0.0077 |
| Dog | 2,496 | 227 | 25 | 0.970 | 0.979 | +0.0098 |
| Human | 3,000 | 1,500 | 0 | 0.974 | 0.974 | +0.0000 |
| Mean, eight species: human included, goat excluded |  |  |  | 0.941 | 0.975 | +0.0336 |
| Pooled | 11,109 | 2,237 | 203 | 0.973 | 0.985 | +0.0118 |

Trait positives are Online Mendelian Inheritance in Animals (OMIA) alleles for traits, such as coat and feather colour, morphology, production traits, blood groups, gait and resistance to infection, rather than disorders; goat keeps one disease allele, so it is not estimated. Removing them raises Evo 2-40B's species-mean AUROC at the 8,192-bp readout from 0.941 to 0.975 over the eight species, human among them, that keep at least five disease alleles. Over the same species and variants, the impact ordering moves from 0.954 to 0.974 and the coding flag from 0.906 to 0.928. AUROC, area under the receiver operating characteristic curve; OMIA, Online Mendelian Inheritance in Animals.

**Table S11. GERP through the same consequence-gradient estimator as Evo 2**

| **Consequence class** | **n positives (GERP)** | **Evo 2 (1,001-bp single token)** | **GERP** | **Evo 2 − GERP** |
| --- | --- | --- | --- | --- |
| nonsense (stop-gain) | 110 | 0.954 | 0.834 | +0.120 |
| splicing | 67 | 0.772 | 0.900 | −0.128 |
| missense | 310 | 0.896 | 0.879 | +0.017 |
| regulatory | 21 | 0.579 | 0.720 | −0.141 |

The same positive-count-weighted estimator as the Evo 2 gradient, with GERP in its place, on the variants GERP reaches and with cattle's positives, which the Evo 2 gradient leaves out, so the Evo 2 − GERP column subtracts AUROCs computed on different variant sets; the n column counts GERP's positives, and the Evo 2 column rests on the 112, 70, 324 and 18 of Note S26. AUROC, area under the receiver operating characteristic curve; GERP, Genomic Evolutionary Rate Profiling.

**Table S12. The human AUROC under other benign draws**

| **Negative set** | **n negatives** | **AUROC** | **95% CI** |
| --- | --- | --- | --- |
| ClinVar B/LB, all | 17,376 | 0.851 | [0.836, 0.866] |
| ClinVar B/LB, flagged coding | 4,377 | 0.811 | [0.795, 0.826] |
| ClinVar B/LB, not flagged coding | 12,999 | 0.865 | [0.850, 0.879] |

Evo 2-40B, the same 1,500 human positives at the 1,001-bp readout on GRCh38 against larger ClinVar Benign/Likely_benign (B/LB) draws; against the atlas panel's own 1,500 benign negatives they give 0.825. A negative is flagged coding when any of its ClinVar molecular-consequence terms contains missense, synonymous, nonsense, stop, frameshift, inframe or coding; "coding" also matches "non-coding transcript variant", so every negative carrying that term is flagged coding whatever its other terms, and the flagged row is not purely coding. Intervals are variant-level percentile bootstraps (2,000 draws). AUROC, area under the receiver operating characteristic curve; CI, confidence interval.

**Table S13. Six splicing datasets: Evo 2-1B, summed over 1,001 bp, against SpliceAI**

| **Dataset** | **n** | **disruptive** | **Evo 2-1B** | **SpliceAI** | **Evo 2 − SpliceAI** |
| --- | --- | --- | --- | --- | --- |
| BRCA1 | 1,386 | 223 | 0.808 | 0.986 | −0.178 |
| FAS exon 6 | 189 | 115 | 0.462 | 0.729 | −0.267 |
| MLH1 | 296 | 160 | 0.692 | 0.966 | −0.274 |
| POU1F1 exon 2 | 941 | 96 | 0.605 | 0.961 | −0.356 |
| RON exon 11 | 598 | 409 | 0.557 | 0.630 | −0.073 |
| WT1 exon 9 | 502 | 57 | 0.372 | 0.912 | −0.540 |

The distribution-free statement is the sign count: six of six, exact two-sided sign test p = 0.031. Against Pangolin the six per-assay deficits are likewise all negative, with an n-weighted mean of −0.259.

**Table S14. Reach by variant class, benchmark and background**

| **Predictor** | **panel** | **Essential Splice** | **Exon Near Junction** | **Intron Near Junction** | **Proximal Intron** | **Deep Exon** |
| --- | --- | --- | --- | --- | --- | --- |
| HAL PSI change (abs) | background | 0.000 | 1.000 | 0.000 | 0.000 | 1.000 |
| HAL PSI change (abs) | benchmark | 0.024 | 0.933 | 0.000 | 0.000 | 0.887 |
| MMSplice logit PSI change (abs) | benchmark | 1.000 | 1.000 | 1.000 | 0.862 | 1.000 |
| SPANR zPSI change (abs) | background | 0.989 | 0.991 | 0.991 | 0.989 | 0.976 |
| SPANR zPSI change (abs) | benchmark | 0.988 | 1.000 | 1.000 | 1.000 | 1.000 |
| SQUIRLS score | background | 0.907 | 0.908 | 0.909 | 0.908 | 0.903 |
| SQUIRLS score | benchmark | 0.679 | 0.975 | 0.975 | 0.999 | 1.000 |
| S-CAP sens minimum (rev) | background | 0.982 | 0.203 | 0.983 | 0.464 | 0.224 |
| S-CAP sens minimum (rev) | benchmark | 0.976 | 0.348 | 0.966 | 0.795 | 0.397 |
| SpliceAI delta max | background | 0.961 | 0.961 | 0.960 | 0.959 | 0.959 |
| ConSpliceML | background | 0.950 | 0.952 | 0.953 | 0.953 | 0.949 |

Reach is the share of variants a predictor scores (overall: Table S28); complete-reach rows are omitted. PSI, percent spliced in; HAL, Hexamer Additive Linear; MMSplice, Modular Modeling of Splicing; SPANR, Splicing-based Analysis of Variants; SQUIRLS, Super Quick Information-content Random-forest Learning of Splice variants; S-CAP, Splicing Clinically Applicable Pathogenicity prediction.

**Table S15. ClinVar consequence rungs at Evo 2-1B: discrimination, power and multiplicity**

| **Consequence** | **positive** | **negative** | **AUROC** | **95% CI** | **MDE** | **p (FWER)** |
| --- | --- | --- | --- | --- | --- | --- |
| 3′ UTR variant | 54 | 250 | 0.7023 | [0.6143, 0.7851] | 0.122 | < 0.00002 |
| 5′ UTR variant | 160 | 250 | 0.6497 | [0.5938, 0.7043] | 0.082 | 0.00042 |
| missense variant | 250 | 250 | 0.8496 | [0.8140, 0.8831] | 0.072 | < 0.00002 |
| non-coding transcript variant | 250 | 250 | 0.7167 | [0.6715, 0.7607] | 0.072 | < 0.00002 |
| nonsense | 250 | 250 | 0.7462 | [0.7020, 0.7885] | 0.072 | < 0.00002 |
| splice acceptor variant | 250 | 250 | 0.7852 | [0.7438, 0.8249] | 0.072 | < 0.00002 |
| splice donor variant | 250 | 250 | 0.7952 | [0.7546, 0.8339] | 0.072 | < 0.00002 |
| synonymous variant | 250 | 250 | 0.7902 | [0.7491, 0.8296] | 0.072 | < 0.00002 |

All values are the Evo 2-1B checkpoint summed over a 1,001-bp window. MDE is the Hanley–McNeil minimum detectable difference from chance at 80% power. p is adjusted for the family-wise error rate (FWER) across the eight rungs by Westfall–Young max-T over 50,000 permutations. Intervals resample variants within class and permutations ignore genes, so both are variant-level. AUROC, area under the receiver operating characteristic curve; CI, confidence interval; UTR, untranslated region.

**Table S16. Calibration transfer per species, leave-one-species-out**

| **Species** | **transferred isotonic** | **trivial global sigmoid** | **in-species oracle** |
| --- | --- | --- | --- |
| Goat | 0.0564 | 0.0481 | 0.0629 |
| Chicken | 0.0203 | 0.0211 | 0.0139 |
| Pig | 0.0375 | 0.0369 | 0.0137 |
| Sheep | 0.0531 | 0.0550 | 0.0136 |
| Horse | 0.0429 | 0.0441 | 0.0133 |
| Cat | 0.0412 | 0.0370 | 0.0058 |
| Cattle | 0.0609 | 0.0707 | 0.0084 |
| Dog | 0.0495 | 0.0491 | 0.0026 |
| Human | 0.1174 | 0.1339 | 0.0073 |
| **Macro** | **0.0533** | **0.0551** | **0.0157** |

Evo 2-40B at the 1,001-bp readout; equal-width 10-bin expected calibration error (ECE). Transferred isotonic, an isotonic map fitted on the six label-rich species (sheep, horse, cat, cattle, dog and human), the target left out where it is one of them; trivial global sigmoid, one two-parameter sigmoid fitted on the same donors; in-species oracle, an isotonic map fitted and scored on the target's own labels, in sample.

**Table S16 (continued)**

| **Calibration set size (variants)** | **mean transferred adaptive ECE** | **SD** |
| --- | --- | --- |
| 25 | 0.0578 | 0.0396 |
| 50 | 0.0322 | 0.0159 |
| 100 | 0.0303 | 0.0133 |
| 200 | 0.0395 | 0.0215 |
| 500 | 0.0353 | 0.0185 |
| 1,000 | 0.0410 | 0.0198 |
| 10,327 | 0.0470 | 0.0146 |

Learning curve: isotonic maps fitted on random draws from the six label-rich species (10,327 variants in all), scored on goat, chicken and pig by adaptive (equal-mass, 10-bin) ECE; mean and standard deviation (SD) over five draws per size and the three species.

**Table S17. Selective prediction per species**

| **Species** | **n** | **errors** | **refused** | **errors removed** | **lift** | **lift 95% CI** | **resolved** |
| --- | --- | --- | --- | --- | --- | --- | --- |
| Goat | 99 | 6 | 15 | 5 | 5.50 | [3.09, 9.90] | not assessed |
| Chicken | 308 | 12 | 47 | 4 | 2.18 | [0.53, 4.18] | no |
| Pig | 396 | 20 | 60 | 12 | 3.96 | [2.54, 5.66] | yes |
| Sheep | 616 | 21 | 93 | 15 | 4.73 | [3.37, 6.34] | yes |
| Horse | 781 | 31 | 117 | 14 | 3.01 | [1.90, 4.24] | yes |
| Cat | 1,365 | 61 | 205 | 30 | 3.27 | [2.52, 4.14] | yes |
| Cattle | 2,068 | 111 | 311 | 59 | 3.53 | [2.93, 4.19] | yes |
| Dog | 2,497 | 86 | 375 | 37 | 2.86 | [2.19, 3.57] | yes |
| Human | 3,000 | 545 | 450 | 140 | 1.71 | [1.52, 1.93] | yes |
| **Pooled** | **11,130** | **893** | **1,673** | **316** | **2.35** | **–** | **–** |

Each species refuses its least-confident 15% of calls on the 1,001-bp Platt posterior (Note S28). Lift is the share of errors removed over the share of calls refused, 1.0 being random refusal; a lift is resolved when its 95% CI excludes 1. The intervals resample variants within species, so they are variant-level and likely too narrow where positives share loci; goat, on two loci, is given no verdict (Note S12). CI, confidence interval.

**Table S18. Per-species operating point at the 0.5 decision threshold**

| **Species** | **sensitivity** | **sensitivity 95% CI** | **specificity** |
| --- | --- | --- | --- |
| Goat | 0.333 | [0.000, 0.667] | 1.000 |
| Chicken | 0.571 | [0.381, 0.760] | 1.000 |
| Pig | 0.444 | [0.275, 0.605] | 1.000 |
| Sheep | 0.786 | [0.673, 0.889] | 0.984 |
| Horse | 0.648 | [0.537, 0.754] | 0.992 |
| Cat | 0.584 | [0.496, 0.670] | 0.993 |
| Cattle | 0.718 | [0.654, 0.782] | 0.969 |
| Dog | 0.683 | [0.623, 0.744] | 0.994 |
| Human | 0.681 | [0.655, 0.705] | 0.956 |
| **Eight non-human (species mean)** | **0.596** | **–** | **–** |
| **Eight non-human (positive-weighted)** | **0.659** | **–** | **–** |

On the 1,001-bp Platt posterior, before any abstention. Intervals as in Table S17; goat's is descriptive only. CI, confidence interval.

**Table S19. The trust layer rebuilt at the 8,192-bp readout**

| **Species** | **ECE (equal-width, 10 bins)** | **errors** | **lift** |
| --- | --- | --- | --- |
| Goat | 0.0514 | 7 | 5.60 |
| Chicken | 0.0272 | 9 | 2.23 |
| Pig | 0.0219 | 11 | 3.05 |
| Sheep | 0.0095 | 17 | 5.12 |
| Horse | 0.0115 | 19 | 2.80 |
| Cat | 0.0132 | 34 | 2.95 |
| Cattle | 0.0232 | 57 | 5.03 |
| Dog | 0.0072 | 54 | 4.70 |
| Human | 0.0226 | 207 | 3.93 |

Built as the 1,001-bp layer of Note S28, but with refusals on an isotonic rather than a Platt posterior and every other species admitted as a donor; errors and lift as in Table S17. The 8,192-bp column refuses the round(0.15 n) least-confident calls in each species (1,665 in all), the 1,001-bp column those inside the strict 15% corridor (1,673); that column's 0.053 is the isotonic arm's calibration error, as Note S28 says, and the Platt posterior behind its other rows reads 0.055. Lifts move with the order of calls tied at the refusal boundary (Note S28): in the atlas table's row order, horse, cat, dog and human read 3.16, 3.14, 4.57 and 3.90. AUROC, area under the receiver operating characteristic curve; ECE, expected calibration error.

**Table S19 (continued)**

|  | **1,001-bp (Note S28)** | **8,192-bp** |
| --- | --- | --- |
| total errors | 893 (8.02%) | 415 (3.74%) |
| errors removed at 15% refusal | 316 (35.4%) | 253 (61.0%) |
| pooled lift | 2.35 | 4.07 |
| macro lift | 3.42 | 3.93 |
| macro ECE (equal-width, 10 bins) | 0.053 | 0.021 |
| macro error-detection AUROC (eight species, human excluded) | 0.736 | 0.771 |
| macro error-detection AUROC (all nine species) | 0.707 | 0.777 |

**Table S20. Evo 2-1B, summed over a 1,001-bp window, against purpose-built splice predictors, by variant class**

| **Variant class** | **n** | **disruptive** | **Evo 2-1B** | **95% CI** | **SpliceAI** | **Pangolin** | **Evo 2 − SpliceAI** | **p (FWER)** |
| --- | --- | --- | --- | --- | --- | --- | --- | --- |
| Essential Splice | 246 | 238 | 0.853 | [0.788, 0.911] | 0.968 | 0.928 | −0.116 | 0.01006 |
| Exon Near Junction | 434 | 148 | 0.554 | [0.496, 0.612] | 0.794 | 0.800 | −0.239 | < 0.00002 |
| Intron Near Junction | 643 | 188 | 0.721 | [0.674, 0.768] | 0.904 | 0.917 | −0.182 | < 0.00002 |
| Proximal Intron | 1,170 | 114 | 0.535 | [0.474, 0.596] | 0.656 | 0.656 | −0.121 | 0.00618 |
| Deep Exon | 1,419 | 372 | 0.471 | [0.438, 0.504] | 0.652 | 0.665 | −0.181 | < 0.00002 |

SpliceAI and Pangolin use roughly 10,000 bases of flanking sequence, 5,000 on each side, so the columns are not matched on input context and the comparison does not separate supervision from context. Essential Splice holds 8 non-disruptive variants (246 less 238), so its interval and p-value are descriptive (Note S2). Evo 2's 95% confidence intervals (CIs) are percentile bootstraps at B = 200,000, stratified within class so that no replicate can lose a class. The p-value is adjusted for the family-wise error rate (FWER) across the five classes by Westfall–Young max-T permutation over 50,000 draws; cells shown as "< 0.00002" sit on the 1/50,001 floor the test can resolve and are bounds rather than estimates. Differences are computed on unrounded AUROCs and can differ by 0.001 from the subtraction of the rounded columns. Pooled over all 3,912 variants Evo 2 reads 0.621 against SpliceAI 0.799 and Pangolin 0.803, a paired difference against SpliceAI of −0.1771 (95% CI [−0.2008, −0.1534], DeLong on the shared variants); that pooled comparison is not covered by the family-wise correction. AUROC, area under the receiver operating characteristic curve.

Table S21. Within-class discrimination at both readouts against GERP

| **Species** | **Class** | **Positives / negatives (all-classes rows: variants)** | **Evo 2, 1,001 bp** | **Evo 2, 8,192 bp** | **GERP** | **Readout gain** | **Margin at 8,192 bp** |
| --- | --- | --- | --- | --- | --- | --- | --- |
| Eight OMIA species, pooled | all classes | 6,649 variants | 0.678 [0.599, 0.759] | 0.773 [0.692, 0.853] | 0.652 [0.554, 0.746] | +0.095 [+0.029, +0.167] | +0.121 [+0.021, +0.239] |
| Human | all classes | 2,880 variants | 0.809 [0.775, 0.841] | 0.850 [0.815, 0.882] | 0.837 [0.799, 0.873] | +0.041 [+0.004, +0.078] | +0.012 [−0.030, +0.056] |
| Eight OMIA species, pooled | missense | 424 / 72 | 0.783 [0.718, 0.842] | 0.746 [0.671, 0.816] | 0.660 [0.573, 0.744] | −0.037 [−0.115, +0.039] | +0.086 [−0.022, +0.186] |
| Eight OMIA species, pooled | splice region | 12 / 16 | 0.578 [0.278, 0.861] | 0.906 [0.724, 1.000] | 0.875 [0.621, 1.000] | +0.328 [−0.012, +0.656] | +0.031 [−0.181, +0.250] |
| Eight OMIA species, pooled | intronic | 27 / 2,140 | 0.534 [0.395, 0.686] | 0.715 [0.560, 0.883] | 0.657 [0.478, 0.830] | +0.181 [+0.046, +0.347] | +0.058 [−0.122, +0.277] |
| Eight OMIA species, pooled | intergenic or flanking | 15 / 2,989 | 0.773 [0.649, 0.903] | 0.854 [0.744, 0.957] | 0.619 [0.420, 0.820] | +0.081 [+0.010, +0.170] | +0.236 [+0.047, +0.442] |
| Human | missense | 566 / 163 | 0.829 [0.794, 0.862] | 0.865 [0.831, 0.898] | 0.843 [0.801, 0.882] | +0.036 [+0.001, +0.072] | +0.023 [−0.020, +0.070] |
| Human | stop gained | 515 / 5 | 0.773 [0.658, 0.878] | 0.449 [0.144, 0.723] | 0.765 [0.684, 0.845] | −0.324 [−0.730, +0.011] | −0.317 [−0.570, −0.037] |
| Human | splice region | 20 / 104 | 0.637 [0.472, 0.806] | 0.934 [0.830, 1.000] | 0.819 [0.686, 0.938] | +0.296 [+0.102, +0.487] | +0.114 [−0.015, +0.263] |

Pairs are formed within species and consequence class (first snpEff term; splice donor and acceptor, 3′ and 5′ UTRs, and intergenic, upstream and downstream each pooled) on the 9,529 variants both readouts and GERP score, pooled by pair count; gain is 8,192 minus 1,001 bp, margin 8,192 bp minus GERP. Positive and negative counts include only species in which the class holds both labels. Class rows need at least ten positives and five negatives; the residual mixed group (intragenic, non-coding exon, and start or stop lost; 11 positives and 148 negatives) is not one class and has no row; human stop-gained, the one margin whose interval lies below zero, rests on five negatives, too few for its interval to be trusted, and human missense's readout gain clears zero by less than its Monte Carlo error. Square brackets hold 95% locus-bootstrap intervals (positive-bearing 100-kb loci drawn whole and negatives individually, B = 2,000); gains and margins are taken on unrounded AUROCs and can differ by 0.001 from the rounded columns. AUROC, area under the receiver operating characteristic curve; GERP, Genomic Evolutionary Rate Profiling; OMIA, Online Mendelian Inheritance in Animals; UTR, untranslated region.

Table S22. Calibrated PP3 and BP4 evidence on ClinVar missense variants, by class

| **Tool** | **ClinVar class** | **Missense variants** | **PP3** | **BP4** | **Indeterminate** | **Unscored** |
| --- | --- | --- | --- | --- | --- | --- |
| REVEL | pathogenic | 58,394 | 82.1% | 3.4% | 14.1% | 0.4% |
| REVEL | benign | 145,797 | 3.2% | 79.7% | 14.7% | 2.4% |
| AlphaMissense | pathogenic | 58,394 | 71.4% | 3.7% | 23.1% | 1.7% |
| AlphaMissense | benign | 145,797 | 2.7% | 73.5% | 16.8% | 7.0% |
| CADD | pathogenic | 58,394 | 75.7% | 6.8% | 17.5% | 0.0% |
| CADD | benign | 145,797 | 8.3% | 80.1% | 11.6% | 0.0% |
| phyloP | pathogenic | 58,394 | 58.4% | 7.7% | 33.9% | 0.0% |
| phyloP | benign | 145,797 | 7.0% | 64.3% | 28.7% | 0.0% |

PP3 and BP4 count evidence at any calibrated strength [18,19]. Indeterminate scores fall between the two bounds; unscored variants receive no evidence. CADD, Combined Annotation Dependent Depletion; REVEL, Rare Exome Variant Ensemble Learner; PP3 and BP4, computational pathogenic and benign evidence.

Table S23. Missense AUROC on ClinGen expert-panel labels, with and without PP3/BP4-dependent labels

| **Tool** | **Pathogenic / benign scored** | **AUROC, all labels** | **AUROC, labels standing without PP3/BP4** | **Difference [95% CI]** |
| --- | --- | --- | --- | --- |
| REVEL | 3,023 / 1,137 | 0.955 | 0.949 | +0.007 [+0.004, +0.010] |
| AlphaMissense | 3,017 / 1,095 | 0.936 | 0.931 | +0.005 [+0.002, +0.009] |
| CADD | 3,024 / 1,144 | 0.898 | 0.894 | +0.004 [+0.001, +0.008] |
| phyloP | 3,024 / 1,144 | 0.809 | 0.801 | +0.008 [+0.002, +0.015] |

A label depends on PP3/BP4 when it applied them and reaches its class only with those points (Note S8); the difference is the AUROC on all labels minus that on labels standing without PP3/BP4, with genes resampled (B = 2,000); differences are taken on unrounded AUROCs. AUROC, area under the receiver operating characteristic curve; CI, confidence interval; CADD, Combined Annotation Dependent Depletion; REVEL, Rare Exome Variant Ensemble Learner; PP3 and BP4, computational pathogenic and benign evidence.

Table S24. Per-species provenance of the atlas

| **Species** | **Assembly** | **GERP track** | **Gene models / snpEff database** | **Source of negatives** | **Positives** | **Genes** | **OMIA phenotypes** | **Trait alleles** |
| --- | --- | --- | --- | --- | --- | --- | --- | --- |
| Goat | ARS1 (GCA_001704415.1) | Ensembl 110, 91 mammals | Ensembl 112 / ARS1 | Ensembl variation 110 | 9 | 2 | 3 | 8 |
| Chicken | GRCg6a (GCA_000002315.5) | Ensembl 106, 27 sauropsids | Ensembl 105 / GRCg6a | Ensembl variation 104 | 28 | 21 | 25 | 20 |
| Pig | Sscrofa11.1 (GCA_000003025.6) | Ensembl 110, 91 mammals | Ensembl 112 / Sscrofa11.1 | PigGTEx v0 genotypes | 36 | 29 | 28 | 13 |
| Sheep | Oar_rambouillet_v1.0 (GCA_002742125.1) | Ensembl 110, 91 mammals | Ensembl 106 / Oar_rambouillet_v1.0 | Ensembl variation 109 | 56 | 37 | 50 | 27 |
| Horse | EquCab3.0 (GCA_002863925.1) | Ensembl 110, 91 mammals | Ensembl 112 / EquCab3.0 | Ensembl variation 110 | 71 | 33 | 43 | 44 |
| Cat | F.catus_Fca126_mat1.0 (GCA_018350175.1) | Ensembl 114, 91 mammals | Ensembl 114 / F.catus_Fca126_mat1.0 | Ensembl variation 114 | 125 | 76 | 86 | 39 |
| Cattle | ARS-UCD1.2 (GCA_002263795.2) | Ensembl 110, 91 mammals | Ensembl 99 / ARS-UCD1.2.99 (pre-built) | Ensembl variation 110 | 188 | 144 | 149 | 28 |
| Dog | CanFam3.1 (GCA_000002285.2) | Ensembl 104, 90 mammals | Ensembl 104 / CanFam3.1 | Ensembl variation 104 | 227 | 178 | 189 | 25 |
| Human | GRCh38 (GCA_000001405.15) | Ensembl 110, 91 mammals | Ensembl 112 / GRCh38 | ClinVar benign, 1:1 | 1,500 | 843 | – | – |

Genes and OMIA phenotypes count the distinct genes and phenotypes among each species' positives. The trait-allele column counts trait alleles among all 740 non-human positives (204); the 737 scored at 8,192 bp hold 203 (Table S10). GERP, Genomic Evolutionary Rate Profiling; OMIA, Online Mendelian Inheritance in Animals; PigGTEx, Pig Genotype-Tissue Expression.

Table S25. Per-predictor provenance on the 49-predictor panel

| **Predictor** | **Scope** | **Training labels** | **Allele-frequency input** | **Covered AUROC** | **Within-gene AUROC** |
| --- | --- | --- | --- | --- | --- |
| SIFT | missense | unsupervised | no | 0.899 | 0.872 |
| SIFT4G | missense | unsupervised | no | 0.897 | 0.897 |
| Polyphen2_HDIV | missense | clinical assertions | no | 0.900 | 0.883 |
| Polyphen2_HVAR | missense | clinical assertions | no | 0.919 | 0.905 |
| MutationTaster | gene-internal variants | clinical assertions | unverified | 0.991 | 0.990 |
| MutationAssessor | missense | unsupervised | no | 0.922 | 0.907 |
| PROVEAN | missense | unsupervised | no | 0.917 | 0.898 |
| VEST4 | missense | clinical assertions | unverified | 0.946 | 0.932 |
| MetaSVM | missense | clinical assertions | yes | 0.936 | 0.886 |
| MetaLR | missense | clinical assertions | yes | 0.937 | 0.903 |
| MetaRNN | rare nsSNVs, all scored | clinical assertions | yes | 0.987 | 0.968 |
| M-CAP | rare missense | clinical assertions | indirect | 0.937 | 0.903 |
| REVEL | missense | clinical assertions | no | 0.971 | 0.944 |
| MutPred2 | missense | clinical assertions | no | 0.961 | 0.944 |
| MVP | missense | clinical assertions | no | 0.949 | 0.913 |
| gMVP | missense | clinical assertions | no | 0.967 | 0.944 |
| MisFit_D | missense | population data | no | 0.945 | 0.934 |
| MisFit_S | missense | population data | yes | 0.880 | 0.928 |
| MPC | missense | clinical assertions | no | 0.854 | 0.861 |
| PrimateAI | missense | population data | no | 0.886 | 0.894 |
| DEOGEN2 | missense | clinical assertions | no | 0.944 | 0.910 |
| BayesDel_addAF | SNVs and indels | clinical assertions | yes | 0.990 | 0.984 |
| BayesDel_noAF | SNVs and indels | clinical assertions | no | 0.983 | 0.976 |
| ClinPred | missense | clinical assertions | yes | 0.983 | 0.969 |
| LIST-S2 | missense | clinical assertions | no | 0.891 | 0.862 |
| VARITY_R | rare missense | clinical assertions | no | 0.968 | 0.950 |
| VARITY_ER | extremely rare missense | clinical assertions | no | 0.964 | 0.944 |
| VARITY_R_LOO | rare missense | clinical assertions | no | 0.967 | 0.947 |
| VARITY_ER_LOO | extremely rare missense | clinical assertions | no | 0.963 | 0.943 |
| ESM1b | missense | unsupervised | no | 0.957 | 0.963 |
| AlphaMissense | missense | population data | no | 0.960 | 0.954 |
| PHACTboost | missense | clinical assertions | no | 0.981 | 0.965 |
| MutFormer | missense | clinical assertions | indirect | 0.935 | 0.883 |
| MutScore | missense | clinical assertions | no | 0.979 | 0.959 |
| popEVE | missense | population data | no | 0.924 | 0.939 |
| CADD_raw | all SNVs | evolutionary proxy | no | 0.968 | 0.976 |
| DANN | all SNVs | evolutionary proxy | no | 0.833 | 0.773 |
| fathmm-XF_coding | coding SNVs | clinical assertions | no | 0.752 | 0.522 |
| Eigen-raw_coding | coding SNVs | unsupervised | no | 0.947 | 0.924 |
| Eigen-PC-raw_coding | coding SNVs | unsupervised | no | 0.922 | 0.882 |
| GERP++_RS | conservation track | none (conservation) | no | 0.801 | 0.693 |
| GERP_92_mammals | conservation track | none (conservation) | no | 0.763 | 0.673 |
| phyloP100way_vertebrate | conservation track | none (conservation) | no | 0.838 | 0.743 |
| phyloP470way_mammalian | conservation track | none (conservation) | no | 0.820 | 0.752 |
| phyloP17way_primate | conservation track | none (conservation) | no | 0.722 | 0.585 |
| phastCons100way_vertebrate | conservation track | none (conservation) | no | 0.787 | 0.685 |
| phastCons470way_mammalian | conservation track | none (conservation) | no | 0.770 | 0.662 |
| phastCons17way_primate | conservation track | none (conservation) | no | 0.691 | 0.580 |
| bStatistic | conservation track | none (conservation) | no | 0.549 | 0.477 |

Scope, training labels and allele-frequency input are from each tool's publication or documentation and the dbNSFP 5.3.1a description (unverified, not confirmed there; indirect, via other predictors' scores; the 'no' of Polyphen2_HDIV, Polyphen2_HVAR, MutPred2 and PHACTboost rests on published feature categories, not a full feature list); both AUROCs are on the whole panel, and within-gene AUROC pools pairs within a gene; reach, penalty, missense values and bounds are in figures/dbnsfp_49_provenance.csv (Additional file 3). Primary references, in row order: SIFT [1], SIFT4G [173], PolyPhen-2 [2], MutationTaster [125], MutationAssessor [174], PROVEAN [175], VEST4 [176], MetaSVM and MetaLR [177], MetaRNN [178], M-CAP [179], REVEL [40], MutPred2 [180], MVP [181], gMVP [182], MisFit [183], MPC [184], PrimateAI [185], DEOGEN2 [186], BayesDel [187], ClinPred [188], LIST-S2 [144], VARITY [189], ESM1b [9], AlphaMissense [41], PHACTboost [190], MutFormer [191], MutScore [192], popEVE [193], CADD [39], DANN [194], fathmm-XF [195], Eigen and Eigen-PC [196], GERP++ [43], GERP [42], phyloP [37], phastCons [38] and bStatistic [197]. SNV, single-nucleotide variant; nsSNV, nonsynonymous SNV; AUROC, area under the receiver operating characteristic curve; dbNSFP, database for nonsynonymous single-nucleotide polymorphisms' functional predictions.

Table S26. Covered-AUROC shift of the 49 predictors on newer ClinVar labels

| **Predictor** | **ClinVar in training** | **Shift** |
| --- | --- | --- |
| SIFT | no | +0.0265 |
| SIFT4G | no | +0.0379 |
| Polyphen2_HDIV | no | +0.0491 |
| Polyphen2_HVAR | no | +0.0394 |
| MutationTaster | yes | −0.0037 |
| MutationAssessor | no | +0.0451 |
| PROVEAN | no | +0.0330 |
| VEST4 | no | −0.0030 |
| MetaSVM | no | +0.0025 |
| MetaLR | no | +0.0035 |
| MetaRNN | yes | −0.0111 |
| M-CAP | no | +0.0293 |
| REVEL | test | +0.0050 |
| MutPred2 | no | +0.0168 |
| MVP | yes | +0.0210 |
| gMVP | yes | +0.0072 |
| MisFit_D | no | +0.0156 |
| MisFit_S | no | −0.0133 |
| MPC | yes | +0.0059 |
| PrimateAI | no | +0.0012 |
| DEOGEN2 | no | +0.0143 |
| BayesDel_addAF | yes | −0.0042 |
| BayesDel_noAF | yes | +0.0047 |
| ClinPred | yes | −0.0135 |
| LIST-S2 | yes | +0.0233 |
| VARITY_R | yes | +0.0101 |
| VARITY_ER | yes | +0.0159 |
| VARITY_R_LOO | yes | +0.0152 |
| VARITY_ER_LOO | yes | +0.0176 |
| ESM1b | no | +0.0154 |
| AlphaMissense | calibration | +0.0067 |
| PHACTboost | yes | −0.0074 |
| MutFormer | no | +0.0001 |
| MutScore | yes | +0.0054 |
| popEVE | evaluation | +0.0217 |
| CADD_raw | no | +0.0172 |
| DANN | no | +0.0825 |
| fathmm-XF_coding | no | +0.0702 |
| Eigen-raw_coding | no | +0.0300 |
| Eigen-PC-raw_coding | no | +0.0373 |
| GERP++_RS | no | +0.0727 |
| GERP_92_mammals | no | +0.0676 |
| phyloP100way_vertebrate | no | +0.0486 |
| phyloP470way_mammalian | no | +0.0465 |
| phyloP17way_primate | no | +0.0960 |
| phastCons100way_vertebrate | no | +0.0563 |
| phastCons470way_mammalian | no | +0.0624 |
| phastCons17way_primate | no | +0.0249 |
| bStatistic | no | −0.0213 |

Shift is the covered AUROC on newer labels minus that on the disjoint older ones (Note S5); test, calibration and evaluation mark predictors that used ClinVar only for that purpose. AUROC, area under the receiver operating characteristic curve.

Table S27. Calibrated PP3 and BP4 levels as likelihood ratios

| **Tool and level** | **Likelihood ratio, all labels (95% CI)** | **Stratum closest to target** |
| --- | --- | --- |
| REVEL, BP4 very strong | 0, no pathogenic variant at the level | 0, all labels |
| REVEL, BP4 strong | 0.0032 (0.0015 to 0.0050) | 0.0053, added 2021 to 2023 |
| REVEL, BP4 moderate | 0.024 (0.021 to 0.028) | 0.034, added 2021 to 2023 |
| REVEL, BP4 supporting | 0.14 (0.12 to 0.16) | 0.19, one star |
| REVEL, unscored | 0.18 (0.11 to 0.27) | – |
| REVEL, PP3 supporting | 6.81 (6.00 to 7.77) | 4.46, added 2021 to 2023 |
| REVEL, PP3 moderate | 30.9 (26.9 to 35.1) | 18.9, added 2021 to 2023 |
| REVEL, PP3 strong | 217 (178 to 263) | 149, added 2021 to 2023 |
| AlphaMissense, BP4 moderate | 0.021 (0.018 to 0.023) | 0.025, added 2021 to 2023 |
| AlphaMissense, BP4 supporting | 0.15 (0.13 to 0.16) | 0.17, added after 2023 |
| AlphaMissense, unscored | 0.25 (0.15 to 0.38) | – |
| AlphaMissense, PP3 supporting | 9.37 (8.38 to 10.5) | 6.95, added 2021 to 2023 |
| AlphaMissense, PP3 moderate | 23.3 (20.9 to 25.9) | 15.2, added 2021 to 2023 |
| AlphaMissense, PP3 strong | 111 (93.9 to 132) | 73.7, added 2021 to 2023 |
| CADD, BP4 strong | 0.0039 (0.0023 to 0.0058) | 0.0078, added 2021 to 2023 |
| CADD, BP4 moderate | 0.029 (0.026 to 0.033) | 0.033, added 2021 to 2023 |
| CADD, BP4 supporting | 0.29 (0.26 to 0.32) | 0.38, added after 2023 |
| CADD, unscored | 0.31 (0 to 0.49) | – |
| CADD, PP3 supporting | 5.76 (5.37 to 6.21) | 4.18, added 2021 to 2023 |
| CADD, PP3 moderate | 15.2 (13.9 to 16.5) | 9.11, added 2021 to 2023 |
| phyloP, BP4 moderate | 0.042 (0.037 to 0.049) | 0.062, carried by the 2021 release |
| phyloP, BP4 supporting | 0.19 (0.18 to 0.21) | 0.22, added 2021 to 2023 |
| phyloP, unscored | 0, no pathogenic variant at the level | – |
| phyloP, PP3 supporting | 7.98 (7.32 to 8.75) | 5.83, added 2021 to 2023 |
| phyloP, PP3 moderate | 11.2 (9.39 to 13.6) | 7.88, added 2021 to 2023 |

Ratios are on the 204,191 ClinVar missense variants at one review star or more, genes resampled (Note S8); declined variants form the unscored level, and the indeterminate level, between the BP4 and PP3 thresholds, is not shown. Stratum closest to target: of six strata (all labels; labels carried by the 2021 release, added 2021 to 2023 or added after 2023; one star; two stars or more), the one whose ratio is nearest its level's point-system odds (Note S8), all labels where strata tie. CADD's unscored ratio rests on one pathogenic variant, so its resampled interval understates its uncertainty; phyloP's unscored level has none. CI, confidence interval; PP3 and BP4, computational pathogenic and benign evidence. The intervals are percentile limits over 2,000 resamples of genes, and their ends move by about 0.01 between runs.

Table S28. What eight splice predictors answer, on a benchmark and in deployment

| **Predictor** | **Benchmark reach** | **Background reach** | **Standardised reach** | **Gap** |
| --- | --- | --- | --- | --- |
| Pangolin | 1.000 | 1.000 | 1.000 | +0.000 |
| MMSplice | 0.959 | 1.000 | 1.000 | −0.041 |
| SPANR | 0.999 | 0.984 | 0.985 | +0.014 |
| SpliceAI | 1.000 | 0.959 | 0.959 | +0.041 |
| ConSpliceML | 1.000 | 0.951 | 0.951 | +0.049 |
| SQUIRLS | 0.973 | 0.906 | 0.906 | +0.066 |
| HAL | 0.427 | 0.435 | 0.474 | −0.047 |
| S-CAP | 0.641 | 0.389 | 0.466 | +0.175 |

The gap is benchmark reach minus background reach standardised to the benchmark's variant-class mix.

Rows follow background reach, ties broken by benchmark reach. HAL, Hexamer Additive Linear; MMSplice, Modular Modeling of Splicing; SPANR, Splicing-based Analysis of Variants; SQUIRLS, Super Quick Information-content Random-forest Learning of Splice variants; S-CAP, Splicing Clinically Applicable Pathogenicity prediction.

Table S29. What conventions for unscored variants decide among the 1,176 comparisons of the 49 predictors, set against the bound

| **Quantity** | **Missense subset** | **Whole panel** |
| --- | --- | --- |
| Variants | 202,643 | 328,328 |
| Comparisons identified by the bound | 429 | 83 |
| Ordered under every common score for declined variants | 676 | 202 |
| Ordered by the least-deleterious fill, point estimates | 1,176 | 1,176 |
| Resolved by the least-deleterious fill | 957 | 1,055 |
| Resolved by the fill and identified by the bound | 427 | 82 |
| Fill orders that reverse under the most-deleterious fill (both orders resolved) | 205 (67) | 721 (608) |
| Spearman correlation between the two fills' rankings | 0.809 | −0.320 |
| Resolved on shared variants (kept by Holm's procedure) | 1,113 (1,057) | 1,118 (1,065) |
| Resolved on shared variants and identified by the bound | 427 | 82 |
| Shared-variant order opposite to the least-deleterious-fill order (both orders resolved) | 188 (104) | 619 (540) |
| Shared-variant order opposite to the side-by-side order (resolved on shared variants) | 6 (0) | 58 (35) |

The least-deleterious fill gives each unscored variant the predictor's lowest dbNSFP rank score on the panel, the most-deleterious fill its highest; on the whole panel the fills also cover variants outside a predictor's declared scope. Shared variants are those both predictors score. A difference in AUROC is resolved when its 95% interval over 2,000 gene resamples excludes zero. Holm's procedure runs over each panel's 1,176 shared-variant differences, on two-sided normal p-values from the resamples' standard deviation. Side by side compares the two covered AUROCs. An identified comparison cannot reverse under any fill or on shared variants (Note S36). Per-pair and per-predictor values are in Additional file 3. AUROC, area under the receiver operating characteristic curve; dbNSFP, database for nonsynonymous single-nucleotide polymorphisms' functional predictions.

Table S30. Variants that receive no PP3 or BP4 points from a calibrated score: outside scope, unscored or indeterminate

| **Variant set** | **Score** | **Variants** | **Outside scope** | **Unscored** | **Indet.** | **Zero points** | **BP4** | **PP3** |
| --- | --- | --- | --- | --- | --- | --- | --- | --- |
| ClinVar single-nucleotide variants of uncertain significance | AlphaMissense | 2,232,064 | 134,277 (6.0%) | 57,494 (2.6%) | 744,749 (33.4%) | 936,520 (42.0%) | 994,888 (44.6%) | 300,656 (13.5%) |
| The missense variants among them | AlphaMissense | 2,097,787 | 0 (0.0%) | 57,494 (2.7%) | 744,749 (35.5%) | 802,243 (38.2%) | 994,888 (47.4%) | 300,656 (14.3%) |
| ClinVar single-nucleotide variants of uncertain significance in the 84 secondary-findings genes | AlphaMissense | 116,064 | 12,213 (10.5%) | 1,857 (1.6%) | 39,700 (34.2%) | 53,770 (46.3%) | 39,102 (33.7%) | 23,192 (20.0%) |
| ClinVar single-nucleotide variants of uncertain significance in the 84 secondary-findings genes | CADD | 116,064 | 12,213 (10.5%) | 0 (0.0%) | 25,556 (22.0%) | 37,769 (32.5%) | 49,268 (42.4%) | 29,027 (25.0%) |
| Every possible missense substitution in the 84 secondary-findings genes | AlphaMissense | 767,747 | 0 (0.0%) | 16,014 (2.1%) | 306,482 (39.9%) | 322,496 (42.0%) | 254,687 (33.2%) | 190,564 (24.8%) |
| Every possible missense substitution in the 84 secondary-findings genes | CADD | 767,747 | 0 (0.0%) | 0 (0.0%) | 160,653 (20.9%) | 160,653 (20.9%) | 443,612 (57.8%) | 163,482 (21.3%) |

Shares are of the row's variants; zero points is the sum of the three columns before it. ClinVar variants are those of uncertain significance at one review star or better in the release that built the reach panel, and a variant is within missense scope when ClinVar's first-listed consequence is missense. AlphaMissense is its canonical-transcript release read by GRCh38 position at its calibrated intervals [19]: scores at or below 0.169 count as BP4 and at or above 0.792 as PP3, and where the release holds several rows for one substitution the last is read. CADD is v1.7, read at the intervals calibrated for an earlier CADD release [18], read in the 84 genes only. Possible substitutions were enumerated from the MANE Select coding sequences of the 84 genes of ACMG SF v3.3, not from either score's file. REVEL was not read on variants of uncertain significance.

Column headings: outside scope, outside missense scope; unscored, unscored within scope; Indet., indeterminate; BP4 and PP3, at any strength. CADD, Combined Annotation Dependent Depletion; REVEL, Rare Exome Variant Ensemble Learner; MANE, Matched Annotation from NCBI and EMBL-EBI; ACMG SF v3.3, the American College of Medical Genetics and Genomics list of genes for reporting secondary findings, version 3.3; PP3 and BP4, computational pathogenic and benign evidence.

Table S31. Secondary-findings genes in which AlphaMissense's canonical-transcript release leaves possible missense substitutions unscored

| **Gene** | **MANE Select transcript** | **Possible missense substitutions** | **Unscored by AlphaMissense, canonical-transcript release** | **Of those, scored by its isoform release** | **Unscored by CADD** |
| --- | --- | --- | --- | --- | --- |
| CALM1 | ENST00000356978.9 | 1,013 | 1,013 (100.0%) | 1,013 | 0 (0.0%) |
| CALM2 | ENST00000272298.12 | 1,017 | 1,017 (100.0%) | 1,017 | 0 (0.0%) |
| WT1 | ENST00000452863.10 | 3,412 | 3,412 (100.0%) | 3,412 | 0 (0.0%) |
| TTN | ENST00000589042.5 | 235,209 | 10,572 (4.5%) | 870 | 0 (0.0%) |
| All 84 genes | – | 767,747 | 16,014 (2.1%) | 6,312 | 0 (0.0%) |

The other genes of ACMG SF v3.3 have no unscored possible missense substitution under either score. Substitutions were enumerated from the MANE Select coding sequence (MANE v1.5). The 9,702 substitutions unscored by both AlphaMissense releases are in TTN. CADD, Combined Annotation Dependent Depletion; MANE, Matched Annotation from NCBI and EMBL-EBI; ACMG SF v3.3, the American College of Medical Genetics and Genomics list of genes for reporting secondary findings, version 3.3.

Table S32. Variant classes and genes that widely used scores leave unscored

| **Class** | **Panel and source of the scores** | **Variants or genes** | **Scores** | **What is scored or unscored** |
| --- | --- | --- | --- | --- |
| Pathogenic initiator-codon variants | dbNSFP panel, dbNSFP 5.3.1a | 1,608 in 749 genes; 1,558 on the MANE Select start codon | MutationAssessor, MisFit_D, MisFit_S, MPC, PrimateAI, AlphaMissense, MutScore | 0.016 to 0.053 scored; the other 25 missense-scope predictors score at least 0.820 |
| Benign initiator-codon variants | dbNSFP panel, dbNSFP 5.3.1a | 202 in 183 genes; 70 on the MANE Select start codon | the same seven | 0.436 to 0.649 scored |
| Mitochondrial genome | ClinVar panel, each tool's own release | 1,717 in 37 genes; 111 pathogenic | CADD, REVEL, AlphaMissense | none scored; phyloP and phastCons score all 1,717 |
| Unlocalised or alternate scaffolds | ClinVar panel, each tool's own release | 12; 0 pathogenic | CADD, phyloP, phastCons, REVEL, AlphaMissense | none scored |
| Pathogenic missense variants outside any MANE coding sequence | dbNSFP panel, dbNSFP 5.3.1a | 91 in 23 genes | mean of the 32 missense-scope predictors | 0.6161 unscored (95% CI 0.5082 to 0.6643), against 0.0367 in internal coding exons |
| Benign missense variants outside any MANE coding sequence | dbNSFP panel, dbNSFP 5.3.1a | 3,117 in 1,059 genes | mean of the 32 missense-scope predictors | 0.4766 unscored (0.4518 to 0.5028), against 0.0717 in internal coding exons |
| Whole genes, AlphaMissense's own release | ClinVar panel, genes with at least 10 ClinVar missense variants | 97 of 4,298 genes | AlphaMissense | none of the gene's ClinVar missense variants scored; the 97 hold 2,877 of its 11,196 unscored missense variants and 645 of its 1,019 unscored pathogenic ones |
| Whole genes, REVEL's own release | ClinVar panel, genes with at least 10 ClinVar missense variants | 27 of 4,298 genes | REVEL | none scored; the 27 hold 631 of its 3,767 unscored missense variants and 130 of its 252 unscored pathogenic ones |
| Whole genes, dbNSFP's REVEL column | dbNSFP panel, genes with at least 10 missense variants | 137 of 4,254 genes | REVEL | none scored in dbNSFP; REVEL's own release scores more than 0.9 of the gene in 95 and none in 30 |
| Whole genes, dbNSFP's AlphaMissense column | dbNSFP panel, genes with at least 10 missense variants | 12 of 4,254 genes | AlphaMissense | none scored in dbNSFP; AlphaMissense's own release scores more than 0.9 of the gene in 5 and none in 6 |

Reach on pathogenic and on benign initiator-codon variants is 0.016 and 0.436 for MutationAssessor, 0.017 and 0.480 for MisFit_D and for MisFit_S, 0.022 and 0.446 for MPC, 0.026 and 0.480 for PrimateAI, 0.053 and 0.495 for AlphaMissense and 0.052 and 0.649 for MutScore. Position is read on MANE v1.5 by coordinate; which transcript each predictor read is not in the public tables, so position is a property of the variant. Intervals resample genes. The largest genes wholly unscored by AlphaMissense's own release, by pathogenic count, are ARSA (179 pathogenic of 190 missense variants), PRPH2 (114 of 119), CACNA1C (51 of 134) and WT1 (46 of 67). CI, confidence interval; dbNSFP, database for nonsynonymous single-nucleotide polymorphisms' functional predictions; MANE, Matched Annotation from NCBI and EMBL-EBI.

Table S33. Flow of records in the systematic search for published benchmarks

| **Step** | **Records** | **Reasons (records)** |
| --- | --- | --- |
| Returned by the PubMed query and screened on title and abstract | 75 |  |
| Excluded on title and abstract | 31 | not a predictor comparison (18); no clinical labels (5); single predictor (4); no ROC analysis (1); review or commentary without own evaluation (1); retracted, erratum or duplicate record (1); not variant-level prediction (1) |
| Full texts sought | 44 |  |
| Not retrievable from an open source | 13 | no open full text (11); open copy only at a publisher's site that refused automated requests (2) |
| Full texts assessed | 31 |  |
| Excluded on full text | 14 | no AUROC printed (8); AUROC shown only as graphics (6) |
| Included: per-predictor AUROCs printed on a clinically labelled panel | 17 |  |
| Assessable: an AUROC ordering stated, and scored counts or coverage reported | 10 |  |
| Included but not assessable | 7 | scored counts not reported (3); compared on variants all predictors score (2); scored counts not reported for the predictors ranked first (1); no AUROC ordering stated (1) |

PubMed was searched once, on 11 November 2026, with this query, fixed before it was run: (benchmark*[ti] OR compar*[ti] OR evaluat*[ti] OR assess*[ti] OR performance[ti]) AND (variant*[ti] OR mutation*[ti] OR indel*[ti] OR substitution*[ti]) AND (predict*[ti] OR "in silico"[ti] OR computational[ti] OR tool*[ti] OR method*[ti] OR algorithm*[ti] OR score*[ti] OR classifier*[ti]) AND (pathogenic*[tiab] OR deleterious*[tiab] OR damaging[tiab] OR "variant effect"[tiab] OR "variant effects"[tiab] OR "disease-causing"[tiab] OR "disease-associated"[tiab]) AND (ClinVar[tiab] OR HGMD[tiab] OR ClinGen[tiab] OR ACMG[tiab] OR "clinical variant*"[tiab] OR "clinically relevant"[tiab] OR "clinical significance"[tiab] OR "clinical classification*"[tiab] OR "clinical interpretation"[tiab] OR "clinically classified"[tiab] OR "known pathogenic"[tiab] OR "pathogenic and benign"[tiab] OR "benign and pathogenic"[tiab] OR "pathogenic or benign"[tiab] OR "benign or pathogenic"[tiab]) AND 2015:2026[dp] AND English[la] NOT preprint[pt]. PubMed reported the last three quoted phrases as absent from its phrase index and ignored them. The query is anchored on the title, so an article that introduces a predictor under a title naming only that predictor lies outside it. Full text was read only from open sources. Checks: a random 20 of the 31 exclusions on title and abstract and all 14 exclusions on full text were screened a second time without disagreement; the three legends of Barbosa et al. [72] (104 entries) and the table of Heo and Kim [73] (28 rows) were extracted a second time, independently; the two extractions differ by one legend entry, which the counts include; of a random 70 of the 467 rows first extracted, 69 were re-read without difference and 1, in an article that is not assessable, agrees within the resolution of its image. Every record, with its decision and reason, is in Additional file 3. AUROC, area under the receiver operating characteristic curve; ROC, receiver operating characteristic.

Table S34. The ten assessable benchmarks of the systematic search: stated orderings by verdict

| **Benchmark** | **Stated** | **Identified** | **Not identified** | **Share** | **Split-dependent** | **Not assessable** | **How stated** | **Coverage reported** |
| --- | --- | --- | --- | --- | --- | --- | --- | --- |
| Barbosa et al. [72] | 1,692 | 961 | 639 | 37.8% | 92 | 0 | three figure legends ranked by AUROC | scored total per tool; class split not printed |
| Heo and Kim [73] | 378 | 106 | 272 | 72.0% | 0 | 0 | one figure legend in descending AUROC; the text names the two best | scored counts per class |
| Li et al. [133] | 330 | 307 | 23 | 7.0% | 0 | 0 | text | missing count with confusion metrics |
| Gu et al. [134] | 187 | 173 | 14 | 7.5% | 0 | 0 | text, tools listed in descending AUROC | counts per class, from confusion counts |
| Wang et al. [132] | 66 | 48 | 18 | 27.3% | 0 | 0 | text and bold type in a table | missing rate with confusion metrics |
| Liu et al. [135] | 63 | 52 | 11 | 17.5% | 0 | 0 | text | missing fraction with confusion metrics |
| Montanucci et al. [136] | 63 | 0 | 60 | 95.2% | 0 | 3 | table sorted by AUROC, and text | scored share only |
| Pshennikova et al. [137] | 23 | 23 | 0 | 0.0% | 0 | 0 | text | every variant scored by every tool |
| Michels et al. [138] | 21 | 16 | 5 | 23.8% | 0 | 0 | table in descending AUROC | counts per class for the one tool scored on fewer |
| Montenegro et al. [139] | 19 | 16 | 3 | 15.8% | 0 | 0 | text | counts of calls per class |
| All ten | 2,842 | 1,702 | 1,045 | 36.8% | 92 | 3 |  |  |

An ordering is stated when the article prints a rank by AUROC, sorts a table or legend by it, marks the best AUROC typographically or says in words that one predictor has the higher AUROC. Share, not identified over stated. Readings that change a row: Heo and Kim's text alone states 52 orderings, all identified. Montenegro et al.'s tables disagree with one another; with counts that reproduce the confusion metrics the article prints, 1 or 0 of its orderings is unidentified. All 14 unidentified orderings of Gu et al. involve M-CAP, entered as scoring 54 of 57 controls from the article's table; if it scored all 57, as the article's text implies, none is unidentified. With these three read strictly, 756 of 2,516 stated orderings are not identified (30.0%), and 6 of the 10 benchmarks state at least one. For Montanucci et al. pair coverage is taken as the printed share squared, an approximation; bracketed over every class split of the implied unscored count, all 60 stay unidentified. Liu et al.'s 11 lie on their somatic and expression quantitative trait locus panels, which are not clinically labelled; on their ClinVar panel all 7 stated orderings are identified. Some benchmarks of both sets, Barbosa et al. and CAGI6 [7] among them, also report coverage-aware analyses, which the check does not address. The seven included benchmarks that are not assessable do not report scored counts for all the predictors they order [198–201], compare on variants every predictor scores [147,202] or state no ordering by AUROC [140] (Table S33).

Column headings: share, share not identified; split-dependent, depends on the class split; how stated, how the ordering is stated; AUROC, area under the receiver operating characteristic curve.

Table S35. The ten orderings that the selection rule of Table 2 gives without its cap of two rows per benchmark

| **#** | **Benchmark (how the ordering is printed)** | **P / B** | **Printed higher: AUROC; variants; bound** | **Printed lower: AUROC; variants; bound** | **λ*** | **Articles** | **Not identified** |
| --- | --- | --- | --- | --- | --- | --- | --- |
| 1 | Heo and Kim [73] (sorted legend) | as Table 2, row 1 |  |  |  |  |  |
| 2 | Heo and Kim [73] (sorted legend) | as Table 2, row 2 |  |  |  |  |  |
| 3 | Heo and Kim [73] (sorted legend) | 1,951 / 2,638 | REVEL: 0.9707; scored 1,784 of 1,951 P, 2,362 of 2,638 B; 0.795 to 0.976 | PolyPhen-2 (HumVar): 0.8961; scored 1,652 of 1,951 P, 2,181 of 2,638 B; 0.627 to 0.927 | 0.360 | 10 | 4 of 16 |
| 4 | Heo and Kim [73] (sorted legend) | 1,951 / 2,638 | REVEL: 0.9707; scored 1,784 of 1,951 P, 2,362 of 2,638 B; 0.795 to 0.976 | CADD: 0.8971; scored 1,951 of 1,951 P, 2,638 of 2,638 B; 0.897 to 0.897 | 0.418 | 10 | 3 of 9 |
| 5 | Barbosa et al. [72] (ranked legend) | as Table 2, row 3 |  |  |  |  |  |
| 6 | Heo and Kim [73] (sorted legend) | 1,951 / 2,638 | REVEL: 0.9707; scored 1,784 of 1,951 P, 2,362 of 2,638 B; 0.795 to 0.976 | SIFT4G: 0.8850; scored 1,856 of 1,951 P, 2,367 of 2,638 B; 0.755 to 0.902 | 0.445 | 9 | 2 of 8 |
| 7 | Barbosa et al. [72] (ranked legend) | as Table 2, row 4 |  |  |  |  |  |
| 8 | Barbosa et al. [72] (ranked legend) | 278 / 193 | CADD: 0.81; 1 of 471 unscored; 0.807 to 0.811 | MutPred: 0.8; 192 of 471 unscored; 0.290 to 0.927 | 0.049 to 0.077 | 7 | 2 of 2 |
| 9 | Heo and Kim [73] (sorted legend) | 1,951 / 2,638 | MutPred: 0.9508; scored 1,595 of 1,951 P, 856 of 2,638 B; 0.252 to 0.987 | SIFT4G: 0.8850; scored 1,856 of 1,951 P, 2,367 of 2,638 B; 0.755 to 0.902 | 0.092 | 7 | 3 of 5 |
| 10 | Barbosa et al. [72] (ranked legend) | 93 / 100 | ClinPred: 0.99; 2 of 193 unscored; 0.970 to 0.990 | REVEL: 0.97; 2 of 193 unscored; 0.951 to 0.971 | 0.912 to 0.980 | 7 | 1 of 5 |

Columns, abbreviations and the selection rule as in Table 2, without the cap, among the stated, unidentified orderings of the thirteen predictor families; candidates are the 75 such orderings whose coverage is printed as counts (28 pairs of families, 4 articles). Ranked legend, a legend whose caption says the tools are ranked by AUROC; sorted legend, a legend in descending AUROC that the caption does not call a ranking, so that under the stricter reading of Note S24 the six rows of Heo and Kim are not stated orderings. Rows 5, 7, 8 and 10 are differences of 0.01 or 0.02 between AUROCs printed to two decimals on panels of 193 and 471 variants.

Column headings: P / B, the panel's pathogenic and benign variants; articles, articles printing the pair; not identified, of the pair's stated orderings.

Table S36. The GPN-Star benchmark suite, dataset by dataset: model pairs resolved on the complete case and identified once unscored variants are charged

| **Dataset (species)** | **Metric the suite prints** | **Score files (unscored)** | **Pairs** | **Resolved (Holm)** | **Touching unscored** | **Of these, resolved** | **Of these, identified** |
| --- | --- | --- | --- | --- | --- | --- | --- |
| clinvar_vs_benign (human) | average precision | 19 (6) | 171 | 167 (165) | 93 | 93 | 77 |
| cosmic (human) | average precision | 17 (8) | 136 | 124 (113) | 100 | 88 | 78 |
| omim_traitgym (human) | average precision | 21 (7) | 210 | 181 (164) | 119 | 103 | 101 |
| ukb_finemapped_coding (human) | average precision | 18 (2) | 153 | 100 (68) | 33 | 27 | 24 |
| ukb_finemapped_nc_traitgym (human) | average precision | 21 (7) | 210 | 166 (144) | 119 | 97 | 93 |
| gnomad_balanced (human) | odds ratio | 12 (8) | 66 | 59 (58) | 60 | 60 | 47 |
| mmrdb (mouse) | average precision | 3 (2) | 3 | 3 (3) | 3 | 3 | 2 |
| wmgp_balanced (mouse) | odds ratio | 3 (3) | 3 | 3 (3) | 3 | 3 | 0 |
| flybase_lethal (fly) | average precision | 3 (2) | 3 | 3 (3) | 3 | 3 | 3 |
| dest (fly) | odds ratio | 3 (2) | 3 | 3 (3) | 3 | 3 | 3 |
| celegans_lethal (worm) | average precision | 3 (2) | 3 | 3 (3) | 3 | 3 | 2 |
| caendr (worm) | odds ratio | 3 (2) | 3 | 3 (3) | 3 | 3 | 2 |
| galbase (chicken) | odds ratio | 3 (2) | 3 | 3 (3) | 3 | 3 | 3 |
| 1001gp (Arabidopsis) | odds ratio | 3 (2) | 3 | 2 (2) | 3 | 3 | 0 |
| All 14 |  | 132 (55) | 970 | 820 (735) | 548 | 492 | 435 |

Datasets under the names of the release that accompanies GPN-Star [30]; no model was rescored. Complete case, the variants every released file of a dataset scores, the rule the article states for its own comparisons. A pair is resolved when its difference clears zero by 1.96 standard errors over 2,000 paired resamples: variants resampled within label class for average precision, as in the article; whole chromosomes, or 1-Mb blocks where an assembly has fewer than ten chromosomes, for the odds ratio of rare against common variants, for which the article gives no interval. A pair is identified when the two files' bounds on whole-panel AUROC are disjoint, with 95% confidence when the Imbens–Manski set of the gap excludes zero. Over all 970 pairs, 831 are resolved in AUROC on the complete case and 780 are identified at 95% once charged (723 and 675 under Holm's procedure). The bound is for AUROC, whereas the suite prints average precision and odds ratios; in 18 pairs the printed metric and AUROC give opposite orders that are both resolved. In the mouse and Arabidopsis population sets (wmgp_balanced, 1001gp) the PhyloP and PhastCons files leave 19.5% and 14.0% of variants unscored. AUROC, area under the receiver operating characteristic curve; GPN-Star, Genomic Pretrained Network with Species Tree and Alignment Representations.

Column headings: score files (unscored), score files (those with unscored variants); resolved (Holm), resolved at 95% by the printed metric (under Holm's procedure); touching unscored, pairs touching a file with unscored variants; of these, resolved, resolved in AUROC on the complete case; of these, identified, identified at 95% once charged.

Table S37. SimpleQA: bounds on the accuracy of eight language models had each answered every question

| **Model** | **Correct (%)** | **Not attempted (%)** | **Incorrect (%)** | **Bound on accuracy had every question been answered (%)** | **Accuracy on attempted questions (%)** | **95% confidence set for the bounded accuracy (%)** |
| --- | --- | --- | --- | --- | --- | --- |
| Claude-3-haiku | 5.1 | 75.3 | 19.6 | 5.1 to 80.4 | 20.6 | 4.5 to 81.4 |
| Claude-3-sonnet | 5.7 | 75.0 | 19.3 | 5.7 to 80.7 | 22.8 | 5.1 to 81.7 |
| Claude-3-opus | 23.5 | 39.6 | 36.9 | 23.5 to 63.1 | 38.9 | 22.4 to 64.3 |
| Claude-3.5-sonnet | 28.9 | 35.0 | 36.1 | 28.9 to 63.9 | 44.5 | 27.8 to 65.1 |
| o1-mini | 8.1 | 28.5 | 63.4 | 8.1 to 36.6 | 11.3 | 7.4 to 37.8 |
| o1-preview | 42.7 | 9.2 | 48.1 | 42.7 to 51.9 | 47.0 | 41.5 to 53.1 |
| GPT-4o | 38.2 | 1.0 | 60.8 | 38.2 to 39.2 | 38.6 | 37.0 to 40.4 |
| GPT-4o-mini | 8.6 | 0.9 | 90.5 | 8.6 to 9.5 | 8.7 | 7.9 to 10.2 |

Shares of the 4,326 questions, from the third table of the SimpleQA article [48], models in the order of the share not attempted. The bound runs from the share correct (every declined question counted wrong) to the share correct plus the share not attempted. Accuracy on attempted questions is the share correct over the shares correct and incorrect, recomputed from the three shares; for GPT-4o the article prints 38.0 (Note S42), and the 38.6 here reproduces its printed F-score of 38.4. Rounding of the printed percentages changes no count below. Confidence sets are of the Imbens–Manski type on binomial standard errors. The point bounds identify seven comparisons (six at 95% confidence, Note S42): o1-preview above GPT-4o, o1-mini and GPT-4o-mini; GPT-4o above o1-mini and GPT-4o-mini; Claude-3.5-sonnet and Claude-3-opus above GPT-4o-mini. Seven can never be identified, the two widths summing to at least one: Claude-3-haiku and Claude-3-sonnet with each other and each with Claude-3-opus, Claude-3.5-sonnet and o1-mini.

Table S38. Evo 2 against the 241-mammal phyloP, pCADD and chCADD: reach, the charge for silence and the margins

| **Species** | **Comparator** | **Positives scored** | **Negatives scored** | **Penalty** | **Margin, 1,001 bp** | **Margin, 8,192 bp** | **Verdict, 8,192 bp (replicates)** |
| --- | --- | --- | --- | --- | --- | --- | --- |
| Goat | phyloP-241 | 8 of 9 (88.9%) | 56 of 90 (62.2%) | 0.115 | +0.212 | +0.225 | identified, Evo 2 ahead (not estimable) |
| Chicken | phyloP-241 | 26 of 28 (92.9%) | 29 of 280 (10.4%) | 0.194 | +0.125 [−0.035, +0.301] | +0.210 [+0.067, +0.377] | not identified (16.8% / 0.0%) |
| Pig | phyloP-241 | 36 of 36 (100.0%) | 158 of 360 (43.9%) | 0.227 | −0.045 [−0.173, +0.062] | −0.036 [−0.163, +0.069] | not identified (1.8% / 0.0%) |
| Sheep | phyloP-241 | 55 of 56 (98.2%) | 284 of 560 (50.7%) | 0.230 | −0.053 [−0.141, +0.025] | +0.005 [−0.042, +0.060] | not identified (22.9% / 0.0%) |
| Horse | phyloP-241 | 71 of 71 (100.0%) | 428 of 710 (60.3%) | 0.178 | −0.056 [−0.133, +0.079] | −0.005 [−0.048, +0.080] | not identified (6.7% / 0.0%) |
| Cat | phyloP-241 | 122 of 125 (97.6%) | 798 of 1,240 (64.4%) | 0.157 | −0.052 [−0.097, −0.005] | −0.012 [−0.056, +0.035] | not identified (1.0% / 0.0%) |
| Cattle | phyloP-241 | 187 of 188 (99.5%) | 912 of 1,880 (48.5%) | 0.225 | −0.016 [−0.053, +0.022] | +0.036 [+0.008, +0.064] | identified, Evo 2 ahead (71.6% / 0.0%) |
| Dog | phyloP-241 | 227 of 227 (100.0%) | 1,275 of 2,270 (56.2%) | 0.201 | −0.063 [−0.095, −0.031] | +0.014 [−0.004, +0.031] | not identified (16.0% / 0.0%) |
| Human | phyloP-241 | 1,500 of 1,500 (100.0%) | 1,500 of 1,500 (100.0%) | 0.000 | −0.089 [−0.109, −0.069] | +0.060 [+0.049, +0.072] | identified, Evo 2 ahead (100.0% / 0.0%) |
| Species mean (9) | phyloP-241 | 97.4% | 55.2% | 0.170 | −0.004 (−0.083 to +0.074) | +0.055 (−0.019 to +0.129) |  |
| Pig | pCADD | 36 of 36 (100.0%) | 360 of 360 (100.0%) | 0.000 | +0.006 [−0.098, +0.098] | +0.021 [−0.083, +0.128] | point identified, Evo 2 ahead (65.8% / 34.2%) |
| Chicken | chCADD | 28 of 28 (100.0%) | 280 of 280 (100.0%) | 0.000 | −0.120 [−0.216, −0.044] | −0.028 [−0.081, +0.014] | point identified, chCADD ahead (10.7% / 89.2%) |

Reach is over every atlas variant of the species, and the must-answer penalty is the covered minus the must-answer AUROC (GERP's is in Table S1; its species mean is 0.102). phyloP-241, the Zoonomia 241-mammal phyloP, is read on GRCh38, non-human variants at the position their base lifts to through the UCSC chain of their own assembly; 55 of chicken's 308 variants reach a human position. Margins are on the variants both Evo 2 readouts and the comparator score (7,669 for phyloP-241, all 396 pig and all 308 chicken variants for pCADD and chCADD); a positive margin favours Evo 2. Square brackets hold 95% locus-bootstrap intervals (positive-bearing 100-kb loci drawn whole, B = 2,000); goat, whose positives occupy two loci, carries none. The mean row's round brackets hold a t interval over species on k − 1 degrees of freedom; without goat the margins are −0.031 (−0.086 to +0.024) and +0.034 (−0.030 to +0.098), and without goat and chicken −0.053 (−0.074 to −0.033) and +0.009 (−0.021 to +0.038). Verdicts place every unscored variant at both extremes; replicates draw loci whole (B = 20,000), and a species whose positives occupy fewer than three loci is not estimable. For a score that answers every variant the bound is a point, so its verdict is the sign of a point difference and the replicate shares carry the evidence. Full tables, including the within-class rows and the margins on the variants GERP also scores: Additional file 3, scripts/recompute_atlas_comparators.py.

Column headings: penalty, must-answer penalty; margin, Evo 2 minus comparator; verdict, 8,192 bp (replicates), verdict at 8,192 bp, with the shares of replicates identified for Evo 2 and for the comparator. AUROC, area under the receiver operating characteristic curve; GERP, Genomic Evolutionary Rate Profiling; UCSC, University of California, Santa Cruz; pCADD and chCADD, Combined Annotation Dependent Depletion (CADD) built for pig and for chicken.

Table S39. Conservation scores on the consequence-matched arm, by arm and consequence class

| **Score** | **Variants** | **Atlas, original negatives** | **Same-class arm** | **Synonymous arm** |
| --- | --- | --- | --- | --- |
| phyloP-241 | species mean (9) | 0.890 | 0.740 (0.631 to 0.850) | 0.892 (0.832 to 0.953) |
| GERP | species mean (9) | 0.830 | 0.658 (0.531 to 0.785) | 0.842 (0.788 to 0.896) |
| phyloP-241 | eight OMIA species, missense, pooled | 0.787 [0.716, 0.851] | 0.811 [0.787, 0.835] | 0.952 [0.939, 0.964] |
| GERP | eight OMIA species, missense, pooled | 0.660 [0.573, 0.744] | 0.712 [0.680, 0.744] | 0.868 [0.846, 0.887] |
| phyloP-241 | eight OMIA species, stop gained, pooled | – | 0.656 [0.589, 0.718] | 0.879 [0.838, 0.918] |
| GERP | eight OMIA species, stop gained, pooled | – | 0.626 [0.559, 0.690] | 0.808 [0.760, 0.853] |
| phyloP-241 | human, missense | 0.867 | 0.841 [0.815, 0.865] | 0.960 [0.947, 0.972] |
| GERP | human, missense | 0.843 | 0.805 [0.777, 0.831] | 0.924 [0.906, 0.940] |
| GPN-Star | chicken | 0.931 [0.871, 0.983] | 0.715 [0.597, 0.826] | 0.904 [0.813, 0.973] |
| GPN-Star | human | 0.969 [0.961, 0.976] | 0.840 [0.823, 0.858] | 0.974 [0.964, 0.982] |
| pCADD | pig | 0.842 [0.724, 0.943] | 0.838 [0.723, 0.926] | 0.933 [0.828, 1.000] |
| chCADD | chicken | 0.981 [0.960, 0.997] | 0.549 [0.423, 0.692] | 1.000 [1.000, 1.000] |

Covered AUROC; Evo 2 is not scored on the arm. The panel holds 2,237 positives, 5,002 same-class negatives and 4,117 synonymous negatives in nine species, matched on consequence class (first snpEff term), reference trinucleotide and alternate base. Same-class arm: each positive against population variants of its own class, pairs ranked within species and class; synonymous arm: each protein-changing positive (missense, stop gained, start lost, stop lost) against synonymous population variants, every pair ranked. Pooled rows pool pairs within species and class by pair count, as Table S21 does. Atlas column: the same score against the atlas's original negatives, every positive included (Note S44's drops compare each arm with the atlas on that arm's own positives, so they differ slightly from this column's differences); for missense, the atlas's own within-class missense rows, which rest on 74 (phyloP-241) and 72 (GERP) OMIA negatives; stop gained has no such row. Round brackets: t intervals over species on k − 1 degrees of freedom; square brackets: 95% locus-bootstrap intervals (positive-bearing 100-kb loci drawn whole, B = 2,000). Charged for silence, the species means are 0.727 (0.628 to 0.826) and 0.875 (0.802 to 0.948) for phyloP-241 and 0.621 (0.523 to 0.720) and 0.770 (0.709 to 0.831) for GERP on the same-class and synonymous arms, against 0.720 and 0.728 on the atlas. Over every pair of the same-class arm the snpEff impact ordering reads 0.506 (0.495 to 0.517) by the species mean, against 0.958 on the atlas. Per-species rows, the other classes and the matching rules: Additional file 3, scripts/recompute_atlas_matched.py. AUROC, area under the receiver operating characteristic curve; GERP, Genomic Evolutionary Rate Profiling; OMIA, Online Mendelian Inheritance in Animals; GPN-Star, Genomic Pretrained Network with Species Tree and Alignment Representations; pCADD and chCADD, as in Table S38.

### References

References 1 to 122 keep the main manuscript's numbers, and only those cited in this file are listed; numbers above 122 are cited in this file only and continue that list.

1. Kumar P, Henikoff S, Ng PC. Predicting the effects of coding non-synonymous variants on protein function using the SIFT algorithm. Nat Protoc. 2009;4(7):1073-1081. doi:10.1038/nprot.2009.86.

2. Adzhubei IA, Schmidt S, Peshkin L, et al. A method and server for predicting damaging missense mutations. Nat Methods. 2010;7(4):248-249. doi:10.1038/nmeth0410-248.

4. Radjasandirane R, Diharce J, Gelly JC, et al. Assessment of variant effect predictors unveils variants difficulty as a critical performance indicator. bioRxiv. 2024. doi:10.1101/2024.07.08.602580 (version 2; published as Insights for variant clinical interpretation based on a benchmark of 65 variant effect predictors, Genomics 2025;117(3):111036, doi:10.1016/j.ygeno.2025.111036).

5. Livesey BJ, Marsh JA. Variant effect predictor correlation with functional assays is reflective of clinical classification performance. Genome Biol. 2025;26(1):104. doi:10.1186/s13059-025-03575-w.

6. Notin P, Kollasch AW, Ritter D, et al. ProteinGym: large-scale benchmarks for protein fitness prediction and design. Adv Neural Inf Process Syst. 2023;36:64331-64379. doi:10.52202/075280-2810.

7. Rastogi R, Chung R, Li S, et al. Critical assessment of missense variant effect predictors on disease-relevant variant data. Hum Genet. 2025;144(2-3):281-293. doi:10.1007/s00439-025-02732-2.

9. Brandes N, Goldman G, Wang CH, et al. Genome-wide prediction of disease variant effects with a deep protein language model. Nat Genet. 2023;55(9):1512-1522. doi:10.1038/s41588-023-01465-0.

17. Harrizi S, Nait Irahal I, Mostafa K, et al. Gene identity, not variant effect, dominates ClinVar benchmarks of missense pathogenicity predictors. bioRxiv. 2026. doi:10.64898/2026.08.21.746180.

18. Pejaver V, Byrne AB, Feng BJ, et al. Calibration of computational tools for missense variant pathogenicity classification and ClinGen recommendations for PP3/BP4 criteria. Am J Hum Genet. 2022;109(12):2163-2177. doi:10.1016/j.ajhg.2022.10.013.

19. Bergquist T, Stenton SL, Nadeau EAW, et al. Calibration of additional computational tools expands ClinGen recommendation options for variant classification with PP3/BP4 criteria. Genet Med. 2025;27(6):101402. doi:10.1016/j.gim.2025.101402.

20. Frazer J, Notin P, Dias M, et al. Disease variant prediction with deep generative models of evolutionary data. Nature. 2021;599(7883):91-95. doi:10.1038/s41586-021-04043-8.

21. Choe YJ, Gangrade A, Ramdas A. Counterfactually comparing abstaining classifiers. Adv Neural Inf Process Syst. 2023;36:28281-28293. doi:10.52202/075280-1229.

23. Fernández-Barrios R, Pastor-López I, González-Santocildes A, et al. When disagreement labels do not identify F1 rankings. arXiv. 2026. doi:10.48550/arXiv.2611.01712.

24. Manski CF. Partial identification of probability distributions. New York: Springer; 2003. doi:10.1007/b97478.

25. Horowitz JL, Manski CF. Identification and robustness with contaminated and corrupted data. Econometrica. 1995;63(2):281-302. doi:10.2307/2951627.

26. Li Y, Engelhard MM. Extending ROC analysis to uncertainty-aware risk prediction with an interval-based AUC (iAUC). arXiv. 2026. doi:10.48550/arXiv.2602.04775.

30. Ye C, Benegas G, Albors C, et al. Predicting genome-wide functional constraints with GPN-Star. Nature. 2026;659(8139):141-150. doi:10.1038/s41586-026-11005-5.

31. Brixi G, Durrant MG, Ku J, et al. Genome modelling and design across all domains of life with Evo 2. Nature. 2026;652(8112):1349-1361. doi:10.1038/s41586-026-10176-5.

34. Jiang J, Tiezzi F, Maltecca C. Cross-species functional variant classification using the Evo 2 genomic language model: implications for farm animal genetics and breeding. Research Square. 2025. doi:10.21203/rs.3.rs-7579108/v1.

37. Pollard KS, Hubisz MJ, Rosenbloom KR, et al. Detection of nonneutral substitution rates on mammalian phylogenies. Genome Res. 2010;20(1):110-121. doi:10.1101/gr.097857.109.

38. Siepel A, Bejerano G, Pedersen JS, et al. Evolutionarily conserved elements in vertebrate, insect, worm, and yeast genomes. Genome Res. 2005;15(8):1034-1050. doi:10.1101/gr.3715005.

39. Schubach M, Maass T, Nazaretyan L, et al. CADD v1.7: using protein language models, regulatory CNNs and other nucleotide-level scores to improve genome-wide variant predictions. Nucleic Acids Res. 2024;52(D1):D1143-D1154. doi:10.1093/nar/gkad989.

40. Ioannidis NM, Rothstein JH, Pejaver V, et al. REVEL: an ensemble method for predicting the pathogenicity of rare missense variants. Am J Hum Genet. 2016;99(4):877-885. doi:10.1016/j.ajhg.2016.08.016.

41. Cheng J, Novati G, Pan J, et al. Accurate proteome-wide missense variant effect prediction with AlphaMissense. Science. 2023;381(6664):eadg7492. doi:10.1126/science.adg7492.

42. Cooper GM, Stone EA, Asimenos G, et al. Distribution and intensity of constraint in mammalian genomic sequence. Genome Res. 2005;15(7):901-913. doi:10.1101/gr.3577405.

43. Davydov EV, Goode DL, Sirota M, et al. Identifying a high fraction of the human genome to be under selective constraint using GERP++. PLoS Comput Biol. 2010;6(12):e1001025. doi:10.1371/journal.pcbi.1001025.

44. Zoonomia Consortium. A comparative genomics multitool for scientific discovery and conservation. Nature. 2020;587(7833):240-245. doi:10.1038/s41586-020-2876-6.

45. Christmas MJ, Kaplow IM, Genereux DP, et al. Evolutionary constraint and innovation across hundreds of placental mammals. Science. 2023;380(6643):eabn3943. doi:10.1126/science.abn3943.

46. Preston CG, Wright MW, Madhavrao R, et al. ClinGen Variant Curation Interface: a variant classification platform for the application of evidence criteria from ACMG/AMP guidelines. Genome Med. 2022;14(1):6. doi:10.1186/s13073-021-01004-8.

47. Benegas G, Eraslan G, Song YS. Benchmarking DNA sequence models for causal regulatory variant prediction in human genetics. bioRxiv. 2025. doi:10.1101/2025.02.11.637758.

48. Wei J, Nguyen K, Chung HW, et al. Measuring short-form factuality in large language models. arXiv. 2024. doi:10.48550/arXiv.2411.04368.

50. Imbens GW, Manski CF. Confidence intervals for partially identified parameters. Econometrica. 2004;72(6):1845-1857. doi:10.1111/j.1468-0262.2004.00555.x.

51. Stoye J. More on confidence intervals for partially identified parameters. Econometrica. 2009;77(4):1299-1315. doi:10.3982/ECTA7347.

54. Masten MA, Poirier A. Inference on breakdown frontiers. Quant Econ. 2020;11(1):41-111. doi:10.3982/QE1288.

55. Richards S, Aziz N, Bale S, et al. Standards and guidelines for the interpretation of sequence variants: a joint consensus recommendation of the American College of Medical Genetics and Genomics and the Association for Molecular Pathology. Genet Med. 2015;17(5):405-424. doi:10.1038/gim.2015.30.

56. Wilcox EH, Sarmady M, Wulf B, et al. Evaluating the impact of in silico predictors on clinical variant classification. Genet Med. 2022;24(4):924-930. doi:10.1016/j.gim.2021.11.018.

57. Lee K, Abul-Husn NS, Amendola LM, et al. ACMG SF v3.3 list for reporting of secondary findings in clinical exome and genome sequencing: a policy statement of the American College of Medical Genetics and Genomics (ACMG). Genet Med. 2025;27(8):101454. doi:10.1016/j.gim.2025.101454.

58. Smith C, Kitzman JO. Benchmarking splice variant prediction algorithms using massively parallel splicing assays. Genome Biol. 2023;24(1):294. doi:10.1186/s13059-023-03144-z.

63. Teng J, Gao Y, Yin H, et al. A compendium of genetic regulatory effects across pig tissues. Nat Genet. 2024;56(1):112-123. doi:10.1038/s41588-023-01585-7.

64. Kumar S, Suleski M, Craig JM, et al. TimeTree 5: an expanded resource for species divergence times. Mol Biol Evol. 2022;39(8):msac174. doi:10.1093/molbev/msac174.

66. Groß C, Derks M, Megens HJ, et al. pCADD: SNV prioritisation in *Sus scrofa*. Genet Sel Evol. 2020;52(1):4. doi:10.1186/s12711-020-0528-9.

67. Groß C, Bortoluzzi C, de Ridder D, et al. Prioritizing sequence variants in conserved non-coding elements in the chicken genome using chCADD. PLoS Genet. 2020;16(9):e1009027. doi:10.1371/journal.pgen.1009027.

69. Hu H, Xie W. Human genomic tools to assess missense variant pathogenicity in domestic mammals using a mock-variant strategy. Open Vet J. 2025;15(10):5183-5191. doi:10.5455/OVJ.2025.v15.i10.35.

70. McLaren W, Gil L, Hunt SE, et al. The Ensembl Variant Effect Predictor. Genome Biol. 2016;17(1):122. doi:10.1186/s13059-016-0974-4.

71. Findlay GM, Daza RM, Martin B, et al. Accurate classification of BRCA1 variants with saturation genome editing. Nature. 2018;562(7726):217-222. doi:10.1038/s41586-018-0461-z.

72. Barbosa P, Ribeiro M, Carmo-Fonseca M, et al. Clinical significance of genetic variation in hypertrophic cardiomyopathy: comparison of computational tools to prioritize missense variants. Front Cardiovasc Med. 2022;9:975478. doi:10.3389/fcvm.2022.975478.

73. Heo JY, Kim JH. Assessing the performance of 28 pathogenicity prediction methods on rare single nucleotide variants in coding regions. BMC Genomics. 2025;26(1):641. doi:10.1186/s12864-025-11787-4.

74. Ravikumar S. The missing "I don't know": why three reasoning-reliability findings converge on calibrated abstention. In: Proceedings of the 15th International Joint Conference on Natural Language Processing and the 5th Conference of the Asia-Pacific Chapter of the Association for Computational Linguistics (Volume 2: Short Papers). Association for Computational Linguistics; 2026. p. 513-524. doi:10.18653/2026.aacl-short.34.

75. Meier J, Rao R, Verkuil R, et al. Language models enable zero-shot prediction of the effects of mutations on protein function. Adv Neural Inf Process Syst. 2021;34:29287-29303.

78. Dalla-Torre H, Gonzalez L, Mendoza-Revilla J, et al. Nucleotide Transformer: building and evaluating robust foundation models for human genomics. Nat Methods. 2025;22(2):287-297. doi:10.1038/s41592-024-02523-z.

85. Warr A, Affara N, Aken B, et al. An improved pig reference genome sequence to enable pig genetics and genomics research. GigaScience. 2020;9(6):giaa051. doi:10.1093/gigascience/giaa051.

88. Cui R, Elzur RA, Kanai M, et al. Improving fine-mapping by modeling infinitesimal effects. Nat Genet. 2024;56(1):162-169. doi:10.1038/s41588-023-01597-3.

95. Zeng T, Li YI. Predicting RNA splicing from DNA sequence using Pangolin. Genome Biol. 2022;23(1):103. doi:10.1186/s13059-022-02664-4.

102. Tavtigian SV, Harrison SM, Boucher KM, et al. Fitting a naturally scaled point system to the ACMG/AMP variant classification guidelines. Hum Mutat. 2020;41(10):1734-1737. doi:10.1002/humu.24088.

107. DeLong ER, DeLong DM, Clarke-Pearson DL. Comparing the areas under two or more correlated receiver operating characteristic curves: a nonparametric approach. Biometrics. 1988;44(3):837-845. doi:10.2307/2531595.

108. Sun X, Xu W. Fast implementation of DeLong's algorithm for comparing the areas under correlated receiver operating characteristic curves. IEEE Signal Process Lett. 2014;21(11):1389-1393. doi:10.1109/LSP.2014.2337313.

109. Morales J, Pujar S, Loveland JE, et al. A joint NCBI and EMBL-EBI transcript set for clinical genomics and research. Nature. 2022;604(7905):310-315. doi:10.1038/s41586-022-04558-8.

110. Karczewski KJ, Francioli LC, Tiao G, et al. The mutational constraint spectrum quantified from variation in 141,456 humans. Nature. 2020;581(7809):434-443. doi:10.1038/s41586-020-2308-7.

115. Liu S, Gao Y, Canela-Xandri O, et al. A multi-tissue atlas of regulatory variants in cattle. Nat Genet. 2022;54(9):1438-1447. doi:10.1038/s41588-022-01153-5.

116. UniProt Consortium. UniProt: the Universal Protein Knowledgebase in 2025. Nucleic Acids Res. 2025;53(D1):D609-D617. doi:10.1093/nar/gkae1010.

117. Pearce MT, Dooms T, Yamamoto R, et al. EVEE: interpretable variant effect prediction from genomic foundation model embeddings. Zenodo. 2026. doi:10.5281/zenodo.19701997.

118. Tabet DR, Coté AG, Lancaster MC, et al. The functional landscape of coding variation in the familial hypercholesterolemia gene LDLR. Science. 2026;391(6787):eady7186. doi:10.1126/science.ady7186.

120. Lin Z, Akin H, Rao R, et al. Evolutionary-scale prediction of atomic-level protein structure with a language model. Science. 2023;379(6637):1123-1130. doi:10.1126/science.ade2574.

123. Stenton SL, Pejaver V, Bergquist T, et al. Assessment of the evidence yield for the calibrated PP3/BP4 computational recommendations. Genet Med. 2024;26(11):101213. doi:10.1016/j.gim.2024.101213.

124. Walsh I, Fishman D, Garcia-Gasulla D, et al. DOME: recommendations for supervised machine learning validation in biology. Nat Methods. 2021;18(10):1122-1127. doi:10.1038/s41592-021-01205-4.

125. Steinhaus R, Proft S, Schuelke M, et al. MutationTaster2021. Nucleic Acids Res. 2021;49(W1):W446-W451. doi:10.1093/nar/gkab266.

126. Gotoh O. An improved algorithm for matching biological sequences. J Mol Biol. 1982;162(3):705-708. doi:10.1016/0022-2836(82)90398-9.

127. Henikoff S, Henikoff JG. Amino acid substitution matrices from protein blocks. Proc Natl Acad Sci U S A. 1992;89(22):10915-10919. doi:10.1073/pnas.89.22.10915.

128. Chou HH, Takematsu H, Diaz S, et al. A mutation in human CMP-sialic acid hydroxylase occurred after the Homo-Pan divergence. Proc Natl Acad Sci U S A. 1998;95(20):11751-11756. doi:10.1073/pnas.95.20.11751.

129. Mantel N, Haenszel W. Statistical aspects of the analysis of data from retrospective studies of disease. J Natl Cancer Inst. 1959;22(4):719-748. doi:10.1093/jnci/22.4.719.

130. Miettinen O, Nurminen M. Comparative analysis of two rates. Stat Med. 1985;4(2):213-226. doi:10.1002/sim.4780040211.

131. Farrington CP, Manning G. Test statistics and sample size formulae for comparative binomial trials with null hypothesis of non-zero risk difference or non-unity relative risk. Stat Med. 1990;9(12):1447-1454. doi:10.1002/sim.4780091208.

132. Wang Z, Zhao G, Li B, et al. Performance comparison of computational methods for the prediction of the function and pathogenicity of non-coding variants. Genomics Proteomics Bioinformatics. 2023;21(3):649-661. doi:10.1016/j.gpb.2022.02.002.

133. Li J, Zhao T, Zhang Y, et al. Performance evaluation of pathogenicity-computation methods for missense variants. Nucleic Acids Res. 2018;46(15):7793-7804. doi:10.1093/nar/gky678.

134. Gu YJ, Wang PY, Fu QQ, et al. Epilepsy-associated CHD2 missense variants and optimization strategies for genetic diagnosis: a comparative analysis of algorithms. Front Neurol. 2025;16:1729387. doi:10.3389/fneur.2025.1729387.

135. Liu X, Gu L, Hao C, et al. Systematic assessment of structural variant annotation tools for genomic interpretation. Life Sci Alliance. 2025;8(3):e202402949. doi:10.26508/lsa.202402949.

136. Montanucci L, Brünger T, Boßelmann CM, et al. Evaluating novel in silico tools for accurate pathogenicity classification in epilepsy-associated genetic missense variants. Epilepsia. 2024;65(12):3655-3663. doi:10.1111/epi.18155.

137. Pshennikova VG, Barashkov NA, Romanov GP, et al. Comparison of predictive in silico tools on missense variants in GJB2, GJB6, and GJB3 genes associated with autosomal recessive deafness 1A (DFNB1A). ScientificWorldJournal. 2019;2019:5198931. doi:10.1155/2019/5198931.

138. Michels M, Matte U, Fraga LR, et al. Determining the pathogenicity of CFTR missense variants: multiple comparisons of in silico predictors and variant annotation databases. Genet Mol Biol. 2019;42(3):560-570. doi:10.1590/1678-4685-GMB-2018-0148.

139. Montenegro LR, Lerário AM, Nishi MY, et al. Performance of mutation pathogenicity prediction tools on missense variants associated with 46,XY differences of sex development. Clinics (Sao Paulo). 2021;76:e2052. doi:10.6061/clinics/2021/e2052.

140. Cannon S, Williams M, Gunning AC, et al. Evaluation of in silico pathogenicity prediction tools for the classification of small in-frame indels. BMC Med Genomics. 2023;16(1):36. doi:10.1186/s12920-023-01454-6.

141. Barbosa P, Savisaar R, Carmo-Fonseca M, et al. Computational prediction of human deep intronic variation. GigaScience. 2023;12:giad085. doi:10.1093/gigascience/giad085.

142. Cubuk C, Garrett A, Choi S, et al. Clinical likelihood ratios and balanced accuracy for 44 in silico tools against multiple large-scale functional assays of cancer susceptibility genes. Genet Med. 2021;23(11):2096-2104. doi:10.1038/s41436-021-01265-z.

143. Strauch Y, Lord J, Niranjan M, et al. CI-SpliceAI—improving machine learning predictions of disease causing splicing variants using curated alternative splice sites. PLoS One. 2022;17(6):e0269159. doi:10.1371/journal.pone.0269159.

144. Malhis N, Jacobson M, Jones SJM, et al. LIST-S2: taxonomy based sorting of deleterious missense mutations across species. Nucleic Acids Res. 2020;48(W1):W154-W161. doi:10.1093/nar/gkaa288.

145. Riepe TV, Khan M, Roosing S, et al. Benchmarking deep learning splice prediction tools using functional splice assays. Hum Mutat. 2021;42(7):799-810. doi:10.1002/humu.24212.

146. Leman R, Parfait B, Vidaud D, et al. SPiP: Splicing Prediction Pipeline, a machine learning tool for massive detection of exonic and intronic variant effects on mRNA splicing. Hum Mutat. 2022;43(12):2308-2323. doi:10.1002/humu.24491.

147. Gunning AC, Fryer V, Fasham J, et al. Assessing performance of pathogenicity predictors using clinically relevant variant datasets. J Med Genet. 2021;58(8):547-555. doi:10.1136/jmedgenet-2020-107003.

148. Rowlands C, Thomas HB, Lord J, et al. Comparison of in silico strategies to prioritize rare genomic variants impacting RNA splicing for the diagnosis of genomic disorders. Sci Rep. 2021;11(1):20607. doi:10.1038/s41598-021-99747-2.

149. Lord J, Oquendo CJ, Wai HA, et al. Predicting the impact of rare variants on RNA splicing in CAGI6. Hum Genet. 2025;144(2-3):243-251. doi:10.1007/s00439-023-02624-3.

150. Rentzsch P, Schubach M, Shendure J, et al. CADD-Splice—improving genome-wide variant effect prediction using deep learning-derived splice scores. Genome Med. 2021;13(1):31. doi:10.1186/s13073-021-00835-9.

151. Caron B, Luo Y, Rausell A. NCBoost classifies pathogenic non-coding variants in Mendelian diseases through supervised learning on purifying selection signals in humans. Genome Biol. 2019;20(1):32. doi:10.1186/s13059-019-1634-2.

152. Zadrozny B, Elkan C. Transforming classifier scores into accurate multiclass probability estimates. In: Proceedings of the Eighth ACM SIGKDD International Conference on Knowledge Discovery and Data Mining (KDD '02). New York: ACM; 2002. p. 694-699. doi:10.1145/775047.775151.

153. Platt JC. Probabilistic outputs for support vector machines and comparisons to regularized likelihood methods. In: Smola AJ, Bartlett P, Schölkopf B, et al., editors. Advances in Large Margin Classifiers. Cambridge (MA): MIT Press; 1999. p. 61-74.

154. Kull M, Silva Filho TM, Flach P. Beta calibration: a well-founded and easily implemented improvement on logistic calibration for binary classifiers. In: Proceedings of the 20th International Conference on Artificial Intelligence and Statistics (AISTATS). Proc Mach Learn Res. 2017;54:623-631.

155. Saerens M, Latinne P, Decaestecker C. Adjusting the outputs of a classifier to new a priori probabilities: a simple procedure. Neural Comput. 2002;14(1):21-41. doi:10.1162/089976602753284446.

156. Elkan C. The foundations of cost-sensitive learning. In: Proceedings of the Seventeenth International Joint Conference on Artificial Intelligence (IJCAI'01). San Francisco (CA): Morgan Kaufmann; 2001. p. 973-978.

157. Vovk V, Gammerman A, Shafer G. Algorithmic learning in a random world. New York: Springer; 2005. doi:10.1007/b106715.

158. Angelopoulos AN, Bates S. Conformal prediction: a gentle introduction. Found Trends Mach Learn. 2023;16(4):494-591. doi:10.1561/2200000101.

159. Barber RF, Candès EJ, Ramdas A, et al. Conformal prediction beyond exchangeability. Ann Stat. 2023;51(2):816-845. doi:10.1214/23-AOS2276.

160. Schiff Y, Kao CH, Gokaslan A, et al. Caduceus: bi-directional equivariant long-range DNA sequence modeling. In: Proceedings of the 41st International Conference on Machine Learning (ICML). Proc Mach Learn Res. 2024;235:43632-43648.

161. Rosen BD, Bickhart DM, Schnabel RD, et al. De novo assembly of the cattle reference genome with single-molecule sequencing. GigaScience. 2020;9(3):giaa021. doi:10.1093/gigascience/giaa021.

162. Davenport KM, Bickhart DM, Worley K, et al. An improved ovine reference genome assembly to facilitate in-depth functional annotation of the sheep genome. GigaScience. 2022;11:giab096. doi:10.1093/gigascience/giab096.

163. Bickhart DM, Rosen BD, Koren S, et al. Single-molecule sequencing and chromatin conformation capture enable de novo reference assembly of the domestic goat genome. Nat Genet. 2017;49(4):643-650. doi:10.1038/ng.3802.

164. Kalbfleisch TS, Rice ES, DePriest MS, et al. Improved reference genome for the domestic horse increases assembly contiguity and composition. Commun Biol. 2018;1:197. doi:10.1038/s42003-018-0199-z.

165. International Chicken Genome Sequencing Consortium. Sequence and comparative analysis of the chicken genome provide unique perspectives on vertebrate evolution. Nature. 2004;432(7018):695-716. doi:10.1038/nature03154.

166. Lindblad-Toh K, Wade CM, Mikkelsen TS, et al. Genome sequence, comparative analysis and haplotype structure of the domestic dog. Nature. 2005;438(7069):803-819. doi:10.1038/nature04338.

167. Hoeppner MP, Lundquist A, Pirun M, et al. An improved canine genome and a comprehensive catalogue of coding genes and non-coding transcripts. PLoS One. 2014;9(3):e91172. doi:10.1371/journal.pone.0091172.

168. Bredemeyer KR, Hillier L, Harris AJ, et al. Single-haplotype comparative genomics provides insights into lineage-specific structural variation during cat evolution. Nat Genet. 2023;55(11):1953-1963. doi:10.1038/s41588-023-01548-y.

169. Keightley PD, Eyre-Walker A. Joint inference of the distribution of fitness effects of deleterious mutations and population demography based on nucleotide polymorphism frequencies. Genetics. 2007;177(4):2251-2261. doi:10.1534/genetics.107.080663.

170. Eyre-Walker A, Keightley PD. The distribution of fitness effects of new mutations. Nat Rev Genet. 2007;8(8):610-618. doi:10.1038/nrg2146.

171. Lek M, Karczewski KJ, Minikel EV, et al.; Exome Aggregation Consortium. Analysis of protein-coding genetic variation in 60,706 humans. Nature. 2016;536(7616):285-291. doi:10.1038/nature19057.

172. Lindblad-Toh K, Garber M, Zuk O, et al. A high-resolution map of human evolutionary constraint using 29 mammals. Nature. 2011;478(7370):476-482. doi:10.1038/nature10530.

173. Vaser R, Adusumalli S, Leng SN, et al. SIFT missense predictions for genomes. Nat Protoc. 2016;11(1):1-9. doi:10.1038/nprot.2015.123.

174. Reva B, Antipin Y, Sander C. Predicting the functional impact of protein mutations: application to cancer genomics. Nucleic Acids Res. 2011;39(17):e118. doi:10.1093/nar/gkr407.

175. Choi Y, Sims GE, Murphy S, et al. Predicting the functional effect of amino acid substitutions and indels. PLoS One. 2012;7(10):e46688. doi:10.1371/journal.pone.0046688.

176. Carter H, Douville C, Stenson PD, et al. Identifying Mendelian disease genes with the Variant Effect Scoring Tool. BMC Genomics. 2013;14(Suppl 3):S3. doi:10.1186/1471-2164-14-S3-S3.

177. Dong C, Wei P, Jian X, et al. Comparison and integration of deleteriousness prediction methods for nonsynonymous SNVs in whole exome sequencing studies. Hum Mol Genet. 2015;24(8):2125-2137. doi:10.1093/hmg/ddu733.

178. Li C, Zhi D, Wang K, et al. MetaRNN: differentiating rare pathogenic and rare benign missense SNVs and InDels using deep learning. Genome Med. 2022;14(1):115. doi:10.1186/s13073-022-01120-z.

179. Jagadeesh KA, Wenger AM, Berger MJ, et al. M-CAP eliminates a majority of variants of uncertain significance in clinical exomes at high sensitivity. Nat Genet. 2016;48(12):1581-1586. doi:10.1038/ng.3703.

180. Pejaver V, Urresti J, Lugo-Martinez J, et al. Inferring the molecular and phenotypic impact of amino acid variants with MutPred2. Nat Commun. 2020;11(1):5918. doi:10.1038/s41467-020-19669-x.

181. Qi H, Zhang H, Zhao Y, et al. MVP predicts the pathogenicity of missense variants by deep learning. Nat Commun. 2021;12(1):510. doi:10.1038/s41467-020-20847-0.

182. Zhang H, Xu MS, Fan X, et al. Predicting functional effect of missense variants using graph attention neural networks. Nat Mach Intell. 2022;4(11):1017-1028. doi:10.1038/s42256-022-00561-w.

183. Zhao Y, Lan T, Zhong G, et al. A probabilistic graphical model for estimating selection coefficients of nonsynonymous variants from human population sequence data. Nat Commun. 2025;16(1):4670. doi:10.1038/s41467-025-59937-2.

184. Samocha KE, Kosmicki JA, Karczewski KJ, et al. Regional missense constraint improves variant deleteriousness prediction. bioRxiv. 2017. doi:10.1101/148353.

185. Sundaram L, Gao H, Padigepati SR, et al. Predicting the clinical impact of human mutation with deep neural networks. Nat Genet. 2018;50(8):1161-1170. doi:10.1038/s41588-018-0167-z.

186. Raimondi D, Tanyalcin I, Ferté J, et al. DEOGEN2: prediction and interactive visualization of single amino acid variant deleteriousness in human proteins. Nucleic Acids Res. 2017;45(W1):W201-W206. doi:10.1093/nar/gkx390.

187. Feng BJ. PERCH: a unified framework for disease gene prioritization. Hum Mutat. 2017;38(3):243-251. doi:10.1002/humu.23158.

188. Alirezaie N, Kernohan KD, Hartley T, et al. ClinPred: prediction tool to identify disease-relevant nonsynonymous single-nucleotide variants. Am J Hum Genet. 2018;103(4):474-483. doi:10.1016/j.ajhg.2018.08.005.

189. Wu Y, Liu H, Li R, et al. Improved pathogenicity prediction for rare human missense variants. Am J Hum Genet. 2021;108(10):1891-1906. doi:10.1016/j.ajhg.2021.08.012.

190. Dereli O, Kuru N, Akkoyun E, et al. PHACTboost: a phylogeny-aware pathogenicity predictor for missense mutations via boosting. Mol Biol Evol. 2024;41(7):msae136. doi:10.1093/molbev/msae136.

191. Jiang TT, Fang L, Wang K. Deciphering "the language of nature": a transformer-based language model for deleterious mutations in proteins. Innovation (Camb). 2023;4(5):100487. doi:10.1016/j.xinn.2023.100487.

192. Quinodoz M, Peter VG, Cisarova K, et al. Analysis of missense variants in the human genome reveals widespread gene-specific clustering and improves prediction of pathogenicity. Am J Hum Genet. 2022;109(3):457-470. doi:10.1016/j.ajhg.2022.01.006.

193. Orenbuch R, Shearer CA, Kollasch AW, et al. Proteome-wide model for human disease genetics. Nat Genet. 2025;57(12):3165-3174. doi:10.1038/s41588-025-02400-1.

194. Quang D, Chen Y, Xie X. DANN: a deep learning approach for annotating the pathogenicity of genetic variants. Bioinformatics. 2015;31(5):761-763. doi:10.1093/bioinformatics/btu703.

195. Rogers MF, Shihab HA, Mort M, et al. FATHMM-XF: accurate prediction of pathogenic point mutations via extended features. Bioinformatics. 2018;34(3):511-513. doi:10.1093/bioinformatics/btx536.

196. Ionita-Laza I, McCallum K, Xu B, et al. A spectral approach integrating functional genomic annotations for coding and noncoding variants. Nat Genet. 2016;48(2):214-220. doi:10.1038/ng.3477.

197. McVicker G, Gordon D, Davis C, et al. Widespread genomic signatures of natural selection in hominid evolution. PLoS Genet. 2009;5(5):e1000471. doi:10.1371/journal.pgen.1000471.

198. Munté E, Muiños F, Marín R, et al. Evaluation of BoostDM, a somatic variant prediction tool, for the interpretation of germline variants in hereditary cancer genes. Eur J Hum Genet. 2026;34(9):1336-1341. doi:10.1038/s41431-026-02077-y.

199. Nelson N, Niaz A, Fairfax K, et al. The performance of in silico prediction tools for variant curation in a panel of cancer genes. Hum Mutat. 2026;2026:8817550. doi:10.1155/humu/8817550.

200. Brock DC, Wang M, Hussain HMJ, et al. Comparative analysis of in-silico tools in identifying pathogenic variants in dominant inherited retinal diseases. Hum Mol Genet. 2024;33(11):945-957. doi:10.1093/hmg/ddae028.

201. Ding X, Singh P, Schimenti K, et al. In vivo versus in silico assessment of potentially pathogenic missense variants in human reproductive genes. Proc Natl Acad Sci U S A. 2023;120(30):e2219925120. doi:10.1073/pnas.2219925120.

202. Holland KD, Bouley TM, Horn PS. Comparison and optimization of in silico algorithms for predicting the pathogenicity of sodium channel variants in epilepsy. Epilepsia. 2017;58(7):1190-1198. doi:10.1111/epi.13798.
