## Supplementary figures and images for "Unscored variants leave many comparisons of variant effect predictors undecided: evidence from ClinVar, published benchmarks and genomic language models"

### banner.png

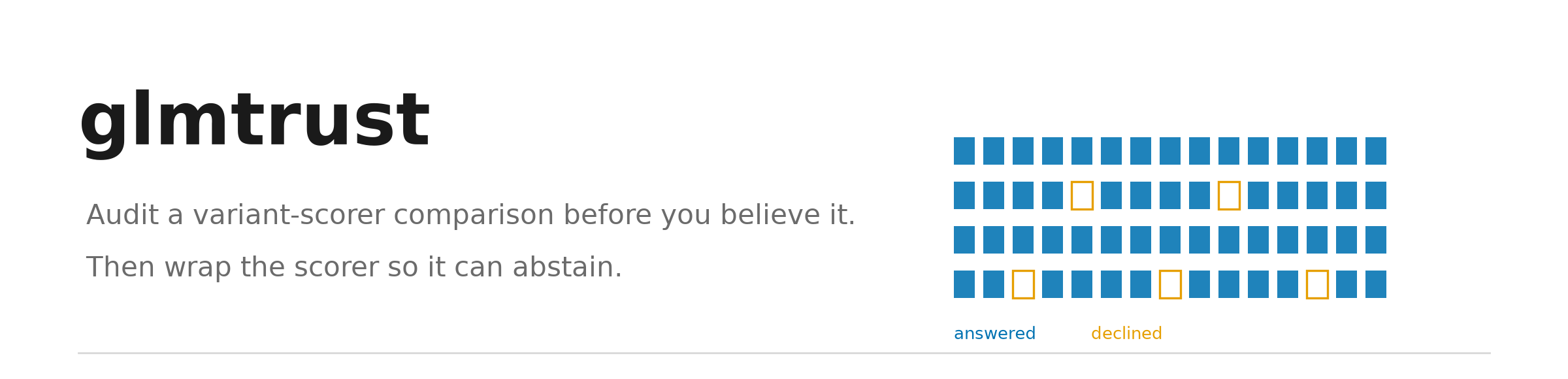

### brca1_brct_evo2.png

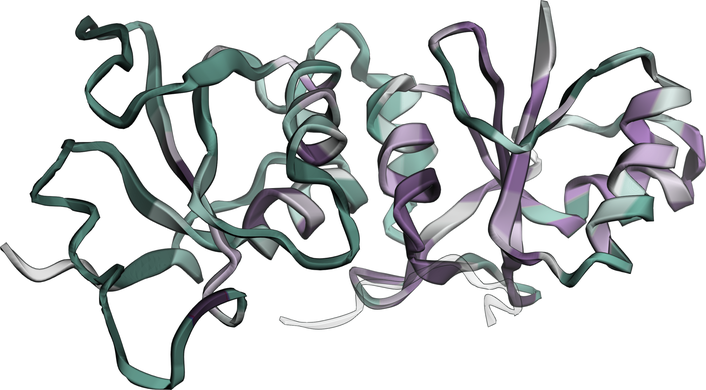

### brca1_brct_sge.png

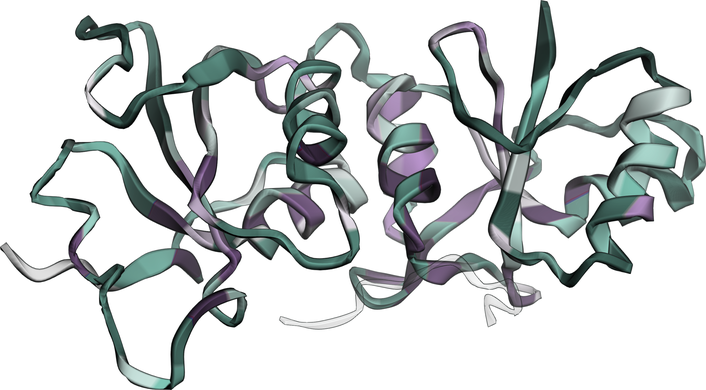

### brca1_ring_evo2.png

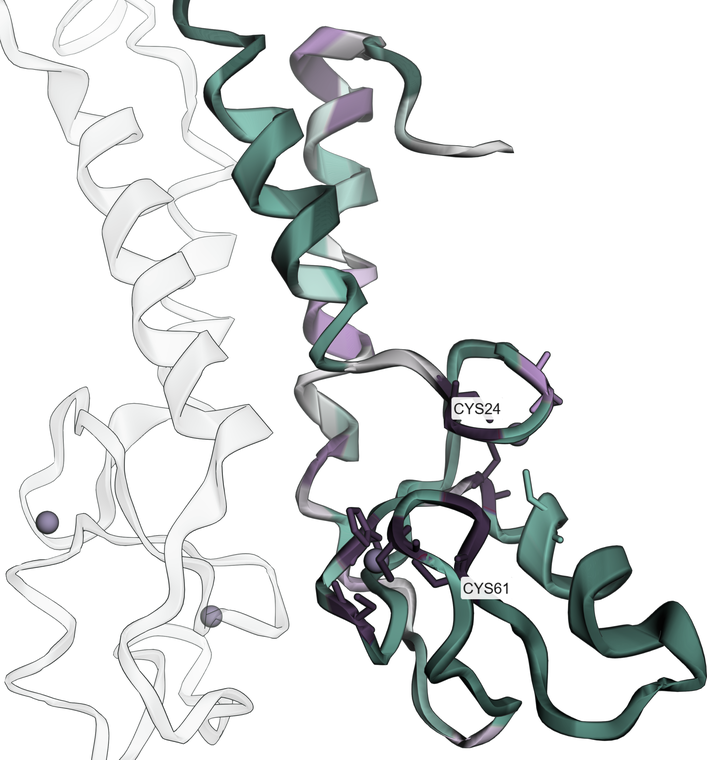

### brca1_ring_sge.png

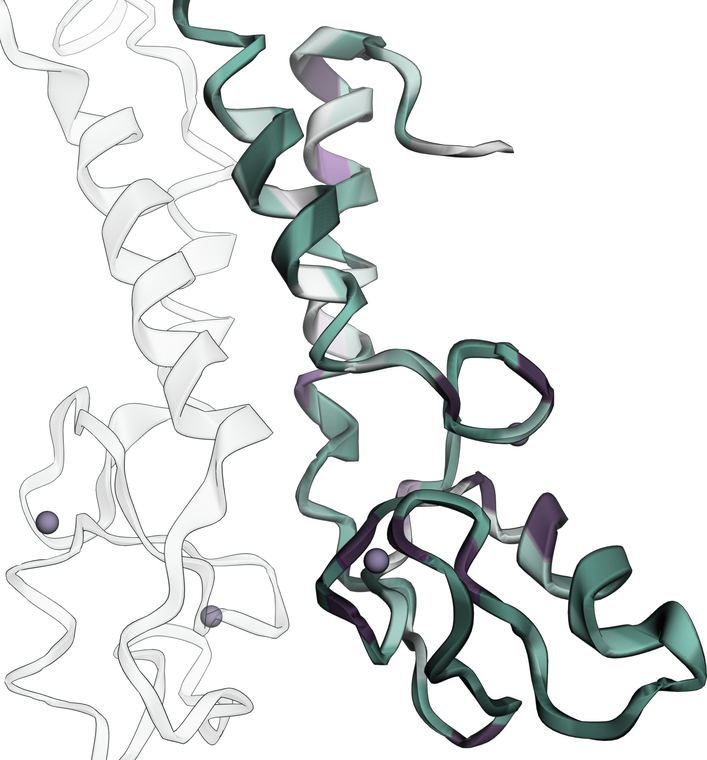

### cat.png

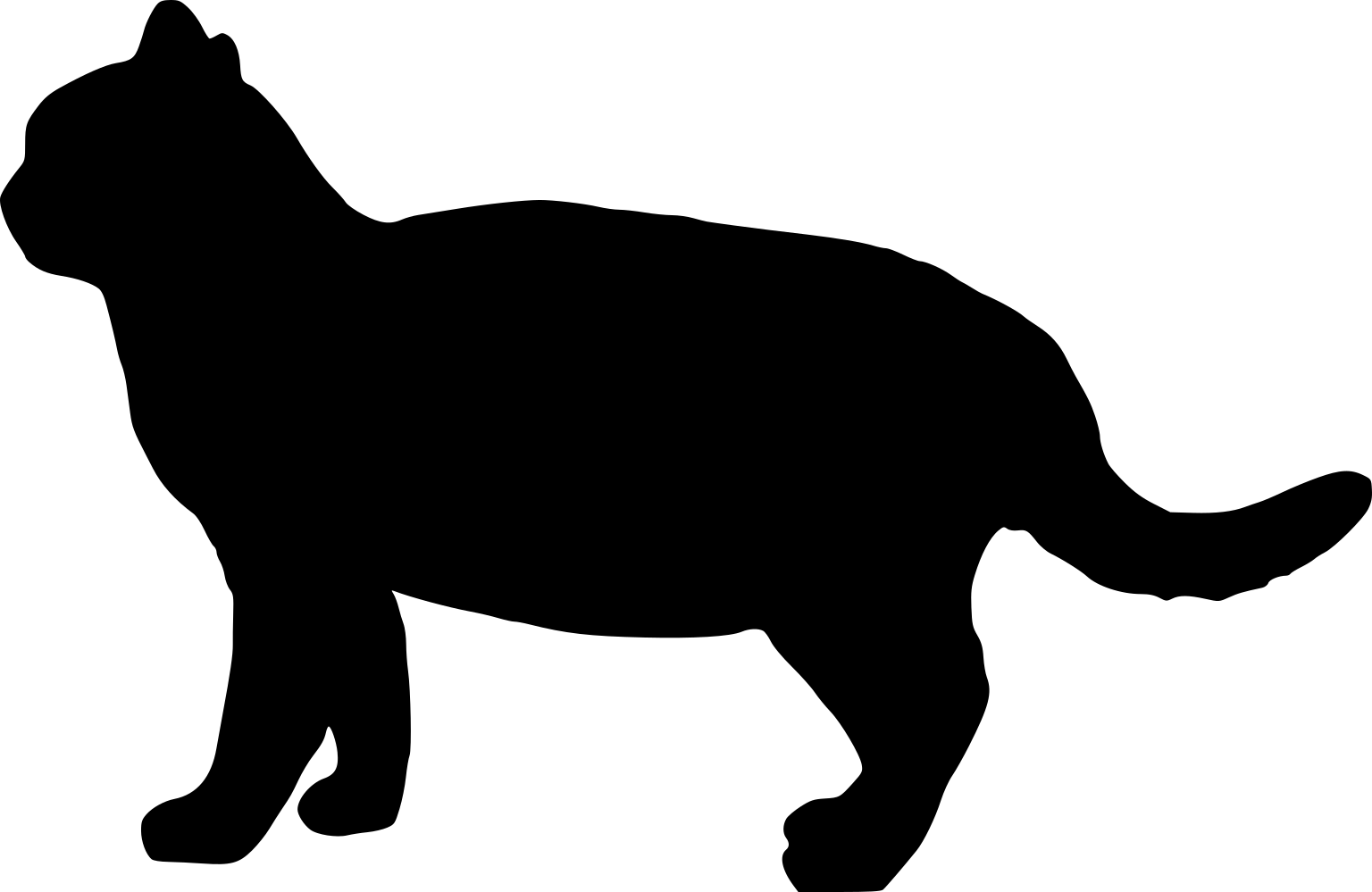

### cattle.png

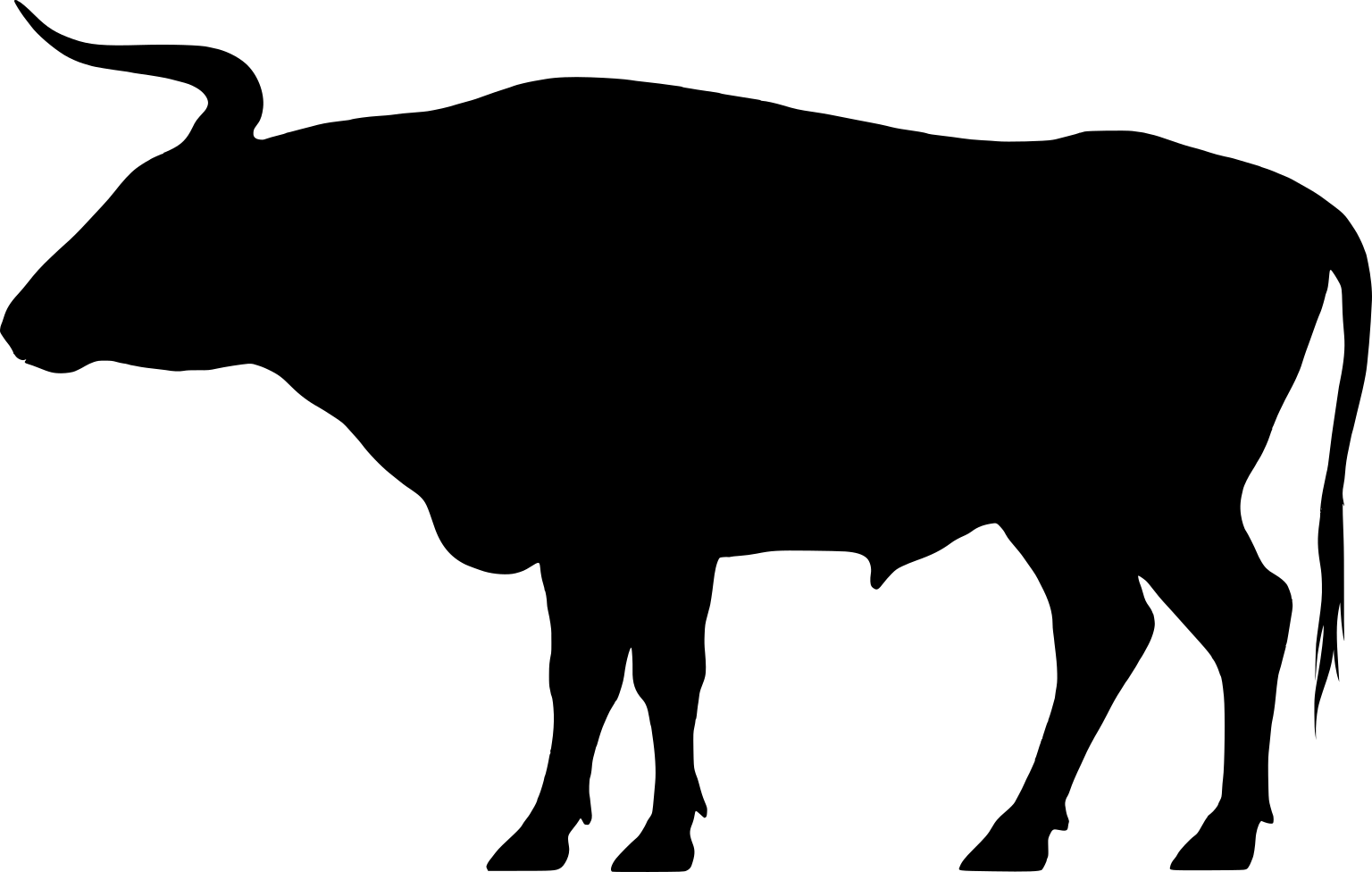

### chicken.png

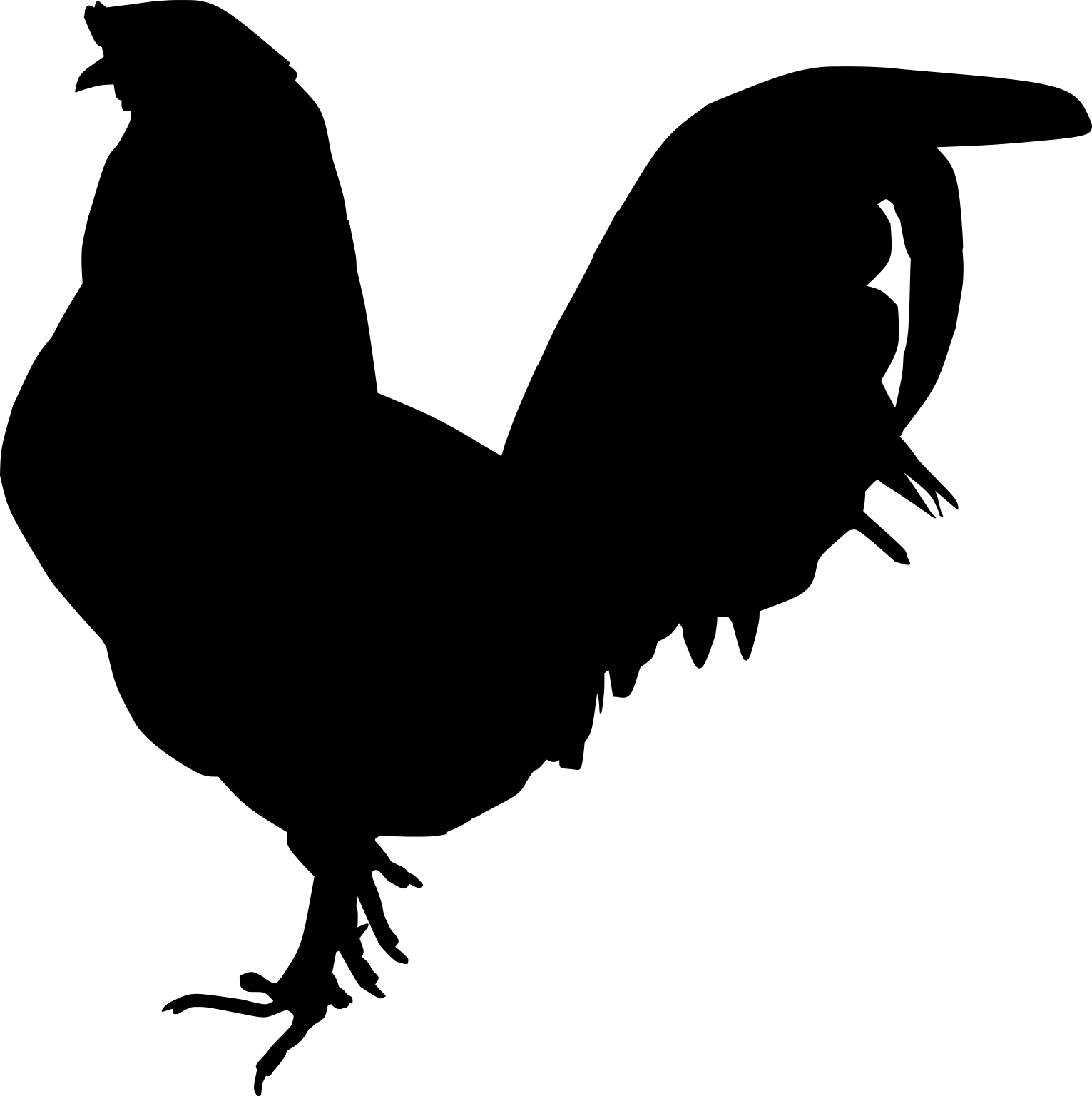

### concept.png

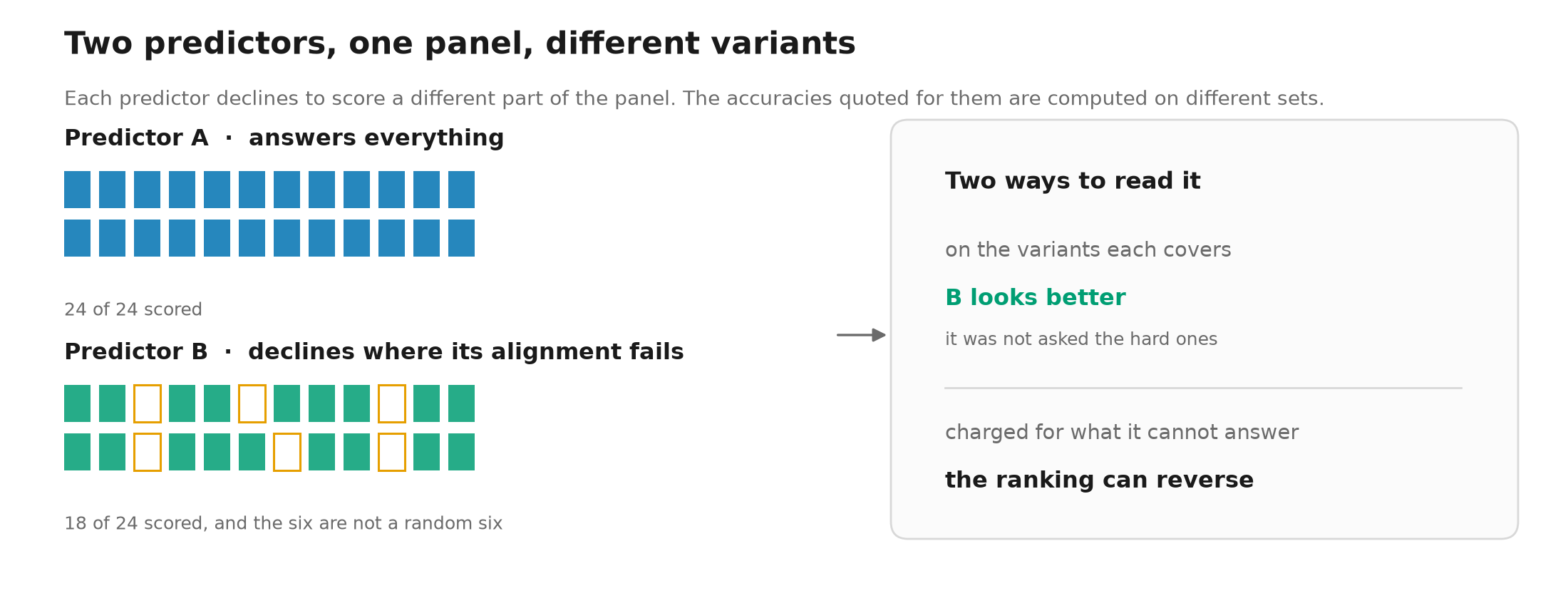

### dog.png

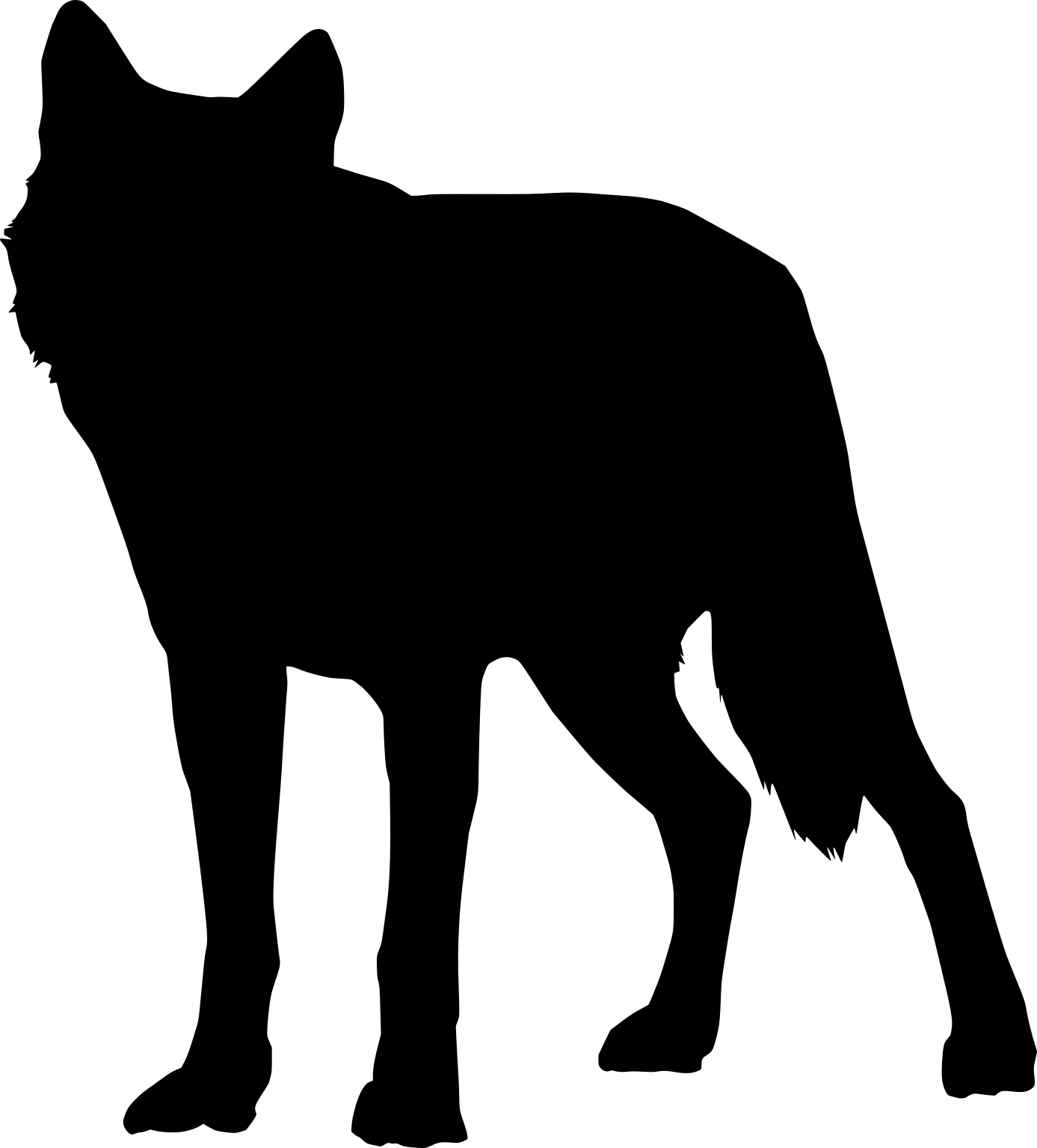

### goat.png

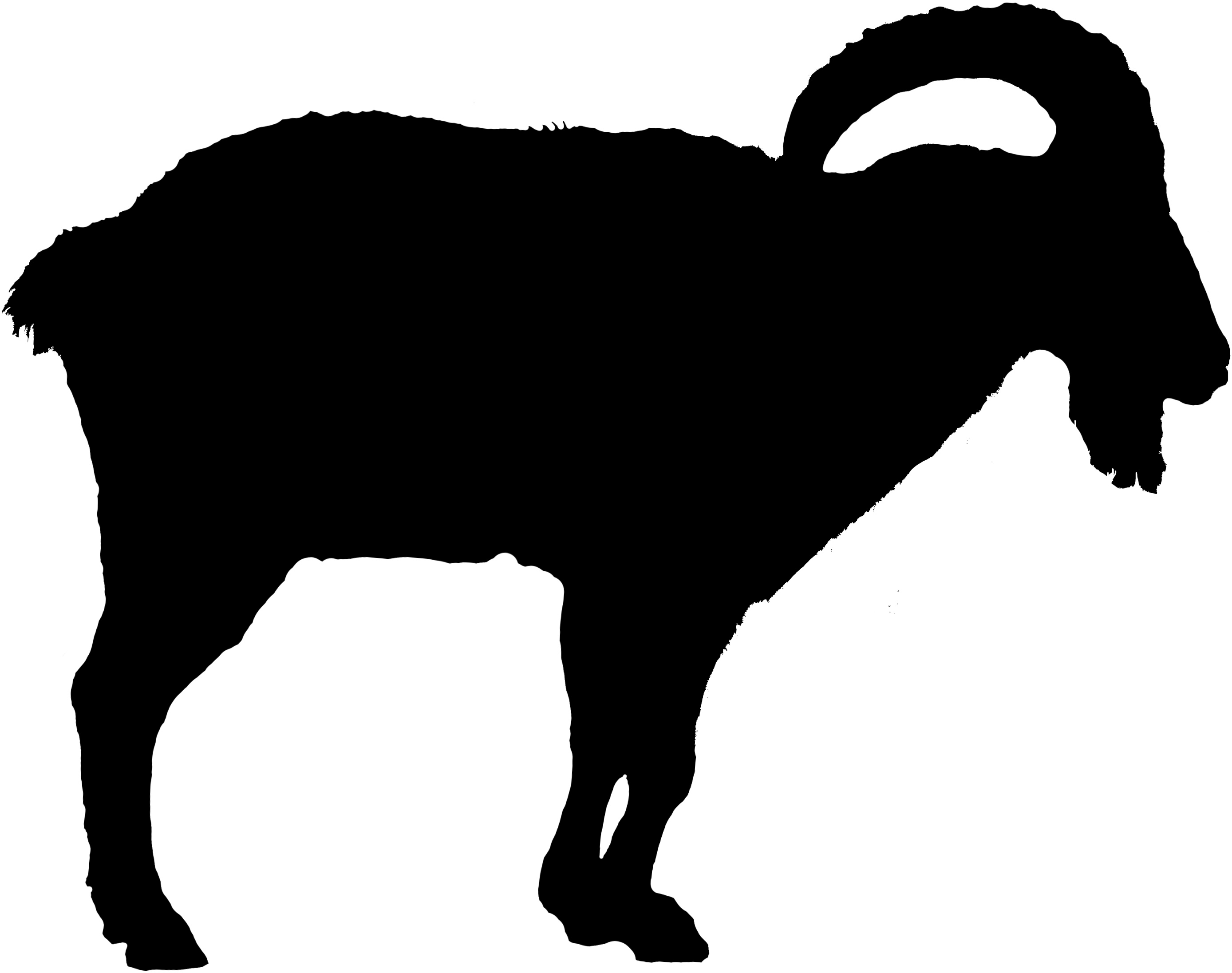

### horse.png

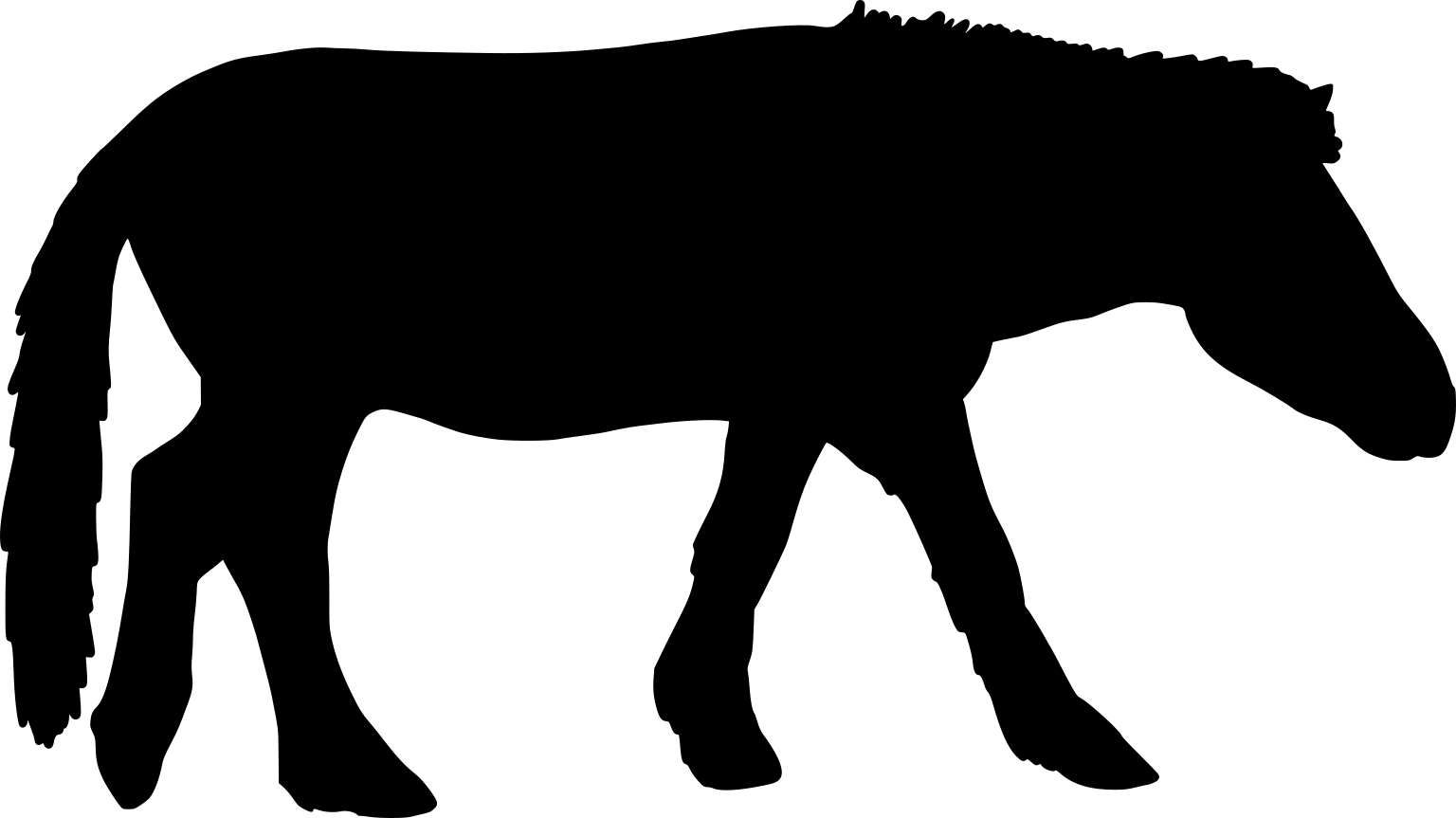

### human.png

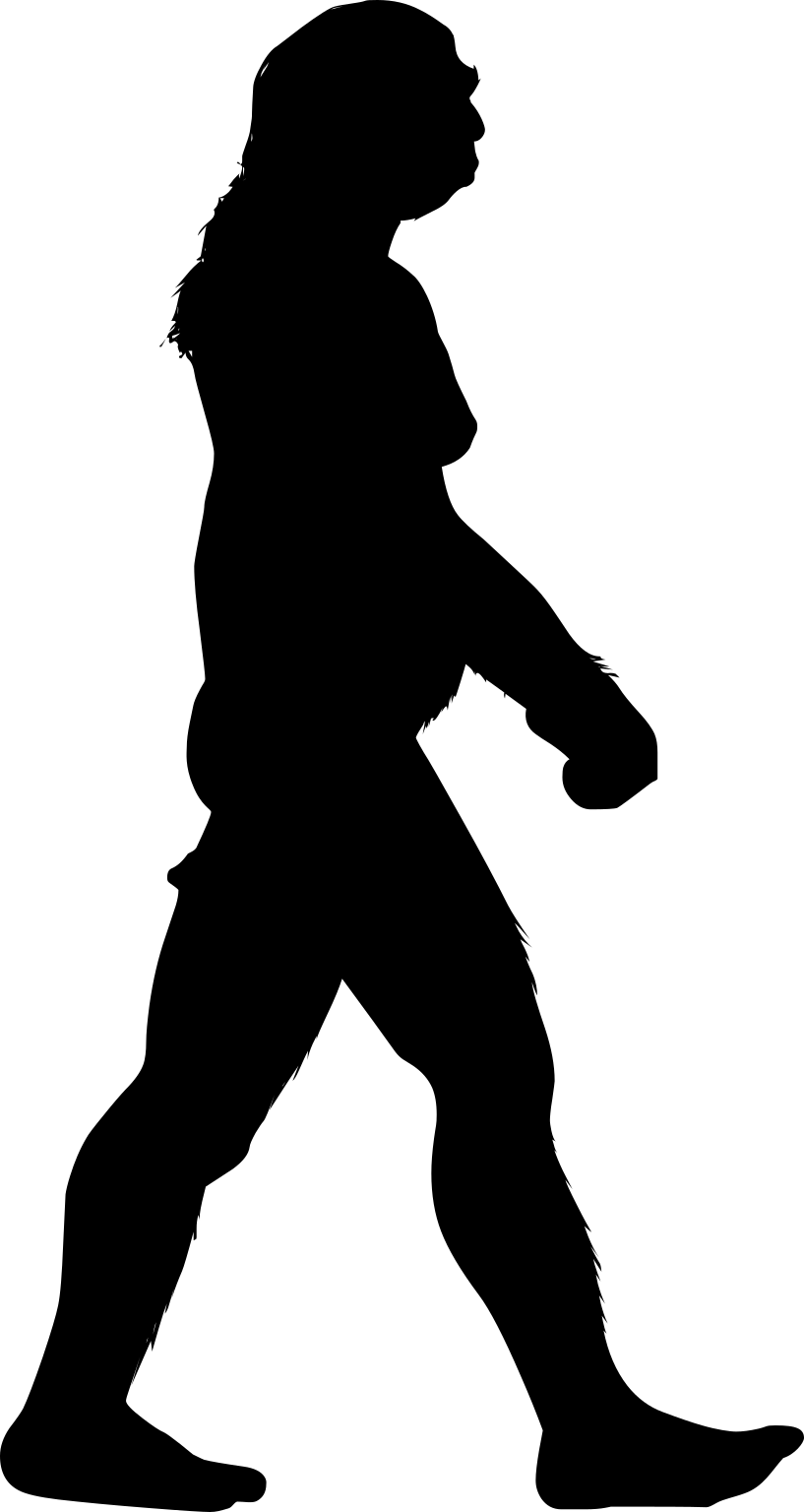

### pig.png

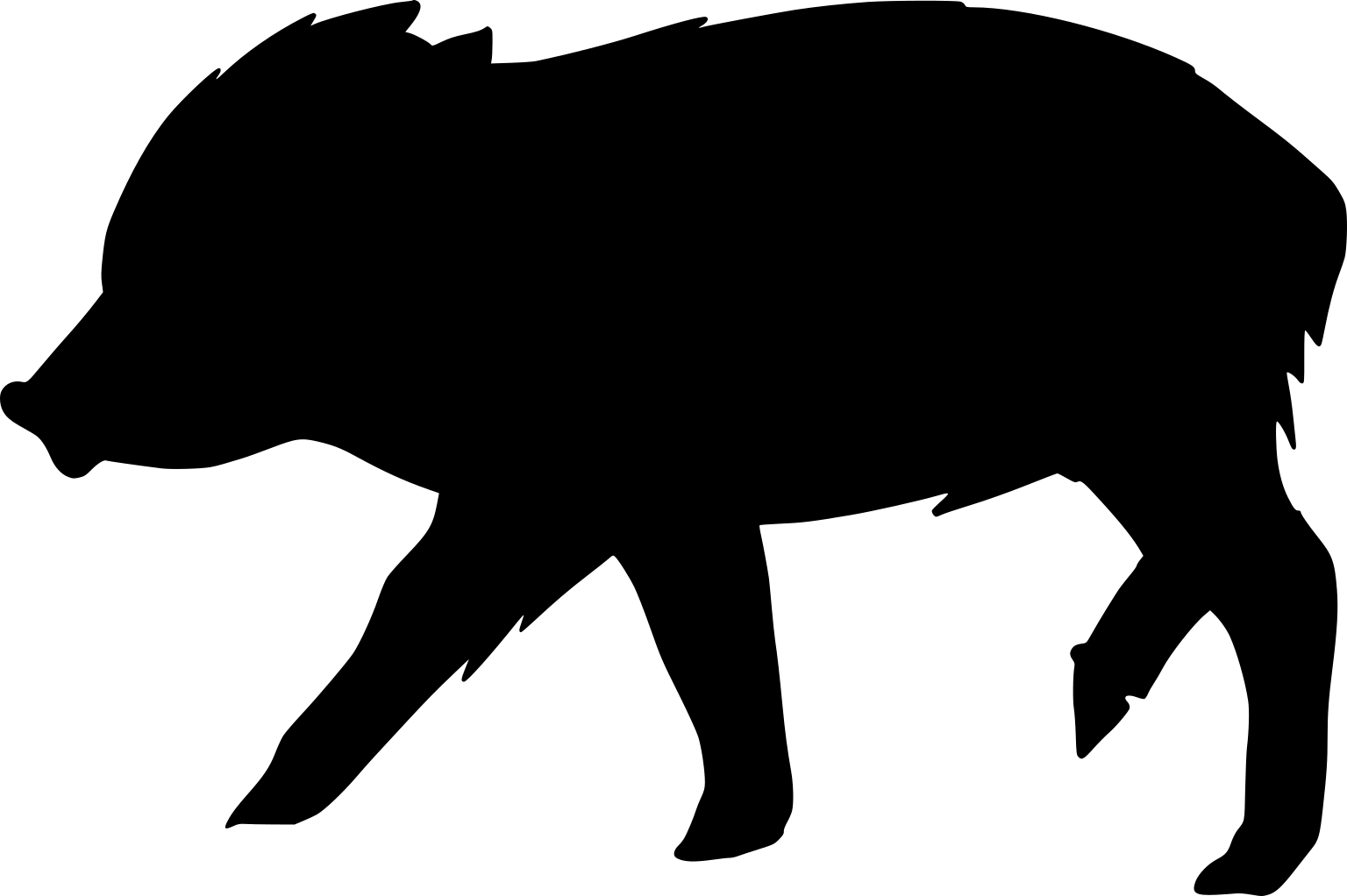

### sheep.png

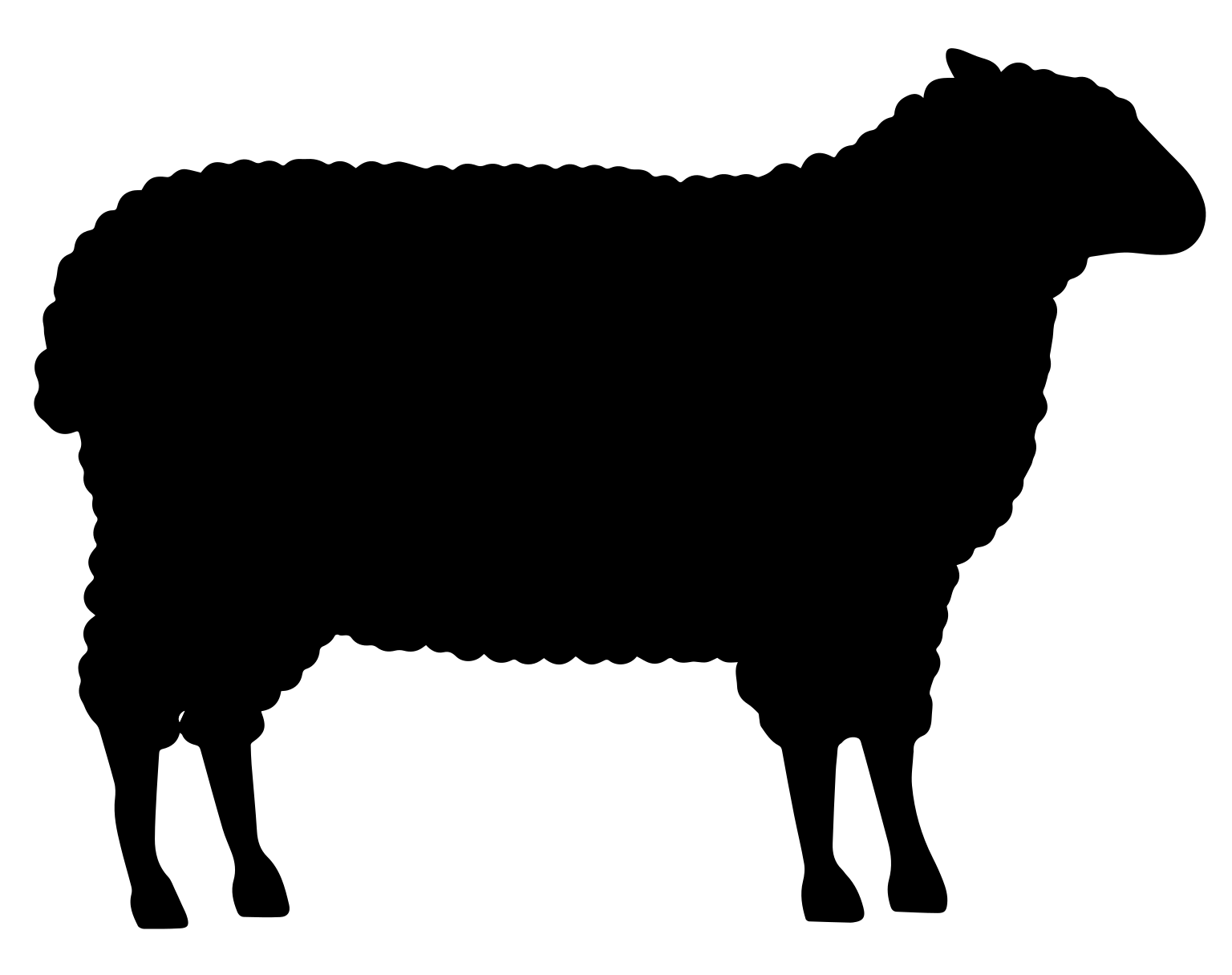
